# Associations between the environment, brain, mental health, and cognition across adolescence

**DOI:** 10.64898/2026.04.27.721046

**Authors:** Yumnah T. Khan, Jakob Seidlitz, Lena Dorfschmidt, Alex Tsompanidis, Carrie Allison, Lifespan Brain Chart Consortium, Ran Barzilay, Aaron Alexander-Bloch, Simon Baron-Cohen, Sarah-Jayne Blakemore, Richard A. I. Bethlehem

## Abstract

Adolescence is a critical period for the development of the brain, cognition, and mental health, which are shaped by a wide range of environmental factors. In the present study, we analysed the Adolescent Brain and Cognitive Development (ABCD) dataset to examine how a range of proximal (e.g., socioeconomic status, familial circumstances) and distal (e.g., neighbourhood conditions, access to healthcare and education) environmental factors are associated with changes in centile-based measures of brain structure, and whether these brain differences subsequently mediate variations in cognition and mental health. We analysed these associations both at baseline (N = 6,911; 3,605 M, 3,606 F; mean age = 9.93) and longitudinally across three timepoints (N = 1,628; 879 M, 749 F; ages 8-15). At baseline, a more advantaged proximal and distal environment was associated with larger volumes across the whole brain relative to age- and sex-matched peers, which, in turn, mediated better mental health outcomes and cognitive performance. In the longitudinal analysis, the childhood environment predicted changes in brain structure across adolescence, and these structural changes predicted changes in mental health and cognition. The childhood environment also predicted cognitive but not mental health changes across adolescence, suggesting that these associations may already be established early in adolescence. These findings provide insight into how environmental and neural factors shape adolescent mental health and cognition, with potential implications for early intervention strategies aimed at promoting positive developmental outcomes.

## Introduction

Adolescence, the period of transition between childhood and adulthood, is widely recognised as a formative period of development. The brain undergoes significant structural and functional reorganisation^1^, which, in turn, is thought to underpin the substantial cognitive development and changes in mental health that occur during this period^2^.While these associations have long been hypothesised, there is surprisingly limited longitudinal research assessing these links across adolescence. Moreover, both brain and behavioural development are shaped by a plethora of environmental factors. Therefore, incorporating the role of the environment is critical for understanding individual differences in adolescent cognitive and mental health trajectories.

It has been suggested that a reorganisation of the brain occurs during adolescence^1^, triggered by a surge of synaptic elimination^3^, axonal growth, and myelination^4^. At a macro-structural level, white matter volumes show linear increases^5^, while gray matter volumes follow a non-linear inverted U-shaped trajectory; initial increases in early childhood, followed by a peak in late childhood and subsequent decreases throughout adolescence^6–8^. Regional gray matter volumes mirror known patterns of behavioural development and generally follow a posterior-to-anterior gradient, where sensory regions mature earlier than higher-order associative regions^5,9^.

It is also during adolescence that the majority of mental health conditions are first diagnosed, including depression, anxiety, eating disorders, and schizophrenia^10,11^. Simultaneously, substantial cognitive development also occurs across various domains such as executive functioning, response inhibition, attention control, language, planning and organisation, emotion regulation, and social cognition^12^. It is hypothesised that these changes in mental health and cognition are underpinned by the structural changes in the brain occurring during adolescence. Consistent with this hypothesis, lower cortical^13^, subcortical^14^, and global gray matter volumes^15^ have been associated with higher scores on measures of psychopathology in adolescence. In other cases, region-specific effects have been observed for mental health^16^ and cognitive outcomes^17,18^, and these effects also appear to vary depending on the developmental stage under investigation. More recent frameworks have also proposed that relative *deviations*^19^ in brain structure from population norms may better explain individual differences in mental health and cognitive outcomes than raw brain volumes, though further large-scale research is required to better characterise these associations in adolescence.

Importantly, measures of brain structure and behaviour during adolescence have both been associated with environmental factors. These can broadly be categorised into distal or macro-environmental factors (e.g., neighbourhood socioeconomic conditions, access to education, access to healthcare) and proximal or micro-environmental factors (e.g., household income, childhood adversity, parenting factors). Given that several of these factors are often tightly inter-related^20^ (e.g., neighbourhood conditions often covary with socioeconomic status), it is prudent to investigate their collective impact on adolescent outcomes. Generally, higher psychopathological traits during adolescence have been associated with adverse environmental conditions, such as low socioeconomic status, harsh or inconsistent parenting, family conflict, and exposure to community violence and disadvantage^21–24^. On the other hand, higher cognitive outcomes have been associated with better environmental conditions, such as having a higher socioeconomic status, reduced exposure to air pollution, and favourable perinatal conditions (e.g., higher birth weight)^25–27^. Environmental factors have also been associated with individual differences in brain structure, such that improved macro-and micro-environmental conditions are generally associated with larger brain volumes^28,29^.

Increasing evidence has suggested that the effects of environmental exposures on mental health and cognition are mediated through influences on the brain detectable via MRI. For instance, family income has been shown to influence neurocognitive functioning through its effects on brain structure, particularly within frontal and temporal regions^25,30^. Similarly, recent work using data from the Adolescent Brain Cognitive Development (ABCD) study demonstrated that childhood family and neighbourhood environments are associated with differences in brain structure during early adolescence, which also mediate the association between environments and psychological outcomes such as externalising traits (i.e., presenting internal distress via outwardly directed behaviours such as aggression) ^14^. Thus, it is possible that environmental factors may be associated with behaviour and cognition partly via their effects on brain structure, and variations in this pathway may partially contribute to individual differences in these outcomes.

While this pathway has previously been investigated, several gaps remain. First, most studies rely on cross-sectional data, which may miss meaningful individual differences in developmental trajectories (i.e., whether brain, cognitive, or behavioural development is accelerating, decelerating, or remaining stable)^31^. Given that adolescence is marked by substantial brain and behavioural development, it may be that these *trajectories* of change are more predictive of individual differences in outcomes^32^. Second, relatively few studies have examined how change with age in one domain (e.g., brain structure) predicts concurrent change in another (e.g., behaviour) during critical periods of development, such as adolescence^33^. Investigating these linked temporal trajectories may be more informative in understanding the temporal timeline of these relationships and improving causal inferences. Third, traditional analyses often rely on raw volumetric measures or absolute changes in brain volume. On the other hand, centile-based^8,34^ approaches indicate where an individual’s brain volume falls relative to same-age and same-sex peers. These centile scores can then be used to show whether an individual’s development remains stable, accelerates, or falls behind over time relative to the population reference. These *relative* changes might be more sensitive compared to absolute changes in explaining individual differences in mental health and cognition.

In the present study, in order to understand the factors associated with changes in mental health and cognition across adolescence, we aimed to investigate the environment-brain-behaviour pathway using centile-based measures of brain volumes. The ABCD Study consists of a large, longitudinal sample with environmental measures, MRI scans, cognitive tests, and mental health trait measures^35^. Here, we examined how the late childhood macro- and micro-environment predict changes in population-referenced brain volumes and, in turn, changes in mental health and cognition across adolescence. To capture baseline effects as well as longitudinal changes over time, we investigated the brain-behaviour-environment mediation pathway in both the baseline pre-adolescent cohort (ages 8-11) as well as a longitudinal cohort subset with data over 3 timepoints across adolescence (ages 8-15).

## RESULTS

A brief overview of the methods is presented in Figure 1, which is described further below and in the Methods section.

**Figure 1.**
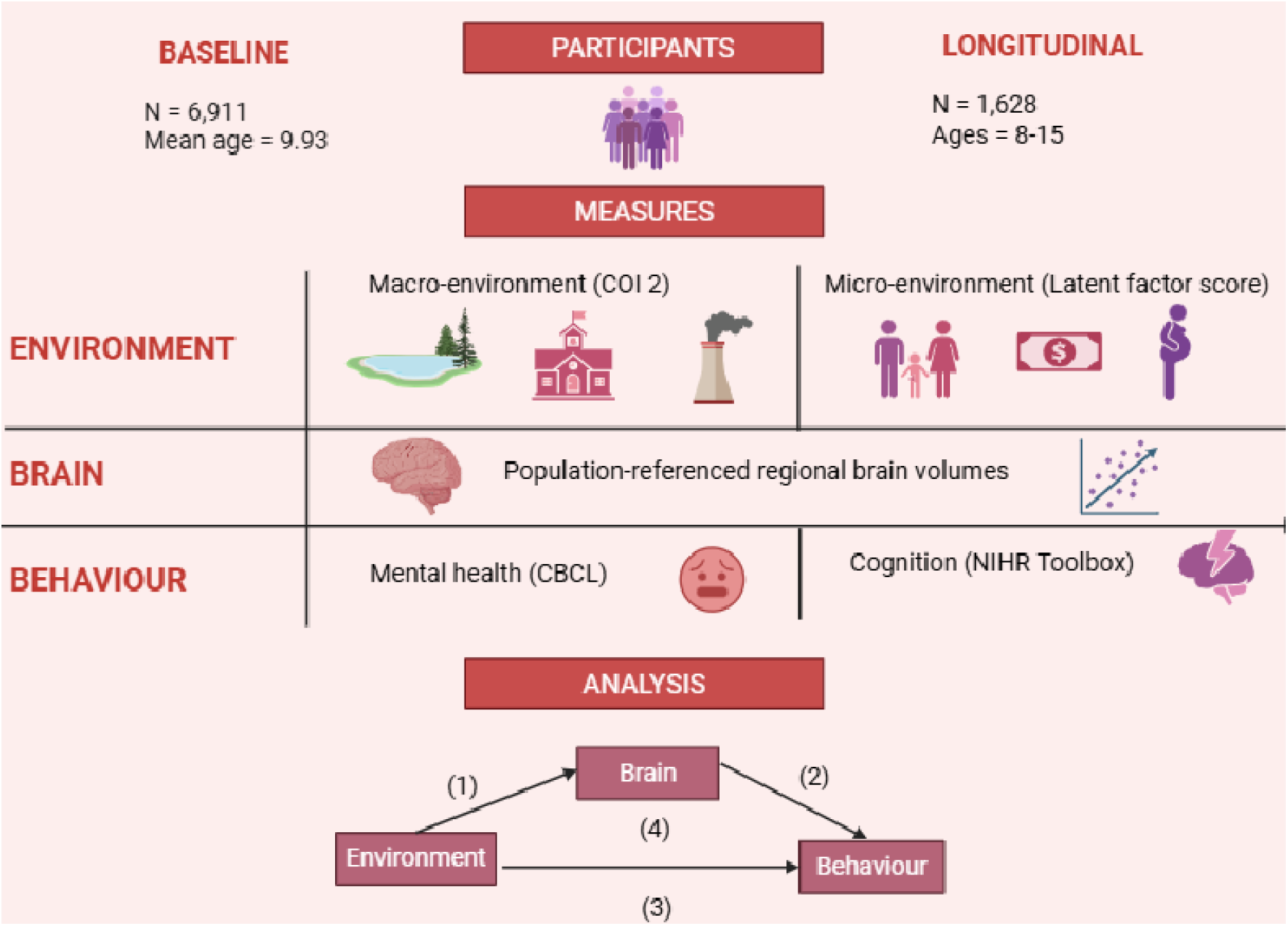
Brief overview of methodology. *Note*. Brief overview of methodology, including participants in the baseline and longitudinal cohort. The macro-environment was measured via the Child Opportunity Index (COI 2.0), which captures a broad range of neighbourhood-level environmental indicators across the social and economic, health and environment, and educational domains. The micro-environment was summarised into a single summary latent factor score, which aggregates a range of proximal factors such economic security, parental characteristics, school/community environment, and adverse childhood experiences. For population-referenced measures of grey matter volume, centile scores were calculated for each individual, which reflect the percentile each individual’s brain volumes fall in relation to age and sex-matched peers. Mental health traits were measured via the Childhood Behaviour Checklist (CBCL), grouped into internalising (e.g., sadness, depression, anxiety, and loneliness) and externalising (e.g., aggression and rule-breaking) traits. Cognition was measured via the NIH Toolbox Cognition Battery (NIH Toolbox®), which consists of various sub-tasks assessing cognitive domains such as executive function, memory, processing speed, vocabulary, and reading.

### Baseline analysis

The ABCD baseline sample used in this study consisted of 6,911 pre-adolescent participants (3,606 female, 3,605 male) aged 8-11 years (Mean = 9.93, SD = 0.63) with complete data from all measurements of interests. For each individual, grey matter volume centile scores were computed for each region, which reflect the percentile each individual’s brain volumes fall in relation to age and sex-matched peers. These scores were computed via a maximum-likelihood estimator and using previously derived population norms and out sample-estimation approach^8^. Each time point was treated as an independent study, and random effects of site were calculated at each time independently, which has been shown to account for site-related variation^8^.

Mixed-effects models were used to examine the relationships between environmental factors, regional brain centile scores, and mental health and cognitive outcomes. Age and sex were modelled as covariates, while testing site and family ID (to account for sibling pairs) were modelled as random intercepts. To provide a standardised measure of effect size, standardised beta coefficients were computed via z-scoring the outcome and predictor of interest. Significant associations were followed up with mediation analyses. False Discovery Rate (FDR) corrections were applied within analyses categories (e.g., for each environmental predictor or mental health/cognitive task) to control for multiple comparisons across the whole brain (see further details in the “Methods” section).

### Environment-brain associations

The macro-environment was measured at baseline via the Child Opportunity Index (COI 2.0)^36,37^, which aggregates a broad range of macro-environmental indicators into three summary domains: Social and Economic (e.g., employment rate, poverty rate, homeownership rate, median household income), Education (e.g., school poverty, high school graduation rate, college enrolment, ECE centres), and Health and Environment (e.g., access to healthy food, access to green space, industrial pollutants, health insurance coverage). The micro-environment was measured via a single summary latent factor score, which aggregates a range of proximal factors across the following domains: economic security, parental characteristics, school/community environment, adverse childhood experiences, physiological health, and perinatal wellbeing (e.g., gestational age at birth, birth weight)^38^.

First, to assess associations between the environment and the brain, we assessed whether micro- and macro-environment scores were associated with centile scores at baseline. Consistent with hypotheses, pre-adolescents with more favourable micro- and macro environments showed larger brain volumes relative to their peers (Supplementary Tables 1-4). These effects were observed across the entire brain, though associations were particularly pronounced in multisensory and sensorimotor regions such as the middle temporal gyrus, lateral occipital gyrus, inferior temporal gyrus, precentral gyrus, and post-central gyrus (Figure 2). Similar patterns were observed across the COI 2.0 subdomains and the micro-environment latent factor scores.

**Figure 2.**
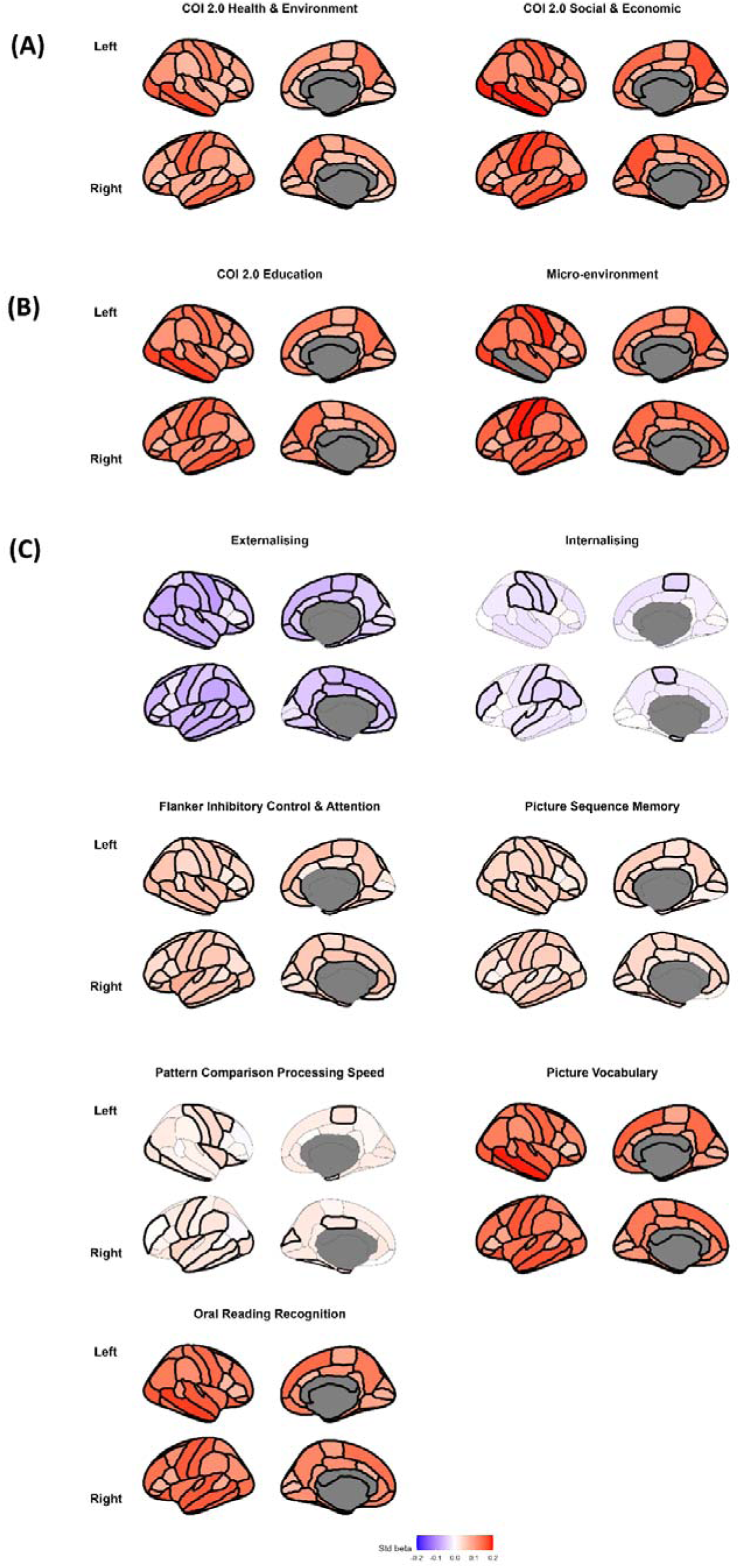
Baseline Associations. *Note.* Brain plots depicting regional standardised beta coefficients for associations between (A) environmental factors and centile scores, (B) centile scores and mental health traits, and (C) centile scores and change scores on cognitive tasks. Regions are colour-coded according to the magnitude and direction of the standardised beta values. Statistically significant regions (p_FDR_ < 0.05) are outlined. Details on the measures can be found in the Methods section.

### Brain-behaviour associations

Next, we examined whether brain centile scores are associated with mental health and cognitive outcomes. Mental health traits were measured via the Childhood Behaviour Checklist (CBCL)^39^, which is grouped into two higher-order summary factors measuring internalising (e.g., sadness, depression, anxiety, and loneliness) and externalising (e.g., aggression and rule-breaking) traits. As predicted, adolescents with lower brain volumes relative to their peers showed greater internalising and externalising traits (Supplementary Tables 4-5). Associations with externalising were particularly pronounced in the frontal pole, temporal pole, transverse temporal gyrus, and pars triangularis – structures that are broadly implicated in higher-order executive functioning and sensory, socio-emotional, and semantic processing. Associations with internalising were particularly pronounced in the supramarginal gyrus, superior temporal gyrus, rostral middle frontal gyrus, and postcentral gyrus – structures that are broadly implicated in higher-order sensory integration, cognition, and motor control (Figure 2).

Cognition was measured via the NIH Toolbox Cognition Battery (NIH Toolbox®)^40^, which includes the following tasks: (a) Flanker Inhibitory Control & Attention: measures attention, cognitive control, executive function, and inhibition of automatic response, (b) Picture Sequence Memory: measures episodic memory and sequencing, (c) Pattern Comparison Processing Speed: measures information processing and processing speed, (d) Picture Vocabulary: measures language vocabulary knowledge, and (e) Oral Reading Recognition: measures language, oral reading skills, and academic achievement. As hypothesised, adolescents with larger volumes across the brain relative to their peers showed higher scores across all cognitive measures (Figure 2).

### Environment-behaviour

Next, associations between environmental factors and cognitive and mental health outcomes were assessed. As previously evidenced, less advantaged macro- and micro-environmental conditions were generally associated with higher internalising and externalising traits and lower scores across all cognitive measures (Table 1).

**Table 1:**
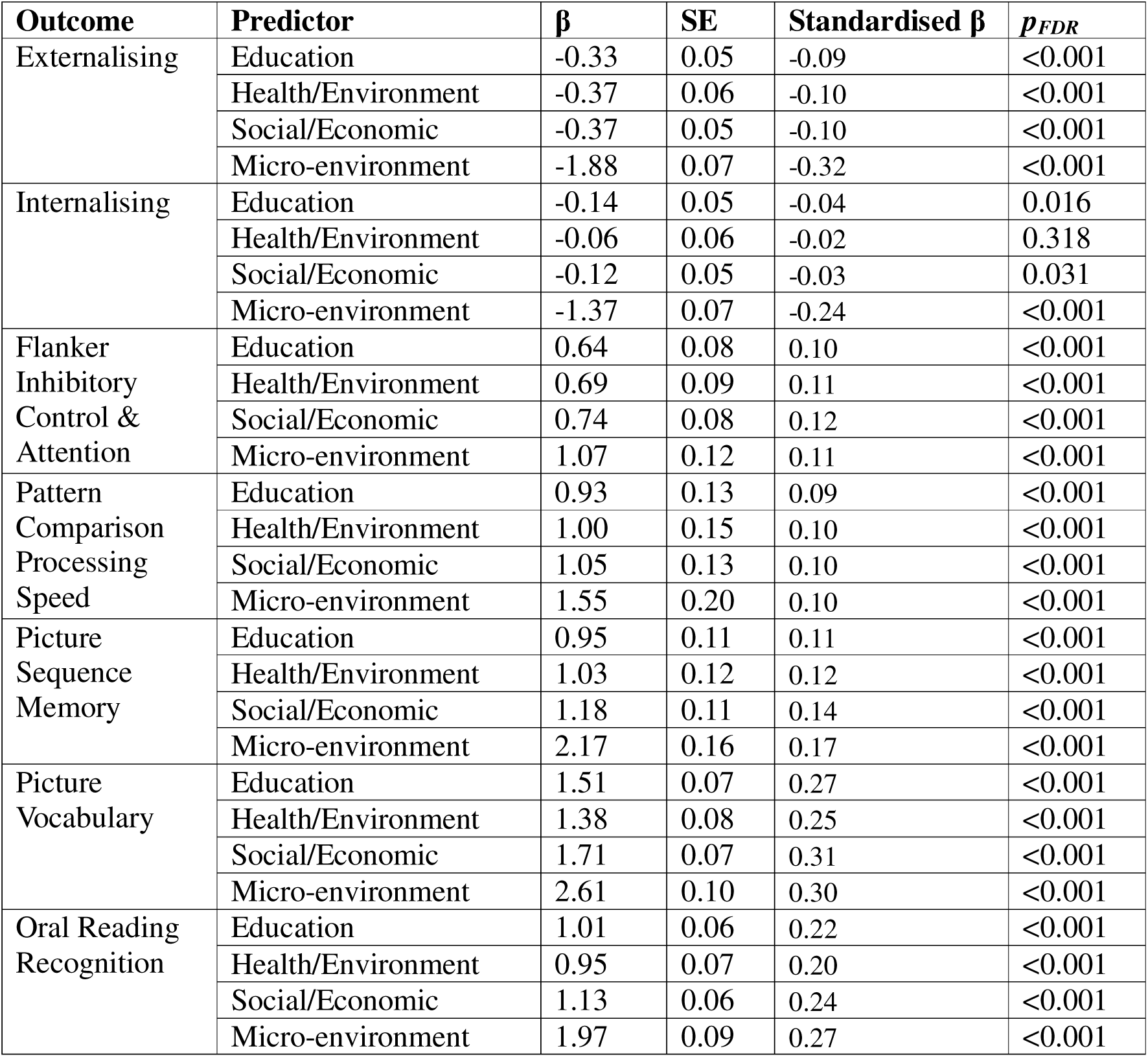
Environment-behaviour associations at baseline. Shown are unstandardised regression coefficients (β), standard errors (SE), standardised coefficients (standardised β), and FDR-corrected *p* values (*p*_FDR_) for all combinations of environment-behaviour associations at baseline.

### Environment-brain-behaviour mediations

Since significant environment-brain, brain-behaviour, and environment-behaviour associations were observed, we next examined whether the associations between the environment and behaviour are partially mediated via the brain. The environment-brain-behaviour mediation pathway was tested for combinations showing significant associations in the preceding analyses (Supplementary Tables 12-39) by pre-selecting the regions that showed associations with both the environment as well as mental health and cognitive outcomes. Results indicated that regional centile scores mediated the associations between (a) all macro-environmental variables and externalising (Supplementary Tables 12-15), (b) the social/economic and educational macro-environment and internalising (Supplementary Tables 17 & 18), (c) all macro-environmental variables and scores on the Flanker Inhibitory Control and Attention Task (Supplementary Tables 20-22), and (d) all macro- and micro-environmental variables and scores on the Oral Reading Recognition Task (Supplementary Tables 36-39). Therefore, at baseline, brain centile scores partially mediate the association between environmental factors and behavioural outcomes.

### Longitudinal analysis

The longitudinal cohort consisted of a subset of participants from the baseline cohort that had complete data across the three timepoints for all measurements of interest, resulting in 1,628 (749 males, 879 females) participants aged 8-15. Demographic characteristics of this longitudinal subset were largely similar to the baseline cohort (see Table 3 in Methods).

Analyses followed the same structure as the baseline analysisis, except that age-adjusted change scores were used for brain and behavioural measures by averaging the differences in scores between consecutive timepoints (see “Methods” section for further details). All models controlled for sex and, where relevant, baseline mental health and cognitive scores.

### Environment-brain associations

The baseline environment, measured at the first timepoint during late childhood, was tested as a predictor for changes in centile scores across adolescence. Across various regions, adolescents with a better childhood macro- and micro-environment showed upward shifts in centile scores (Supplementary Tables 40-44), moving higher up in their centile rankings across adolescence. In other words, the brain volumes of these adolescents increased faster or decreased slower than what was expected based on their initial rankings. Associations were particularly pronounced in higher-order association and multimodal integration regions, such as the precuneus, insula, pericalcarine gyrus, lingual gyrus, inferior parietal gyrus, and middle temporal gyrus (Figure 3). Associations were also observed amongst regions of the associative network, visual network, and default mode network.

**Figure 3.**
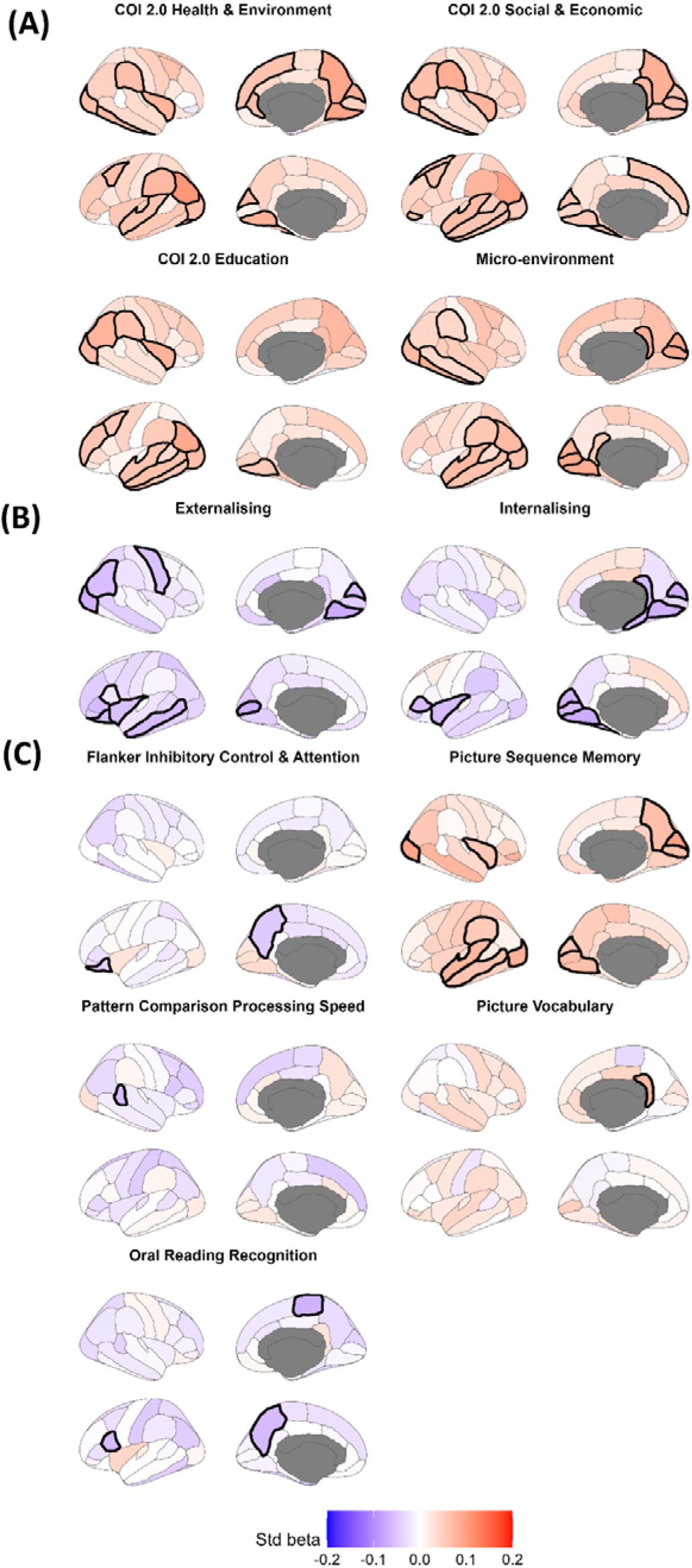
Longitudinal Associations. *Note*. Brain plots depicting regional standardised beta coefficients for longitudinal associations between (A) environmental factors and changes in centile scores, (B) centile change scores and changes in mental health, and (C) centile change scores and changes in cognition. Regions are colour-coded according to the magnitude and direction of the standardised beta values. Statistically significant regions (p_FDR_ < 0.05) are outlined.

### Brain-behaviour associations

Longitudinally, adolescents who showed downward shifts in centile scores (i.e., decreases in their population-referenced brain volume percentiles across adolescence) show increased traits of poor mental health (Supplementary Tables 44 & 45). Internalising and externalising showed both overlapping as well as distinct neural correlates. For instance, both were associated with downward shifts in regions such as the left insula, right cuneus, left pars triangularis, right lingual, and left pericalcarine gyrus, which have been implicated in processes such as self-awareness, emotion regulation, inhibitory control, and visual and semantic processing. Externalising-specific associations were observed in the left lateral orbitofrontal cortex, right lateral occipital cortex, left middle temporal gyrus, right inferior parietal lobule, and right precentral gyrus. Internalising-specific associations, on the other hand, were observed in the left fusiform gyrus, right parahippocampal gyrus, and right isthmus cingulate, previously implicated in visual recognition, memory, and self-referential processing.

With respect to cognition, increases in scores on the Flanker Inhibitory Control and Attention Task (Supplementary Table 46) were associated with upward centile shifts in the left lateral orbitofrontal cortex and left precuneus, previously implicated in inhibitory control, self-referential-processing, and attention shifting. Improvements on the Picture Sequence Memory task were associated with upward centile shifts across the brain (Supplementary Table 47), while improvements on the Oral Reading Recognition task were associated with downward shifts in language regions such as the left pars opercularis, as well as the left precuneus and right paracentral gyrus (Supplementary Table 48). Improvements on the Pattern Comparison Processing Speed task were associated with downward centile shifts in the right superior temporal sulcus, implicated in various broad aspects of cognition (Supplementary Table 49), while increases on the Picture Vocabulary Task were associated with upward shifts in the right isthmus cingulate cortex, previously linked with aspects of memory (Supplementary Table 50).

### Environment – behaviour

The childhood environment was tested as a predictor of changes in mental health and cognition across adolescence. A better macro- and micro-environment was associated with increases in scores across all of the measured cognitive tasks. There were, however, no associations between environmental factors and changes in internalising and externalising (Table 2).

**Table 2:** Longitudinal environment-behaviour associations.

| Outcome | Predictor | $\beta$ | SE | Standardised $\beta$ | $p_{\text{FDR}}$ |
| --- | --- | --- | --- | --- | --- |
| Externalising | Education | -0.01 | 0.02 | -0.01 | 0.707 |
|  | Health/Environment | 0.00 | 0.02 | -0.00 | 0.894 |
|  | Social/Economic | -0.00 | 0.02 | -0.00 | 0.929 |
|  | Micro-environment | -0.05 | 0.03 | -0.04 | 0.145 |
| Internalising | Education | -0.02 | 0.02 | -0.02 | 0.623 |
|  | Health/Environment | -0.01 | 0.03 | -0.01 | 0.707 |
|  | Social/Economic | 0.02 | 0.02 | 0.02 | 0.550 |
|  | Micro-environment | -0.02 | 0.04 | -0.01 | 0.670 |
| Flanker Inhibitory Control & Attention | Education | 0.21 | 0.03 | 0.13 | <0.001 |
|  | Health/Environment | 0.18 | 0.03 | 0.10 | <0.001 |
|  | Social/Economic | 0.22 | 0.03 | 0.13 | <0.001 |
|  | Micro-environment | 0.42 | 0.05 | 0.16 | <0.001 |
| Pattern Comparison Processing Speed | Education | 0.25 | 0.07 | 0.08 | <0.001 |
|  | Health/Environment | 0.22 | 0.08 | 0.07 | 0.008 |
|  | Social/Economic | 0.27 | 0.07 | 0.09 | <0.001 |
|  | Micro-environment | 0.29 | 0.11 | 0.06 | 0.017 |
| Picture Sequence Memory | Education | 0.19 | 0.06 | 0.07 | 0.008 |
|  | Health/Environment | 0.34 | 0.07 | 0.12 | <0.001 |
|  | Social/Economic | 0.27 | 0.07 | 0.09 | <0.001 |
|  | Micro-environment | 0.61 | 0.10 | 0.13 | <0.001 |
| Picture Vocabulary | Education | 0.20 | 0.03 | 0.17 | <0.001 |
|  | Health/Environment | 0.15 | 0.03 | 0.13 | <0.001 |
|  | Social/Economic | 0.19 | 0.03 | 0.16 | <0.001 |
|  | Micro-environment | 0.35 | 0.05 | 0.19 | <0.001 |
| Oral Reading Recognition | Education | 0.05 | 0.02 | 0.05 | 0.063 |
|  | Health/Environment | 0.02 | 0.03 | 0.02 | 0.670 |
|  | Social/Economic | 0.05 | 0.03 | 0.05 | 0.069 |
|  | Micro-environment | 0.18 | 0.04 | 0.12 | <0.001 |

*Note.* Shown are unstandardised regression coefficients ( $\beta$ ), standard errors (SE), standardised coefficients (standardised $\beta$ ), and FDR-corrected $p$ values ( $p_{\text{FDR}}$ ) for all combinations of environment-behaviour associations at baseline.

### Mediation

Mediation analyses were conducted to explore whether the environment-brain-behaviour pathway might explain longitudinal changes in mental health and cognition across adolescence. Outcomes showing significant associations with environmental factors (Table 2) were further tested for the environment-brain-behaviour mediation pathway, and only regions that showed associations with both the environment as well as cognitive outcomes were pre-selected. As a result, mediations were tested for the Picture Sequence Memory Task and the Picture Vocabulary Task. Various significant effects were identified for the Picture Sequence Memory Task. For instance, centile scores in regions such as the cuneus and lateral occipital gyrus mediated the relationship between environmental measures and changes on the Picture Sequence Memory Task (Supplementary Table 51). Additionally, changes in the right isthmus of the cingulate cortex mediated the relationship between the micro-environment and scores on the Picture Vocabulary Task (Supplementary Table 52).

### Sensitivity analysis

To evaluate the robustness of the observed findings, a series of additional sensitivity analyses were conducted. First, associations were re-examined using an alternative indicator of the macro-environment - the Area Deprivation Index (ADI). The ADI provides a neighbourhood-level composite score based on census-based indicators of income, education, employment, and housing conditions. The measure most closely matches the COI 2.0 Social and Economic and Education domains. Pearson’s correlations on standardised beta coefficients indicated a high concordance between results from the ADI and COI 2.0 (Supplementary Table 53 & Figures 1-2).

Second, to account for the potential confounding effects of puberty, analyses were reconducted using pubertal stage (for baseline analysis) or change in pubertal stage (for longitudinal analysis) as a covariate. Although in some cases fewer regions reached statistical significance in these adjusted models, the overall direction and magnitude of effects generally remained consistent and high correlations were observed with the primary findings (Supplementary Table 54 & Figures 3-4).

## Discussion

The present study examined baseline and longitudinal associations between the environment, brain, and behaviour in adolescence. At baseline, we found that children with a better macro-and micro-environment showed larger brain volumes relative to their peers. Both better environments and higher brain centile scores were also associated with better mental health and cognitive outcomes. Importantly, in several cases, the relationship between environmental factors and cognitive and mental health outcomes was partially mediated via the brain. Longitudinally, we found that adolescents with a better childhood macro- and micro-environment moved higher up in their centile rankings across adolescence. Moreover, adolescents who moved upwards in their centile scores in targeted regions showed improvements in mental health and cognition. The childhood macro- and micro-environments were associated with changes in cognition, but not in mental health, across adolescence. The environment-brain-behaviour pathway therefore offered some explanatory power in predicting longitudinal changes in cognition across adolescence.

First, the findings provide confirmatory evidence for the associations between the environment and brain structure. At baseline, those with a better macro- and micro-environment showed higher brain volumes relative to their peers. Associations were observed across the brain and were particularly pronounced in multisensory and sensorimotor regions (e.g., middle temporal, lateral occipital, inferior temporal, precentral and postcentral gyri). Longitudinally, those with better environments showed increases in their relative centile rankings across adolescence. Associations were particularly pronounced in higher-order associative and multi-modal integrative regions (e.g., precuneus, insula, pericalcarine, lingual, inferior parietal, and middle temporal gyri). This trend aligns with known patterns of cortical maturation, where sensory regions develop first followed later by higher-order associative regions^9^. Environmental associations with brain structure therefore align with the general trajectory of cortical development.

Our findings further demonstrate that, at baseline, a better macro- and micro-environment was associated with better cognitive and mental health outcomes across all measured domains, consistent with a large body of previous literature^15,31,41^. Longitudinally, a better childhood micro- and macro-environment was associated with increases on all the measured cognitive tasks across adolescence. However, the childhood environment did not predict changes in mental health traits. There are several potential explanations for these findings. Firstly, given that environmental factors are dynamic and may change at the individual-level, it is possible that the childhood environment is not a reliable predictor of mental health changes across adolescence, which may be more sensitive to the current environment. As such, measures of the environment that are more temporally aligned with mental health measures might result in more prominent associations. It is also possible that, while environment plays a significant role in shaping baseline effects, it may not play a further additive role in shaping longitudinal change across adolescence. Instead, these relationships may already become established pre-adolescence and, subsequently, the initial effects of the environment on the brain and mental health may become self-reinforcing in driving longitudinal change. Alternatively, it is also possible that the substantial reorganisation of the brain during adolescence may overshadow the impact of baseline environmental effects.

Indeed, as expected, brain structure significantly predicted mental health and cognitive scores, both at baseline and longitudinally. Higher baseline centile scores across multiple brain regions predicted lower internalising and externalising traits. Longitudinally, individuals showing decreases in their relative centile rankings across adolescence showed increased traits of poor mental health. Prominent regions showing associations with externalising included structures such as the insula and lateral orbitofrontal cortex, which have previously been implicated in processes such as self-awareness^42^, emotion regulation^43^, impulse control^44^, and reward processing^43^. Structural variations within the insula, in particular, are implicated across psychiatric conditions, and more specifically with externalising psychopathologies^45^. Associations were also observed across a wide range of regions, including the visual and associative regions (e.g., cuneus, pericalcarine, lateral occipital, and lingual cortex) as well as semantic and motor regions (e.g., middle temporal gyrus, pars triangularis, inferior parietal cortex, and precentral gyrus). Although these regions have been linked to a broad set of functions, some research suggests that various of them may also be involved externalising-relevant processes such as inhibitory control^45–48^, attention control^49^, and impulsivity^49^. Indeed, structural variations in several of these regions have also previously been associated with externalising traits^53–55^. For instance, largely consistent with the findings of the present study, it has previously been shown that externalising traits are linked to negative deviations in gray matter volumes from peers across structures such as the cuneus, inferior parietal lobe, lingual gyrus, and precentral gyrus in childhood and adolescence^50^.

Several regions associated with externalising also overlapped with those associated with internalising, including the insula, cuneus, lingual gyrus, pars triangularis, and pericalcarine gyrus. This is consistent with prior evidence suggesting that internalising and externalising show both overlapping as well as distinct neural correlates^51,52^. These regions have also been implicated in internalising-relevant traits and exhibit structural variations in internalising conditions such as depression^53–55^. The insula, for instance, has been linked to interoception (potentially explaining somatic aspects of internalising) and emotion processing^42^, with smaller volumes and reduced surface areas showing associations with internalising^51,54^. Regions specific to internalising included the fusiform gyrus and default mode regions such as the isthmus cingulate and parahippocampal gyrus – previously implicated in memory and self-referential cognition^53^. Likewise, reduced volume, surface area, cortical thickness and gray matter density across these regions have been linked to depression and anxiety^55–57^.

With respect to cognition, at baseline, children whose brain volumes were larger relative to age and sex-matched peers showed higher performance across the range of tested cognitive tasks measured. Longitudinally, more region-specific and task-dependent associations were observed with mixed directionality. For instance, increases on the Flanker Inhibitory Control and Attention Task were associated with downward shifts in the lateral orbitofrontal cortex, implicated in inhibitory control^44^, as well as the precuneus, implicated in self-referential processing^58^ and attention-shifting^59^. Increases on the Oral Reading Recognition task were likewise associated with downward shifts in the precuneus and the pars ocupalaris, both of which have been linked to aspects of language and semantic processing^60,61^. On the other hand, increases on the Picture Memory Task, which assesses episodic memory and sequencing, were associated with upward shifts in centile scores across the brain. Increases on the Picture Vocabulary Task were also associated with upward shifts in the isthmus of the cingulate cortex, while performance on the Pattern Comparison Processing Speed Task was associated with downward shifts in the superior temporal sulcus, implicated in various broad aspects of cognition (e.g., sensory integration, attention control)^62^. Although seemingly counterintuitive, it is possible that these downward shifts, in fact, reflect more advanced maturation, since overall reductions in gray matter volumes are typical during adolescent development^63^. These effects may be region and task-dependent, with certain tasks requiring increasing centiles in specific regions for better performance while others requiring decreased centiles. It is also important to note that since these analyses are based primarily on structural measures, any functional interpretations are largely speculative and should be interpreted with caution.

Finally, a key aim of the present study was to investigate the environment-brain-behaviour mediation pathway. At baseline, several such mediation effects were identified - for instance, an improved macro-environment was associated with higher centile scores, which, in turn, predicted lower internalising and externalising. Centile scores also mediated the association between the macro-environment and scores on the Flanker Inhibitory Control and Attention task, as well as the association between both the micro- and macro-environment and scores on the Oral Reading Recognition Task. These results suggest that the associations between the environment and behaviour are mediated, at least partially, via the brain.

Longitudinally, changes in regions such as the cuneus and lateral occipital gyrus mediated the relationship between environmental measures and changes on the Picture Memory Task. Similarly, changes in the cingulate cortex mediated the relationship between the micro-environment and scores on the Picture Sequence Memory Task. These findings suggest that the childhood environment predicts aspects of cognitive development across adolescence partly through modulating changes in brain structure. Besides these associations, no other longitudinal mediation effects were observed for the other outcomes examined. This can largely be attributed to the fact that the baseline environment was not predictive of longitudinal changes across outcome such as internalising and externalising, which precluded further mediation analyses.

There are a number of limitations to consider when interpreting these findings. First, mediation analyses typically require high statistical power. In the present study, the longitudinal sample, although still large, was smaller than the baseline cohort. As a result, some subtle mediation effects may not have been detected in the longitudinal analyses due to comparatively lower power. Secondly, while brain and behavioural changes were measured longitudinally, environmental factors were measured only at baseline. It is possible that *changes* in the environment may be more informative in predicting changes in the brain, mental health, and cognition across adolescence. This is particularly important considering that the COVID-19 pandemic occurred during data collection and resulted in various substantial changes to the health, social, economic, and educational environment. Third, macro-environmental variables were derived from geospatial data, which hold limited spatial and temporal resolution. Fourth, the ABCD sample is concentrated in US metropolitan areas, which may limit the global generalisability of these findings. Fifth, because several environmental factors are highly collinear and interrelated, we used summary-level scores as a method to parsimoniously obtain a broad overview of the impact of the environment on brain development and behaviour. However, as a result, this approach limited inference on the specific contributions of individual environmental predictors. Additionally, some other important aspects of the adolescent micro-environment, such as cyberbullying or discrimination, were not considered as these measures were not fully available for all timepoints in the ABCD dataset. Finally, several environmental factors that were investigated are not fully separable from genetic factors. For instance, micro-environmental factors such as parental education can be highly heritable ^64^ and are also linked with macro-environmental factors such as neighbourhood characteristics. It will therefore be important for future work to assess the extent to which the observed associations persist after accounting for genetic factors.

In light of these limitations, future studies may consider disentangling the roles of individual environmental predictors on outcomes of interest. Future research should also consider incorporating both repeated measures of the environment as well as additional brain-based measures (e.g., white matter microstructure, functional connectivity). Collectively, these measures will produce more explanatory mediation pathways. Studies should also consider integrating genetic data into longitudinal environment-brain-behaviour frameworks to clarify how genetic predispositions may moderate sensitivity to environmental effects on the brain and behaviour. Finally, since adolescence is also a critical period for sex differences in the brain and behaviour^65^, sex differences in the environment-brain-behaviour mediation pathway should also be explored further.

In summary, the present study identified both baseline and longitudinal associations between the environment, brain, and behaviour during adolescence. Key strengths include its large sample size, longitudinal design, and use of brain metrics benchmarked against population-based norms. The findings represent an important step towards the identification of brain markers and enviro-types that shape cognition and mental health during this critical period of development. In turn, this may support the development of both neurobehavioral interventions and policy initiatives aimed at improving the environment to promote positive outcomes for adolescents.

## Methods

### Participants

The ABCD Study is a longitudinal study spanning across 21 research sites in the United States^35^. The study obtained ethical approval from a central Institutional Review Board at the University of California, San Diego, as well as local approval from each of the participating sites. General exclusion criteria for the ABCD study included an MRI contraindication, major neurological disorder, gestational age at birth <28 weeks, birth weight <1200g, birth complications that required hospitalisation for >1 month, uncorrected vision, and a diagnosis of schizophrenia, autism, learning disability, and alcohol or substance use disorder at the time of enrolment. Participants were also excluded if they did not pass an MRI quality control check (https://wiki.abcdstudy.org/release-notes/imaging/qualitycontrol.html).

The baseline sample consists of 6,911 participants aged 8-11 years (Mean = 9.93, SD = 0.63). Participants with any missing measurements of interest (structural MRI, environmental data, cognitive tests, mental health questionnaires) were excluded from the analysis. Sample demographic characteristics are presented in Table 3.

**Table 3.**
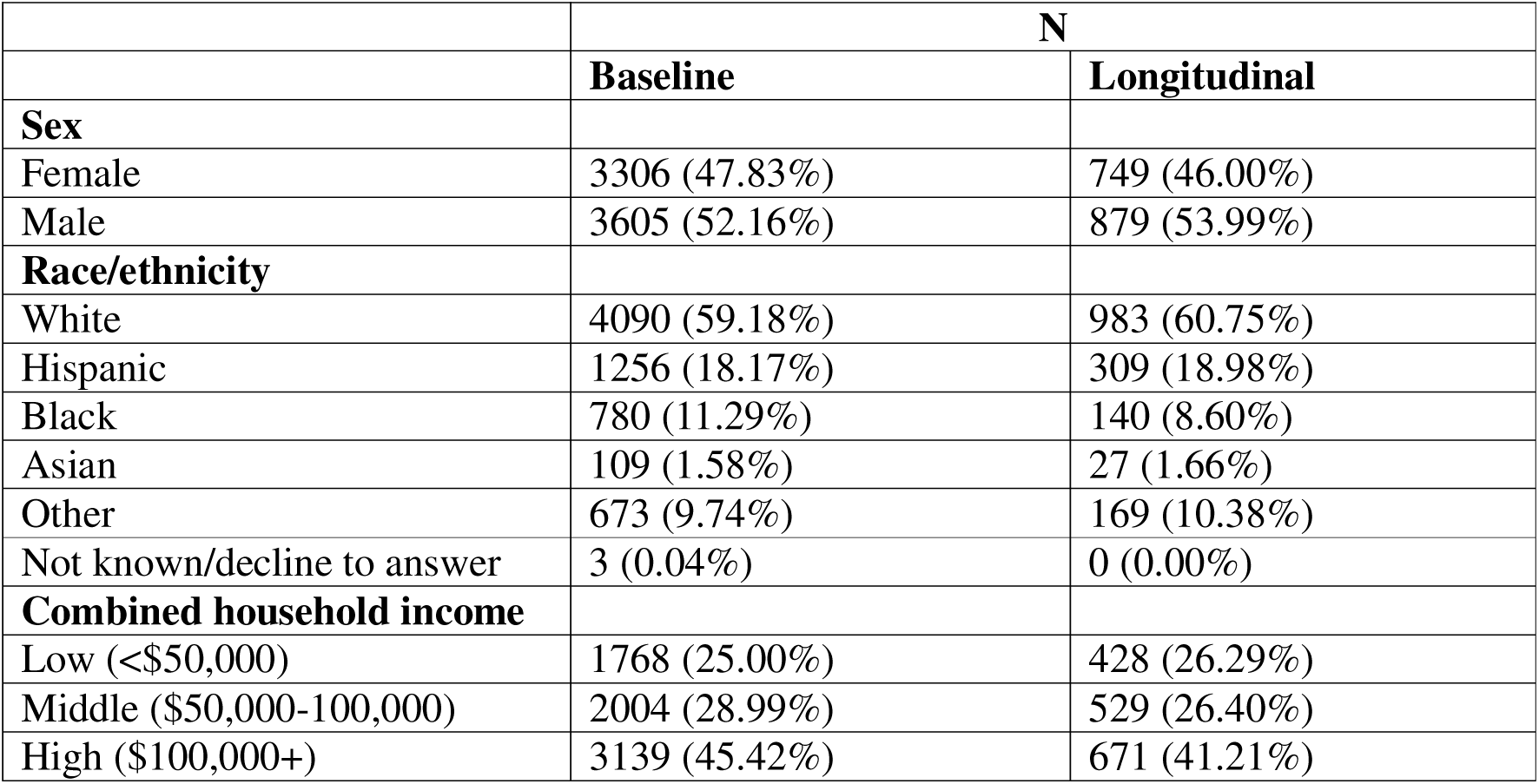
Sample demographic characteristics. The table presents the number and percent of participants by sex, race/ethnicity, and combined household income, separately for the baseline and longitudinal data.

|  | N |  |
| --- | --- | --- |
|  | Baseline | Longitudinal |
| **Sex** |  |  |
| Female | 3306 (47.83%) | 749 (46.00%) |
| Male | 3605 (52.16%) | 879 (53.99%) |
| **Race/ethnicity** |  |  |
| White | 4090 (59.18%) | 983 (60.75%) |
| Hispanic | 1256 (18.17%) | 309 (18.98%) |
| Black | 780 (11.29%) | 140 (8.60%) |
| Asian | 109 (1.58%) | 27 (1.66%) |
| Other | 673 (9.74%) | 169 (10.38%) |
| Not known/decline to answer | 3 (0.04%) | 0 (0.00%) |
| **Combined household income** |  |  |
| Low (<\$50,000) | 1768 (25.00%) | 428 (26.29%) |
| Middle (\$50,000-100,000) | 2004 (28.99%) | 529 (26.40%) |
| High (\$100,000+) | 3139 (45.42%) | 671 (41.21%) |

The longitudinal cohort consisted of data from 3 timepoints spanning ages 8-15 (timepoint 1: ages 8-11, timepoint 2: ages 10-13, timepoint 3: ages 12-15). Only participants that had all measures of interest across all 3 timepoints were included in the analysis. The sample consisted of 1,628 participants. Sample demographic characteristics are presented in Table 3. Both the baseline and longitudinal samples presented with largely similar descriptive characteristics.

### Measures

#### Macro-environment

The Child Opportunity Index (COI 2.0) provides linked external data with various area-level indices of the macro-environment^36,37^. It consists of 29 indicators summarised into three summary domains: Social and Economic (e.g., employment rate, poverty rate, homeownership rate, median household income), Education (e.g., school poverty, high school graduation rate, college enrolment, ECE centres), and Health and Environment (e.g., access to healthy food, access to green space, industrial pollutants, health insurance coverage). This information was obtained from publicly available sources (e.g., the American Community Survey, U.S. Department of Agriculture) and aggregated at the census tract level based on the participants’ baseline residential addresses. Details are provided in Table 4.

**Table 4.**
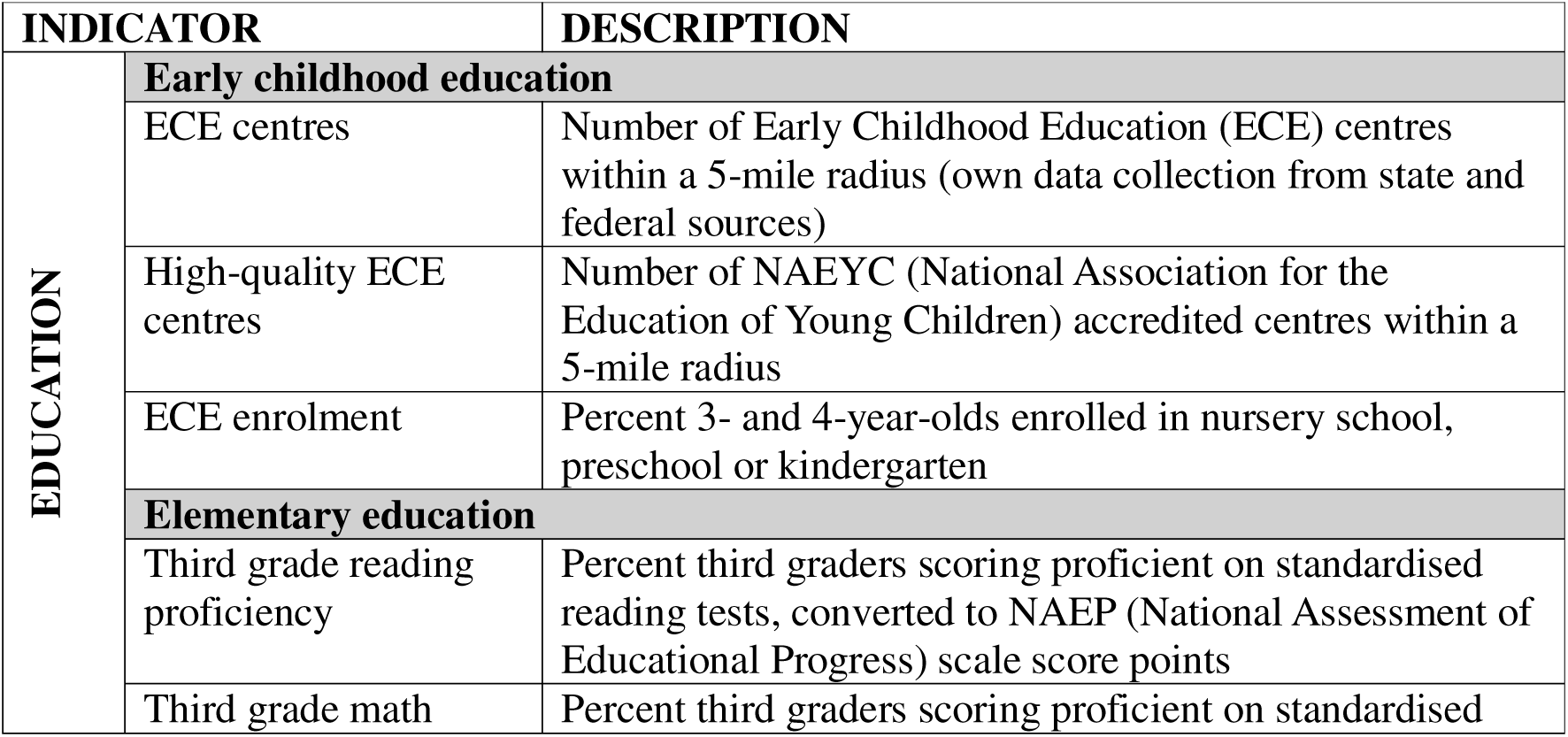
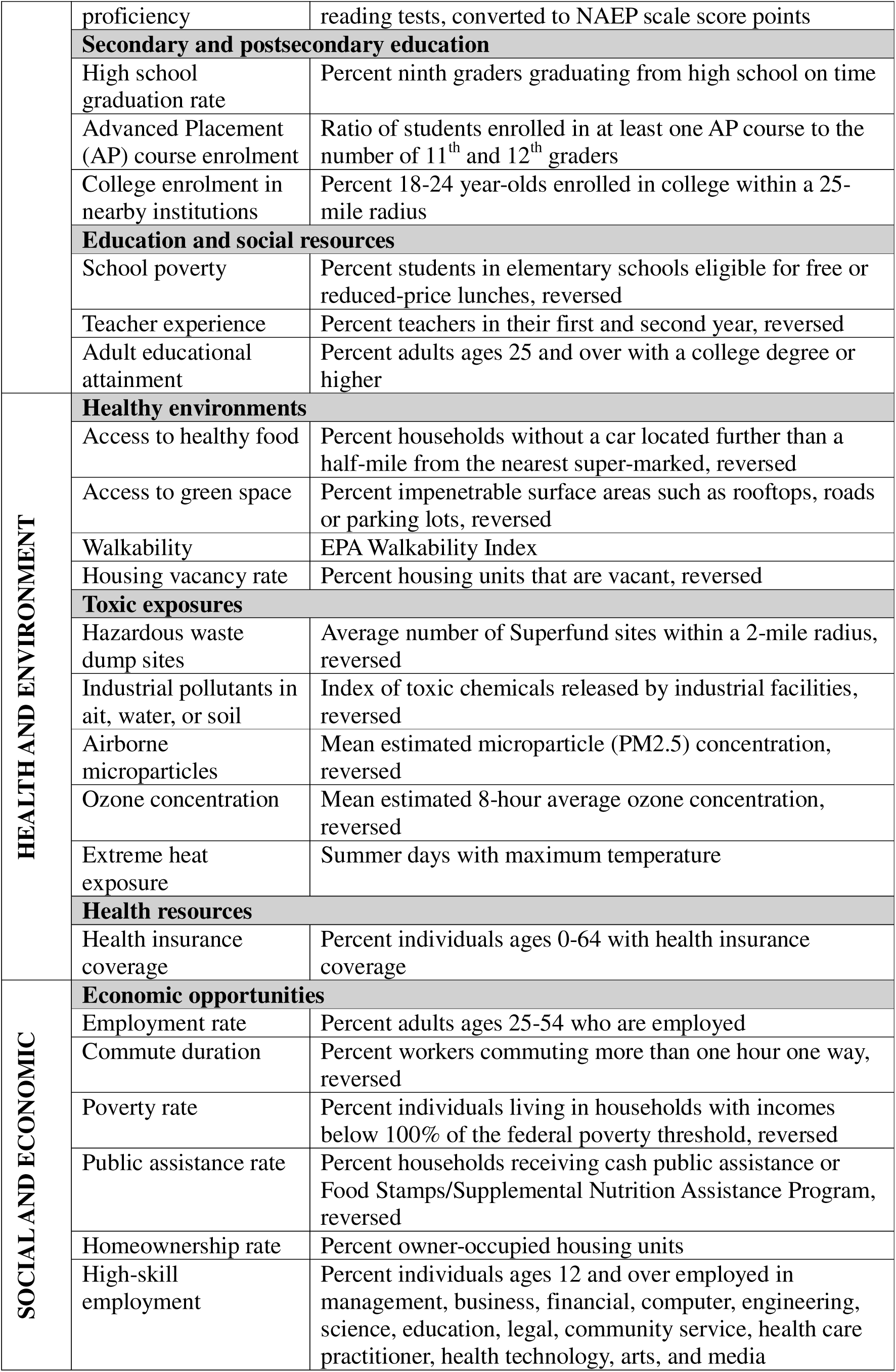
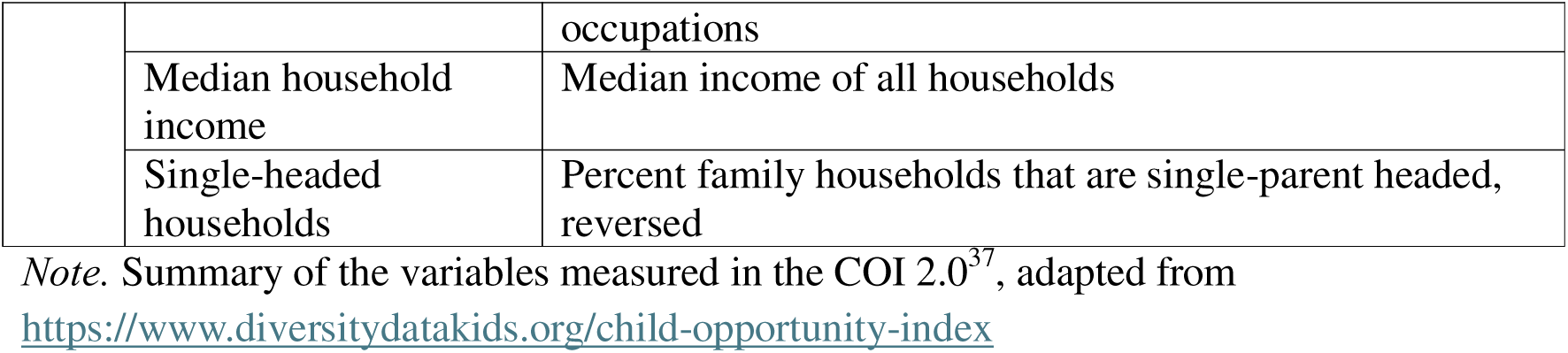
Childhood Opportunity Index.

**Table 4.** Micro-environment summary latent factor score.

| Groups | Measures |
| --- | --- |
| **Economic** | Food security, housing security, ability to pay bills, access to medical/dental |
| **Parental** | Parental education, total caregiver warmth, parental monitoring, dual parent households |
| **ACEs** | Family conflict, history of a traumatic event, parent psychopathology |
| **School/community** | Neighbourhood safety, positive school environment, school engagement |
| **Physiological** | Sleep hours, BMI |
| **Perinatal** | Total prenatal conditions, planned pregnancy, maternal age at birth, history of prenatal substance use, gestational age (weeks), birth weight (kg) |

#### Micro-environment

A latent factor score, derived from a previous study via group factor analysis^38^, was used as a summary measure of the micro-environment. The latent factor score was generated from 22 proximal measures across six domains: economic security (e.g., food security, housing security, ability to pay bills, and access to medical care), parental characteristics (e.g., education, dual parent households, parental monitoring, caregiver warmth), school/community environment (neighbourhood safety, positive school environment), adverse childhood experiences (e.g., family conflict, history of a traumatic event, parental psychopathology), physiological health (e.g., sleep hours, BMI), and perinatal wellbeing (e.g., gestational age at birth, birth weight). Further details are provided in Table 4.

#### Mental Health

The Childhood Behaviour Checklist (CBCL) is a validated, parent-report questionnaire that measures psychopathological traits^39^. The measure can be grouped into two higher-order summary factors: internalising, which assesses sadness, depression, anxiety, and loneliness traits (e.g., “feels worthless or inferior” or “complains of loneliness”), and externalising, which assesses aggression and rule-breaking traits (“cruelty, bullying, or meanness to others” or “gets in many fights”).

#### Cognition

The NIH Toolbox Cognition Battery (NIH Toolbox®) is a standardised assessment measuring neurocognition across a range of key domains^40^. In the ABCD study, the following tasks were measured longitudinally across the three timepoints: (a) Flanker Inhibitory Control & Attention Task: measures attention, cognitive control, executive function, and inhibition of automatic response, (b) Picture Sequence Memory Task: measures episodic memory and sequencing, (c) Pattern Comparison Processing Speed Task: measures information processing and processing speed, (d) Picture Vocabulary Task: measures language vocabulary knowledge, and (e) Oral Reading Recognition Task: measures language, oral reading skills, and academic achievement. Together, these serve as representative measures of cognitive functioning across a range of domains. In line with the recommendations for longitudinal analysis, age-uncorrected scores were used in the present study.

#### Mental health and cognition change scores

To measure longitudinal change in mental health and cognition across the three timepoints, we calculated an age-normalised change metric. This metric was calculated by averaging the change in scores between the timepoints (e.g., change between T1 and T2, change between T2 and T3) and adjusting for the age differences between timepoints. The following calculation was computed for each participant:

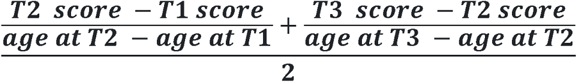

#### Centile scores

To compute centile scores, we benchmarked each participant’s regional brain volumes against a population-based reference. The reference model was derived from a previous large-scale study that modelled sex-stratified trajectories of brain volumes using GAMLSS in over 100,000 individuals across the lifespan^8^. We then computed, for each individual, the relative distance of each regional brain volume from the median of the age and sex-specific distribution provided by the reference model. This score, expressed in percentiles, reflected where an individual’s brain volume fell in comparison to same-age and same-sex peers.

For the longitudinal analysis, we averaged the change in centile scores between the timepoints (change between T1 and T2 and change between T2 and T3) as follows:

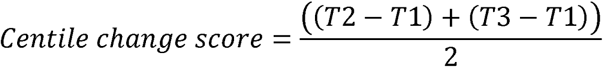

Given that centile scores are already age-adjusted, we did not further correct this measure for age.

#### Statistical Analysis

All associations were examined in both the baseline data as well as the longitudinal data from all three timepoints. In the baseline analysis, mixed effects models were used to assess (1) The associations between each of the four environmental variables with each baseline regional centile score, (2) The association between each baseline regional centile with each baseline measure of mental health and cognition, (3) The association between each of the four environmental variables with each baseline measure of mental health and cognition, and (4) The environment – brain – behaviour mediation pathway for all combinations of environment, brain, mental health, and cognitive measures that showed significant associations in steps 1-3. These analysis steps are visualised in Figure 1.

For each analysis, testing site and family ID (to account for sibling pairs) were modelled as random intercepts, while age and sex were modelled as covariates. Benjamini-Hochberg False Discovery Rate (FDR) corrections were applied within the following steps of analyses – (a) For analysis steps 1 and 2, FDR corrections were applied separately for each variable of interest to encompass all regional brain volumes, (b) For analysis step 3, a single FDR correction was applied across all analyses to encompass all combinations of environmental and cognitive/mental health measures, and (c) for analysis step 4, FDR corrections were applied within each mediation pathway to encompass all pre-selected regional volumes of interest.

In the longitudinal analysis, mixed effects models were used to assess (1) The associations between each of the four environmental variables with each regional centile change score, (2) (2) The association between each regional centile change score with each longitudinal change measure of mental health and cognition (3) The association between each of the four environmental variables with each longitudinal measure of mental health and cognition, and (4) The longitudinal environment – brain – behaviour mediation pathway for all combinations of environment, brain, mental health, and cognitive measures that showed significant associations in steps 1-3. Similar to the baseline analyses, each analysis modelled random intercepts for testing site and family ID. Additionally, all analyses were controlled for sex but not for age, as ages were already taken into consideration when deriving the change scores. Analysis steps 2-4 also further controlled for baseline mental health and cognition scores. FDR corrections were applied in the same manner as the baseline analysis.

We also conducted additional sensitivity analyses to assess the consistency of the observed results. First, we re-conducted the analyses controlling all for pubertal stage, measured via the Pubertal Development Scale (PDS)^66^. The PDS measures the development of secondary sexual characteristics (e.g., body hair) and also includes sex-specific measures for males (e.g., facial hair, voice changes) and females (e.g., breast development and menarche). Based on this information, a 5-point scale is used to represent the pubertal stage of each participant (i.e., pre-puberty, early puberty, mid-puberty, late puberty). In the baseline analysis, this pubertal stage was modelled as a covariate. For the longitudinal analysis, a puberty change metric was derived by calculating the difference in PDS scores between the first and final timepoints to approximate an adolescent’s pubertal progression over the studied period. To assess concordance, Pearson’s correlations were computed on the standardised beta coefficients from the main analysis and puberty-controlled analysis.

To assess the replicability of findings across different measures of the macro-environment, we also conducted additional analyses using the Area Deprivation Index (ADI). Like the COI 2.0, the ADI provides neighbourhood-level composite score based on census-based indicators of income, education, employment, and housing conditions. Pearson’s correlations were computed on the standardised beta coefficients of the COI 2.0 and ADI to determine concordance.

## Acknowledgements

Data used in the preparation of this article were obtained from the Adolescent Brain Cognitive Development^SM^ (ABCD) Study (https://abcdstudy.org), held in the NIMH Data Archive (NDA). This is a multisite, longitudinal study designed to recruit more than 10,000 children age 9-10 and follow them over 10 years into early adulthood. The ABCD Study® is supported by the National Institutes of Health and additional federal partners under award numbers U01DA041048, U01DA050989, U01DA051016, U01DA041022, U01DA051018, U01DA051037, U01DA050987, U01DA041174, U01DA041106, U01DA041117, U01DA041028, U01DA041134, U01DA050988, U01DA051039, U01DA041156, U01DA041025, U01DA041120, U01DA051038, U01DA041148, U01DA041093, U01DA041089, U24DA041123, U24DA041147. A full list of supporters is available at https://abcdstudy.org/federal-partners.html. A listing of participating sites and a complete listing of the study investigators can be found at https://abcdstudy.org/consortium_members/. ABCD consortium investigators designed and implemented the study and/or provided data but did not necessarily participate in the analysis or writing of this report. This manuscript reflects the views of the authors and may not reflect the opinions or views of the NIH or ABCD consortium investigators. We are also grateful to Manya Gupta for assisting with organising the supplementary materials for this article.

## Author Contributions

Conceptualisation: YTK and RAIB. Methodology: YTK, RAIB, RB. Formal analysis: YTK. Writing – original draft: YTK. Writing – Review & Editing: YTK, RAIB, SJB, RB, JS, AAB, AT, CA, LD, and SBC.

