## Supplementary Materials for "Associations between the environment, brain, mental health, and cognition across adolescence"

### Environmental factors influencing brain development, mental health, and cognition across adolescence

### Table of Contents

**Supplementary Table 1:** Baseline associations between regional centile scores and COI 2.0 health and environment summary score………………………………………………………………….5-6

**Supplementary Table 2:** Baseline associations between regional centile scores and COI 2.0 social and economic summary score……………………………………………………………………..7-8

**Supplementary Table 3:** Baseline associations between regional centile scores and COI 2.0 education summary score…………………………………………………………………………9-10

**Supplementary Table 4:** Baseline associations between regional centile scores and micro-environment latent factor score…………………………………………………………………...11-12

**Supplementary Table 5:** Baseline associations between regional centile scores and CBCL externalising traits………………………………………………………………………………...13-14

**Supplementary Table 6:** Baseline associations between regional centile scores and CBCL internalising traits…………………………………………………………………………………15-16

**Supplementary Table 7:** Baseline associations between regional centile scores and scores on the Flanker Inhibitory Control & Attention Task …………………………………………………….17-18

**Supplementary Table 8:** Baseline associations between regional centile scores and scores on the Oral Reading Recognition Task ……………………………………………………………………….19-20

**Supplementary Table 9:** Baseline associations between regional centile scores and scores on the Picture Vocabulary Task ………………………………………………………………………….21-22

**Supplementary Table 10:** Baseline associations between regional centile scores and scores on the Picture Vocabulary Task ………………………………………………………………………….23-24

**Supplementary Table 11:** Baseline associations between regional centile scores and scores on the Pattern Comparison Processing Speed Task……………………………………………………...25-26

**Supplementary Table 12:** Regions mediating the associations between COI 2.0 health/environment scores and externalising scores at baseline……………………………………………………….27-28

**Supplementary Table 13:** Regions mediating the associations between COI 2.0 social/economic scores and externalising scores at baseline……………………………………………………….29-30

**Supplementary Table 14:** Regions mediating the associations between COI 2.0 education scores and externalising scores at baseline ………………………………………………………………....31-32

**Supplementary Table 15:** Regions mediating the associations between micro-environment scores and externalising scores at baseline……………………………………………………………..33-34

**Supplementary Table 16:** Regions mediating the associations between COI 2.0 health/environment scores and internalising scores at baseline…………………………………………………………35

**Supplementary Table 17:** Regions mediating the associations between COI 2.0 social/economic scores and internalising scores at baseline………………………………………………………...36

**Supplementary Table 18:** Regions mediating the associations between COI 2.0 education scores and internalising scores at baseline…………………………………………………………………….37

**Supplementary Table 19:** Regions mediating the associations between micro-environment scores and internalising scores at baseline………………………………………………………………...38

**Supplementary Table 20:** Regions mediating the associations between health/environment scores and Flanker Inhibitory Control & Attention scores at baseline…………………………………39-40

**Supplementary Table 21:** Regions mediating the associations between social/economic scores and Flanker Inhibitory Control & Attention scores at baseline……………………………………...41-42

**Supplementary Table 22:** Regions mediating the associations between education scores and Flanker Inhibitory Control & Attention scores at baseline……………………………………………....43-44

**Supplementary Table 23:** Regions mediating the associations between micro-environment scores and Flanker Inhibitory Control & Attention scores at baseline………………………………....45-46

**Supplementary Table 24:** Regions mediating the associations between health/environment scores and Pattern Comparison Processing Speed scores at baseline………………………………….47-48

**Supplementary Table 25:** Regions mediating the associations between social/economic scores and Pattern Comparison Processing Speed scores at baseline……………………………………....49-50

**Supplementary Table 26:** Regions mediating the associations between education scores and Pattern Comparison Processing Speed scores at baseline……………………………………………….51-52

**Supplementary Table 27:** Regions mediating the associations between micro-environment scores and Pattern Comparison Processing Speed scores at baseline…………………………………….53

**Supplementary Table 28:** Regions mediating the associations between COI 2.0 health/environment scores and Picture Vocabulary scores at baseline……………………………………………….54-55

**Supplementary Table 29:** Regions mediating the associations between COI 2.0 social/economic scores and Picture Vocabulary scores at baseline……………………………………………….56-57

**Supplementary Table 30:** Regions mediating the associations between COI 2.0 education scores and Picture Vocabulary scores at baseline…………………………………………………………...58-59

**Supplementary Table 31:** Regions mediating the associations between micro-environment scores and Picture Vocabulary scores at baseline……………………………………………………....60-61

**Supplementary Table 32:** Regions mediating the associations between COI 2.0 health/environment scores and Picture Sequence Memory scores at baseline……………………………………….62-63

**Supplementary Table 33:** Regions mediating the associations between COI 2.0 social/economic scores and Picture Sequence Memory scores at baseline……………………………………….64-65

**Supplementary Table 34:** Regions mediating the associations between COI 2.0 education scores and Picture Sequence Memory scores at baseline…………………………………………………...66-67

**Supplementary Table 35:** Regions mediating the associations between micro-environment scores and Picture Sequence Memory scores at baseline………………………………………………68-69

**Supplementary Table 36:** Regions mediating the associations between COI 2.0 health/environment scores and Oral Reading Recognition scores at baseline………………………………………..70-71

**Supplementary Table 37:** Regions mediating the associations between COI 2.0 social/economic scores and Oral Reading Recognition scores at baseline………………………………………..72-73

**Supplementary Table 38:** Regions mediating the associations between COI 2.0 education scores and Oral Reading Recognition scores at baseline…………………………………………………....74-75

**Supplementary Table 39:** Regions mediating the associations between micro-environment scores and Oral Reading Recognition scores at baseline………………………………………………..76-77

**Supplementary Table 40:** Longitudinal associations between regional centile change scores and COI 2.0 health and environment summary change scores…………………………………………….78-79

**Supplementary Table 41:** Longitudinal associations between regional centile change scores and COI 2.0 social and economic summary change scores………………………………………………..80-81

**Supplementary Table 42:** Longitudinal associations between regional centile change scores and COI 2.0 education summary change scores…………………………………………………………...82-83

**Supplementary Table 43:** Longitudinal associations between regional centile change scores and micro-environment latent factor change scores………………………………………………….84-85

**Supplementary Table 44:** Longitudinal associations between regional centile change scores and CBCL internalising change scores………………………………………………………………86-87

**Supplementary Table 45:** Longitudinal associations between regional centile change scores and CBCL externalising change scores……………………………………………………………...88-89

**Supplementary Table 46:** Longitudinal associations between regional centile change scores and Flanker Inhibitory Control and Attention Task change scores………………………………….90-91

**Supplementary Table 47:** Longitudinal associations between regional centile change scores and Picture Sequence Memory Task change scores…………………………………………………92-93

**Supplementary Table 48:** Longitudinal associations between regional centile change scores and Oral Reading Recognition Task change scores………………………………………………………94-95

**Supplementary Table 49:** Longitudinal associations between regional centile change scores and Pattern Comparison Processing Speed change scores………………………………………….96-97

**Supplementary Table 50:** Longitudinal associations between regional centile change scores and Picture Vocabulary Task change scores………………………………………………………….98-99

**Supplementary Table 51:** Regions mediating the associations between environmental variables and changes in the Picture Sequence Memory Task……………………………………………….100-101

**Supplementary Table 52:** Regions mediating the associations between environmental variables and changes on the Picture Vocabulary Task…...………………………………………………………102

**Supplementary Table 53:** Correlations between COI 2.0 and ADI analysis………………….…..103

**Supplementary Table 54:** Correlations between main analysis and puberty-controlled analysis...104

**Supplementary Figure 1:** ADI Baseline Figure…………………………………………………..105

**Supplementary Figure 2:** ADI Longitudinal Figure……………………………..……………….106

**Supplementary Figure 3:** Puberty Baseline Figure……………………………..….…………….107

**Supplementary Figure 4:** Puberty Longitudinal Figure……………………………..……..…….109

**Supplementary Table 1: Baseline associations between regional centile scores and COI 2.0 health and environment summary score**

Values shown are unstandardised and standardised beta coefficients with associated standard errors (SE) for each region. P values are false discovery rate (FDR) corrected to account for multiple comparisons. All analyses controlled for age and sex and included random intercepts for testing site and family ID (to account for sibling pairs).

| **Region** | **Unstandardised estimate (SE)** | **Standardised estimate (SE)** | ***p*_FDR_** |
| --- | --- | --- | --- |
| Left Banks of the Superior Temporal Sulcus | 0.013 (0.002) | 0.065 (0.012) | 0.000 |
| Left Caudal Anterior Cingulate Cortex | 0.010 (0.003) | 0.047 (0.013) | 0.000 |
| Left Caudal Middle Frontal Gyrus | 0.019 (0.003) | 0.092 (0.013) | 0.000 |
| Left Cuneus | 0.021 (0.003) | 0.102 (0.014) | 0.000 |
| Left Entorhinal Cortex | 0.018 (0.003) | 0.091 (0.013) | 0.000 |
| Left Frontal Pole | 0.013 (0.003) | 0.067 (0.013) | 0.000 |
| Left Fusiform Gyrus | 0.022 (0.003) | 0.109 (0.014) | 0.000 |
| Left Inferior Parietal Lobule | 0.013 (0.003) | 0.065 (0.013) | 0.000 |
| Left Inferior Temporal Gyrus | 0.029 (0.003) | 0.145 (0.014) | 0.000 |
| Left Insular Cortex | 0.017 (0.003) | 0.086 (0.013) | 0.000 |
| Left Isthmus of the Cingulate Cortex | 0.015 (0.003) | 0.076 (0.013) | 0.000 |
| Left Lateral Occipital Cortex | 0.028 (0.003) | 0.136 (0.014) | 0.000 |
| Left Lateral Orbitofrontal Cortex | 0.026 (0.003) | 0.127 (0.014) | 0.000 |
| Left Lingual Gyrus | 0.017 (0.003) | 0.087 (0.014) | 0.000 |
| Left Medial Orbitofrontal Cortex | 0.010 (0.003) | 0.049 (0.012) | 0.000 |
| Left Middle Temporal Gyrus | 0.030 (0.003) | 0.147 (0.014) | 0.000 |
| Left Paracentral Lobule | 0.017 (0.003) | 0.084 (0.013) | 0.000 |
| Left Parahippocampal Gyrus | 0.014 (0.003) | 0.068 (0.013) | 0.000 |
| Left Pars Opercularis | 0.015 (0.002) | 0.073 (0.012) | 0.000 |
| Left Pars Orbitalis | 0.018 (0.003) | 0.089 (0.012) | 0.000 |
| Left Pars Triangularis | 0.009 (0.002) | 0.043 (0.012) | 0.001 |
| Left Pericalcarine Cortex | 0.011 (0.003) | 0.057 (0.013) | 0.000 |
| Left Postcentral Gyrus | 0.029 (0.003) | 0.143 (0.014) | 0.000 |
| Left Posterior Cingulate Cortex | 0.014 (0.003) | 0.070 (0.013) | 0.000 |
| Left Precentral Gyrus | 0.031 (0.003) | 0.154 (0.014) | 0.000 |
| Left Precuneus | 0.027 (0.003) | 0.133 (0.014) | 0.000 |
| Left Rostral Anterior Cingulate Cortex | 0.012 (0.003) | 0.060 (0.013) | 0.000 |
| Left Rostral Middle Frontal Gyrus | 0.018 (0.003) | 0.089 (0.013) | 0.000 |
| Left Superior Frontal Gyrus | 0.022 (0.003) | 0.107 (0.014) | 0.000 |
| Left Superior Parietal Lobule | 0.023 (0.003) | 0.116 (0.014) | 0.000 |
| Left Superior Temporal Gyrus | 0.016 (0.003) | 0.080 (0.013) | 0.000 |
| Left Supramarginal Gyrus | 0.021 (0.003) | 0.103 (0.014) | 0.000 |
| Left Temporal Pole | 0.010 (0.003) | 0.047 (0.012) | 0.000 |
| Left Transverse Temporal Gyrus (Heschl’s Gyrus) | 0.013 (0.002) | 0.066 (0.012) | 0.000 |
| Right Banks of the Superior Temporal Sulcus | 0.015 (0.003) | 0.076 (0.013) | 0.000 |
| Right Caudal Anterior Cingulate Cortex | 0.011 (0.003) | 0.055 (0.012) | 0.000 |
| Right Caudal Middle Frontal Gyrus | 0.015 (0.002) | 0.073 (0.012) | 0.000 |
| Right Cuneus | 0.020 (0.003) | 0.096 (0.013) | 0.000 |
| Right Entorhinal Cortex | 0.017 (0.003) | 0.087 (0.013) | 0.000 |
| Right Frontal Pole | 0.015 (0.002) | 0.073 (0.012) | 0.000 |
| Right Fusiform Gyrus | 0.022 (0.003) | 0.107 (0.014) | 0.000 |
| Right Inferior Parietal Lobule | 0.022 (0.003) | 0.105 (0.014) | 0.000 |
| Right Inferior Temporal Gyrus | 0.027 (0.003) | 0.134 (0.014) | 0.000 |
| Right Insular Cortex | 0.014 (0.003) | 0.067 (0.012) | 0.000 |
| Right Isthmus of the Cingulate Cortex | 0.009 (0.002) | 0.046 (0.012) | 0.000 |
| Right Lateral Occipital Cortex | 0.032 (0.003) | 0.157 (0.015) | 0.000 |
| Right Lateral Orbitofrontal Cortex | 0.027 (0.003) | 0.131 (0.014) | 0.000 |
| Right Lingual Gyrus | 0.013 (0.003) | 0.064 (0.013) | 0.000 |
| Right Medial Orbitofrontal Cortex | 0.015 (0.003) | 0.074 (0.012) | 0.000 |
| Right Middle Temporal Gyrus | 0.035 (0.003) | 0.174 (0.015) | 0.000 |
| Right Paracentral Lobule | 0.015 (0.003) | 0.073 (0.013) | 0.000 |
| Right Parahippocampal Gyrus | 0.014 (0.003) | 0.068 (0.013) | 0.000 |
| Right Pars Opercularis | 0.015 (0.002) | 0.073 (0.012) | 0.000 |
| Right Pars Orbitalis | 0.022 (0.003) | 0.111 (0.013) | 0.000 |
| Right Pars Triangularis | 0.009 (0.002) | 0.046 (0.012) | 0.000 |
| Right Pericalcarine Cortex | 0.008 (0.003) | 0.040 (0.013) | 0.002 |
| Right Postcentral Gyrus | 0.023 (0.003) | 0.115 (0.014) | 0.000 |
| Right Posterior Cingulate Cortex | 0.014 (0.003) | 0.068 (0.013) | 0.000 |
| Right Precentral Gyrus | 0.025 (0.003) | 0.128 (0.014) | 0.000 |
| Right Precuneus | 0.027 (0.003) | 0.136 (0.014) | 0.000 |
| Right Rostral Anterior Cingulate Cortex | 0.011 (0.002) | 0.058 (0.013) | 0.000 |
| Right Rostral Middle Frontal Gyrus | 0.019 (0.003) | 0.094 (0.014) | 0.000 |
| Right Superior Frontal Gyrus | 0.021 (0.003) | 0.105 (0.014) | 0.000 |
| Right Superior Parietal Lobule | 0.025 (0.003) | 0.126 (0.014) | 0.000 |
| Right Superior Temporal Gyrus | 0.020 (0.003) | 0.099 (0.014) | 0.000 |
| Right Supramarginal Gyrus | 0.015 (0.003) | 0.073 (0.013) | 0.000 |
| Right Temporal Pole | 0.006 (0.002) | 0.031 (0.012) | 0.017 |
| Right Transverse Temporal Gyrus (Heschl’s Gyrus) | 0.020 (0.003) | 0.097 (0.013) | 0.000 |

**Supplementary Table 2: Baseline associations between regional centile scores and COI 2.0 social and economic summary score**

Values shown are unstandardised and standardised beta coefficients with associated standard errors (SE) for each region. P values are false discovery rate (FDR) corrected to account for multiple comparisons. All analyses controlled for age and sex and included random intercepts for testing site and family ID (to account for sibling pairs).

| **Region** | **Unstandardised estimate (SE)** | **Standardised estimate (SE)** | ***p*_FDR_** |
| --- | --- | --- | --- |
| Left Banks of the Superior Temporal Sulcus | 0.016 (0.002) | 0.081 (0.012) | 0.000 |
| Left Caudal Anterior Cingulate Cortex | 0.019 (0.003) | 0.092 (0.012) | 0.000 |
| Left Caudal Middle Frontal Gyrus | 0.024 (0.003) | 0.118 (0.012) | 0.000 |
| Left Cuneus | 0.025 (0.003) | 0.124 (0.013) | 0.000 |
| Left Entorhinal Cortex | 0.025 (0.002) | 0.129 (0.013) | 0.000 |
| Left Frontal Pole | 0.017 (0.002) | 0.085 (0.012) | 0.000 |
| Left Fusiform Gyrus | 0.032 (0.003) | 0.158 (0.013) | 0.000 |
| Left Inferior Parietal Lobule | 0.016 (0.003) | 0.082 (0.013) | 0.000 |
| Left Inferior Temporal Gyrus | 0.036 (0.003) | 0.176 (0.013) | 0.000 |
| Left Insular Cortex | 0.024 (0.003) | 0.119 (0.013) | 0.000 |
| Left Isthmus of the Cingulate Cortex | 0.021 (0.003) | 0.104 (0.013) | 0.000 |
| Left Lateral Occipital Cortex | 0.035 (0.003) | 0.170 (0.013) | 0.000 |
| Left Lateral Orbitofrontal Cortex | 0.035 (0.003) | 0.173 (0.013) | 0.000 |
| Left Lingual Gyrus | 0.024 (0.003) | 0.122 (0.013) | 0.000 |
| Left Medial Orbitofrontal Cortex | 0.016 (0.002) | 0.079 (0.012) | 0.000 |
| Left Middle Temporal Gyrus | 0.035 (0.003) | 0.174 (0.013) | 0.000 |
| Left Paracentral Lobule | 0.022 (0.003) | 0.110 (0.013) | 0.000 |
| Left Parahippocampal Gyrus | 0.023 (0.003) | 0.113 (0.012) | 0.000 |
| Left Pars Opercularis | 0.018 (0.002) | 0.090 (0.012) | 0.000 |
| Left Pars Orbitalis | 0.024 (0.002) | 0.120 (0.012) | 0.000 |
| Left Pars Triangularis | 0.012 (0.002) | 0.059 (0.012) | 0.000 |
| Left Pericalcarine Cortex | 0.015 (0.002) | 0.077 (0.013) | 0.000 |
| Left Postcentral Gyrus | 0.037 (0.003) | 0.182 (0.013) | 0.000 |
| Left Posterior Cingulate Cortex | 0.016 (0.003) | 0.078 (0.012) | 0.000 |
| Left Precentral Gyrus | 0.037 (0.003) | 0.185 (0.013) | 0.000 |
| Left Precuneus | 0.034 (0.003) | 0.166 (0.013) | 0.000 |
| Left Rostral Anterior Cingulate Cortex | 0.021 (0.003) | 0.106 (0.012) | 0.000 |
| Left Rostral Middle Frontal Gyrus | 0.027 (0.003) | 0.133 (0.013) | 0.000 |
| Left Superior Frontal Gyrus | 0.027 (0.003) | 0.130 (0.013) | 0.000 |
| Left Superior Parietal Lobule | 0.031 (0.003) | 0.154 (0.013) | 0.000 |
| Left Superior Temporal Gyrus | 0.026 (0.003) | 0.130 (0.013) | 0.000 |
| Left Supramarginal Gyrus | 0.029 (0.003) | 0.142 (0.013) | 0.000 |
| Left Temporal Pole | 0.017 (0.002) | 0.086 (0.012) | 0.000 |
| Left Transverse Temporal Gyrus (Heschl’s Gyrus) | 0.021 (0.002) | 0.103 (0.012) | 0.000 |
| Right Banks of the Superior Temporal Sulcus | 0.020 (0.003) | 0.100 (0.013) | 0.000 |
| Right Caudal Anterior Cingulate Cortex | 0.015 (0.003) | 0.075 (0.012) | 0.000 |
| Right Caudal Middle Frontal Gyrus | 0.022 (0.002) | 0.109 (0.012) | 0.000 |
| Right Cuneus | 0.026 (0.003) | 0.128 (0.013) | 0.000 |
| Right Entorhinal Cortex | 0.025 (0.002) | 0.124 (0.012) | 0.000 |
| Right Frontal Pole | 0.022 (0.003) | 0.109 (0.013) | 0.000 |
| Right Fusiform Gyrus | 0.031 (0.003) | 0.152 (0.013) | 0.000 |
| Right Inferior Parietal Lobule | 0.025 (0.003) | 0.122 (0.013) | 0.000 |
| Right Inferior Temporal Gyrus | 0.034 (0.003) | 0.169 (0.013) | 0.000 |
| Right Insular Cortex | 0.020 (0.002) | 0.098 (0.012) | 0.000 |
| Right Isthmus of the Cingulate Cortex | 0.013 (0.002) | 0.067 (0.012) | 0.000 |
| Right Lateral Occipital Cortex | 0.039 (0.003) | 0.193 (0.013) | 0.000 |
| Right Lateral Orbitofrontal Cortex | 0.033 (0.003) | 0.161 (0.013) | 0.000 |
| Right Lingual Gyrus | 0.019 (0.003) | 0.094 (0.012) | 0.000 |
| Right Medial Orbitofrontal Cortex | 0.024 (0.003) | 0.117 (0.012) | 0.000 |
| Right Middle Temporal Gyrus | 0.040 (0.003) | 0.197 (0.013) | 0.000 |
| Right Paracentral Lobule | 0.018 (0.003) | 0.088 (0.013) | 0.000 |
| Right Parahippocampal Gyrus | 0.020 (0.003) | 0.098 (0.012) | 0.000 |
| Right Pars Opercularis | 0.020 (0.002) | 0.097 (0.012) | 0.000 |
| Right Pars Orbitalis | 0.027 (0.003) | 0.136 (0.013) | 0.000 |
| Right Pars Triangularis | 0.010 (0.002) | 0.049 (0.012) | 0.000 |
| Right Pericalcarine Cortex | 0.014 (0.003) | 0.067 (0.012) | 0.000 |
| Right Postcentral Gyrus | 0.031 (0.003) | 0.154 (0.013) | 0.000 |
| Right Posterior Cingulate Cortex | 0.020 (0.002) | 0.099 (0.012) | 0.000 |
| Right Precentral Gyrus | 0.033 (0.003) | 0.167 (0.013) | 0.000 |
| Right Precuneus | 0.033 (0.003) | 0.166 (0.013) | 0.000 |
| Right Rostral Anterior Cingulate Cortex | 0.017 (0.002) | 0.086 (0.012) | 0.000 |
| Right Rostral Middle Frontal Gyrus | 0.025 (0.003) | 0.126 (0.013) | 0.000 |
| Right Superior Frontal Gyrus | 0.030 (0.003) | 0.148 (0.013) | 0.000 |
| Right Superior Parietal Lobule | 0.031 (0.003) | 0.155 (0.013) | 0.000 |
| Right Superior Temporal Gyrus | 0.028 (0.003) | 0.140 (0.013) | 0.000 |
| Right Supramarginal Gyrus | 0.021 (0.003) | 0.101 (0.013) | 0.000 |
| Right Temporal Pole | 0.012 (0.002) | 0.061 (0.012) | 0.000 |
| Right Transverse Temporal Gyrus (Heschl’s Gyrus) | 0.025 (0.003) | 0.122 (0.013) | 0.000 |

**Supplementary Table 3: Baseline associations between regional centile scores and COI 2.0 education summary score**

Values shown are unstandardised and standardised beta coefficients with associated standard errors (SE) for each region. P values are false discovery rate (FDR) corrected to account for multiple comparisons. All analyses controlled for age and sex and included random intercepts for testing site and family ID (to account for sibling pairs).

| **Region** | **Unstandardised estimate (SE)** | **Standardised estimate (SE)** | ***p*_FDR_** |
| --- | --- | --- | --- |
| Left Banks of the Superior Temporal Sulcus | 0.017 (0.002) | 0.085 (0.012) | 0.000 |
| Left Caudal Anterior Cingulate Cortex | 0.016 (0.003) | 0.077 (0.012) | 0.000 |
| Left Caudal Middle Frontal Gyrus | 0.021 (0.003) | 0.101 (0.012) | 0.000 |
| Left Cuneus | 0.024 (0.003) | 0.117 (0.013) | 0.000 |
| Left Entorhinal Cortex | 0.024 (0.002) | 0.120 (0.012) | 0.000 |
| Left Frontal Pole | 0.018 (0.002) | 0.090 (0.012) | 0.000 |
| Left Fusiform Gyrus | 0.032 (0.003) | 0.155 (0.012) | 0.000 |
| Left Inferior Parietal Lobule | 0.017 (0.003) | 0.084 (0.012) | 0.000 |
| Left Inferior Temporal Gyrus | 0.034 (0.003) | 0.166 (0.012) | 0.000 |
| Left Insular Cortex | 0.025 (0.003) | 0.120 (0.012) | 0.000 |
| Left Isthmus of the Cingulate Cortex | 0.021 (0.003) | 0.105 (0.012) | 0.000 |
| Left Lateral Occipital Cortex | 0.032 (0.003) | 0.157 (0.013) | 0.000 |
| Left Lateral Orbitofrontal Cortex | 0.034 (0.003) | 0.163 (0.013) | 0.000 |
| Left Lingual Gyrus | 0.024 (0.003) | 0.121 (0.013) | 0.000 |
| Left Medial Orbitofrontal Cortex | 0.018 (0.003) | 0.087 (0.012) | 0.000 |
| Left Middle Temporal Gyrus | 0.033 (0.003) | 0.161 (0.013) | 0.000 |
| Left Paracentral Lobule | 0.018 (0.003) | 0.085 (0.012) | 0.000 |
| Left Parahippocampal Gyrus | 0.024 (0.003) | 0.115 (0.012) | 0.000 |
| Left Pars Opercularis | 0.019 (0.002) | 0.094 (0.012) | 0.000 |
| Left Pars Orbitalis | 0.024 (0.003) | 0.119 (0.012) | 0.000 |
| Left Pars Triangularis | 0.014 (0.003) | 0.069 (0.012) | 0.000 |
| Left Pericalcarine Cortex | 0.016 (0.003) | 0.082 (0.013) | 0.000 |
| Left Postcentral Gyrus | 0.034 (0.003) | 0.164 (0.013) | 0.000 |
| Left Posterior Cingulate Cortex | 0.017 (0.003) | 0.085 (0.012) | 0.000 |
| Left Precentral Gyrus | 0.031 (0.003) | 0.153 (0.013) | 0.000 |
| Left Precuneus | 0.030 (0.003) | 0.146 (0.012) | 0.000 |
| Left Rostral Anterior Cingulate Cortex | 0.018 (0.003) | 0.088 (0.012) | 0.000 |
| Left Rostral Middle Frontal Gyrus | 0.027 (0.003) | 0.134 (0.012) | 0.000 |
| Left Superior Frontal Gyrus | 0.024 (0.003) | 0.116 (0.013) | 0.000 |
| Left Superior Parietal Lobule | 0.029 (0.003) | 0.143 (0.012) | 0.000 |
| Left Superior Temporal Gyrus | 0.025 (0.003) | 0.123 (0.013) | 0.000 |
| Left Supramarginal Gyrus | 0.026 (0.003) | 0.125 (0.013) | 0.000 |
| Left Temporal Pole | 0.017 (0.003) | 0.084 (0.012) | 0.000 |
| Left Transverse Temporal Gyrus (Heschl’s Gyrus) | 0.020 (0.003) | 0.099 (0.012) | 0.000 |
| Right Banks of the Superior Temporal Sulcus | 0.018 (0.003) | 0.089 (0.012) | 0.000 |
| Right Caudal Anterior Cingulate Cortex | 0.014 (0.003) | 0.068 (0.012) | 0.000 |
| Right Caudal Middle Frontal Gyrus | 0.021 (0.003) | 0.104 (0.012) | 0.000 |
| Right Cuneus | 0.025 (0.003) | 0.124 (0.012) | 0.000 |
| Right Entorhinal Cortex | 0.021 (0.002) | 0.105 (0.012) | 0.000 |
| Right Frontal Pole | 0.024 (0.002) | 0.118 (0.012) | 0.000 |
| Right Fusiform Gyrus | 0.032 (0.003) | 0.156 (0.012) | 0.000 |
| Right Inferior Parietal Lobule | 0.022 (0.003) | 0.107 (0.013) | 0.000 |
| Right Inferior Temporal Gyrus | 0.031 (0.003) | 0.150 (0.013) | 0.000 |
| Right Insular Cortex | 0.020 (0.003) | 0.098 (0.012) | 0.000 |
| Right Isthmus of the Cingulate Cortex | 0.013 (0.002) | 0.064 (0.012) | 0.000 |
| Right Lateral Occipital Cortex | 0.036 (0.003) | 0.176 (0.013) | 0.000 |
| Right Lateral Orbitofrontal Cortex | 0.030 (0.003) | 0.149 (0.013) | 0.000 |
| Right Lingual Gyrus | 0.021 (0.003) | 0.101 (0.012) | 0.000 |
| Right Medial Orbitofrontal Cortex | 0.025 (0.003) | 0.123 (0.012) | 0.000 |
| Right Middle Temporal Gyrus | 0.038 (0.003) | 0.184 (0.013) | 0.000 |
| Right Paracentral Lobule | 0.017 (0.003) | 0.081 (0.012) | 0.000 |
| Right Parahippocampal Gyrus | 0.023 (0.003) | 0.111 (0.012) | 0.000 |
| Right Pars Opercularis | 0.017 (0.003) | 0.082 (0.012) | 0.000 |
| Right Pars Orbitalis | 0.026 (0.003) | 0.126 (0.012) | 0.000 |
| Right Pars Triangularis | 0.010 (0.002) | 0.053 (0.012) | 0.000 |
| Right Pericalcarine Cortex | 0.014 (0.003) | 0.067 (0.012) | 0.000 |
| Right Postcentral Gyrus | 0.030 (0.003) | 0.145 (0.013) | 0.000 |
| Right Posterior Cingulate Cortex | 0.016 (0.002) | 0.082 (0.012) | 0.000 |
| Right Precentral Gyrus | 0.028 (0.003) | 0.140 (0.013) | 0.000 |
| Right Precuneus | 0.029 (0.003) | 0.144 (0.012) | 0.000 |
| Right Rostral Anterior Cingulate Cortex | 0.014 (0.002) | 0.071 (0.012) | 0.000 |
| Right Rostral Middle Frontal Gyrus | 0.023 (0.003) | 0.112 (0.013) | 0.000 |
| Right Superior Frontal Gyrus | 0.027 (0.003) | 0.130 (0.012) | 0.000 |
| Right Superior Parietal Lobule | 0.028 (0.003) | 0.139 (0.013) | 0.000 |
| Right Superior Temporal Gyrus | 0.027 (0.003) | 0.134 (0.012) | 0.000 |
| Right Supramarginal Gyrus | 0.022 (0.003) | 0.107 (0.012) | 0.000 |
| Right Temporal Pole | 0.012 (0.003) | 0.062 (0.012) | 0.000 |
| Right Transverse Temporal Gyrus (Heschl’s Gyrus) | 0.022 (0.003) | 0.109 (0.012) | 0.000 |

**Supplementary Table 4: Baseline associations between regional centile scores and micro-environment latent factor score**

Values shown are unstandardised and standardised beta coefficients with associated standard errors (SE) for each region. P values are false discovery rate (FDR) corrected to account for multiple comparisons. All analyses controlled for age and sex and included random intercepts for testing site and family ID (to account for sibling pairs).

| **Region** | **Unstandardised estimate (SE)** | **Standardised estimate (SE)** | ***p*_FDR_** |
| --- | --- | --- | --- |
| Left Banks of the Superior Temporal Sulcus | 0.035 (0.004) | 0.114 (0.012) | 0.000 |
| Left Caudal Anterior Cingulate Cortex | 0.033 (0.004) | 0.105 (0.012) | 0.000 |
| Left Caudal Middle Frontal Gyrus | 0.038 (0.004) | 0.121 (0.012) | 0.000 |
| Left Cuneus | 0.038 (0.004) | 0.120 (0.012) | 0.000 |
| Left Entorhinal Cortex | 0.045 (0.004) | 0.148 (0.012) | 0.000 |
| Left Frontal Pole | 0.031 (0.004) | 0.100 (0.012) | 0.000 |
| Left Fusiform Gyrus | 0.055 (0.004) | 0.176 (0.012) | 0.000 |
| Left Inferior Parietal Lobule | 0.037 (0.004) | 0.118 (0.012) | 0.000 |
| Left Inferior Temporal Gyrus | 0.057 (0.004) | 0.183 (0.012) | 0.000 |
| Left Insular Cortex | 0.040 (0.004) | 0.129 (0.012) | 0.000 |
| Left Isthmus of the Cingulate Cortex | 0.034 (0.004) | 0.109 (0.012) | 0.000 |
| Left Lateral Occipital Cortex | 0.048 (0.004) | 0.151 (0.012) | 0.000 |
| Left Lateral Orbitofrontal Cortex | 0.055 (0.004) | 0.175 (0.012) | 0.000 |
| Left Lingual Gyrus | 0.038 (0.004) | 0.123 (0.012) | 0.000 |
| Left Medial Orbitofrontal Cortex | 0.035 (0.004) | 0.113 (0.012) | 0.000 |
| Left Middle Temporal Gyrus | 0.060 (0.004) | 0.192 (0.012) | 0.000 |
| Left Paracentral Lobule | 0.045 (0.004) | 0.143 (0.012) | 0.000 |
| Left Parahippocampal Gyrus | 0.040 (0.004) | 0.126 (0.012) | 0.000 |
| Left Pars Opercularis | 0.029 (0.004) | 0.094 (0.012) | 0.000 |
| Left Pars Orbitalis | 0.045 (0.004) | 0.143 (0.012) | 0.000 |
| Left Pars Triangularis | 0.023 (0.004) | 0.074 (0.012) | 0.000 |
| Left Pericalcarine Cortex | 0.026 (0.004) | 0.086 (0.012) | 0.000 |
| Left Postcentral Gyrus | 0.061 (0.004) | 0.196 (0.012) | 0.000 |
| Left Posterior Cingulate Cortex | 0.034 (0.004) | 0.107 (0.012) | 0.000 |
| Left Precentral Gyrus | 0.061 (0.004) | 0.195 (0.012) | 0.000 |
| Left Precuneus | 0.051 (0.004) | 0.164 (0.012) | 0.000 |
| Left Rostral Anterior Cingulate Cortex | 0.041 (0.004) | 0.132 (0.012) | 0.000 |
| Left Rostral Middle Frontal Gyrus | 0.047 (0.004) | 0.150 (0.012) | 0.000 |
| Left Superior Frontal Gyrus | 0.049 (0.004) | 0.155 (0.012) | 0.000 |
| Left Superior Parietal Lobule | 0.050 (0.004) | 0.160 (0.012) | 0.000 |
| Left Superior Temporal Gyrus | 0.046 (0.004) | 0.146 (0.012) | 0.000 |
| Left Supramarginal Gyrus | 0.046 (0.004) | 0.149 (0.012) | 0.000 |
| Left Temporal Pole | 0.031 (0.004) | 0.100 (0.012) | 0.000 |
| Left Transverse Temporal Gyrus (Heschl’s Gyrus) | 0.033 (0.004) | 0.106 (0.012) | 0.000 |
| Right Banks of the Superior Temporal Sulcus | 0.038 (0.004) | 0.122 (0.012) | 0.000 |
| Right Caudal Anterior Cingulate Cortex | 0.027 (0.004) | 0.086 (0.012) | 0.000 |
| Right Caudal Middle Frontal Gyrus | 0.045 (0.004) | 0.146 (0.012) | 0.000 |
| Right Cuneus | 0.033 (0.004) | 0.105 (0.012) | 0.000 |
| Right Entorhinal Cortex | 0.041 (0.004) | 0.132 (0.012) | 0.000 |
| Right Frontal Pole | 0.036 (0.004) | 0.117 (0.012) | 0.000 |
| Right Fusiform Gyrus | 0.055 (0.004) | 0.176 (0.012) | 0.000 |
| Right Inferior Parietal Lobule | 0.044 (0.004) | 0.140 (0.012) | 0.000 |
| Right Inferior Temporal Gyrus | 0.057 (0.004) | 0.185 (0.012) | 0.000 |
| Right Insular Cortex | 0.038 (0.004) | 0.122 (0.012) | 0.000 |
| Right Isthmus of the Cingulate Cortex | 0.030 (0.004) | 0.096 (0.012) | 0.000 |
| Right Lateral Occipital Cortex | 0.055 (0.004) | 0.177 (0.012) | 0.000 |
| Right Lateral Orbitofrontal Cortex | 0.052 (0.004) | 0.168 (0.012) | 0.000 |
| Right Lingual Gyrus | 0.035 (0.004) | 0.111 (0.012) | 0.000 |
| Right Medial Orbitofrontal Cortex | 0.045 (0.004) | 0.143 (0.012) | 0.000 |
| Right Middle Temporal Gyrus | 0.066 (0.004) | 0.209 (0.012) | 0.000 |
| Right Paracentral Lobule | 0.032 (0.004) | 0.103 (0.012) | 0.000 |
| Right Parahippocampal Gyrus | 0.038 (0.004) | 0.123 (0.012) | 0.000 |
| Right Pars Opercularis | 0.029 (0.004) | 0.094 (0.012) | 0.000 |
| Right Pars Orbitalis | 0.044 (0.004) | 0.140 (0.012) | 0.000 |
| Right Pars Triangularis | 0.019 (0.004) | 0.064 (0.012) | 0.000 |
| Right Pericalcarine Cortex | 0.024 (0.004) | 0.077 (0.012) | 0.000 |
| Right Postcentral Gyrus | 0.055 (0.004) | 0.175 (0.012) | 0.000 |
| Right Posterior Cingulate Cortex | 0.033 (0.004) | 0.108 (0.012) | 0.000 |
| Right Precentral Gyrus | 0.059 (0.004) | 0.192 (0.012) | 0.000 |
| Right Precuneus | 0.050 (0.004) | 0.162 (0.012) | 0.000 |
| Right Rostral Anterior Cingulate Cortex | 0.032 (0.004) | 0.104 (0.012) | 0.000 |
| Right Rostral Middle Frontal Gyrus | 0.042 (0.004) | 0.136 (0.012) | 0.000 |
| Right Superior Frontal Gyrus | 0.052 (0.004) | 0.166 (0.012) | 0.000 |
| Right Superior Parietal Lobule | 0.046 (0.004) | 0.149 (0.012) | 0.000 |
| Right Superior Temporal Gyrus | 0.045 (0.004) | 0.146 (0.012) | 0.000 |
| Right Supramarginal Gyrus | 0.041 (0.004) | 0.131 (0.012) | 0.000 |
| Right Temporal Pole | 0.022 (0.004) | 0.073 (0.012) | 0.000 |
| Right Transverse Temporal Gyrus (Heschl’s Gyrus) | 0.034 (0.004) | 0.109 (0.012) | 0.000 |

**Supplementary Table 5: Baseline associations between regional centile scores and CBCL externalising traits**

Values shown are unstandardised and standardised beta coefficients with associated standard errors (SE) for each region. P values are false discovery rate (FDR) corrected to account for multiple comparisons. All analyses controlled for age and sex and included random intercepts for testing site and family ID (to account for sibling pairs).

| **Region** | **Unstandardised estimate (SE)** | **Standardised estimate (SE)** | ***p*_FDR_** |
| --- | --- | --- | --- |
| Left Banks of the Superior Temporal Sulcus | -0.752 (0.222) | -0.040 (0.012) | 0.001 |
| Left Caudal Anterior Cingulate Cortex | -0.704 (0.218) | -0.038 (0.012) | 0.002 |
| Left Caudal Middle Frontal Gyrus | -0.376 (0.220) | -0.049 (0.012) | 0.118 |
| Left Cuneus | -0.401 (0.224) | -0.010 (0.012) | 0.101 |
| Left Entorhinal Cortex | -0.948 (0.227) | -0.019 (0.012) | 0.000 |
| Left Frontal Pole | -0.514 (0.223) | -0.027 (0.012) | 0.030 |
| Left Fusiform Gyrus | -1.181 (0.222) | -0.064 (0.012) | 0.000 |
| Left Inferior Parietal Lobule | -1.222 (0.225) | -0.065 (0.012) | 0.000 |
| Left Inferior Temporal Gyrus | -1.303 (0.223) | -0.070 (0.012) | 0.000 |
| Left Insular Cortex | -1.121 (0.225) | -0.060 (0.012) | 0.000 |
| Left Isthmus of the Cingulate Cortex | -0.969 (0.224) | -0.051 (0.012) | 0.000 |
| Left Lateral Occipital Cortex | -0.811 (0.222) | -0.044 (0.012) | 0.000 |
| Left Lateral Orbitofrontal Cortex | -1.036 (0.223) | -0.056 (0.012) | 0.000 |
| Left Lingual Gyrus | -0.850 (0.230) | -0.044 (0.012) | 0.000 |
| Left Medial Orbitofrontal Cortex | -0.727 (0.224) | -0.039 (0.012) | 0.002 |
| Left Middle Temporal Gyrus | -1.279 (0.225) | -0.068 (0.012) | 0.000 |
| Left Paracentral Lobule | -0.980 (0.222) | -0.052 (0.012) | 0.000 |
| Left Parahippocampal Gyrus | -0.758 (0.217) | -0.041 (0.012) | 0.001 |
| Left Pars Opercularis | -0.459 (0.226) | -0.036 (0.012) | 0.059 |
| Left Pars Orbitalis | -0.665 (0.222) | -0.019 (0.012) | 0.004 |
| Left Pars Triangularis | -0.516 (0.222) | -0.027 (0.012) | 0.029 |
| Left Pericalcarine Cortex | -0.388 (0.231) | -0.047 (0.012) | 0.122 |
| Left Postcentral Gyrus | -0.871 (0.222) | -0.019 (0.012) | 0.000 |
| Left Posterior Cingulate Cortex | -1.067 (0.220) | -0.057 (0.012) | 0.000 |
| Left Precentral Gyrus | -1.280 (0.223) | -0.068 (0.012) | 0.000 |
| Left Precuneus | -0.895 (0.224) | -0.048 (0.012) | 0.000 |
| Left Rostral Anterior Cingulate Cortex | -0.899 (0.224) | -0.048 (0.012) | 0.000 |
| Left Rostral Middle Frontal Gyrus | -1.197 (0.225) | -0.064 (0.012) | 0.000 |
| Left Superior Frontal Gyrus | -1.252 (0.222) | -0.067 (0.012) | 0.000 |
| Left Superior Parietal Lobule | -0.931 (0.224) | -0.049 (0.012) | 0.000 |
| Left Superior Temporal Gyrus | -1.357 (0.224) | -0.072 (0.012) | 0.000 |
| Left Supramarginal Gyrus | -1.435 (0.222) | -0.077 (0.012) | 0.000 |
| Left Temporal Pole | -0.661 (0.223) | -0.035 (0.012) | 0.004 |
| Left Transverse Temporal Gyrus (Heschl’s Gyrus) | -0.759 (0.225) | -0.040 (0.012) | 0.001 |
| Right Banks of the Superior Temporal Sulcus | -0.767 (0.222) | -0.041 (0.012) | 0.001 |
| Right Caudal Anterior Cingulate Cortex | -0.735 (0.219) | -0.040 (0.012) | 0.001 |
| Right Caudal Middle Frontal Gyrus | -0.924 (0.223) | -0.049 (0.012) | 0.000 |
| Right Cuneus | -0.370 (0.224) | -0.054 (0.012) | 0.123 |
| Right Entorhinal Cortex | -1.025 (0.226) | -0.020 (0.012) | 0.000 |
| Right Frontal Pole | -0.641 (0.224) | -0.034 (0.012) | 0.006 |
| Right Fusiform Gyrus | -1.201 (0.222) | -0.064 (0.012) | 0.000 |
| Right Inferior Parietal Lobule | -1.310 (0.221) | -0.071 (0.012) | 0.000 |
| Right Inferior Temporal Gyrus | -1.233 (0.225) | -0.066 (0.012) | 0.000 |
| Right Insular Cortex | -0.948 (0.226) | -0.051 (0.012) | 0.000 |
| Right Isthmus of the Cingulate Cortex | -0.354 (0.225) | -0.050 (0.012) | 0.123 |
| Right Lateral Occipital Cortex | -0.929 (0.224) | -0.020 (0.012) | 0.000 |
| Right Lateral Orbitofrontal Cortex | -1.282 (0.225) | -0.068 (0.012) | 0.000 |
| Right Lingual Gyrus | -0.792 (0.224) | -0.042 (0.012) | 0.001 |
| Right Medial Orbitofrontal Cortex | -1.173 (0.224) | -0.063 (0.012) | 0.000 |
| Right Middle Temporal Gyrus | -1.310 (0.224) | -0.070 (0.012) | 0.000 |
| Right Paracentral Lobule | -1.076 (0.223) | -0.057 (0.012) | 0.000 |
| Right Parahippocampal Gyrus | -0.822 (0.222) | -0.044 (0.012) | 0.000 |
| Right Pars Opercularis | -0.364 (0.221) | -0.036 (0.012) | 0.123 |
| Right Pars Orbitalis | -0.669 (0.223) | -0.020 (0.012) | 0.004 |
| Right Pars Triangularis | -0.187 (0.228) | -0.063 (0.012) | 0.413 |
| Right Pericalcarine Cortex | -0.425 (0.224) | -0.022 (0.012) | 0.081 |
| Right Postcentral Gyrus | -1.170 (0.222) | -0.023 (0.012) | 0.000 |
| Right Posterior Cingulate Cortex | -1.105 (0.227) | -0.058 (0.012) | 0.000 |
| Right Precentral Gyrus | -1.623 (0.226) | -0.085 (0.012) | 0.000 |
| Right Precuneus | -0.778 (0.226) | -0.041 (0.012) | 0.001 |
| Right Rostral Anterior Cingulate Cortex | -0.838 (0.228) | -0.044 (0.012) | 0.000 |
| Right Rostral Middle Frontal Gyrus | -0.924 (0.226) | -0.049 (0.012) | 0.000 |
| Right Superior Frontal Gyrus | -1.177 (0.222) | -0.063 (0.012) | 0.000 |
| Right Superior Parietal Lobule | -0.702 (0.225) | -0.037 (0.012) | 0.003 |
| Right Superior Temporal Gyrus | -1.233 (0.226) | -0.066 (0.012) | 0.000 |
| Right Supramarginal Gyrus | -1.268 (0.222) | -0.068 (0.012) | 0.000 |
| Right Temporal Pole | -0.363 (0.225) | -0.031 (0.012) | 0.123 |
| Right Transverse Temporal Gyrus (Heschl’s Gyrus) | -0.589 (0.223) | -0.024 (0.012) | 0.012 |

**Supplementary Table 6: Baseline associations between regional centile scores and CBCL internalising traits**

Values shown are unstandardised and standardised beta coefficients with associated standard errors (SE) for each region. P values are false discovery rate (FDR) corrected to account for multiple comparisons. All analyses controlled for age and sex and included random intercepts for testing site and family ID (to account for sibling pairs).

| **Region** | **Unstandardised estimate (SE)** | **Standardised estimate (SE)** | ***p*_FDR_** |
| --- | --- | --- | --- |
| Left Banks of the Superior Temporal Sulcus | -0.272 (0.217) | -0.015 (0.012) | 0.285 |
| Left Caudal Anterior Cingulate Cortex | -0.411 (0.214) | -0.023 (0.012) | 0.096 |
| Left Caudal Middle Frontal Gyrus | -0.194 (0.215) | -0.011 (0.012) | 0.392 |
| Left Cuneus | 0.069 (0.221) | 0.004 (0.012) | 0.774 |
| Left Entorhinal Cortex | -0.605 (0.223) | -0.032 (0.012) | 0.026 |
| Left Frontal Pole | 0.166 (0.218) | 0.009 (0.012) | 0.470 |
| Left Fusiform Gyrus | -0.273 (0.218) | -0.015 (0.012) | 0.285 |
| Left Inferior Parietal Lobule | -0.627 (0.221) | -0.034 (0.012) | 0.026 |
| Left Inferior Temporal Gyrus | -0.457 (0.219) | -0.025 (0.012) | 0.067 |
| Left Insular Cortex | -0.294 (0.222) | -0.016 (0.012) | 0.279 |
| Left Isthmus of the Cingulate Cortex | -0.263 (0.220) | -0.014 (0.012) | 0.285 |
| Left Lateral Occipital Cortex | -0.219 (0.219) | -0.012 (0.012) | 0.343 |
| Left Lateral Orbitofrontal Cortex | -0.095 (0.220) | -0.005 (0.012) | 0.687 |
| Left Lingual Gyrus | -0.033 (0.227) | -0.002 (0.012) | 0.891 |
| Left Medial Orbitofrontal Cortex | -0.408 (0.220) | -0.022 (0.012) | 0.111 |
| Left Middle Temporal Gyrus | -0.335 (0.222) | -0.018 (0.012) | 0.217 |
| Left Paracentral Lobule | -0.601 (0.218) | -0.033 (0.012) | 0.026 |
| Left Parahippocampal Gyrus | -0.172 (0.213) | -0.010 (0.012) | 0.444 |
| Left Pars Opercularis | 0.053 (0.221) | 0.003 (0.012) | 0.824 |
| Left Pars Orbitalis | -0.173 (0.218) | -0.009 (0.012) | 0.451 |
| Left Pars Triangularis | -0.012 (0.218) | -0.001 (0.012) | 0.960 |
| Left Pericalcarine Cortex | 0.060 (0.228) | 0.003 (0.012) | 0.809 |
| Left Postcentral Gyrus | -0.570 (0.218) | -0.031 (0.012) | 0.026 |
| Left Posterior Cingulate Cortex | -0.323 (0.216) | -0.018 (0.012) | 0.220 |
| Left Precentral Gyrus | -0.338 (0.220) | -0.018 (0.012) | 0.208 |
| Left Precuneus | -0.289 (0.221) | -0.016 (0.012) | 0.280 |
| Left Rostral Anterior Cingulate Cortex | -0.326 (0.220) | -0.018 (0.012) | 0.225 |
| Left Rostral Middle Frontal Gyrus | -0.566 (0.222) | -0.031 (0.012) | 0.026 |
| Left Superior Frontal Gyrus | -0.272 (0.219) | -0.015 (0.012) | 0.285 |
| Left Superior Parietal Lobule | -0.301 (0.221) | -0.016 (0.012) | 0.270 |
| Left Superior Temporal Gyrus | -0.508 (0.221) | -0.028 (0.012) | 0.040 |
| Left Supramarginal Gyrus | -0.646 (0.218) | -0.035 (0.012) | 0.026 |
| Left Temporal Pole | -0.339 (0.219) | -0.018 (0.012) | 0.204 |
| Left Transverse Temporal Gyrus (Heschl’s Gyrus) | -0.162 (0.221) | -0.009 (0.012) | 0.485 |
| Right Banks of the Superior Temporal Sulcus | -0.220 (0.217) | -0.012 (0.012) | 0.340 |
| Right Caudal Anterior Cingulate Cortex | -0.325 (0.214) | -0.018 (0.012) | 0.213 |
| Right Caudal Middle Frontal Gyrus | -0.364 (0.218) | -0.020 (0.012) | 0.162 |
| Right Cuneus | 0.056 (0.220) | 0.003 (0.012) | 0.812 |
| Right Entorhinal Cortex | -0.475 (0.221) | -0.025 (0.012) | 0.059 |
| Right Frontal Pole | -0.362 (0.220) | -0.019 (0.012) | 0.169 |
| Right Fusiform Gyrus | -0.218 (0.219) | -0.012 (0.012) | 0.347 |
| Right Inferior Parietal Lobule | -0.317 (0.218) | -0.017 (0.012) | 0.234 |
| Right Inferior Temporal Gyrus | -0.411 (0.222) | -0.022 (0.012) | 0.112 |
| Right Insular Cortex | -0.263 (0.223) | -0.014 (0.012) | 0.285 |
| Right Isthmus of the Cingulate Cortex | 0.098 (0.220) | 0.005 (0.012) | 0.680 |
| Right Lateral Occipital Cortex | -0.003 (0.220) | 0.000 (0.012) | 0.988 |
| Right Lateral Orbitofrontal Cortex | -0.406 (0.222) | -0.022 (0.012) | 0.116 |
| Right Lingual Gyrus | -0.275 (0.221) | -0.015 (0.012) | 0.285 |
| Right Medial Orbitofrontal Cortex | -0.186 (0.220) | -0.010 (0.012) | 0.425 |
| Right Middle Temporal Gyrus | -0.439 (0.221) | -0.024 (0.012) | 0.084 |
| Right Paracentral Lobule | -0.674 (0.218) | -0.036 (0.012) | 0.026 |
| Right Parahippocampal Gyrus | -0.259 (0.219) | -0.014 (0.012) | 0.285 |
| Right Pars Opercularis | 0.158 (0.216) | 0.009 (0.012) | 0.485 |
| Right Pars Orbitalis | -0.388 (0.219) | -0.021 (0.012) | 0.132 |
| Right Pars Triangularis | 0.110 (0.223) | 0.006 (0.012) | 0.646 |
| Right Pericalcarine Cortex | 0.132 (0.221) | 0.007 (0.012) | 0.574 |
| Right Postcentral Gyrus | -0.629 (0.218) | -0.034 (0.012) | 0.026 |
| Right Posterior Cingulate Cortex | -0.390 (0.223) | -0.021 (0.012) | 0.136 |
| Right Precentral Gyrus | -0.616 (0.223) | -0.033 (0.012) | 0.026 |
| Right Precuneus | -0.264 (0.223) | -0.014 (0.012) | 0.285 |
| Right Rostral Anterior Cingulate Cortex | -0.219 (0.223) | -0.012 (0.012) | 0.352 |
| Right Rostral Middle Frontal Gyrus | -0.414 (0.223) | -0.022 (0.012) | 0.110 |
| Right Superior Frontal Gyrus | -0.443 (0.219) | -0.024 (0.012) | 0.079 |
| Right Superior Parietal Lobule | -0.211 (0.222) | -0.011 (0.012) | 0.367 |
| Right Superior Temporal Gyrus | -0.428 (0.223) | -0.023 (0.012) | 0.097 |
| Right Supramarginal Gyrus | -0.506 (0.218) | -0.028 (0.012) | 0.038 |
| Right Temporal Pole | -0.044 (0.221) | -0.002 (0.012) | 0.850 |
| Right Transverse Temporal Gyrus (Heschl’s Gyrus) | -0.266 (0.219) | -0.014 (0.012) | 0.285 |

**Supplementary Table 7: Baseline associations between regional centile scores and scores on the Flanker Inhibitory Control & Attention Task**

Values shown are unstandardised and standardised beta coefficients with associated standard errors (SE) for each region. P values are false discovery rate (FDR) corrected to account for multiple comparisons. All analyses controlled for age and sex and included random intercepts for testing site and family ID (to account for sibling pairs).

| **Region** | **Unstandardised estimate (SE)** | **Standardised estimate (SE)** | ***p*_FDR_** |
| --- | --- | --- | --- |
| Left Banks of the Superior Temporal Sulcus | 0.814 (0.358) | 0.027 (0.012) | 0.032 |
| Left Caudal Anterior Cingulate Cortex | 1.358 (0.351) | 0.045 (0.012) | 0.000 |
| Left Caudal Middle Frontal Gyrus | 1.307 (0.353) | 0.043 (0.012) | 0.000 |
| Left Cuneus | 1.269 (0.358) | 0.042 (0.012) | 0.001 |
| Left Entorhinal Cortex | 1.853 (0.365) | 0.060 (0.012) | 0.000 |
| Left Frontal Pole | 0.679 (0.360) | 0.022 (0.012) | 0.075 |
| Left Fusiform Gyrus | 2.232 (0.354) | 0.074 (0.012) | 0.000 |
| Left Inferior Parietal Lobule | 0.958 (0.361) | 0.031 (0.012) | 0.011 |
| Left Inferior Temporal Gyrus | 1.554 (0.358) | 0.051 (0.012) | 0.000 |
| Left Insular Cortex | 2.584 (0.360) | 0.085 (0.012) | 0.000 |
| Left Isthmus of the Cingulate Cortex | 1.148 (0.360) | 0.038 (0.012) | 0.002 |
| Left Lateral Occipital Cortex | 1.784 (0.354) | 0.059 (0.012) | 0.000 |
| Left Lateral Orbitofrontal Cortex | 1.936 (0.356) | 0.065 (0.012) | 0.000 |
| Left Lingual Gyrus | 1.158 (0.368) | 0.037 (0.012) | 0.002 |
| Left Medial Orbitofrontal Cortex | 0.977 (0.359) | 0.032 (0.012) | 0.009 |
| Left Middle Temporal Gyrus | 2.113 (0.360) | 0.070 (0.012) | 0.000 |
| Left Paracentral Lobule | 1.143 (0.357) | 0.038 (0.012) | 0.002 |
| Left Parahippocampal Gyrus | 1.659 (0.348) | 0.056 (0.012) | 0.000 |
| Left Pars Opercularis | 0.990 (0.363) | 0.032 (0.012) | 0.009 |
| Left Pars Orbitalis | 0.983 (0.357) | 0.032 (0.012) | 0.009 |
| Left Pars Triangularis | 1.354 (0.358) | 0.044 (0.012) | 0.000 |
| Left Pericalcarine Cortex | 0.686 (0.369) | 0.022 (0.012) | 0.078 |
| Left Postcentral Gyrus | 1.717 (0.355) | 0.057 (0.012) | 0.000 |
| Left Posterior Cingulate Cortex | 1.806 (0.353) | 0.060 (0.012) | 0.000 |
| Left Precentral Gyrus | 1.795 (0.358) | 0.059 (0.012) | 0.000 |
| Left Precuneus | 1.703 (0.357) | 0.056 (0.012) | 0.000 |
| Left Rostral Anterior Cingulate Cortex | 2.075 (0.359) | 0.068 (0.012) | 0.000 |
| Left Rostral Middle Frontal Gyrus | 1.161 (0.361) | 0.038 (0.012) | 0.002 |
| Left Superior Frontal Gyrus | 1.528 (0.355) | 0.051 (0.012) | 0.000 |
| Left Superior Parietal Lobule | 1.321 (0.359) | 0.043 (0.012) | 0.000 |
| Left Superior Temporal Gyrus | 1.686 (0.359) | 0.056 (0.012) | 0.000 |
| Left Supramarginal Gyrus | 1.711 (0.357) | 0.057 (0.012) | 0.000 |
| Left Temporal Pole | 1.204 (0.358) | 0.040 (0.012) | 0.001 |
| Left Transverse Temporal Gyrus (Heschl’s Gyrus) | 1.560 (0.361) | 0.051 (0.012) | 0.000 |
| Right Banks of the Superior Temporal Sulcus | 0.924 (0.356) | 0.030 (0.012) | 0.013 |
| Right Caudal Anterior Cingulate Cortex | 0.596 (0.353) | 0.020 (0.012) | 0.101 |
| Right Caudal Middle Frontal Gyrus | 1.494 (0.358) | 0.049 (0.012) | 0.000 |
| Right Cuneus | 0.694 (0.358) | 0.023 (0.012) | 0.067 |
| Right Entorhinal Cortex | 0.968 (0.363) | 0.031 (0.012) | 0.011 |
| Right Frontal Pole | 0.408 (0.361) | 0.013 (0.012) | 0.259 |
| Right Fusiform Gyrus | 1.818 (0.356) | 0.060 (0.012) | 0.000 |
| Right Inferior Parietal Lobule | 1.746 (0.354) | 0.058 (0.012) | 0.000 |
| Right Inferior Temporal Gyrus | 1.440 (0.361) | 0.047 (0.012) | 0.000 |
| Right Insular Cortex | 2.017 (0.361) | 0.066 (0.012) | 0.000 |
| Right Isthmus of the Cingulate Cortex | 1.110 (0.361) | 0.036 (0.012) | 0.003 |
| Right Lateral Occipital Cortex | 2.118 (0.357) | 0.070 (0.012) | 0.000 |
| Right Lateral Orbitofrontal Cortex | 2.014 (0.359) | 0.066 (0.012) | 0.000 |
| Right Lingual Gyrus | 1.318 (0.359) | 0.044 (0.012) | 0.000 |
| Right Medial Orbitofrontal Cortex | 1.458 (0.358) | 0.048 (0.012) | 0.000 |
| Right Middle Temporal Gyrus | 2.233 (0.357) | 0.074 (0.012) | 0.000 |
| Right Paracentral Lobule | 1.487 (0.358) | 0.049 (0.012) | 0.000 |
| Right Parahippocampal Gyrus | 1.539 (0.356) | 0.051 (0.012) | 0.000 |
| Right Pars Opercularis | 0.758 (0.355) | 0.025 (0.012) | 0.044 |
| Right Pars Orbitalis | 1.025 (0.359) | 0.034 (0.012) | 0.006 |
| Right Pars Triangularis | 0.621 (0.368) | 0.020 (0.012) | 0.101 |
| Right Pericalcarine Cortex | 0.313 (0.358) | 0.010 (0.012) | 0.383 |
| Right Postcentral Gyrus | 1.440 (0.355) | 0.048 (0.012) | 0.000 |
| Right Posterior Cingulate Cortex | 0.920 (0.364) | 0.030 (0.012) | 0.016 |
| Right Precentral Gyrus | 1.732 (0.363) | 0.056 (0.012) | 0.000 |
| Right Precuneus | 1.690 (0.360) | 0.055 (0.012) | 0.000 |
| Right Rostral Anterior Cingulate Cortex | 1.704 (0.367) | 0.055 (0.012) | 0.000 |
| Right Rostral Middle Frontal Gyrus | 1.351 (0.362) | 0.044 (0.012) | 0.000 |
| Right Superior Frontal Gyrus | 2.115 (0.355) | 0.070 (0.012) | 0.000 |
| Right Superior Parietal Lobule | 1.200 (0.361) | 0.039 (0.012) | 0.001 |
| Right Superior Temporal Gyrus | 1.851 (0.362) | 0.061 (0.012) | 0.000 |
| Right Supramarginal Gyrus | 1.225 (0.356) | 0.041 (0.012) | 0.001 |
| Right Temporal Pole | 0.862 (0.361) | 0.028 (0.012) | 0.024 |
| Right Transverse Temporal Gyrus (Heschl’s Gyrus) | 1.544 (0.357) | 0.051 (0.012) | 0.000 |

**Supplementary Table 8: Baseline associations between regional centile scores and scores on the Oral Reading Recognition Task**

Values shown are unstandardised and standardised beta coefficients with associated standard errors (SE) for each region. P values are false discovery rate (FDR) corrected to account for multiple comparisons. All analyses controlled for age and sex and included random intercepts for testing site and family ID (to account for sibling pairs).

| **Region** | **Unstandardised estimate (SE)** | **Standardised estimate (SE)** | ***p*_FDR_** |
| --- | --- | --- | --- |
| Left Banks of the Superior Temporal Sulcus | 2.069 (0.262) | 0.089 (0.011) | 0.000 |
| Left Caudal Anterior Cingulate Cortex | 1.838 (0.258) | 0.081 (0.011) | 0.000 |
| Left Caudal Middle Frontal Gyrus | 2.560 (0.259) | 0.112 (0.011) | 0.000 |
| Left Cuneus | 1.884 (0.266) | 0.082 (0.012) | 0.000 |
| Left Entorhinal Cortex | 2.532 (0.269) | 0.107 (0.011) | 0.000 |
| Left Frontal Pole | 1.936 (0.263) | 0.083 (0.011) | 0.000 |
| Left Fusiform Gyrus | 3.720 (0.261) | 0.163 (0.011) | 0.000 |
| Left Inferior Parietal Lobule | 2.614 (0.266) | 0.113 (0.011) | 0.000 |
| Left Inferior Temporal Gyrus | 3.414 (0.263) | 0.149 (0.011) | 0.000 |
| Left Insular Cortex | 3.086 (0.266) | 0.134 (0.012) | 0.000 |
| Left Isthmus of the Cingulate Cortex | 2.296 (0.265) | 0.100 (0.012) | 0.000 |
| Left Lateral Occipital Cortex | 3.215 (0.262) | 0.141 (0.012) | 0.000 |
| Left Lateral Orbitofrontal Cortex | 3.598 (0.263) | 0.159 (0.012) | 0.000 |
| Left Lingual Gyrus | 2.618 (0.273) | 0.111 (0.012) | 0.000 |
| Left Medial Orbitofrontal Cortex | 2.792 (0.264) | 0.122 (0.012) | 0.000 |
| Left Middle Temporal Gyrus | 3.687 (0.266) | 0.160 (0.012) | 0.000 |
| Left Paracentral Lobule | 2.239 (0.263) | 0.098 (0.011) | 0.000 |
| Left Parahippocampal Gyrus | 2.590 (0.256) | 0.115 (0.011) | 0.000 |
| Left Pars Opercularis | 2.053 (0.267) | 0.088 (0.011) | 0.000 |
| Left Pars Orbitalis | 2.602 (0.262) | 0.113 (0.011) | 0.000 |
| Left Pars Triangularis | 1.810 (0.263) | 0.078 (0.011) | 0.000 |
| Left Pericalcarine Cortex | 1.574 (0.276) | 0.067 (0.012) | 0.000 |
| Left Postcentral Gyrus | 3.520 (0.261) | 0.154 (0.011) | 0.000 |
| Left Posterior Cingulate Cortex | 2.599 (0.260) | 0.114 (0.011) | 0.000 |
| Left Precentral Gyrus | 3.672 (0.263) | 0.160 (0.011) | 0.000 |
| Left Precuneus | 2.817 (0.265) | 0.123 (0.012) | 0.000 |
| Left Rostral Anterior Cingulate Cortex | 2.699 (0.265) | 0.117 (0.012) | 0.000 |
| Left Rostral Middle Frontal Gyrus | 3.228 (0.266) | 0.140 (0.012) | 0.000 |
| Left Superior Frontal Gyrus | 3.208 (0.263) | 0.141 (0.012) | 0.000 |
| Left Superior Parietal Lobule | 2.595 (0.266) | 0.112 (0.012) | 0.000 |
| Left Superior Temporal Gyrus | 3.705 (0.264) | 0.161 (0.011) | 0.000 |
| Left Supramarginal Gyrus | 2.606 (0.263) | 0.114 (0.011) | 0.000 |
| Left Temporal Pole | 2.049 (0.264) | 0.089 (0.011) | 0.000 |
| Left Transverse Temporal Gyrus (Heschl’s Gyrus) | 2.670 (0.266) | 0.115 (0.011) | 0.000 |
| Right Banks of the Superior Temporal Sulcus | 2.530 (0.262) | 0.110 (0.011) | 0.000 |
| Right Caudal Anterior Cingulate Cortex | 1.692 (0.259) | 0.074 (0.011) | 0.000 |
| Right Caudal Middle Frontal Gyrus | 2.506 (0.263) | 0.109 (0.011) | 0.000 |
| Right Cuneus | 2.232 (0.265) | 0.098 (0.012) | 0.000 |
| Right Entorhinal Cortex | 1.882 (0.267) | 0.080 (0.011) | 0.000 |
| Right Frontal Pole | 2.191 (0.265) | 0.094 (0.011) | 0.000 |
| Right Fusiform Gyrus | 3.707 (0.262) | 0.162 (0.011) | 0.000 |
| Right Inferior Parietal Lobule | 3.014 (0.262) | 0.133 (0.011) | 0.000 |
| Right Inferior Temporal Gyrus | 3.264 (0.267) | 0.142 (0.012) | 0.000 |
| Right Insular Cortex | 3.063 (0.267) | 0.133 (0.012) | 0.000 |
| Right Isthmus of the Cingulate Cortex | 2.034 (0.266) | 0.088 (0.011) | 0.000 |
| Right Lateral Occipital Cortex | 3.681 (0.263) | 0.161 (0.012) | 0.000 |
| Right Lateral Orbitofrontal Cortex | 3.550 (0.265) | 0.154 (0.012) | 0.000 |
| Right Lingual Gyrus | 2.126 (0.266) | 0.093 (0.012) | 0.000 |
| Right Medial Orbitofrontal Cortex | 2.989 (0.264) | 0.131 (0.012) | 0.000 |
| Right Middle Temporal Gyrus | 4.110 (0.263) | 0.180 (0.012) | 0.000 |
| Right Paracentral Lobule | 2.105 (0.263) | 0.091 (0.011) | 0.000 |
| Right Parahippocampal Gyrus | 2.654 (0.263) | 0.116 (0.011) | 0.000 |
| Right Pars Opercularis | 1.684 (0.261) | 0.074 (0.011) | 0.000 |
| Right Pars Orbitalis | 2.715 (0.264) | 0.118 (0.011) | 0.000 |
| Right Pars Triangularis | 1.733 (0.270) | 0.073 (0.011) | 0.000 |
| Right Pericalcarine Cortex | 1.844 (0.267) | 0.081 (0.012) | 0.000 |
| Right Postcentral Gyrus | 3.067 (0.262) | 0.135 (0.012) | 0.000 |
| Right Posterior Cingulate Cortex | 2.572 (0.268) | 0.110 (0.011) | 0.000 |
| Right Precentral Gyrus | 3.423 (0.267) | 0.147 (0.011) | 0.000 |
| Right Precuneus | 3.053 (0.268) | 0.132 (0.012) | 0.000 |
| Right Rostral Anterior Cingulate Cortex | 2.543 (0.269) | 0.108 (0.011) | 0.000 |
| Right Rostral Middle Frontal Gyrus | 2.985 (0.267) | 0.130 (0.012) | 0.000 |
| Right Superior Frontal Gyrus | 3.367 (0.263) | 0.148 (0.012) | 0.000 |
| Right Superior Parietal Lobule | 2.768 (0.267) | 0.119 (0.012) | 0.000 |
| Right Superior Temporal Gyrus | 3.715 (0.267) | 0.161 (0.012) | 0.000 |
| Right Supramarginal Gyrus | 2.615 (0.263) | 0.114 (0.011) | 0.000 |
| Right Temporal Pole | 1.576 (0.267) | 0.068 (0.011) | 0.000 |
| Right Transverse Temporal Gyrus (Heschl’s Gyrus) | 2.448 (0.264) | 0.106 (0.011) | 0.000 |

**Supplementary Table 9: Baseline associations between regional centile scores and scores on the Picture Vocabulary Task**

Values shown are unstandardised and standardised beta coefficients with associated standard errors (SE) for each region. P values are false discovery rate (FDR) corrected to account for multiple comparisons. All analyses controlled for age and sex and included random intercepts for testing site and family ID (to account for sibling pairs).

| **Region** | **Unstandardised estimate (SE)** | **Standardised estimate (SE)** | ***p*_FDR_** |
| --- | --- | --- | --- |
| Left Banks of the Superior Temporal Sulcus | 0.845 (0.500) | 0.020 (0.012) | 0.095 |
| Left Caudal Anterior Cingulate Cortex | 0.856 (0.491) | 0.021 (0.012) | 0.085 |
| Left Caudal Middle Frontal Gyrus | 1.557 (0.494) | 0.037 (0.012) | 0.002 |
| Left Cuneus | 1.659 (0.501) | 0.040 (0.012) | 0.001 |
| Left Entorhinal Cortex | 1.992 (0.511) | 0.046 (0.012) | 0.000 |
| Left Frontal Pole | 0.030 (0.502) | 0.001 (0.012) | 0.953 |
| Left Fusiform Gyrus | 2.241 (0.497) | 0.054 (0.012) | 0.000 |
| Left Inferior Parietal Lobule | 1.347 (0.505) | 0.032 (0.012) | 0.008 |
| Left Inferior Temporal Gyrus | 2.930 (0.500) | 0.070 (0.012) | 0.000 |
| Left Insular Cortex | 2.078 (0.505) | 0.049 (0.012) | 0.000 |
| Left Isthmus of the Cingulate Cortex | 1.094 (0.503) | 0.026 (0.012) | 0.031 |
| Left Lateral Occipital Cortex | 1.826 (0.497) | 0.044 (0.012) | 0.000 |
| Left Lateral Orbitofrontal Cortex | 2.531 (0.499) | 0.061 (0.012) | 0.000 |
| Left Lingual Gyrus | 1.392 (0.515) | 0.032 (0.012) | 0.008 |
| Left Medial Orbitofrontal Cortex | 0.592 (0.502) | 0.014 (0.012) | 0.242 |
| Left Middle Temporal Gyrus | 2.535 (0.505) | 0.060 (0.012) | 0.000 |
| Left Paracentral Lobule | 1.623 (0.499) | 0.039 (0.012) | 0.001 |
| Left Parahippocampal Gyrus | 2.373 (0.487) | 0.058 (0.012) | 0.000 |
| Left Pars Opercularis | 0.804 (0.508) | 0.019 (0.012) | 0.117 |
| Left Pars Orbitalis | 1.930 (0.499) | 0.046 (0.012) | 0.000 |
| Left Pars Triangularis | 0.470 (0.500) | 0.011 (0.012) | 0.349 |
| Left Pericalcarine Cortex | 1.240 (0.517) | 0.029 (0.012) | 0.018 |
| Left Postcentral Gyrus | 2.126 (0.497) | 0.051 (0.012) | 0.000 |
| Left Posterior Cingulate Cortex | 1.263 (0.495) | 0.030 (0.012) | 0.012 |
| Left Precentral Gyrus | 2.407 (0.501) | 0.057 (0.012) | 0.000 |
| Left Precuneus | 1.752 (0.501) | 0.042 (0.012) | 0.001 |
| Left Rostral Anterior Cingulate Cortex | 1.863 (0.503) | 0.044 (0.012) | 0.000 |
| Left Rostral Middle Frontal Gyrus | 1.642 (0.506) | 0.039 (0.012) | 0.001 |
| Left Superior Frontal Gyrus | 1.753 (0.498) | 0.042 (0.012) | 0.001 |
| Left Superior Parietal Lobule | 2.355 (0.503) | 0.056 (0.012) | 0.000 |
| Left Superior Temporal Gyrus | 1.516 (0.503) | 0.036 (0.012) | 0.003 |
| Left Supramarginal Gyrus | 2.170 (0.499) | 0.052 (0.012) | 0.000 |
| Left Temporal Pole | 1.622 (0.500) | 0.038 (0.012) | 0.001 |
| Left Transverse Temporal Gyrus (Heschl’s Gyrus) | 0.843 (0.506) | 0.020 (0.012) | 0.099 |
| Right Banks of the Superior Temporal Sulcus | 1.102 (0.498) | 0.026 (0.012) | 0.028 |
| Right Caudal Anterior Cingulate Cortex | 1.494 (0.492) | 0.036 (0.012) | 0.003 |
| Right Caudal Middle Frontal Gyrus | 1.668 (0.500) | 0.040 (0.012) | 0.001 |
| Right Cuneus | 1.532 (0.501) | 0.037 (0.012) | 0.003 |
| Right Entorhinal Cortex | 1.521 (0.507) | 0.036 (0.012) | 0.003 |
| Right Frontal Pole | 0.306 (0.504) | 0.007 (0.012) | 0.546 |
| Right Fusiform Gyrus | 2.339 (0.499) | 0.056 (0.012) | 0.000 |
| Right Inferior Parietal Lobule | 1.747 (0.496) | 0.042 (0.012) | 0.001 |
| Right Inferior Temporal Gyrus | 2.946 (0.505) | 0.070 (0.012) | 0.000 |
| Right Insular Cortex | 2.050 (0.507) | 0.049 (0.012) | 0.000 |
| Right Isthmus of the Cingulate Cortex | 0.705 (0.505) | 0.017 (0.012) | 0.167 |
| Right Lateral Occipital Cortex | 1.940 (0.501) | 0.046 (0.012) | 0.000 |
| Right Lateral Orbitofrontal Cortex | 2.536 (0.503) | 0.060 (0.012) | 0.000 |
| Right Lingual Gyrus | 0.647 (0.503) | 0.015 (0.012) | 0.202 |
| Right Medial Orbitofrontal Cortex | 1.171 (0.502) | 0.028 (0.012) | 0.021 |
| Right Middle Temporal Gyrus | 2.350 (0.501) | 0.056 (0.012) | 0.000 |
| Right Paracentral Lobule | 1.235 (0.500) | 0.029 (0.012) | 0.015 |
| Right Parahippocampal Gyrus | 1.106 (0.499) | 0.026 (0.012) | 0.028 |
| Right Pars Opercularis | 0.514 (0.496) | 0.012 (0.012) | 0.304 |
| Right Pars Orbitalis | 1.498 (0.501) | 0.036 (0.012) | 0.003 |
| Right Pars Triangularis | 0.925 (0.513) | 0.021 (0.012) | 0.074 |
| Right Pericalcarine Cortex | 1.225 (0.502) | 0.029 (0.012) | 0.016 |
| Right Postcentral Gyrus | 1.980 (0.497) | 0.048 (0.012) | 0.000 |
| Right Posterior Cingulate Cortex | 1.218 (0.509) | 0.028 (0.012) | 0.018 |
| Right Precentral Gyrus | 2.079 (0.508) | 0.049 (0.012) | 0.000 |
| Right Precuneus | 1.967 (0.506) | 0.047 (0.012) | 0.000 |
| Right Rostral Anterior Cingulate Cortex | 2.287 (0.512) | 0.053 (0.012) | 0.000 |
| Right Rostral Middle Frontal Gyrus | 1.322 (0.508) | 0.031 (0.012) | 0.010 |
| Right Superior Frontal Gyrus | 2.130 (0.499) | 0.051 (0.012) | 0.000 |
| Right Superior Parietal Lobule | 1.827 (0.505) | 0.043 (0.012) | 0.000 |
| Right Superior Temporal Gyrus | 1.926 (0.507) | 0.046 (0.012) | 0.000 |
| Right Supramarginal Gyrus | 1.702 (0.498) | 0.041 (0.012) | 0.001 |
| Right Temporal Pole | 0.764 (0.504) | 0.018 (0.012) | 0.133 |
| Right Transverse Temporal Gyrus (Heschl’s Gyrus) | 1.014 (0.500) | 0.024 (0.012) | 0.045 |

**Supplementary Table 10: Baseline associations between regional centile scores and scores on the Picture Vocabulary Task**

Values shown are unstandardised and standardised beta coefficients with associated standard errors (SE) for each region. P values are false discovery rate (FDR) corrected to account for multiple comparisons. All analyses controlled for age and sex and included random intercepts for testing site and family ID (to account for sibling pairs).

| **Region** | **Unstandardised estimate (SE)** | **Standardised estimate (SE)** | ***p*_FDR_** |
| --- | --- | --- | --- |
| Left Banks of the Superior Temporal Sulcus | 2.484 (0.307) | 0.089 (0.011) | 0.000 |
| Left Caudal Anterior Cingulate Cortex | 2.135 (0.302) | 0.078 (0.011) | 0.000 |
| Left Caudal Middle Frontal Gyrus | 3.470 (0.303) | 0.127 (0.011) | 0.000 |
| Left Cuneus | 2.311 (0.311) | 0.084 (0.011) | 0.000 |
| Left Entorhinal Cortex | 3.421 (0.314) | 0.121 (0.011) | 0.000 |
| Left Frontal Pole | 2.068 (0.309) | 0.074 (0.011) | 0.000 |
| Left Fusiform Gyrus | 4.605 (0.304) | 0.168 (0.011) | 0.000 |
| Left Inferior Parietal Lobule | 2.800 (0.312) | 0.100 (0.011) | 0.000 |
| Left Inferior Temporal Gyrus | 4.311 (0.307) | 0.157 (0.011) | 0.000 |
| Left Insular Cortex | 3.819 (0.311) | 0.138 (0.011) | 0.000 |
| Left Isthmus of the Cingulate Cortex | 3.347 (0.310) | 0.121 (0.011) | 0.000 |
| Left Lateral Occipital Cortex | 3.800 (0.306) | 0.139 (0.011) | 0.000 |
| Left Lateral Orbitofrontal Cortex | 4.589 (0.307) | 0.168 (0.011) | 0.000 |
| Left Lingual Gyrus | 3.098 (0.319) | 0.110 (0.011) | 0.000 |
| Left Medial Orbitofrontal Cortex | 3.105 (0.310) | 0.113 (0.011) | 0.000 |
| Left Middle Temporal Gyrus | 4.635 (0.310) | 0.168 (0.011) | 0.000 |
| Left Paracentral Lobule | 2.907 (0.308) | 0.106 (0.011) | 0.000 |
| Left Parahippocampal Gyrus | 3.092 (0.300) | 0.115 (0.011) | 0.000 |
| Left Pars Opercularis | 2.621 (0.313) | 0.093 (0.011) | 0.000 |
| Left Pars Orbitalis | 3.430 (0.307) | 0.124 (0.011) | 0.000 |
| Left Pars Triangularis | 1.846 (0.308) | 0.067 (0.011) | 0.000 |
| Left Pericalcarine Cortex | 2.013 (0.322) | 0.071 (0.011) | 0.000 |
| Left Postcentral Gyrus | 4.137 (0.305) | 0.151 (0.011) | 0.000 |
| Left Posterior Cingulate Cortex | 3.062 (0.305) | 0.112 (0.011) | 0.000 |
| Left Precentral Gyrus | 4.512 (0.307) | 0.163 (0.011) | 0.000 |
| Left Precuneus | 3.682 (0.309) | 0.134 (0.011) | 0.000 |
| Left Rostral Anterior Cingulate Cortex | 3.306 (0.310) | 0.120 (0.011) | 0.000 |
| Left Rostral Middle Frontal Gyrus | 3.877 (0.311) | 0.140 (0.011) | 0.000 |
| Left Superior Frontal Gyrus | 4.104 (0.307) | 0.150 (0.011) | 0.000 |
| Left Superior Parietal Lobule | 3.325 (0.311) | 0.120 (0.011) | 0.000 |
| Left Superior Temporal Gyrus | 4.426 (0.309) | 0.160 (0.011) | 0.000 |
| Left Supramarginal Gyrus | 3.673 (0.307) | 0.133 (0.011) | 0.000 |
| Left Temporal Pole | 2.510 (0.309) | 0.091 (0.011) | 0.000 |
| Left Transverse Temporal Gyrus (Heschl’s Gyrus) | 3.311 (0.311) | 0.119 (0.011) | 0.000 |
| Right Banks of the Superior Temporal Sulcus | 3.278 (0.306) | 0.119 (0.011) | 0.000 |
| Right Caudal Anterior Cingulate Cortex | 2.309 (0.303) | 0.084 (0.011) | 0.000 |
| Right Caudal Middle Frontal Gyrus | 3.471 (0.307) | 0.125 (0.011) | 0.000 |
| Right Cuneus | 2.547 (0.311) | 0.093 (0.011) | 0.000 |
| Right Entorhinal Cortex | 2.393 (0.313) | 0.085 (0.011) | 0.000 |
| Right Frontal Pole | 2.802 (0.311) | 0.100 (0.011) | 0.000 |
| Right Fusiform Gyrus | 4.518 (0.306) | 0.164 (0.011) | 0.000 |
| Right Inferior Parietal Lobule | 3.565 (0.306) | 0.130 (0.011) | 0.000 |
| Right Inferior Temporal Gyrus | 4.376 (0.311) | 0.158 (0.011) | 0.000 |
| Right Insular Cortex | 3.479 (0.313) | 0.126 (0.011) | 0.000 |
| Right Isthmus of the Cingulate Cortex | 2.412 (0.312) | 0.086 (0.011) | 0.000 |
| Right Lateral Occipital Cortex | 4.312 (0.308) | 0.157 (0.011) | 0.000 |
| Right Lateral Orbitofrontal Cortex | 4.504 (0.309) | 0.163 (0.011) | 0.000 |
| Right Lingual Gyrus | 2.801 (0.311) | 0.102 (0.011) | 0.000 |
| Right Medial Orbitofrontal Cortex | 3.732 (0.309) | 0.136 (0.011) | 0.000 |
| Right Middle Temporal Gyrus | 5.306 (0.306) | 0.194 (0.011) | 0.000 |
| Right Paracentral Lobule | 2.508 (0.308) | 0.091 (0.011) | 0.000 |
| Right Parahippocampal Gyrus | 3.274 (0.308) | 0.119 (0.011) | 0.000 |
| Right Pars Opercularis | 2.598 (0.305) | 0.094 (0.011) | 0.000 |
| Right Pars Orbitalis | 3.452 (0.308) | 0.125 (0.011) | 0.000 |
| Right Pars Triangularis | 1.619 (0.316) | 0.057 (0.011) | 0.000 |
| Right Pericalcarine Cortex | 2.034 (0.313) | 0.074 (0.011) | 0.000 |
| Right Postcentral Gyrus | 3.725 (0.307) | 0.136 (0.011) | 0.000 |
| Right Posterior Cingulate Cortex | 3.011 (0.314) | 0.107 (0.011) | 0.000 |
| Right Precentral Gyrus | 4.267 (0.312) | 0.152 (0.011) | 0.000 |
| Right Precuneus | 4.070 (0.312) | 0.147 (0.011) | 0.000 |
| Right Rostral Anterior Cingulate Cortex | 3.063 (0.314) | 0.108 (0.011) | 0.000 |
| Right Rostral Middle Frontal Gyrus | 3.362 (0.313) | 0.121 (0.011) | 0.000 |
| Right Superior Frontal Gyrus | 4.266 (0.307) | 0.156 (0.011) | 0.000 |
| Right Superior Parietal Lobule | 3.634 (0.311) | 0.130 (0.011) | 0.000 |
| Right Superior Temporal Gyrus | 4.531 (0.312) | 0.164 (0.011) | 0.000 |
| Right Supramarginal Gyrus | 3.103 (0.308) | 0.113 (0.011) | 0.000 |
| Right Temporal Pole | 2.263 (0.312) | 0.081 (0.011) | 0.000 |
| Right Transverse Temporal Gyrus (Heschl’s Gyrus) | 3.174 (0.308) | 0.115 (0.011) | 0.000 |

**Supplementary Table 11: Baseline associations between regional centile scores and scores on the Pattern Comparison Processing Speed Task**

Values shown are unstandardised and standardised beta coefficients with associated standard errors (SE) for each region. P values are false discovery rate (FDR) corrected to account for multiple comparisons. All analyses controlled for age and sex and included random intercepts for testing site and family ID (to account for sibling pairs).

| **Region** | **Unstandardised estimate (SE)** | **Standardised estimate (SE)** | ***p*_FDR_** |
| --- | --- | --- | --- |
| Left Banks of the Superior Temporal Sulcus | -0.013 (0.591) | -0.000 (0.012) | 0.982 |
| Left Caudal Anterior Cingulate Cortex | 1.127 (0.580) | 0.023 (0.012) | 0.060 |
| Left Caudal Middle Frontal Gyrus | 1.029 (0.583) | 0.020 (0.012) | 0.087 |
| Left Cuneus | 1.376 (0.591) | 0.027 (0.012) | 0.025 |
| Left Entorhinal Cortex | 2.139 (0.603) | 0.041 (0.012) | 0.001 |
| Left Frontal Pole | -0.051 (0.594) | -0.001 (0.012) | 0.935 |
| Left Fusiform Gyrus | 1.410 (0.587) | 0.028 (0.012) | 0.020 |
| Left Inferior Parietal Lobule | -0.209 (0.596) | -0.004 (0.012) | 0.741 |
| Left Inferior Temporal Gyrus | 1.340 (0.591) | 0.027 (0.012) | 0.028 |
| Left Insular Cortex | 2.244 (0.596) | 0.044 (0.012) | 0.000 |
| Left Isthmus of the Cingulate Cortex | 0.920 (0.594) | 0.018 (0.012) | 0.135 |
| Left Lateral Occipital Cortex | 1.687 (0.586) | 0.034 (0.012) | 0.005 |
| Left Lateral Orbitofrontal Cortex | 1.186 (0.589) | 0.024 (0.012) | 0.052 |
| Left Lingual Gyrus | 1.169 (0.608) | 0.023 (0.012) | 0.063 |
| Left Medial Orbitofrontal Cortex | -0.095 (0.593) | -0.002 (0.012) | 0.882 |
| Left Middle Temporal Gyrus | 1.492 (0.596) | 0.029 (0.012) | 0.016 |
| Left Paracentral Lobule | 0.518 (0.589) | 0.010 (0.012) | 0.398 |
| Left Parahippocampal Gyrus | 1.150 (0.575) | 0.023 (0.012) | 0.054 |
| Left Pars Opercularis | 0.772 (0.599) | 0.015 (0.012) | 0.215 |
| Left Pars Orbitalis | 1.146 (0.589) | 0.023 (0.012) | 0.060 |
| Left Pars Triangularis | 0.634 (0.590) | 0.012 (0.012) | 0.299 |
| Left Pericalcarine Cortex | 0.917 (0.610) | 0.018 (0.012) | 0.145 |
| Left Postcentral Gyrus | 0.681 (0.587) | 0.014 (0.012) | 0.263 |
| Left Posterior Cingulate Cortex | 1.237 (0.584) | 0.025 (0.012) | 0.041 |
| Left Precentral Gyrus | 1.498 (0.592) | 0.030 (0.012) | 0.014 |
| Left Precuneus | 1.195 (0.591) | 0.024 (0.012) | 0.052 |
| Left Rostral Anterior Cingulate Cortex | 1.251 (0.594) | 0.025 (0.012) | 0.042 |
| Left Rostral Middle Frontal Gyrus | 0.074 (0.597) | 0.001 (0.012) | 0.909 |
| Left Superior Frontal Gyrus | 0.809 (0.588) | 0.016 (0.012) | 0.184 |
| Left Superior Parietal Lobule | 1.124 (0.594) | 0.022 (0.012) | 0.067 |
| Left Superior Temporal Gyrus | 1.307 (0.594) | 0.026 (0.012) | 0.034 |
| Left Supramarginal Gyrus | 1.368 (0.589) | 0.027 (0.012) | 0.025 |
| Left Temporal Pole | 1.118 (0.591) | 0.022 (0.012) | 0.067 |
| Left Transverse Temporal Gyrus (Heschl’s Gyrus) | 1.165 (0.597) | 0.023 (0.012) | 0.060 |
| Right Banks of the Superior Temporal Sulcus | -0.205 (0.588) | -0.004 (0.012) | 0.741 |
| Right Caudal Anterior Cingulate Cortex | 0.109 (0.582) | 0.002 (0.012) | 0.864 |
| Right Caudal Middle Frontal Gyrus | 1.763 (0.591) | 0.035 (0.012) | 0.004 |
| Right Cuneus | 1.201 (0.591) | 0.024 (0.012) | 0.050 |
| Right Entorhinal Cortex | 1.639 (0.599) | 0.032 (0.012) | 0.008 |
| Right Frontal Pole | -0.399 (0.595) | -0.008 (0.012) | 0.520 |
| Right Fusiform Gyrus | 1.143 (0.589) | 0.023 (0.012) | 0.061 |
| Right Inferior Parietal Lobule | 1.037 (0.585) | 0.021 (0.012) | 0.087 |
| Right Inferior Temporal Gyrus | 1.629 (0.596) | 0.032 (0.012) | 0.008 |
| Right Insular Cortex | 2.105 (0.598) | 0.042 (0.012) | 0.001 |
| Right Isthmus of the Cingulate Cortex | 0.624 (0.596) | 0.012 (0.012) | 0.311 |
| Right Lateral Occipital Cortex | 1.454 (0.591) | 0.029 (0.012) | 0.017 |
| Right Lateral Orbitofrontal Cortex | 1.545 (0.594) | 0.031 (0.012) | 0.012 |
| Right Lingual Gyrus | 0.992 (0.593) | 0.020 (0.012) | 0.105 |
| Right Medial Orbitofrontal Cortex | 1.021 (0.592) | 0.020 (0.012) | 0.095 |
| Right Middle Temporal Gyrus | 0.710 (0.591) | 0.014 (0.012) | 0.246 |
| Right Paracentral Lobule | 1.387 (0.591) | 0.027 (0.012) | 0.023 |
| Right Parahippocampal Gyrus | 1.144 (0.588) | 0.023 (0.012) | 0.060 |
| Right Pars Opercularis | 0.440 (0.586) | 0.009 (0.012) | 0.473 |
| Right Pars Orbitalis | 0.238 (0.592) | 0.005 (0.012) | 0.706 |
| Right Pars Triangularis | 0.422 (0.606) | 0.008 (0.012) | 0.505 |
| Right Pericalcarine Cortex | 1.087 (0.592) | 0.022 (0.012) | 0.075 |
| Right Postcentral Gyrus | 1.362 (0.587) | 0.027 (0.012) | 0.025 |
| Right Posterior Cingulate Cortex | 0.725 (0.601) | 0.014 (0.012) | 0.245 |
| Right Precentral Gyrus | 1.966 (0.599) | 0.038 (0.012) | 0.001 |
| Right Precuneus | 0.419 (0.597) | 0.008 (0.012) | 0.503 |
| Right Rostral Anterior Cingulate Cortex | 0.925 (0.605) | 0.018 (0.012) | 0.139 |
| Right Rostral Middle Frontal Gyrus | -0.322 (0.599) | -0.006 (0.012) | 0.609 |
| Right Superior Frontal Gyrus | 0.959 (0.589) | 0.019 (0.012) | 0.115 |
| Right Superior Parietal Lobule | 0.644 (0.596) | 0.013 (0.012) | 0.297 |
| Right Superior Temporal Gyrus | 0.744 (0.599) | 0.015 (0.012) | 0.232 |
| Right Supramarginal Gyrus | 0.645 (0.589) | 0.013 (0.012) | 0.291 |
| Right Temporal Pole | 1.440 (0.595) | 0.028 (0.012) | 0.019 |
| Right Transverse Temporal Gyrus (Heschl’s Gyrus) | 0.890 (0.591) | 0.018 (0.012) | 0.145 |

**Supplementary Table 12: Regions mediating the associations between COI 2.0 health/environment scores and externalising scores at baseline**

The table summarises the mediation analyses ran across regions that showed both environment-brain and brain-behaviour associations. The ACME (Average Causal Mediation Effect) estimate is reported, which reflects the indirect effect of the predictor on the outcome that is transmitted through the mediator, as well as proportion mediated values, which indicate the estimated fraction of the total effect explained by the mediator. Associated ACME confidence intervals and FDR-corrected p values are also reported. P values are false discovery rate (FDR) corrected to account for multiple comparisons. Analyses controlled for age and sex and included random intercepts for testing site and family ID (to account for sibling pairs).

| **Region** | **ACME estimate** | **ACME CI** | **Proportion mediated** | ***p*_FDR_** |
| --- | --- | --- | --- | --- |
| Left Banks of the Superior Temporal Sulcus | -0.010 | -0.017 - -0.004 | 0.024 | 0.000 |
| Left Caudal Anterior Cingulate Cortex | -0.007 | -0.013 - -0.002 | 0.018 | 0.000 |
| Left Entorhinal Cortex | -0.012 | -0.021 - -0.005 | 0.030 | 0.000 |
| Left Frontal Pole | -0.022 | -0.031 - -0.013 | 0.013 | 0.082 |
| Left Fusiform Gyrus | -0.014 | -0.022 - -0.007 | 0.054 | 0.000 |
| Left Inferior Parietal Lobule | -0.028 | -0.040 - -0.017 | 0.035 | 0.000 |
| Left Inferior Temporal Gyrus | -0.013 | -0.021 - -0.006 | 0.070 | 0.000 |
| Left Insular Cortex | -0.015 | -0.024 - -0.006 | 0.043 | 0.000 |
| Left Isthmus of the Cingulate Cortex | -0.020 | -0.030 - -0.011 | 0.032 | 0.000 |
| Left Lateral Occipital Cortex | -0.009 | -0.018 - -0.003 | 0.037 | 0.000 |
| Left Lateral Orbitofrontal Cortex | -0.008 | -0.015 - -0.003 | 0.051 | 0.000 |
| Left Lingual Gyrus | -0.026 | -0.038 - -0.015 | 0.023 | 0.002 |
| Left Medial Orbitofrontal Cortex | -0.010 | -0.017 - -0.004 | 0.020 | 0.000 |
| Left Middle Temporal Gyrus | -0.014 | -0.022 - -0.007 | 0.065 | 0.000 |
| Left Paracentral Lobule | -0.010 | -0.020 - -0.003 | 0.035 | 0.000 |
| Left Parahippocampal Gyrus | -0.004 | -0.008 - 0.000 | 0.024 | 0.002 |
| Left Pars Orbitalis | -0.020 | -0.031 - -0.009 | 0.025 | 0.006 |
| Left Pars Triangularis | -0.013 | -0.022 - -0.007 | 0.009 | 0.043 |
| Left Postcentral Gyrus | -0.029 | -0.041 - -0.018 | 0.050 | 0.000 |
| Left Posterior Cingulate Cortex | -0.016 | -0.026 - -0.007 | 0.033 | 0.000 |
| Left Precentral Gyrus | -0.010 | -0.017 - -0.004 | 0.073 | 0.000 |
| Left Precuneus | -0.019 | -0.028 - -0.010 | 0.040 | 0.000 |
| Left Rostral Anterior Cingulate Cortex | -0.022 | -0.032 - -0.012 | 0.026 | 0.002 |
| Left Rostral Middle Frontal Gyrus | -0.015 | -0.024 - -0.006 | 0.047 | 0.000 |
| Left Superior Frontal Gyrus | -0.018 | -0.028 - -0.010 | 0.054 | 0.000 |
| Left Superior Parietal Lobule | -0.021 | -0.030 - -0.012 | 0.037 | 0.000 |
| Left Superior Temporal Gyrus | -0.005 | -0.011 - 0.001 | 0.045 | 0.000 |
| Left Supramarginal Gyrus | -0.006 | -0.012 - -0.002 | 0.052 | 0.000 |
| Left Temporal Pole | -0.010 | -0.017 - -0.004 | 0.016 | 0.000 |
| Left Transverse Temporal Gyrus (Heschl’s Gyrus) | -0.017 | -0.026 - -0.010 | 0.025 | 0.002 |
| Right Banks of the Superior Temporal Sulcus | -0.010 | -0.017 - -0.004 | 0.024 | 0.000 |
| Right Caudal Anterior Cingulate Cortex | -0.008 | -0.015 - -0.003 | 0.021 | 0.000 |
| Right Caudal Middle Frontal Gyrus | -0.013 | -0.021 - -0.007 | 0.033 | 0.000 |
| Right Entorhinal Cortex | -0.015 | -0.025 - -0.008 | 0.038 | 0.000 |
| Right Frontal Pole | -0.021 | -0.031 - -0.013 | 0.020 | 0.023 |
| Right Fusiform Gyrus | -0.020 | -0.030 - -0.011 | 0.053 | 0.000 |
| Right Inferior Parietal Lobule | -0.023 | -0.034 - -0.013 | 0.050 | 0.000 |
| Right Inferior Temporal Gyrus | -0.018 | -0.029 - -0.008 | 0.057 | 0.000 |
| Right Insular Cortex | -0.025 | -0.037 - -0.014 | 0.032 | 0.000 |
| Right Lateral Occipital Cortex | -0.008 | -0.016 - -0.003 | 0.046 | 0.000 |
| Right Lateral Orbitofrontal Cortex | -0.017 | -0.027 - -0.009 | 0.063 | 0.000 |
| Right Lingual Gyrus | -0.030 | -0.043 - -0.018 | 0.021 | 0.004 |
| Right Medial Orbitofrontal Cortex | -0.010 | -0.017 - -0.004 | 0.043 | 0.000 |
| Right Middle Temporal Gyrus | -0.013 | -0.021 - -0.007 | 0.076 | 0.000 |
| Right Paracentral Lobule | -0.011 | -0.021 - -0.003 | 0.034 | 0.000 |
| Right Parahippocampal Gyrus | -0.022 | -0.034 - -0.013 | 0.024 | 0.000 |
| Right Pars Orbitalis | -0.014 | -0.022 - -0.007 | 0.029 | 0.004 |
| Right Postcentral Gyrus | -0.031 | -0.042 - -0.019 | 0.056 | 0.000 |
| Right Posterior Cingulate Cortex | -0.014 | -0.025 - -0.005 | 0.034 | 0.000 |
| Right Precentral Gyrus | -0.009 | -0.015 - -0.003 | 0.078 | 0.000 |
| Right Precuneus | -0.014 | -0.022 - -0.006 | 0.036 | 0.002 |
| Right Rostral Anterior Cingulate Cortex | -0.020 | -0.030 - -0.011 | 0.022 | 0.000 |
| Right Rostral Middle Frontal Gyrus | -0.011 | -0.021 - -0.003 | 0.034 | 0.000 |
| Right Superior Frontal Gyrus | -0.020 | -0.030 - -0.011 | 0.051 | 0.000 |
| Right Superior Parietal Lobule | -0.016 | -0.024 - -0.008 | 0.028 | 0.004 |
| Right Superior Temporal Gyrus | -0.008 | -0.016 - -0.001 | 0.049 | 0.000 |
| Right Supramarginal Gyrus | -0.008 | -0.016 - -0.001 | 0.040 | 0.000 |
| Right Transverse Temporal Gyrus (Heschl’s Gyrus) | -0.013 | -0.021 - -0.006 | 0.021 | 0.025 |

**Supplementary Table 13: Regions mediating the associations between COI 2.0 social/economic scores and externalising scores at baseline**

The table summarises the mediation analyses ran across regions that showed both environment-brain and brain-behaviour associations. The ACME (Average Causal Mediation Effect) estimate is reported, which reflects the indirect effect of the predictor on the outcome that is transmitted through the mediator, as well as proportion mediated values, which indicate the estimated fraction of the total effect explained by the mediator. Associated ACME confidence intervals and FDR-corrected p values are also reported. P values are false discovery rate (FDR) corrected to account for multiple comparisons. Analyses controlled for age and sex and included random intercepts for testing site and family ID (to account for sibling pairs).

| **Region** | **ACME estimate** | **ACME CI** | **Proportion mediated** | ***p*_FDR_** |
| --- | --- | --- | --- | --- |
| Left Banks of the Superior Temporal Sulcus | -0.011 | -0.020 - -0.003 | 0.029 | 0.000 |
| Left Caudal Anterior Cingulate Cortex | -0.013 | -0.022 - -0.004 | 0.032 | 0.003 |
| Left Entorhinal Cortex | -0.017 | -0.028 - -0.006 | 0.042 | 0.005 |
| Left Frontal Pole | -0.006 | -0.015 - 0.002 | 0.015 | 0.114 |
| Left Fusiform Gyrus | -0.031 | -0.046 - -0.018 | 0.078 | 0.000 |
| Left Inferior Parietal Lobule | -0.018 | -0.027 - -0.010 | 0.044 | 0.000 |
| Left Inferior Temporal Gyrus | -0.037 | -0.052 - -0.022 | 0.092 | 0.000 |
| Left Insular Cortex | -0.024 | -0.035 - -0.012 | 0.060 | 0.000 |
| Left Isthmus of the Cingulate Cortex | -0.017 | -0.028 - -0.008 | 0.042 | 0.000 |
| Left Lateral Occipital Cortex | -0.019 | -0.032 - -0.006 | 0.047 | 0.005 |
| Left Lateral Orbitofrontal Cortex | -0.029 | -0.045 - -0.014 | 0.073 | 0.003 |
| Left Lingual Gyrus | -0.013 | -0.024 - -0.002 | 0.033 | 0.013 |
| Left Medial Orbitofrontal Cortex | -0.011 | -0.020 - -0.004 | 0.029 | 0.000 |
| Left Middle Temporal Gyrus | -0.034 | -0.048 - -0.019 | 0.085 | 0.000 |
| Left Paracentral Lobule | -0.020 | -0.031 - -0.010 | 0.049 | 0.000 |
| Left Parahippocampal Gyrus | -0.014 | -0.025 - -0.005 | 0.036 | 0.003 |
| Left Pars Orbitalis | -0.012 | -0.023 - -0.001 | 0.030 | 0.036 |
| Left Pars Triangularis | -0.005 | -0.011 - 0.000 | 0.012 | 0.073 |
| Left Postcentral Gyrus | -0.025 | -0.041 - -0.011 | 0.063 | 0.003 |
| Left Posterior Cingulate Cortex | -0.015 | -0.024 - -0.007 | 0.038 | 0.000 |
| Left Precentral Gyrus | -0.038 | -0.053 - -0.024 | 0.096 | 0.000 |
| Left Precuneus | -0.021 | -0.035 - -0.007 | 0.052 | 0.000 |
| Left Rostral Anterior Cingulate Cortex | -0.017 | -0.026 - -0.008 | 0.042 | 0.000 |
| Left Rostral Middle Frontal Gyrus | -0.026 | -0.038 - -0.015 | 0.066 | 0.000 |
| Left Superior Frontal Gyrus | -0.028 | -0.040 - -0.017 | 0.069 | 0.000 |
| Left Superior Parietal Lobule | -0.020 | -0.033 - -0.007 | 0.051 | 0.000 |
| Left Superior Temporal Gyrus | -0.029 | -0.043 - -0.017 | 0.073 | 0.000 |
| Left Supramarginal Gyrus | -0.032 | -0.045 - -0.020 | 0.081 | 0.000 |
| Left Temporal Pole | -0.010 | -0.019 - -0.002 | 0.024 | 0.013 |
| Left Transverse Temporal Gyrus (Heschl’s Gyrus) | -0.014 | -0.025 - -0.005 | 0.035 | 0.005 |
| Right Banks of the Superior Temporal Sulcus | -0.012 | -0.021 - -0.004 | 0.031 | 0.005 |
| Right Caudal Anterior Cingulate Cortex | -0.011 | -0.018 - -0.004 | 0.027 | 0.000 |
| Right Caudal Middle Frontal Gyrus | -0.017 | -0.028 - -0.007 | 0.044 | 0.000 |
| Right Entorhinal Cortex | -0.020 | -0.032 - -0.008 | 0.050 | 0.000 |
| Right Frontal Pole | -0.010 | -0.020 - -0.001 | 0.024 | 0.036 |
| Right Fusiform Gyrus | -0.031 | -0.045 - -0.018 | 0.078 | 0.000 |
| Right Inferior Parietal Lobule | -0.026 | -0.037 - -0.015 | 0.065 | 0.000 |
| Right Inferior Temporal Gyrus | -0.032 | -0.047 - -0.018 | 0.080 | 0.000 |
| Right Insular Cortex | -0.018 | -0.027 - -0.009 | 0.044 | 0.000 |
| Right Lateral Occipital Cortex | -0.024 | -0.040 - -0.008 | 0.059 | 0.003 |
| Right Lateral Orbitofrontal Cortex | -0.034 | -0.049 - -0.020 | 0.086 | 0.000 |
| Right Lingual Gyrus | -0.011 | -0.021 - -0.003 | 0.028 | 0.009 |
| Right Medial Orbitofrontal Cortex | -0.025 | -0.037 - -0.014 | 0.064 | 0.000 |
| Right Middle Temporal Gyrus | -0.038 | -0.054 - -0.022 | 0.095 | 0.000 |
| Right Paracentral Lobule | -0.017 | -0.026 - -0.009 | 0.043 | 0.000 |
| Right Parahippocampal Gyrus | -0.013 | -0.022 - -0.005 | 0.032 | 0.009 |
| Right Pars Orbitalis | -0.013 | -0.025 - -0.002 | 0.034 | 0.009 |
| Right Postcentral Gyrus | -0.030 | -0.043 - -0.017 | 0.074 | 0.000 |
| Right Posterior Cingulate Cortex | -0.019 | -0.029 - -0.009 | 0.049 | 0.000 |
| Right Precentral Gyrus | -0.044 | -0.060 - -0.030 | 0.112 | 0.000 |
| Right Precuneus | -0.017 | -0.032 - -0.004 | 0.044 | 0.003 |
| Right Rostral Anterior Cingulate Cortex | -0.013 | -0.022 - -0.005 | 0.032 | 0.000 |
| Right Rostral Middle Frontal Gyrus | -0.019 | -0.031 - -0.008 | 0.047 | 0.000 |
| Right Superior Frontal Gyrus | -0.029 | -0.043 - -0.016 | 0.073 | 0.000 |
| Right Superior Parietal Lobule | -0.015 | -0.027 - -0.003 | 0.037 | 0.011 |
| Right Superior Temporal Gyrus | -0.028 | -0.041 - -0.016 | 0.069 | 0.000 |
| Right Supramarginal Gyrus | -0.022 | -0.032 - -0.013 | 0.055 | 0.000 |
| Right Transverse Temporal Gyrus (Heschl’s Gyrus) | -0.010 | -0.020 - 0.000 | 0.026 | 0.062 |

**Supplementary Table 14: Regions mediating the associations between COI 2.0 education scores and externalising scores at baseline**

The table summarises the mediation analyses ran across regions that showed both environment-brain and brain-behaviour associations. The ACME (Average Causal Mediation Effect) estimate is reported, which reflects the indirect effect of the predictor on the outcome that is transmitted through the mediator, as well as proportion mediated values, which indicate the estimated fraction of the total effect explained by the mediator. Associated ACME confidence intervals and FDR-corrected p values are also reported. P values are false discovery rate (FDR) corrected to account for multiple comparisons. Analyses controlled for age and sex and included random intercepts for testing site and family ID (to account for sibling pairs).

| **Region** | **ACME estimate** | **ACME CI** | **Proportion mediated** | ***p*_FDR_** |
| --- | --- | --- | --- | --- |
| Left Banks of the Superior Temporal Sulcus | -0.012 | -0.022 - -0.005 | 0.002 | 0.003 |
| Left Caudal Anterior Cingulate Cortex | -0.012 | -0.020 - -0.004 | 0.000 | 0.000 |
| Left Entorhinal Cortex | -0.018 | -0.029 - -0.006 | 0.004 | 0.005 |
| Left Frontal Pole | -0.007 | -0.016 - 0.002 | 0.116 | 0.116 |
| Left Fusiform Gyrus | -0.033 | -0.047 - -0.019 | 0.000 | 0.000 |
| Left Inferior Parietal Lobule | -0.020 | -0.030 - -0.012 | 0.000 | 0.000 |
| Left Inferior Temporal Gyrus | -0.039 | -0.055 - -0.024 | 0.000 | 0.000 |
| Left Insular Cortex | -0.025 | -0.039 - -0.015 | 0.000 | 0.000 |
| Left Isthmus of the Cingulate Cortex | -0.018 | -0.029 - -0.008 | 0.000 | 0.000 |
| Left Lateral Occipital Cortex | -0.020 | -0.033 - -0.007 | 0.002 | 0.003 |
| Left Lateral Orbitofrontal Cortex | -0.030 | -0.045 - -0.015 | 0.000 | 0.000 |
| Left Lingual Gyrus | -0.014 | -0.025 - -0.003 | 0.010 | 0.011 |
| Left Medial Orbitofrontal Cortex | -0.013 | -0.022 - -0.005 | 0.002 | 0.003 |
| Left Middle Temporal Gyrus | -0.035 | -0.052 - -0.021 | 0.000 | 0.000 |
| Left Paracentral Lobule | -0.017 | -0.027 - -0.009 | 0.000 | 0.000 |
| Left Parahippocampal Gyrus | -0.016 | -0.026 - -0.006 | 0.002 | 0.003 |
| Left Pars Orbitalis | -0.013 | -0.025 - -0.002 | 0.024 | 0.026 |
| Left Pars Triangularis | -0.006 | -0.013 - 0.000 | 0.078 | 0.079 |
| Left Postcentral Gyrus | -0.026 | -0.041 - -0.012 | 0.000 | 0.000 |
| Left Posterior Cingulate Cortex | -0.017 | -0.027 - -0.008 | 0.000 | 0.000 |
| Left Precentral Gyrus | -0.037 | -0.051 - -0.023 | 0.000 | 0.000 |
| Left Precuneus | -0.021 | -0.036 - -0.008 | 0.000 | 0.000 |
| Left Rostral Anterior Cingulate Cortex | -0.015 | -0.024 - -0.007 | 0.000 | 0.000 |
| Left Rostral Middle Frontal Gyrus | -0.029 | -0.043 - -0.015 | 0.000 | 0.000 |
| Left Superior Frontal Gyrus | -0.027 | -0.039 - -0.017 | 0.000 | 0.000 |
| Left Superior Parietal Lobule | -0.021 | -0.035 - -0.009 | 0.000 | 0.000 |
| Left Superior Temporal Gyrus | -0.030 | -0.042 - -0.018 | 0.000 | 0.000 |
| Left Supramarginal Gyrus | -0.032 | -0.046 - -0.020 | 0.000 | 0.000 |
| Left Temporal Pole | -0.010 | -0.018 - -0.002 | 0.012 | 0.013 |
| Left Transverse Temporal Gyrus (Heschl’s Gyrus) | -0.014 | -0.024 - -0.005 | 0.002 | 0.003 |
| Right Banks of the Superior Temporal Sulcus | -0.012 | -0.022 - -0.005 | 0.000 | 0.000 |
| Right Caudal Anterior Cingulate Cortex | -0.010 | -0.018 - -0.004 | 0.002 | 0.003 |
| Right Caudal Middle Frontal Gyrus | -0.017 | -0.027 - -0.009 | 0.000 | 0.000 |
| Right Entorhinal Cortex | -0.019 | -0.030 - -0.009 | 0.002 | 0.003 |
| Right Frontal Pole | -0.011 | -0.023 - 0.000 | 0.046 | 0.048 |
| Right Fusiform Gyrus | -0.034 | -0.049 - -0.020 | 0.000 | 0.000 |
| Right Inferior Parietal Lobule | -0.026 | -0.037 - -0.016 | 0.000 | 0.000 |
| Right Inferior Temporal Gyrus | -0.031 | -0.047 - -0.018 | 0.000 | 0.000 |
| Right Insular Cortex | -0.018 | -0.029 - -0.010 | 0.000 | 0.000 |
| Right Lateral Occipital Cortex | -0.024 | -0.041 - -0.009 | 0.002 | 0.003 |
| Right Lateral Orbitofrontal Cortex | -0.035 | -0.049 - -0.021 | 0.000 | 0.000 |
| Right Lingual Gyrus | -0.013 | -0.023 - -0.003 | 0.006 | 0.007 |
| Right Medial Orbitofrontal Cortex | -0.028 | -0.041 - -0.016 | 0.000 | 0.000 |
| Right Middle Temporal Gyrus | -0.040 | -0.058 - -0.024 | 0.000 | 0.000 |
| Right Paracentral Lobule | -0.017 | -0.026 - -0.009 | 0.000 | 0.000 |
| Right Parahippocampal Gyrus | -0.015 | -0.026 - -0.005 | 0.008 | 0.009 |
| Right Pars Orbitalis | -0.014 | -0.026 - -0.004 | 0.010 | 0.011 |
| Right Postcentral Gyrus | -0.031 | -0.046 - -0.018 | 0.000 | 0.000 |
| Right Posterior Cingulate Cortex | -0.017 | -0.027 - -0.009 | 0.000 | 0.000 |
| Right Precentral Gyrus | -0.042 | -0.058 - -0.028 | 0.000 | 0.000 |
| Right Precuneus | -0.018 | -0.032 - -0.005 | 0.002 | 0.003 |
| Right Rostral Anterior Cingulate Cortex | -0.011 | -0.019 - -0.005 | 0.000 | 0.000 |
| Right Rostral Middle Frontal Gyrus | -0.019 | -0.029 - -0.009 | 0.000 | 0.000 |
| Right Superior Frontal Gyrus | -0.028 | -0.042 - -0.016 | 0.000 | 0.000 |
| Right Superior Parietal Lobule | -0.016 | -0.029 - -0.004 | 0.006 | 0.007 |
| Right Superior Temporal Gyrus | -0.030 | -0.045 - -0.017 | 0.000 | 0.000 |
| Right Supramarginal Gyrus | -0.025 | -0.037 - -0.015 | 0.000 | 0.000 |
| Right Transverse Temporal Gyrus (Heschl’s Gyrus) | -0.010 | -0.020 - -0.001 | 0.042 | 0.044 |

**Supplementary Table 15: Regions mediating the associations between micro-environment scores and externalising scores at baseline**

The table summarises the mediation analyses ran across regions that showed both environment-brain and brain-behaviour associations. The ACME (Average Causal Mediation Effect) estimate is reported, which reflects the indirect effect of the predictor on the outcome that is transmitted through the mediator, as well as proportion mediated values, which indicate the estimated fraction of the total effect explained by the mediator. Associated ACME confidence intervals and FDR-corrected p values are also reported. P values are false discovery rate (FDR) corrected to account for multiple comparisons. Analyses controlled for age and sex and included random intercepts for testing site and family ID (to account for sibling pairs).

| **Region** | **ACME estimate** | **ACME CI** | **Proportion mediated** | ***p*_FDR_** |
| --- | --- | --- | --- | --- |
| Left Banks of the Superior Temporal Sulcus | -0.023 | -0.038 - -0.011 | -4.364 | 0.003 |
| Left Caudal Anterior Cingulate Cortex | -0.008 | -0.017 - -0.001 | -1.480 | 0.000 |
| Left Entorhinal Cortex | -0.008 | -0.017 - -0.001 | -1.527 | 0.005 |
| Left Frontal Pole | -0.006 | -0.014 - -0.001 | -1.040 | 0.116 |
| Left Fusiform Gyrus | -0.027 | -0.041 - -0.014 | -4.945 | 0.000 |
| Left Inferior Parietal Lobule | -0.046 | -0.066 - -0.028 | -8.600 | 0.000 |
| Left Inferior Temporal Gyrus | -0.032 | -0.048 - -0.019 | -6.032 | 0.000 |
| Left Insular Cortex | -0.049 | -0.069 - -0.029 | -9.048 | 0.000 |
| Left Isthmus of the Cingulate Cortex | -0.019 | -0.032 - -0.008 | -3.523 | 0.000 |
| Left Lateral Occipital Cortex | -0.005 | -0.012 - 0.002 | -0.866 | 0.003 |
| Left Lateral Orbitofrontal Cortex | -0.038 | -0.057 - -0.022 | -7.068 | 0.000 |
| Left Lingual Gyrus | -0.008 | -0.017 - -0.001 | -1.471 | 0.011 |
| Left Medial Orbitofrontal Cortex | -0.011 | -0.021 - -0.003 | -2.053 | 0.003 |
| Left Middle Temporal Gyrus | -0.059 | -0.082 - -0.038 | -10.947 | 0.000 |
| Left Paracentral Lobule | -0.027 | -0.041 - -0.014 | -4.939 | 0.000 |
| Left Parahippocampal Gyrus | 0.000 | -0.007 - 0.007 | 0.053 | 0.003 |
| Left Pars Orbitalis | -0.019 | -0.034 - -0.007 | -3.572 | 0.026 |
| Left Pars Triangularis | -0.012 | -0.025 - -0.002 | -2.266 | 0.079 |
| Left Postcentral Gyrus | -0.028 | -0.046 - -0.015 | -5.240 | 0.000 |
| Left Posterior Cingulate Cortex | -0.027 | -0.042 - -0.014 | -5.006 | 0.000 |
| Left Precentral Gyrus | -0.036 | -0.052 - -0.022 | -6.652 | 0.000 |
| Left Precuneus | -0.020 | -0.034 - -0.010 | -3.697 | 0.000 |
| Left Rostral Anterior Cingulate Cortex | -0.027 | -0.043 - -0.014 | -5.016 | 0.000 |
| Left Rostral Middle Frontal Gyrus | -0.025 | -0.040 - -0.013 | -4.714 | 0.000 |
| Left Superior Frontal Gyrus | -0.036 | -0.052 - -0.022 | -6.625 | 0.000 |
| Left Superior Parietal Lobule | -0.025 | -0.040 - -0.012 | -4.572 | 0.000 |
| Left Superior Temporal Gyrus | -0.024 | -0.038 - -0.012 | -4.367 | 0.000 |
| Left Supramarginal Gyrus | -0.044 | -0.064 - -0.029 | -8.144 | 0.000 |
| Left Temporal Pole | 0.003 | -0.003 - 0.009 | 0.583 | 0.013 |
| Left Transverse Temporal Gyrus (Heschl’s Gyrus) | -0.011 | -0.021 - -0.004 | -2.122 | 0.003 |
| Right Banks of the Superior Temporal Sulcus | -0.023 | -0.037 - -0.010 | -4.264 | 0.000 |
| Right Caudal Anterior Cingulate Cortex | -0.005 | -0.012 - 0.001 | -0.893 | 0.003 |
| Right Caudal Middle Frontal Gyrus | -0.020 | -0.034 - -0.010 | -3.783 | 0.000 |
| Right Entorhinal Cortex | -0.006 | -0.015 - 0.002 | -1.118 | 0.003 |
| Right Frontal Pole | -0.004 | -0.011 - 0.001 | -0.732 | 0.048 |
| Right Fusiform Gyrus | -0.029 | -0.044 - -0.016 | -5.305 | 0.000 |
| Right Inferior Parietal Lobule | -0.046 | -0.066 - -0.030 | -8.603 | 0.000 |
| Right Inferior Temporal Gyrus | -0.019 | -0.031 - -0.008 | -3.521 | 0.000 |
| Right Insular Cortex | -0.039 | -0.058 - -0.022 | -7.226 | 0.000 |
| Right Lateral Occipital Cortex | -0.009 | -0.018 - -0.001 | -1.675 | 0.003 |
| Right Lateral Orbitofrontal Cortex | -0.052 | -0.070 - -0.033 | -9.637 | 0.000 |
| Right Lingual Gyrus | -0.011 | -0.022 - -0.003 | -2.072 | 0.007 |
| Right Medial Orbitofrontal Cortex | -0.027 | -0.041 - -0.015 | -5.084 | 0.000 |
| Right Middle Temporal Gyrus | -0.064 | -0.090 - -0.045 | -11.918 | 0.000 |
| Right Paracentral Lobule | -0.026 | -0.040 - -0.014 | -4.832 | 0.000 |
| Right Parahippocampal Gyrus | -0.013 | -0.023 - -0.005 | -2.407 | 0.009 |
| Right Pars Orbitalis | -0.021 | -0.036 - -0.008 | -3.946 | 0.011 |
| Right Postcentral Gyrus | -0.035 | -0.052 - -0.020 | -6.458 | 0.000 |
| Right Posterior Cingulate Cortex | -0.024 | -0.039 - -0.013 | -4.492 | 0.000 |
| Right Precentral Gyrus | -0.031 | -0.048 - -0.017 | -5.709 | 0.000 |
| Right Precuneus | -0.017 | -0.029 - -0.007 | -3.108 | 0.003 |
| Right Rostral Anterior Cingulate Cortex | -0.019 | -0.032 - -0.009 | -3.470 | 0.000 |
| Right Rostral Middle Frontal Gyrus | -0.022 | -0.035 - -0.011 | -4.082 | 0.000 |
| Right Superior Frontal Gyrus | -0.034 | -0.050 - -0.020 | -6.297 | 0.000 |
| Right Superior Parietal Lobule | -0.019 | -0.033 - -0.007 | -3.437 | 0.007 |
| Right Superior Temporal Gyrus | -0.024 | -0.039 - -0.013 | -4.464 | 0.000 |
| Right Supramarginal Gyrus | -0.037 | -0.055 - -0.022 | -6.905 | 0.000 |
| Right Transverse Temporal Gyrus (Heschl’s Gyrus) | -0.016 | -0.030 - -0.004 | -3.031 | 0.044 |

**Supplementary Table 16: Regions mediating the associations between COI 2.0 health/environment scores and internalising scores at baseline**

The table summarises the mediation analyses ran across regions that showed both environment-brain and brain-behaviour associations. The ACME (Average Causal Mediation Effect) estimate is reported, which reflects the indirect effect of the predictor on the outcome that is transmitted through the mediator, as well as proportion mediated values, which indicate the estimated fraction of the total effect explained by the mediator. Associated ACME confidence intervals and FDR-corrected p values are also reported. P values are false discovery rate (FDR) corrected to account for multiple comparisons. Analyses controlled for age and sex and included random intercepts for testing site and family ID (to account for sibling pairs).

| **Region** | **ACME estimate** | **ACME CI** | **Proportion mediated** | ***p*_FDR_** |
| --- | --- | --- | --- | --- |
| Left Entorhinal Cortex | -0.008 | -0.015 - -0.001 | 0.096 | 0.092 |
| Left Inferior Parietal Lobule | -0.008 | -0.015 - -0.003 | 0.101 | 0.092 |
| Left Paracentral Lobule | -0.009 | -0.017 - -0.002 | 0.105 | 0.092 |
| Left Postcentral Gyrus | -0.016 | -0.028 - -0.006 | 0.195 | 0.092 |
| Left Rostral Middle Frontal Gyrus | -0.010 | -0.018 - -0.003 | 0.119 | 0.092 |
| Left Superior Temporal Gyrus | -0.008 | -0.015 - -0.002 | 0.095 | 0.092 |
| Left Supramarginal Gyrus | -0.010 | -0.018 - -0.003 | 0.120 | 0.092 |
| Right Paracentral Lobule | -0.009 | -0.016 - -0.003 | 0.108 | 0.092 |
| Right Postcentral Gyrus | -0.013 | -0.023 - -0.004 | 0.156 | 0.092 |
| Right Precentral Gyrus | -0.013 | -0.024 - -0.004 | 0.154 | 0.092 |
| Right Supramarginal Gyrus | -0.007 | -0.014 - -0.001 | 0.082 | 0.092 |

**Supplementary Table 17: Regions mediating the associations between COI 2.0 social/economic scores and internalising scores at baseline**

The table summarises the mediation analyses ran across regions that showed both environment-brain and brain-behaviour associations. The ACME (Average Causal Mediation Effect) estimate is reported, which reflects the indirect effect of the predictor on the outcome that is transmitted through the mediator, as well as proportion mediated values, which indicate the estimated fraction of the total effect explained by the mediator. Associated ACME confidence intervals and FDR-corrected p values are also reported. P values are false discovery rate (FDR) corrected to account for multiple comparisons. Analyses controlled for age and sex and included random intercepts for testing site and family ID (to account for sibling pairs).

| **Region** | **ACME estimate** | **ACME CI** | **Proportion mediated** | ***p*_FDR_** |
| --- | --- | --- | --- | --- |
| Left Entorhinal Cortex | -0.012 | -0.022 - -0.001 | 0.103 | 0.046 |
| Left Inferior Parietal Lobule | -0.011 | -0.019 - -0.004 | 0.093 | 0.035 |
| Left Paracentral Lobule | -0.012 | -0.023 - -0.003 | 0.110 | 0.035 |
| Left Postcentral Gyrus | -0.022 | -0.037 - -0.007 | 0.190 | 0.035 |
| Left Rostral Middle Frontal Gyrus | -0.014 | -0.026 - -0.003 | 0.125 | 0.035 |
| Left Superior Temporal Gyrus | -0.013 | -0.025 - -0.002 | 0.116 | 0.035 |
| Left Supramarginal Gyrus | -0.016 | -0.027 - -0.004 | 0.138 | 0.035 |
| Right Paracentral Lobule | -0.012 | -0.020 - -0.004 | 0.102 | 0.035 |
| Right Postcentral Gyrus | -0.017 | -0.030 - -0.005 | 0.153 | 0.035 |
| Right Precentral Gyrus | -0.018 | -0.033 - -0.004 | 0.161 | 0.035 |

**Supplementary Table 18: Regions mediating the associations between COI 2.0 education scores and internalising scores at baseline**

The table summarises the mediation analyses ran across regions that showed both environment-brain and brain-behaviour associations. The ACME (Average Causal Mediation Effect) estimate is reported, which reflects the indirect effect of the predictor on the outcome that is transmitted through the mediator, as well as proportion mediated values, which indicate the estimated fraction of the total effect explained by the mediator. Associated ACME confidence intervals and FDR-corrected p values are also reported. P values are false discovery rate (FDR) corrected to account for multiple comparisons. Analyses controlled for age and sex and included random intercepts for testing site and family ID (to account for sibling pairs).

| **Region** | **ACME estimate** | **ACME CI** | **Proportion mediated** | ***p*_FDR_** |
| --- | --- | --- | --- | --- |
| Left Entorhinal Cortex | -0.011 | -0.022 - -0.002 | 0.083 | 0.024 |
| Left Inferior Parietal Lobule | -0.012 | -0.020 - -0.004 | 0.085 | 0.011 |
| Left Paracentral Lobule | -0.010 | -0.019 - -0.002 | 0.075 | 0.022 |
| Left Postcentral Gyrus | -0.021 | -0.037 - -0.006 | 0.150 | 0.022 |
| Left Rostral Middle Frontal Gyrus | -0.015 | -0.027 - -0.003 | 0.108 | 0.022 |
| Left Superior Temporal Gyrus | -0.013 | -0.025 - -0.001 | 0.094 | 0.036 |
| Left Supramarginal Gyrus | -0.015 | -0.026 - -0.004 | 0.108 | 0.022 |
| Right Paracentral Lobule | -0.011 | -0.019 - -0.004 | 0.081 | 0.011 |
| Right Postcentral Gyrus | -0.017 | -0.031 - -0.005 | 0.127 | 0.022 |
| Right Precentral Gyrus | -0.017 | -0.029 - -0.004 | 0.120 | 0.022 |
| Right Supramarginal Gyrus | -0.010 | -0.020 - -0.001 | 0.075 | 0.024 |

**Supplementary Table 19: Regions mediating the associations between micro-environment scores and internalising scores at baseline**

The table summarises the mediation analyses ran across regions that showed both environment-brain and brain-behaviour associations. The ACME (Average Causal Mediation Effect) estimate is reported, which reflects the indirect effect of the predictor on the outcome that is transmitted through the mediator, as well as proportion mediated values, which indicate the estimated fraction of the total effect explained by the mediator. Associated ACME confidence intervals and FDR-corrected p values are also reported. P values are false discovery rate (FDR) corrected to account for multiple comparisons. Analyses controlled for age and sex and included random intercepts for testing site and family ID (to account for sibling pairs).

| **Region** | **ACME estimate** | **ACME CI** | **Proportion mediated** | ***p*_FDR_** |
| --- | --- | --- | --- | --- |
| Left Entorhinal Cortex | -0.005 | -0.012 - 0.000 | 0.114 | 0.624 |
| Left Inferior Parietal Lobule | -0.025 | -0.043 - -0.009 | 0.602 | 0.624 |
| Left Paracentral Lobule | -0.015 | -0.027 - -0.005 | 0.353 | 0.624 |
| Left Postcentral Gyrus | -0.019 | -0.033 - -0.007 | 0.454 | 0.624 |
| Left Rostral Middle Frontal Gyrus | -0.012 | -0.023 - -0.004 | 0.293 | 0.624 |
| Left Superior Temporal Gyrus | -0.010 | -0.019 - -0.002 | 0.233 | 0.624 |
| Left Supramarginal Gyrus | -0.019 | -0.035 - -0.008 | 0.461 | 0.624 |
| Right Paracentral Lobule | -0.016 | -0.028 - -0.006 | 0.380 | 0.624 |
| Right Postcentral Gyrus | -0.018 | -0.031 - -0.006 | 0.426 | 0.624 |
| Right Precentral Gyrus | -0.012 | -0.022 - -0.003 | 0.281 | 0.624 |
| Right Supramarginal Gyrus | -0.015 | -0.029 - -0.004 | 0.349 | 0.624 |

**Supplementary Table 20: Regions mediating the associations between health/environment scores and Flanker Inhibitory Control & Attention scores at baseline**

The table summarises the mediation analyses ran across regions that showed both environment-brain and brain-behaviour associations. The ACME (Average Causal Mediation Effect) estimate is reported, which reflects the indirect effect of the predictor on the outcome that is transmitted through the mediator, as well as proportion mediated values, which indicate the estimated fraction of the total effect explained by the mediator. Associated ACME confidence intervals and FDR-corrected p values are also reported. P values are false discovery rate (FDR) corrected to account for multiple comparisons. Analyses controlled for age and sex and included random intercepts for testing site and family ID (to account for sibling pairs).

| **Region** | **Proportion mediated** | **95% confidence interval** | ***p*_FDR_** |
| --- | --- | --- | --- |
| Left Banks of the Superior Temporal Sulcus | 0.012 | 0.000 - 0.028 | 0.060 |
| Left Caudal Anterior Cingulate Cortex | 0.016 | 0.006 - 0.031 | 0.000 |
| Left Caudal Middle Frontal Gyrus | 0.026 | 0.009 - 0.046 | 0.008 |
| Left Cuneus | 0.022 | 0.006 - 0.041 | 0.011 |
| Left Entorhinal Cortex | 0.031 | 0.016 - 0.051 | 0.000 |
| Left Fusiform Gyrus | 0.050 | 0.030 - 0.076 | 0.000 |
| Left Inferior Parietal Lobule | 0.013 | 0.001 - 0.027 | 0.021 |
| Left Inferior Temporal Gyrus | 0.039 | 0.016 - 0.065 | 0.000 |
| Left Insular Cortex | 0.049 | 0.028 - 0.075 | 0.000 |
| Left Isthmus of the Cingulate Cortex | 0.018 | 0.004 - 0.035 | 0.021 |
| Left Lateral Occipital Cortex | 0.041 | 0.022 - 0.067 | 0.000 |
| Left Lateral Orbitofrontal Cortex | 0.046 | 0.025 - 0.071 | 0.000 |
| Left Lingual Gyrus | 0.018 | 0.004 - 0.035 | 0.017 |
| Left Medial Orbitofrontal Cortex | 0.013 | 0.003 - 0.026 | 0.015 |
| Left Middle Temporal Gyrus | 0.055 | 0.032 - 0.082 | 0.000 |
| Left Paracentral Lobule | 0.019 | 0.006 - 0.037 | 0.008 |
| Left Parahippocampal Gyrus | 0.026 | 0.012 - 0.046 | 0.000 |
| Left Pars Opercularis | 0.017 | 0.001 - 0.034 | 0.034 |
| Left Pars Orbitalis | 0.020 | 0.003 - 0.040 | 0.030 |
| Left Pars Triangularis | 0.014 | 0.004 - 0.026 | 0.000 |
| Left Postcentral Gyrus | 0.045 | 0.022 - 0.074 | 0.000 |
| Left Posterior Cingulate Cortex | 0.029 | 0.015 - 0.048 | 0.000 |
| Left Precentral Gyrus | 0.047 | 0.025 - 0.074 | 0.000 |
| Left Precuneus | 0.040 | 0.020 - 0.065 | 0.000 |
| Left Rostral Anterior Cingulate Cortex | 0.030 | 0.015 - 0.051 | 0.000 |
| Left Rostral Middle Frontal Gyrus | 0.023 | 0.008 - 0.042 | 0.004 |
| Left Superior Frontal Gyrus | 0.033 | 0.016 - 0.054 | 0.000 |
| Left Superior Parietal Lobule | 0.026 | 0.008 - 0.048 | 0.015 |
| Left Superior Temporal Gyrus | 0.028 | 0.014 - 0.048 | 0.000 |
| Left Supramarginal Gyrus | 0.029 | 0.015 - 0.049 | 0.000 |
| Left Temporal Pole | 0.014 | 0.003 - 0.030 | 0.008 |
| Left Transverse Temporal Gyrus (Heschl’s Gyrus) | 0.025 | 0.010 - 0.043 | 0.000 |
| Right Banks of the Superior Temporal Sulcus | 0.014 | 0.001 - 0.030 | 0.040 |
| Right Caudal Middle Frontal Gyrus | 0.027 | 0.011 - 0.047 | 0.004 |
| Right Entorhinal Cortex | 0.015 | 0.000 - 0.032 | 0.062 |
| Right Fusiform Gyrus | 0.038 | 0.020 - 0.060 | 0.000 |
| Right Inferior Parietal Lobule | 0.031 | 0.014 - 0.053 | 0.000 |
| Right Inferior Temporal Gyrus | 0.032 | 0.012 - 0.054 | 0.000 |
| Right Insular Cortex | 0.033 | 0.017 - 0.053 | 0.000 |
| Right Isthmus of the Cingulate Cortex | 0.012 | 0.002 - 0.025 | 0.015 |
| Right Lateral Occipital Cortex | 0.055 | 0.033 - 0.084 | 0.000 |
| Right Lateral Orbitofrontal Cortex | 0.048 | 0.027 - 0.076 | 0.000 |
| Right Lingual Gyrus | 0.020 | 0.008 - 0.035 | 0.000 |
| Right Medial Orbitofrontal Cortex | 0.027 | 0.012 - 0.046 | 0.000 |
| Right Middle Temporal Gyrus | 0.065 | 0.039 - 0.097 | 0.000 |
| Right Paracentral Lobule | 0.022 | 0.009 - 0.038 | 0.000 |
| Right Parahippocampal Gyrus | 0.024 | 0.010 - 0.042 | 0.000 |
| Right Pars Opercularis | 0.012 | -0.004 - 0.027 | 0.144 |
| Right Pars Orbitalis | 0.021 | 0.004 - 0.044 | 0.014 |
| Right Postcentral Gyrus | 0.032 | 0.013 - 0.054 | 0.004 |
| Right Posterior Cingulate Cortex | 0.013 | 0.000 - 0.028 | 0.055 |
| Right Precentral Gyrus | 0.039 | 0.019 - 0.065 | 0.000 |
| Right Precuneus | 0.040 | 0.020 - 0.066 | 0.000 |
| Right Rostral Anterior Cingulate Cortex | 0.023 | 0.011 - 0.039 | 0.000 |
| Right Rostral Middle Frontal Gyrus | 0.026 | 0.010 - 0.043 | 0.004 |
| Right Superior Frontal Gyrus | 0.046 | 0.027 - 0.072 | 0.000 |
| Right Superior Parietal Lobule | 0.022 | 0.005 - 0.042 | 0.017 |
| Right Superior Temporal Gyrus | 0.037 | 0.019 - 0.061 | 0.000 |
| Right Supramarginal Gyrus | 0.019 | 0.007 - 0.036 | 0.000 |
| Right Temporal Pole | 0.006 | 0.000 - 0.016 | 0.060 |
| Right Transverse Temporal Gyrus (Heschl’s Gyrus) | 0.031 | 0.014 - 0.052 | 0.000 |

**Supplementary Table 21: Regions mediating the associations between social/economic scores and Flanker Inhibitory Control & Attention scores at baseline**

The table summarises the mediation analyses ran across regions that showed both environment-brain and brain-behaviour associations. The ACME (Average Causal Mediation Effect) estimate is reported, which reflects the indirect effect of the predictor on the outcome that is transmitted through the mediator, as well as proportion mediated values, which indicate the estimated fraction of the total effect explained by the mediator. Associated ACME confidence intervals and FDR-corrected p values are also reported. P values are false discovery rate (FDR) corrected to account for multiple comparisons. Analyses controlled for age and sex and included random intercepts for testing site and family ID (to account for sibling pairs).

| **Region** | **ACME estimate** | **ACME CI** | **Proportion mediated** | ***p*_FDR_** |
| --- | --- | --- | --- | --- |
| Left Banks of the Superior Temporal Sulcus | 0.009 | -0.002 - 0.022 | 0.011 | 0.109 |
| Left Caudal Anterior Cingulate Cortex | 0.020 | 0.008 - 0.036 | 0.024 | 0.000 |
| Left Caudal Middle Frontal Gyrus | 0.022 | 0.006 - 0.041 | 0.026 | 0.003 |
| Left Cuneus | 0.019 | 0.003 - 0.037 | 0.022 | 0.038 |
| Left Entorhinal Cortex | 0.031 | 0.014 - 0.051 | 0.037 | 0.000 |
| Left Fusiform Gyrus | 0.052 | 0.030 - 0.076 | 0.061 | 0.000 |
| Left Inferior Parietal Lobule | 0.011 | 0.000 - 0.023 | 0.013 | 0.055 |
| Left Inferior Temporal Gyrus | 0.033 | 0.009 - 0.057 | 0.038 | 0.003 |
| Left Insular Cortex | 0.050 | 0.030 - 0.069 | 0.058 | 0.000 |
| Left Isthmus of the Cingulate Cortex | 0.016 | 0.003 - 0.030 | 0.019 | 0.019 |
| Left Lateral Occipital Cortex | 0.038 | 0.016 - 0.062 | 0.045 | 0.003 |
| Left Lateral Orbitofrontal Cortex | 0.046 | 0.023 - 0.071 | 0.054 | 0.000 |
| Left Lingual Gyrus | 0.017 | 0.001 - 0.034 | 0.020 | 0.046 |
| Left Medial Orbitofrontal Cortex | 0.013 | 0.001 - 0.025 | 0.015 | 0.045 |
| Left Middle Temporal Gyrus | 0.050 | 0.028 - 0.074 | 0.058 | 0.000 |
| Left Paracentral Lobule | 0.018 | 0.004 - 0.033 | 0.021 | 0.021 |
| Left Parahippocampal Gyrus | 0.028 | 0.012 - 0.044 | 0.033 | 0.003 |
| Left Pars Opercularis | 0.013 | 0.000 - 0.028 | 0.015 | 0.061 |
| Left Pars Orbitalis | 0.016 | -0.001 - 0.033 | 0.019 | 0.070 |
| Left Pars Triangularis | 0.013 | 0.005 - 0.024 | 0.016 | 0.000 |
| Left Postcentral Gyrus | 0.040 | 0.017 - 0.062 | 0.046 | 0.000 |
| Left Posterior Cingulate Cortex | 0.024 | 0.013 - 0.037 | 0.028 | 0.000 |
| Left Precentral Gyrus | 0.041 | 0.019 - 0.065 | 0.048 | 0.000 |
| Left Precuneus | 0.036 | 0.015 - 0.061 | 0.042 | 0.000 |
| Left Rostral Anterior Cingulate Cortex | 0.036 | 0.020 - 0.052 | 0.042 | 0.000 |
| Left Rostral Middle Frontal Gyrus | 0.021 | 0.003 - 0.040 | 0.025 | 0.028 |
| Left Superior Frontal Gyrus | 0.029 | 0.012 - 0.047 | 0.034 | 0.000 |
| Left Superior Parietal Lobule | 0.023 | 0.004 - 0.044 | 0.027 | 0.019 |
| Left Superior Temporal Gyrus | 0.031 | 0.014 - 0.052 | 0.037 | 0.000 |
| Left Supramarginal Gyrus | 0.030 | 0.012 - 0.051 | 0.036 | 0.000 |
| Left Temporal Pole | 0.016 | 0.005 - 0.030 | 0.018 | 0.012 |
| Left Transverse Temporal Gyrus (Heschl’s Gyrus) | 0.025 | 0.010 - 0.042 | 0.029 | 0.000 |
| Right Banks of the Superior Temporal Sulcus | 0.012 | -0.001 - 0.026 | 0.014 | 0.082 |
| Right Caudal Middle Frontal Gyrus | 0.026 | 0.010 - 0.044 | 0.030 | 0.000 |
| Right Entorhinal Cortex | 0.012 | -0.005 - 0.030 | 0.014 | 0.167 |
| Right Fusiform Gyrus | 0.039 | 0.017 - 0.062 | 0.045 | 0.000 |
| Right Inferior Parietal Lobule | 0.029 | 0.013 - 0.047 | 0.034 | 0.000 |
| Right Inferior Temporal Gyrus | 0.028 | 0.005 - 0.052 | 0.033 | 0.012 |
| Right Insular Cortex | 0.033 | 0.018 - 0.051 | 0.038 | 0.000 |
| Right Isthmus of the Cingulate Cortex | 0.012 | 0.003 - 0.022 | 0.013 | 0.021 |
| Right Lateral Occipital Cortex | 0.052 | 0.026 - 0.078 | 0.061 | 0.000 |
| Right Lateral Orbitofrontal Cortex | 0.046 | 0.024 - 0.070 | 0.054 | 0.000 |
| Right Lingual Gyrus | 0.019 | 0.006 - 0.034 | 0.023 | 0.000 |
| Right Medial Orbitofrontal Cortex | 0.027 | 0.010 - 0.045 | 0.031 | 0.000 |
| Right Middle Temporal Gyrus | 0.057 | 0.030 - 0.082 | 0.067 | 0.000 |
| Right Paracentral Lobule | 0.020 | 0.008 - 0.034 | 0.023 | 0.000 |
| Right Parahippocampal Gyrus | 0.023 | 0.009 - 0.040 | 0.027 | 0.000 |
| Right Pars Opercularis | 0.009 | -0.005 - 0.023 | 0.010 | 0.202 |
| Right Pars Orbitalis | 0.017 | 0.000 - 0.036 | 0.020 | 0.062 |
| Right Postcentral Gyrus | 0.029 | 0.009 - 0.049 | 0.033 | 0.003 |
| Right Posterior Cingulate Cortex | 0.011 | -0.002 - 0.026 | 0.013 | 0.112 |
| Right Precentral Gyrus | 0.036 | 0.014 - 0.059 | 0.042 | 0.000 |
| Right Precuneus | 0.035 | 0.014 - 0.057 | 0.041 | 0.000 |
| Right Rostral Anterior Cingulate Cortex | 0.024 | 0.012 - 0.040 | 0.028 | 0.000 |
| Right Rostral Middle Frontal Gyrus | 0.024 | 0.007 - 0.042 | 0.028 | 0.000 |
| Right Superior Frontal Gyrus | 0.048 | 0.026 - 0.069 | 0.056 | 0.000 |
| Right Superior Parietal Lobule | 0.019 | -0.001 - 0.039 | 0.022 | 0.083 |
| Right Superior Temporal Gyrus | 0.037 | 0.017 - 0.057 | 0.043 | 0.000 |
| Right Supramarginal Gyrus | 0.018 | 0.005 - 0.033 | 0.021 | 0.014 |
| Right Temporal Pole | 0.007 | -0.001 - 0.017 | 0.008 | 0.088 |
| Right Transverse Temporal Gyrus (Heschl’s Gyrus) | 0.027 | 0.011 - 0.044 | 0.032 | 0.000 |

**Supplementary Table 22: Regions mediating the associations between education scores and Flanker Inhibitory Control & Attention scores at baseline**

The table summarises the mediation analyses ran across regions that showed both environment-brain and brain-behaviour associations. The ACME (Average Causal Mediation Effect) estimate is reported, which reflects the indirect effect of the predictor on the outcome that is transmitted through the mediator, as well as proportion mediated values, which indicate the estimated fraction of the total effect explained by the mediator. Associated ACME confidence intervals and FDR-corrected p values are also reported. P values are false discovery rate (FDR) corrected to account for multiple comparisons. Analyses controlled for age and sex and included random intercepts for testing site and family ID (to account for sibling pairs).

| **Region** | **ACME estimate** | **ACME CI** | **Proportion mediated** | ***p*_FDR_** |
| --- | --- | --- | --- | --- |
| Left Banks of the Superior Temporal Sulcus | 0.011 | -0.001 - 0.023 | 0.015 | 0.078 |
| Left Caudal Anterior Cingulate Cortex | 0.019 | 0.008 - 0.034 | 0.027 | 0.000 |
| Left Caudal Middle Frontal Gyrus | 0.022 | 0.009 - 0.039 | 0.032 | 0.006 |
| Left Cuneus | 0.022 | 0.005 - 0.039 | 0.031 | 0.015 |
| Left Entorhinal Cortex | 0.034 | 0.017 - 0.053 | 0.048 | 0.000 |
| Left Fusiform Gyrus | 0.057 | 0.036 - 0.081 | 0.080 | 0.000 |
| Left Inferior Parietal Lobule | 0.013 | 0.002 - 0.025 | 0.018 | 0.030 |
| Left Inferior Temporal Gyrus | 0.038 | 0.014 - 0.063 | 0.053 | 0.000 |
| Left Insular Cortex | 0.054 | 0.036 - 0.073 | 0.077 | 0.000 |
| Left Isthmus of the Cingulate Cortex | 0.018 | 0.004 - 0.034 | 0.026 | 0.013 |
| Left Lateral Occipital Cortex | 0.042 | 0.022 - 0.065 | 0.059 | 0.000 |
| Left Lateral Orbitofrontal Cortex | 0.050 | 0.028 - 0.077 | 0.071 | 0.000 |
| Left Lingual Gyrus | 0.019 | 0.003 - 0.036 | 0.027 | 0.022 |
| Left Medial Orbitofrontal Cortex | 0.015 | 0.002 - 0.029 | 0.021 | 0.032 |
| Left Middle Temporal Gyrus | 0.054 | 0.031 - 0.080 | 0.077 | 0.000 |
| Left Paracentral Lobule | 0.017 | 0.005 - 0.031 | 0.024 | 0.009 |
| Left Parahippocampal Gyrus | 0.031 | 0.014 - 0.049 | 0.044 | 0.000 |
| Left Pars Opercularis | 0.015 | 0.002 - 0.030 | 0.021 | 0.022 |
| Left Pars Orbitalis | 0.018 | 0.000 - 0.037 | 0.026 | 0.049 |
| Left Pars Triangularis | 0.016 | 0.006 - 0.028 | 0.023 | 0.003 |
| Left Postcentral Gyrus | 0.044 | 0.022 - 0.068 | 0.062 | 0.000 |
| Left Posterior Cingulate Cortex | 0.027 | 0.015 - 0.043 | 0.038 | 0.000 |
| Left Precentral Gyrus | 0.043 | 0.022 - 0.065 | 0.060 | 0.000 |
| Left Precuneus | 0.039 | 0.019 - 0.061 | 0.055 | 0.000 |
| Left Rostral Anterior Cingulate Cortex | 0.033 | 0.020 - 0.049 | 0.047 | 0.000 |
| Left Rostral Middle Frontal Gyrus | 0.025 | 0.006 - 0.044 | 0.035 | 0.011 |
| Left Superior Frontal Gyrus | 0.030 | 0.015 - 0.047 | 0.043 | 0.000 |
| Left Superior Parietal Lobule | 0.026 | 0.006 - 0.048 | 0.037 | 0.013 |
| Left Superior Temporal Gyrus | 0.034 | 0.016 - 0.053 | 0.048 | 0.000 |
| Left Supramarginal Gyrus | 0.032 | 0.014 - 0.053 | 0.046 | 0.000 |
| Left Temporal Pole | 0.017 | 0.004 - 0.030 | 0.024 | 0.006 |
| Left Transverse Temporal Gyrus (Heschl’s Gyrus) | 0.026 | 0.011 - 0.042 | 0.037 | 0.000 |
| Right Banks of the Superior Temporal Sulcus | 0.013 | 0.001 - 0.026 | 0.018 | 0.041 |
| Right Caudal Middle Frontal Gyrus | 0.027 | 0.012 - 0.044 | 0.038 | 0.000 |
| Right Entorhinal Cortex | 0.014 | -0.002 - 0.030 | 0.019 | 0.094 |
| Right Fusiform Gyrus | 0.044 | 0.021 - 0.068 | 0.062 | 0.000 |
| Right Inferior Parietal Lobule | 0.030 | 0.014 - 0.046 | 0.042 | 0.000 |
| Right Inferior Temporal Gyrus | 0.031 | 0.009 - 0.053 | 0.044 | 0.011 |
| Right Insular Cortex | 0.035 | 0.020 - 0.054 | 0.049 | 0.000 |
| Right Isthmus of the Cingulate Cortex | 0.012 | 0.002 - 0.024 | 0.017 | 0.009 |
| Right Lateral Occipital Cortex | 0.056 | 0.032 - 0.081 | 0.079 | 0.000 |
| Right Lateral Orbitofrontal Cortex | 0.049 | 0.028 - 0.072 | 0.069 | 0.000 |
| Right Lingual Gyrus | 0.023 | 0.008 - 0.038 | 0.032 | 0.006 |
| Right Medial Orbitofrontal Cortex | 0.031 | 0.013 - 0.050 | 0.043 | 0.000 |
| Right Middle Temporal Gyrus | 0.064 | 0.038 - 0.091 | 0.090 | 0.000 |
| Right Paracentral Lobule | 0.020 | 0.009 - 0.034 | 0.028 | 0.000 |
| Right Parahippocampal Gyrus | 0.028 | 0.012 - 0.047 | 0.039 | 0.000 |
| Right Pars Opercularis | 0.010 | -0.002 - 0.022 | 0.014 | 0.104 |
| Right Pars Orbitalis | 0.019 | 0.002 - 0.037 | 0.027 | 0.029 |
| Right Postcentral Gyrus | 0.032 | 0.011 - 0.054 | 0.046 | 0.006 |
| Right Posterior Cingulate Cortex | 0.012 | 0.000 - 0.025 | 0.017 | 0.051 |
| Right Precentral Gyrus | 0.038 | 0.018 - 0.059 | 0.053 | 0.000 |
| Right Precuneus | 0.038 | 0.019 - 0.062 | 0.054 | 0.000 |
| Right Rostral Anterior Cingulate Cortex | 0.022 | 0.010 - 0.036 | 0.031 | 0.000 |
| Right Rostral Middle Frontal Gyrus | 0.025 | 0.011 - 0.042 | 0.036 | 0.000 |
| Right Superior Frontal Gyrus | 0.048 | 0.029 - 0.068 | 0.068 | 0.000 |
| Right Superior Parietal Lobule | 0.022 | 0.000 - 0.041 | 0.031 | 0.050 |
| Right Superior Temporal Gyrus | 0.041 | 0.021 - 0.062 | 0.058 | 0.000 |
| Right Supramarginal Gyrus | 0.021 | 0.006 - 0.036 | 0.030 | 0.011 |
| Right Temporal Pole | 0.008 | -0.001 - 0.018 | 0.011 | 0.087 |
| Right Transverse Temporal Gyrus (Heschl’s Gyrus) | 0.028 | 0.012 - 0.045 | 0.040 | 0.000 |

**Supplementary Table 23: Regions mediating the associations between micro-environment scores and Flanker Inhibitory Control & Attention scores at baseline**

The table summarises the mediation analyses ran across regions that showed both environment-brain and brain-behaviour associations. The ACME (Average Causal Mediation Effect) estimate is reported, which reflects the indirect effect of the predictor on the outcome that is transmitted through the mediator, as well as proportion mediated values, which indicate the estimated fraction of the total effect explained by the mediator. Associated ACME confidence intervals and FDR-corrected p values are also reported. P values are false discovery rate (FDR) corrected to account for multiple comparisons. Analyses controlled for age and sex and included random intercepts for testing site and family ID (to account for sibling pairs).

| **Region** | **ACME estimate** | **ACME CI** | **Proportion mediated** | ***p*_FDR_** |
| --- | --- | --- | --- | --- |
| Left Banks of the Superior Temporal Sulcus | 0.009 | 0.001 - 0.021 | -0.295 | 0.998 |
| Left Caudal Anterior Cingulate Cortex | 0.000 | -0.013 - 0.013 | -0.015 | 0.998 |
| Left Caudal Middle Frontal Gyrus | -0.011 | -0.026 - 0.002 | 0.360 | 0.998 |
| Left Cuneus | -0.007 | -0.019 - 0.004 | 0.221 | 0.998 |
| Left Entorhinal Cortex | -0.015 | -0.033 - -0.001 | 0.513 | 0.998 |
| Left Fusiform Gyrus | -0.006 | -0.025 - 0.013 | 0.188 | 0.998 |
| Left Inferior Parietal Lobule | 0.005 | -0.004 - 0.015 | -0.167 | 0.998 |
| Left Inferior Temporal Gyrus | 0.001 | -0.013 - 0.016 | -0.037 | 0.998 |
| Left Insular Cortex | 0.016 | -0.005 - 0.040 | -0.548 | 0.998 |
| Left Isthmus of the Cingulate Cortex | 0.006 | -0.004 - 0.018 | -0.193 | 0.998 |
| Left Lateral Occipital Cortex | -0.014 | -0.033 - 0.001 | 0.469 | 0.998 |
| Left Lateral Orbitofrontal Cortex | -0.005 | -0.024 - 0.012 | 0.177 | 0.998 |
| Left Lingual Gyrus | 0.001 | -0.009 - 0.012 | -0.044 | 0.998 |
| Left Medial Orbitofrontal Cortex | -0.003 | -0.014 - 0.006 | 0.102 | 0.998 |
| Left Middle Temporal Gyrus | 0.013 | -0.005 - 0.035 | -0.436 | 0.998 |
| Left Paracentral Lobule | 0.000 | -0.011 - 0.011 | -0.002 | 0.998 |
| Left Parahippocampal Gyrus | 0.002 | -0.013 - 0.017 | -0.072 | 0.998 |
| Left Pars Opercularis | -0.008 | -0.021 - 0.001 | 0.271 | 0.998 |
| Left Pars Orbitalis | -0.007 | -0.020 - 0.003 | 0.237 | 0.998 |
| Left Pars Triangularis | -0.008 | -0.022 - 0.005 | 0.263 | 0.998 |
| Left Postcentral Gyrus | 0.002 | -0.013 - 0.019 | -0.061 | 0.998 |
| Left Posterior Cingulate Cortex | 0.001 | -0.014 - 0.017 | -0.034 | 0.998 |
| Left Precentral Gyrus | -0.001 | -0.017 - 0.015 | 0.021 | 0.998 |
| Left Precuneus | -0.010 | -0.026 - 0.004 | 0.335 | 0.998 |
| Left Rostral Anterior Cingulate Cortex | -0.008 | -0.027 - 0.011 | 0.270 | 0.998 |
| Left Rostral Middle Frontal Gyrus | -0.007 | -0.020 - 0.005 | 0.236 | 0.998 |
| Left Superior Frontal Gyrus | -0.003 | -0.017 - 0.011 | 0.087 | 0.998 |
| Left Superior Parietal Lobule | -0.004 | -0.016 - 0.007 | 0.133 | 0.998 |
| Left Superior Temporal Gyrus | 0.002 | -0.014 - 0.018 | -0.063 | 0.998 |
| Left Supramarginal Gyrus | -0.002 | -0.016 - 0.012 | 0.058 | 0.998 |
| Left Temporal Pole | -0.012 | -0.026 - -0.002 | 0.406 | 0.998 |
| Left Transverse Temporal Gyrus (Heschl’s Gyrus) | 0.000 | -0.014 - 0.013 | -0.008 | 0.998 |
| Right Banks of the Superior Temporal Sulcus | 0.011 | 0.003 - 0.025 | -0.388 | 0.998 |
| Right Caudal Middle Frontal Gyrus | -0.014 | -0.030 - 0.000 | 0.477 | 0.998 |
| Right Entorhinal Cortex | -0.008 | -0.020 - 0.001 | 0.256 | 0.998 |
| Right Fusiform Gyrus | -0.003 | -0.020 - 0.013 | 0.115 | 0.998 |
| Right Inferior Parietal Lobule | 0.001 | -0.014 - 0.016 | -0.027 | 0.998 |
| Right Inferior Temporal Gyrus | -0.008 | -0.022 - 0.004 | 0.260 | 0.998 |
| Right Insular Cortex | 0.012 | -0.006 - 0.030 | -0.397 | 0.998 |
| Right Isthmus of the Cingulate Cortex | 0.003 | -0.007 - 0.013 | -0.086 | 0.998 |
| Right Lateral Occipital Cortex | -0.025 | -0.048 - -0.007 | 0.840 | 0.998 |
| Right Lateral Orbitofrontal Cortex | -0.005 | -0.023 - 0.012 | 0.177 | 0.998 |
| Right Lingual Gyrus | -0.003 | -0.015 - 0.009 | 0.089 | 0.998 |
| Right Medial Orbitofrontal Cortex | -0.001 | -0.016 - 0.014 | 0.043 | 0.998 |
| Right Middle Temporal Gyrus | -0.007 | -0.027 - 0.012 | 0.231 | 0.998 |
| Right Paracentral Lobule | 0.000 | -0.012 - 0.014 | -0.017 | 0.998 |
| Right Parahippocampal Gyrus | 0.008 | -0.005 - 0.022 | -0.263 | 0.998 |
| Right Pars Opercularis | 0.000 | -0.009 - 0.008 | -0.003 | 0.998 |
| Right Pars Orbitalis | -0.008 | -0.020 - 0.001 | 0.274 | 0.998 |
| Right Postcentral Gyrus | -0.001 | -0.015 - 0.013 | 0.034 | 0.998 |
| Right Posterior Cingulate Cortex | -0.002 | -0.011 - 0.007 | 0.054 | 0.998 |
| Right Precentral Gyrus | -0.005 | -0.020 - 0.010 | 0.167 | 0.998 |
| Right Precuneus | -0.004 | -0.018 - 0.011 | 0.125 | 0.998 |
| Right Rostral Anterior Cingulate Cortex | 0.001 | -0.013 - 0.016 | -0.039 | 0.998 |
| Right Rostral Middle Frontal Gyrus | -0.012 | -0.028 - 0.000 | 0.412 | 0.998 |
| Right Superior Frontal Gyrus | -0.009 | -0.028 - 0.010 | 0.288 | 0.998 |
| Right Superior Parietal Lobule | 0.003 | -0.007 - 0.015 | -0.115 | 0.998 |
| Right Superior Temporal Gyrus | 0.011 | -0.005 - 0.029 | -0.383 | 0.998 |
| Right Supramarginal Gyrus | 0.012 | 0.001 - 0.028 | -0.407 | 0.998 |
| Right Temporal Pole | -0.008 | -0.019 - 0.000 | 0.254 | 0.998 |
| Right Transverse Temporal Gyrus (Heschl’s Gyrus) | 0.007 | -0.007 - 0.022 | -0.229 | 0.998 |

**Supplementary Table 24: Regions mediating the associations between health/environment scores and Pattern Comparison Processing Speed scores at baseline**

The table summarises the mediation analyses ran across regions that showed both environment-brain and brain-behaviour associations. The ACME (Average Causal Mediation Effect) estimate is reported, which reflects the indirect effect of the predictor on the outcome that is transmitted through the mediator, as well as proportion mediated values, which indicate the estimated fraction of the total effect explained by the mediator. Associated ACME confidence intervals and FDR-corrected p values are also reported. P values are false discovery rate (FDR) corrected to account for multiple comparisons. Analyses controlled for age and sex and included random intercepts for testing site and family ID (to account for sibling pairs).

| **Region** | **ACME estimate** | **ACME CI** | **Proportion mediated** | ***p*_FDR_** |
| --- | --- | --- | --- | --- |
| Left Banks of the Superior Temporal Sulcus | -0.002 | -0.017 - 0.014 | -0.001 | 0.875 |
| Left Caudal Anterior Cingulate Cortex | 0.010 | 0.000 - 0.024 | 0.009 | 0.612 |
| Left Caudal Middle Frontal Gyrus | 0.012 | -0.010 - 0.032 | 0.010 | 0.612 |
| Left Frontal Pole | -0.004 | -0.020 - 0.011 | -0.004 | 0.754 |
| Left Inferior Parietal Lobule | -0.004 | -0.018 - 0.009 | -0.003 | 0.759 |
| Left Isthmus of the Cingulate Cortex | 0.010 | -0.007 - 0.027 | 0.008 | 0.612 |
| Left Lateral Orbitofrontal Cortex | 0.013 | -0.010 - 0.037 | 0.012 | 0.612 |
| Left Lingual Gyrus | 0.011 | -0.006 - 0.029 | 0.010 | 0.612 |
| Left Medial Orbitofrontal Cortex | -0.002 | -0.015 - 0.010 | -0.002 | 0.875 |
| Left Paracentral Lobule | 0.002 | -0.015 - 0.019 | 0.002 | 0.875 |
| Left Parahippocampal Gyrus | 0.012 | -0.003 - 0.028 | 0.010 | 0.612 |
| Left Pars Opercularis | 0.007 | -0.010 - 0.027 | 0.006 | 0.692 |
| Left Pars Orbitalis | 0.015 | -0.008 - 0.037 | 0.013 | 0.612 |
| Left Pars Triangularis | 0.004 | -0.006 - 0.015 | 0.004 | 0.658 |
| Left Pericalcarine Cortex | 0.006 | -0.005 - 0.020 | 0.006 | 0.619 |
| Left Postcentral Gyrus | 0.005 | -0.022 - 0.032 | 0.004 | 0.845 |
| Left Precuneus | 0.015 | -0.008 - 0.040 | 0.013 | 0.612 |
| Left Rostral Middle Frontal Gyrus | -0.003 | -0.021 - 0.017 | -0.002 | 0.875 |
| Left Superior Frontal Gyrus | 0.008 | -0.014 - 0.027 | 0.007 | 0.754 |
| Left Superior Parietal Lobule | 0.013 | -0.008 - 0.034 | 0.011 | 0.612 |
| Left Temporal Pole | 0.009 | -0.003 - 0.022 | 0.008 | 0.612 |
| Left Transverse Temporal Gyrus (Heschl’s Gyrus) | 0.012 | -0.004 - 0.030 | 0.010 | 0.612 |
| Right Banks of the Superior Temporal Sulcus | -0.004 | -0.019 - 0.011 | -0.003 | 0.754 |
| Right Caudal Anterior Cingulate Cortex | -0.001 | -0.015 - 0.012 | -0.001 | 0.875 |
| Right Cuneus | 0.012 | -0.007 - 0.032 | 0.010 | 0.612 |
| Right Frontal Pole | -0.011 | -0.030 - 0.007 | -0.009 | 0.612 |
| Right Fusiform Gyrus | 0.013 | -0.008 - 0.037 | 0.012 | 0.612 |
| Right Inferior Parietal Lobule | 0.012 | -0.006 - 0.032 | 0.011 | 0.612 |
| Right Isthmus of the Cingulate Cortex | 0.004 | -0.006 - 0.016 | 0.003 | 0.754 |
| Right Lingual Gyrus | 0.009 | -0.006 - 0.025 | 0.008 | 0.612 |
| Right Medial Orbitofrontal Cortex | 0.011 | -0.006 - 0.029 | 0.010 | 0.612 |
| Right Middle Temporal Gyrus | 0.008 | -0.023 - 0.040 | 0.007 | 0.754 |
| Right Parahippocampal Gyrus | 0.011 | -0.004 - 0.028 | 0.010 | 0.612 |
| Right Pars Opercularis | 0.004 | -0.015 - 0.023 | 0.003 | 0.845 |
| Right Pars Orbitalis | 0.001 | -0.022 - 0.024 | 0.001 | 0.908 |
| Right Pars Triangularis | 0.004 | -0.008 - 0.016 | 0.003 | 0.729 |
| Right Pericalcarine Cortex | 0.006 | -0.002 - 0.018 | 0.006 | 0.612 |
| Right Posterior Cingulate Cortex | 0.005 | -0.010 - 0.020 | 0.004 | 0.754 |
| Right Precuneus | -0.002 | -0.028 - 0.024 | -0.002 | 0.875 |
| Right Rostral Anterior Cingulate Cortex | 0.007 | -0.005 - 0.021 | 0.006 | 0.619 |
| Right Rostral Middle Frontal Gyrus | -0.009 | -0.028 - 0.008 | -0.007 | 0.658 |
| Right Superior Frontal Gyrus | 0.011 | -0.008 - 0.032 | 0.010 | 0.612 |
| Right Superior Parietal Lobule | 0.004 | -0.016 - 0.027 | 0.004 | 0.875 |
| Right Superior Temporal Gyrus | 0.009 | -0.011 - 0.030 | 0.008 | 0.658 |
| Right Supramarginal Gyrus | 0.005 | -0.011 - 0.020 | 0.004 | 0.754 |
| Right Transverse Temporal Gyrus (Heschl’s Gyrus) | 0.010 | -0.010 - 0.032 | 0.008 | 0.624 |

**Supplementary Table 25: Regions mediating the associations between social/economic scores and Pattern Comparison Processing Speed scores at baseline**

The table summarises the mediation analyses ran across regions that showed both environment-brain and brain-behaviour associations. The ACME (Average Causal Mediation Effect) estimate is reported, which reflects the indirect effect of the predictor on the outcome that is transmitted through the mediator, as well as proportion mediated values, which indicate the estimated fraction of the total effect explained by the mediator. Associated ACME confidence intervals and FDR-corrected p values are also reported. P values are false discovery rate (FDR) corrected to account for multiple comparisons. Analyses controlled for age and sex and included random intercepts for testing site and family ID (to account for sibling pairs).

| **Region** | **ACME estimate** | **ACME CI** | **Proportion mediated** | ***p*_FDR_** |
| --- | --- | --- | --- | --- |
| Left Banks of the Superior Temporal Sulcus | -0.004 | -0.022 - 0.015 | -0.003 | 0.844 |
| Left Caudal Anterior Cingulate Cortex | 0.015 | -0.004 - 0.039 | 0.013 | 0.813 |
| Left Caudal Middle Frontal Gyrus | 0.012 | -0.015 - 0.039 | 0.010 | 0.813 |
| Left Frontal Pole | -0.008 | -0.028 - 0.011 | -0.007 | 0.813 |
| Left Inferior Parietal Lobule | -0.006 | -0.024 - 0.010 | -0.005 | 0.813 |
| Left Isthmus of the Cingulate Cortex | 0.010 | -0.012 - 0.034 | 0.009 | 0.813 |
| Left Lateral Orbitofrontal Cortex | 0.010 | -0.026 - 0.047 | 0.009 | 0.836 |
| Left Lingual Gyrus | 0.012 | -0.017 - 0.040 | 0.010 | 0.813 |
| Left Medial Orbitofrontal Cortex | -0.006 | -0.025 - 0.011 | -0.005 | 0.813 |
| Left Paracentral Lobule | -0.002 | -0.025 - 0.022 | -0.002 | 0.935 |
| Left Parahippocampal Gyrus | 0.014 | -0.009 - 0.041 | 0.012 | 0.813 |
| Left Pars Opercularis | 0.007 | -0.014 - 0.029 | 0.006 | 0.813 |
| Left Pars Orbitalis | 0.015 | -0.013 - 0.043 | 0.013 | 0.813 |
| Left Pars Triangularis | 0.005 | -0.007 - 0.019 | 0.004 | 0.813 |
| Left Pericalcarine Cortex | 0.007 | -0.010 - 0.024 | 0.006 | 0.813 |
| Left Postcentral Gyrus | -0.002 | -0.041 - 0.036 | -0.002 | 0.935 |
| Left Precuneus | 0.014 | -0.022 - 0.051 | 0.012 | 0.813 |
| Left Rostral Middle Frontal Gyrus | -0.010 | -0.041 - 0.017 | -0.009 | 0.813 |
| Left Superior Frontal Gyrus | 0.006 | -0.021 - 0.035 | 0.005 | 0.836 |
| Left Superior Parietal Lobule | 0.012 | -0.020 - 0.043 | 0.010 | 0.813 |
| Left Temporal Pole | 0.012 | -0.009 - 0.033 | 0.010 | 0.813 |
| Left Transverse Temporal Gyrus (Heschl’s Gyrus) | 0.014 | -0.009 - 0.039 | 0.012 | 0.813 |
| Right Banks of the Superior Temporal Sulcus | -0.008 | -0.031 - 0.014 | -0.007 | 0.813 |
| Right Caudal Anterior Cingulate Cortex | -0.003 | -0.021 - 0.015 | -0.003 | 0.844 |
| Right Cuneus | 0.012 | -0.014 - 0.040 | 0.010 | 0.813 |
| Right Frontal Pole | -0.019 | -0.044 - 0.004 | -0.016 | 0.813 |
| Right Fusiform Gyrus | 0.012 | -0.019 - 0.044 | 0.011 | 0.813 |
| Right Inferior Parietal Lobule | 0.013 | -0.010 - 0.039 | 0.011 | 0.813 |
| Right Isthmus of the Cingulate Cortex | 0.004 | -0.010 - 0.019 | 0.003 | 0.836 |
| Right Lingual Gyrus | 0.010 | -0.012 - 0.032 | 0.008 | 0.813 |
| Right Medial Orbitofrontal Cortex | 0.012 | -0.017 - 0.039 | 0.010 | 0.813 |
| Right Middle Temporal Gyrus | 0.002 | -0.037 - 0.041 | 0.002 | 0.942 |
| Right Parahippocampal Gyrus | 0.013 | -0.010 - 0.036 | 0.011 | 0.813 |
| Right Pars Opercularis | 0.002 | -0.020 - 0.024 | 0.002 | 0.935 |
| Right Pars Orbitalis | -0.003 | -0.033 - 0.028 | -0.003 | 0.935 |
| Right Pars Triangularis | 0.004 | -0.008 - 0.016 | 0.003 | 0.813 |
| Right Pericalcarine Cortex | 0.009 | -0.007 - 0.025 | 0.008 | 0.813 |
| Right Posterior Cingulate Cortex | 0.004 | -0.018 - 0.026 | 0.003 | 0.844 |
| Right Precuneus | -0.010 | -0.042 - 0.026 | -0.008 | 0.836 |
| Right Rostral Anterior Cingulate Cortex | 0.008 | -0.011 - 0.030 | 0.007 | 0.813 |
| Right Rostral Middle Frontal Gyrus | -0.018 | -0.045 - 0.011 | -0.015 | 0.813 |
| Right Superior Frontal Gyrus | 0.009 | -0.023 - 0.044 | 0.008 | 0.836 |
| Right Superior Parietal Lobule | 0.000 | -0.032 - 0.031 | 0.000 | 0.982 |
| Right Superior Temporal Gyrus | 0.008 | -0.021 - 0.039 | 0.007 | 0.836 |
| Right Supramarginal Gyrus | 0.004 | -0.018 - 0.025 | 0.003 | 0.844 |
| Right Transverse Temporal Gyrus (Heschl’s Gyrus) | 0.009 | -0.015 - 0.036 | 0.008 | 0.813 |

**Supplementary Table 26: Regions mediating the associations between education scores and Pattern Comparison Processing Speed scores at baseline**

The table summarises the mediation analyses ran across regions that showed both environment-brain and brain-behaviour associations. The ACME (Average Causal Mediation Effect) estimate is reported, which reflects the indirect effect of the predictor on the outcome that is transmitted through the mediator, as well as proportion mediated values, which indicate the estimated fraction of the total effect explained by the mediator. Associated ACME confidence intervals and FDR-corrected p values are also reported. P values are false discovery rate (FDR) corrected to account for multiple comparisons. Analyses controlled for age and sex and included random intercepts for testing site and family ID (to account for sibling pairs).

| **Region** | **ACME estimate** | **ACME CI** | **Proportion mediated** | ***p*_FDR_** |
| --- | --- | --- | --- | --- |
| Left Banks of the Superior Temporal Sulcus | -0.004 | -0.024 - 0.017 | -0.003 | 0.861 |
| Left Caudal Anterior Cingulate Cortex | 0.015 | -0.004 - 0.035 | 0.014 | 0.733 |
| Left Caudal Middle Frontal Gyrus | 0.013 | -0.009 - 0.038 | 0.012 | 0.733 |
| Left Frontal Pole | -0.008 | -0.031 - 0.013 | -0.007 | 0.733 |
| Left Inferior Parietal Lobule | -0.007 | -0.028 - 0.012 | -0.007 | 0.733 |
| Left Isthmus of the Cingulate Cortex | 0.011 | -0.015 - 0.033 | 0.010 | 0.733 |
| Left Lateral Orbitofrontal Cortex | 0.013 | -0.025 - 0.050 | 0.012 | 0.733 |
| Left Lingual Gyrus | 0.013 | -0.016 - 0.041 | 0.012 | 0.733 |
| Left Medial Orbitofrontal Cortex | -0.007 | -0.027 - 0.014 | -0.007 | 0.733 |
| Left Paracentral Lobule | 0.001 | -0.019 - 0.023 | 0.001 | 0.903 |
| Left Parahippocampal Gyrus | 0.016 | -0.010 - 0.044 | 0.015 | 0.733 |
| Left Pars Opercularis | 0.008 | -0.014 - 0.031 | 0.008 | 0.733 |
| Left Pars Orbitalis | 0.017 | -0.010 - 0.046 | 0.016 | 0.733 |
| Left Pars Triangularis | 0.006 | -0.010 - 0.023 | 0.005 | 0.733 |
| Left Pericalcarine Cortex | 0.008 | -0.011 - 0.027 | 0.007 | 0.733 |
| Left Postcentral Gyrus | 0.001 | -0.039 - 0.037 | 0.001 | 0.982 |
| Left Precuneus | 0.017 | -0.016 - 0.050 | 0.015 | 0.733 |
| Left Rostral Middle Frontal Gyrus | -0.011 | -0.045 - 0.021 | -0.010 | 0.733 |
| Left Superior Frontal Gyrus | 0.008 | -0.020 - 0.034 | 0.007 | 0.733 |
| Left Superior Parietal Lobule | 0.014 | -0.021 - 0.047 | 0.013 | 0.733 |
| Left Temporal Pole | 0.013 | -0.008 - 0.034 | 0.012 | 0.733 |
| Left Transverse Temporal Gyrus (Heschl’s Gyrus) | 0.015 | -0.008 - 0.041 | 0.014 | 0.733 |
| Right Banks of the Superior Temporal Sulcus | -0.007 | -0.029 - 0.014 | -0.006 | 0.733 |
| Right Caudal Anterior Cingulate Cortex | -0.002 | -0.017 - 0.014 | -0.002 | 0.903 |
| Right Cuneus | 0.013 | -0.015 - 0.041 | 0.012 | 0.733 |
| Right Frontal Pole | -0.022 | -0.054 - 0.005 | -0.020 | 0.733 |
| Right Fusiform Gyrus | 0.014 | -0.020 - 0.050 | 0.013 | 0.733 |
| Right Inferior Parietal Lobule | 0.014 | -0.011 - 0.042 | 0.013 | 0.733 |
| Right Isthmus of the Cingulate Cortex | 0.004 | -0.010 - 0.020 | 0.004 | 0.733 |
| Right Lingual Gyrus | 0.011 | -0.012 - 0.035 | 0.010 | 0.733 |
| Right Medial Orbitofrontal Cortex | 0.013 | -0.017 - 0.043 | 0.012 | 0.733 |
| Right Middle Temporal Gyrus | 0.004 | -0.038 - 0.043 | 0.003 | 0.903 |
| Right Parahippocampal Gyrus | 0.015 | -0.013 - 0.043 | 0.013 | 0.733 |
| Right Pars Opercularis | 0.004 | -0.016 - 0.023 | 0.004 | 0.813 |
| Right Pars Orbitalis | -0.002 | -0.031 - 0.027 | -0.001 | 0.955 |
| Right Pars Triangularis | 0.004 | -0.008 - 0.020 | 0.004 | 0.733 |
| Right Pericalcarine Cortex | 0.010 | -0.008 - 0.028 | 0.010 | 0.733 |
| Right Posterior Cingulate Cortex | 0.005 | -0.013 - 0.026 | 0.005 | 0.733 |
| Right Precuneus | -0.007 | -0.041 - 0.026 | -0.006 | 0.813 |
| Right Rostral Anterior Cingulate Cortex | 0.008 | -0.007 - 0.026 | 0.008 | 0.733 |
| Right Rostral Middle Frontal Gyrus | -0.015 | -0.040 - 0.012 | -0.014 | 0.733 |
| Right Superior Frontal Gyrus | 0.012 | -0.018 - 0.043 | 0.011 | 0.733 |
| Right Superior Parietal Lobule | 0.001 | -0.030 - 0.034 | 0.001 | 0.920 |
| Right Superior Temporal Gyrus | 0.009 | -0.024 - 0.040 | 0.008 | 0.733 |
| Right Supramarginal Gyrus | 0.004 | -0.021 - 0.028 | 0.003 | 0.867 |
| Right Transverse Temporal Gyrus (Heschl’s Gyrus) | 0.011 | -0.012 - 0.036 | 0.010 | 0.733 |

**Supplementary Table 27: Regions mediating the associations between micro-environment scores and Pattern Comparison Processing Speed scores at baseline**

The table summarises the mediation analyses ran across regions that showed both environment-brain and brain-behaviour associations. The ACME (Average Causal Mediation Effect) estimate is reported, which reflects the indirect effect of the predictor on the outcome that is transmitted through the mediator, as well as proportion mediated values, which indicate the estimated fraction of the total effect explained by the mediator. Associated ACME confidence intervals and FDR-corrected p values are also reported. P values are false discovery rate (FDR) corrected to account for multiple comparisons. Analyses controlled for age and sex and included random intercepts for testing site and family ID (to account for sibling pairs).

| **Region** | **ACME estimate** | **ACME CI** | **Proportion mediated** | ***p*_FDR_** |
| --- | --- | --- | --- | --- |
| Left Cuneus | 0.028 | -0.016 - 0.073 | 0.015 | 0.449 |
| Left Entorhinal Cortex | 0.060 | 0.007 - 0.114 | 0.032 | 0.215 |
| Left Fusiform Gyrus | 0.023 | -0.042 - 0.090 | 0.012 | 0.510 |
| Left Inferior Temporal Gyrus | 0.029 | -0.035 - 0.091 | 0.015 | 0.449 |
| Left Insular Cortex | 0.066 | 0.022 - 0.115 | 0.035 | 0.138 |
| Left Lateral Occipital Cortex | 0.040 | -0.017 - 0.095 | 0.021 | 0.429 |
| Left Middle Temporal Gyrus | 0.037 | -0.032 - 0.106 | 0.020 | 0.449 |
| Left Posterior Cingulate Cortex | 0.021 | -0.017 - 0.060 | 0.011 | 0.449 |
| Left Precentral Gyrus | 0.032 | -0.033 - 0.102 | 0.017 | 0.449 |
| Left Rostral Anterior Cingulate Cortex | 0.024 | -0.024 - 0.076 | 0.013 | 0.449 |
| Left Superior Temporal Gyrus | 0.025 | -0.026 - 0.080 | 0.013 | 0.449 |
| Left Supramarginal Gyrus | 0.029 | -0.025 - 0.086 | 0.015 | 0.449 |
| Right Caudal Middle Frontal Gyrus | 0.043 | -0.008 - 0.100 | 0.023 | 0.429 |
| Right Entorhinal Cortex | 0.037 | -0.012 - 0.090 | 0.020 | 0.429 |
| Right Inferior Temporal Gyrus | 0.040 | -0.026 - 0.110 | 0.021 | 0.449 |
| Right Insular Cortex | 0.060 | 0.011 - 0.110 | 0.032 | 0.138 |
| Right Lateral Occipital Cortex | 0.023 | -0.042 - 0.089 | 0.012 | 0.449 |
| Right Lateral Orbitofrontal Cortex | 0.034 | -0.024 - 0.096 | 0.018 | 0.449 |
| Right Paracentral Lobule | 0.025 | -0.010 - 0.068 | 0.013 | 0.429 |
| Right Postcentral Gyrus | 0.021 | -0.048 - 0.088 | 0.011 | 0.572 |
| Right Precentral Gyrus | 0.051 | -0.022 - 0.121 | 0.027 | 0.429 |
| Right Temporal Pole | 0.022 | -0.003 - 0.051 | 0.012 | 0.429 |

**Supplementary Table 28: Regions mediating the associations between COI 2.0 health/environment scores and Picture Vocabulary scores at baseline**

The table summarises the mediation analyses ran across regions that showed both environment-brain and brain-behaviour associations. The ACME (Average Causal Mediation Effect) estimate is reported, which reflects the indirect effect of the predictor on the outcome that is transmitted through the mediator, as well as proportion mediated values, which indicate the estimated fraction of the total effect explained by the mediator. Associated ACME confidence intervals and FDR-corrected p values are also reported. P values are false discovery rate (FDR) corrected to account for multiple comparisons. Analyses controlled for age and sex and included random intercepts for testing site and family ID (to account for sibling pairs).

| **Region** | **ACME estimate** | **ACME CI** | **Proportion mediated** | ***p*_FDR_** |
| --- | --- | --- | --- | --- |
| Left Banks of the Superior Temporal Sulcus | 0.031 | 0.018 - 0.045 | 0.022 | 0.000 |
| Left Caudal Anterior Cingulate Cortex | 0.020 | 0.009 - 0.032 | 0.014 | 0.000 |
| Left Caudal Middle Frontal Gyrus | 0.055 | 0.037 - 0.074 | 0.038 | 0.000 |
| Left Cuneus | 0.030 | 0.018 - 0.044 | 0.021 | 0.000 |
| Left Entorhinal Cortex | 0.046 | 0.029 - 0.064 | 0.032 | 0.000 |
| Left Frontal Pole | 0.025 | 0.014 - 0.038 | 0.018 | 0.000 |
| Left Fusiform Gyrus | 0.082 | 0.060 - 0.105 | 0.057 | 0.000 |
| Left Inferior Parietal Lobule | 0.030 | 0.016 - 0.043 | 0.021 | 0.000 |
| Left Inferior Temporal Gyrus | 0.091 | 0.068 - 0.115 | 0.063 | 0.000 |
| Left Insular Cortex | 0.054 | 0.036 - 0.075 | 0.038 | 0.000 |
| Left Isthmus of the Cingulate Cortex | 0.044 | 0.028 - 0.061 | 0.031 | 0.000 |
| Left Lateral Occipital Cortex | 0.068 | 0.050 - 0.090 | 0.047 | 0.000 |
| Left Lateral Orbitofrontal Cortex | 0.085 | 0.062 - 0.112 | 0.059 | 0.000 |
| Left Lingual Gyrus | 0.039 | 0.025 - 0.056 | 0.027 | 0.000 |
| Left Medial Orbitofrontal Cortex | 0.032 | 0.016 - 0.050 | 0.022 | 0.000 |
| Left Middle Temporal Gyrus | 0.093 | 0.070 - 0.119 | 0.065 | 0.000 |
| Left Paracentral Lobule | 0.036 | 0.022 - 0.052 | 0.025 | 0.000 |
| Left Parahippocampal Gyrus | 0.039 | 0.023 - 0.057 | 0.027 | 0.000 |
| Left Pars Opercularis | 0.034 | 0.021 - 0.050 | 0.023 | 0.000 |
| Left Pars Orbitalis | 0.058 | 0.041 - 0.077 | 0.040 | 0.000 |
| Left Pars Triangularis | 0.015 | 0.006 - 0.026 | 0.010 | 0.000 |
| Left Pericalcarine Cortex | 0.017 | 0.007 - 0.028 | 0.012 | 0.000 |
| Left Postcentral Gyrus | 0.085 | 0.065 - 0.109 | 0.059 | 0.000 |
| Left Posterior Cingulate Cortex | 0.038 | 0.023 - 0.055 | 0.026 | 0.000 |
| Left Precentral Gyrus | 0.095 | 0.072 - 0.117 | 0.066 | 0.000 |
| Left Precuneus | 0.067 | 0.048 - 0.087 | 0.047 | 0.000 |
| Left Rostral Anterior Cingulate Cortex | 0.036 | 0.020 - 0.053 | 0.025 | 0.000 |
| Left Rostral Middle Frontal Gyrus | 0.059 | 0.040 - 0.077 | 0.041 | 0.000 |
| Left Superior Frontal Gyrus | 0.067 | 0.047 - 0.088 | 0.047 | 0.000 |
| Left Superior Parietal Lobule | 0.054 | 0.037 - 0.073 | 0.038 | 0.000 |
| Left Superior Temporal Gyrus | 0.057 | 0.037 - 0.078 | 0.039 | 0.000 |
| Left Supramarginal Gyrus | 0.051 | 0.034 - 0.070 | 0.035 | 0.000 |
| Left Temporal Pole | 0.023 | 0.011 - 0.037 | 0.016 | 0.000 |
| Left Transverse Temporal Gyrus (Heschl’s Gyrus) | 0.042 | 0.026 - 0.058 | 0.029 | 0.000 |
| Right Banks of the Superior Temporal Sulcus | 0.041 | 0.026 - 0.059 | 0.028 | 0.000 |
| Right Caudal Anterior Cingulate Cortex | 0.023 | 0.012 - 0.037 | 0.016 | 0.000 |
| Right Caudal Middle Frontal Gyrus | 0.048 | 0.032 - 0.067 | 0.034 | 0.000 |
| Right Cuneus | 0.034 | 0.021 - 0.048 | 0.024 | 0.000 |
| Right Entorhinal Cortex | 0.034 | 0.020 - 0.051 | 0.024 | 0.000 |
| Right Frontal Pole | 0.038 | 0.024 - 0.054 | 0.026 | 0.000 |
| Right Fusiform Gyrus | 0.077 | 0.054 - 0.100 | 0.054 | 0.000 |
| Right Inferior Parietal Lobule | 0.052 | 0.034 - 0.073 | 0.036 | 0.000 |
| Right Inferior Temporal Gyrus | 0.080 | 0.060 - 0.105 | 0.056 | 0.000 |
| Right Insular Cortex | 0.044 | 0.028 - 0.062 | 0.030 | 0.000 |
| Right Isthmus of the Cingulate Cortex | 0.021 | 0.009 - 0.035 | 0.015 | 0.002 |
| Right Lateral Occipital Cortex | 0.089 | 0.066 - 0.111 | 0.062 | 0.000 |
| Right Lateral Orbitofrontal Cortex | 0.085 | 0.063 - 0.108 | 0.059 | 0.000 |
| Right Lingual Gyrus | 0.031 | 0.018 - 0.047 | 0.021 | 0.000 |
| Right Medial Orbitofrontal Cortex | 0.054 | 0.034 - 0.074 | 0.037 | 0.000 |
| Right Middle Temporal Gyrus | 0.122 | 0.095 - 0.151 | 0.085 | 0.000 |
| Right Paracentral Lobule | 0.026 | 0.015 - 0.039 | 0.018 | 0.000 |
| Right Parahippocampal Gyrus | 0.041 | 0.024 - 0.059 | 0.028 | 0.000 |
| Right Pars Opercularis | 0.037 | 0.023 - 0.053 | 0.026 | 0.000 |
| Right Pars Orbitalis | 0.061 | 0.043 - 0.080 | 0.042 | 0.000 |
| Right Pars Triangularis | 0.015 | 0.007 - 0.026 | 0.010 | 0.000 |
| Right Pericalcarine Cortex | 0.013 | 0.004 - 0.024 | 0.009 | 0.000 |
| Right Postcentral Gyrus | 0.065 | 0.048 - 0.086 | 0.045 | 0.000 |
| Right Posterior Cingulate Cortex | 0.035 | 0.021 - 0.052 | 0.024 | 0.000 |
| Right Precentral Gyrus | 0.077 | 0.056 - 0.101 | 0.054 | 0.000 |
| Right Precuneus | 0.075 | 0.055 - 0.096 | 0.052 | 0.000 |
| Right Rostral Anterior Cingulate Cortex | 0.032 | 0.016 - 0.048 | 0.022 | 0.000 |
| Right Rostral Middle Frontal Gyrus | 0.046 | 0.030 - 0.063 | 0.032 | 0.000 |
| Right Superior Frontal Gyrus | 0.072 | 0.051 - 0.095 | 0.050 | 0.000 |
| Right Superior Parietal Lobule | 0.058 | 0.041 - 0.077 | 0.040 | 0.000 |
| Right Superior Temporal Gyrus | 0.070 | 0.049 - 0.094 | 0.049 | 0.000 |
| Right Supramarginal Gyrus | 0.038 | 0.023 - 0.055 | 0.027 | 0.000 |
| Right Temporal Pole | 0.014 | 0.002 - 0.026 | 0.010 | 0.014 |
| Right Transverse Temporal Gyrus (Heschl’s Gyrus) | 0.050 | 0.034 - 0.066 | 0.034 | 0.000 |

**Supplementary Table 29: Regions mediating the associations between COI 2.0 social/economic scores and Picture Vocabulary scores at baseline**

The table summarises the mediation analyses ran across regions that showed both environment-brain and brain-behaviour associations. The ACME (Average Causal Mediation Effect) estimate is reported, which reflects the indirect effect of the predictor on the outcome that is transmitted through the mediator, as well as proportion mediated values, which indicate the estimated fraction of the total effect explained by the mediator. Associated ACME confidence intervals and FDR-corrected p values are also reported. P values are false discovery rate (FDR) corrected to account for multiple comparisons. Analyses controlled for age and sex and included random intercepts for testing site and family ID (to account for sibling pairs).

| **Region** | **ACME estimate** | **ACME CI** | **Proportion mediated** | ***p*_FDR_** |
| --- | --- | --- | --- | --- |
| Left Banks of the Superior Temporal Sulcus | 0.034 | 0.021 - 0.051 | 0.019 | <0.001 |
| Left Caudal Anterior Cingulate Cortex | 0.031 | 0.017 - 0.044 | 0.017 | <0.001 |
| Left Caudal Middle Frontal Gyrus | 0.065 | 0.048 - 0.084 | 0.036 | <0.001 |
| Left Cuneus | 0.032 | 0.018 - 0.048 | 0.017 | <0.001 |
| Left Entorhinal Cortex | 0.059 | 0.040 - 0.077 | 0.032 | <0.001 |
| Left Frontal Pole | 0.028 | 0.016 - 0.042 | 0.015 | <0.001 |
| Left Fusiform Gyrus | 0.111 | 0.086 - 0.136 | 0.060 | <0.001 |
| Left Inferior Parietal Lobule | 0.034 | 0.021 - 0.049 | 0.019 | <0.001 |
| Left Inferior Temporal Gyrus | 0.108 | 0.086 - 0.133 | 0.059 | <0.001 |
| Left Insular Cortex | 0.070 | 0.052 - 0.090 | 0.038 | <0.001 |
| Left Isthmus of the Cingulate Cortex | 0.054 | 0.038 - 0.073 | 0.029 | <0.001 |
| Left Lateral Occipital Cortex | 0.082 | 0.063 - 0.105 | 0.045 | <0.001 |
| Left Lateral Orbitofrontal Cortex | 0.115 | 0.091 - 0.140 | 0.063 | <0.001 |
| Left Lingual Gyrus | 0.051 | 0.034 - 0.069 | 0.028 | <0.001 |
| Left Medial Orbitofrontal Cortex | 0.043 | 0.029 - 0.061 | 0.024 | <0.001 |
| Left Middle Temporal Gyrus | 0.112 | 0.088 - 0.136 | 0.061 | <0.001 |
| Left Paracentral Lobule | 0.045 | 0.030 - 0.063 | 0.025 | <0.001 |
| Left Parahippocampal Gyrus | 0.053 | 0.037 - 0.069 | 0.029 | <0.001 |
| Left Pars Opercularis | 0.037 | 0.023 - 0.053 | 0.020 | <0.001 |
| Left Pars Orbitalis | 0.067 | 0.048 - 0.088 | 0.036 | <0.001 |
| Left Pars Triangularis | 0.018 | 0.009 - 0.029 | 0.010 | <0.001 |
| Left Pericalcarine Cortex | 0.020 | 0.010 - 0.032 | 0.011 | <0.001 |
| Left Postcentral Gyrus | 0.098 | 0.076 - 0.124 | 0.054 | <0.001 |
| Left Posterior Cingulate Cortex | 0.041 | 0.027 - 0.057 | 0.023 | <0.001 |
| Left Precentral Gyrus | 0.114 | 0.091 - 0.142 | 0.062 | <0.001 |
| Left Precuneus | 0.079 | 0.057 - 0.102 | 0.043 | <0.001 |
| Left Rostral Anterior Cingulate Cortex | 0.054 | 0.038 - 0.070 | 0.029 | <0.001 |
| Left Rostral Middle Frontal Gyrus | 0.076 | 0.057 - 0.099 | 0.042 | <0.001 |
| Left Superior Frontal Gyrus | 0.079 | 0.059 - 0.102 | 0.043 | <0.001 |
| Left Superior Parietal Lobule | 0.067 | 0.047 - 0.087 | 0.037 | <0.001 |
| Left Superior Temporal Gyrus | 0.087 | 0.065 - 0.109 | 0.047 | <0.001 |
| Left Supramarginal Gyrus | 0.070 | 0.052 - 0.090 | 0.038 | <0.001 |
| Left Temporal Pole | 0.033 | 0.021 - 0.047 | 0.018 | <0.001 |
| Left Transverse Temporal Gyrus (Heschl’s Gyrus) | 0.055 | 0.040 - 0.075 | 0.030 | <0.001 |
| Right Banks of the Superior Temporal Sulcus | 0.050 | 0.035 - 0.066 | 0.027 | <0.001 |
| Right Caudal Anterior Cingulate Cortex | 0.027 | 0.016 - 0.041 | 0.015 | <0.001 |
| Right Caudal Middle Frontal Gyrus | 0.061 | 0.044 - 0.081 | 0.033 | <0.001 |
| Right Cuneus | 0.038 | 0.021 - 0.054 | 0.021 | <0.001 |
| Right Entorhinal Cortex | 0.037 | 0.022 - 0.054 | 0.020 | <0.001 |
| Right Frontal Pole | 0.045 | 0.030 - 0.061 | 0.024 | <0.001 |
| Right Fusiform Gyrus | 0.106 | 0.083 - 0.133 | 0.058 | <0.001 |
| Right Inferior Parietal Lobule | 0.062 | 0.044 - 0.082 | 0.034 | <0.001 |
| Right Inferior Temporal Gyrus | 0.102 | 0.079 - 0.127 | 0.056 | <0.001 |
| Right Insular Cortex | 0.056 | 0.039 - 0.076 | 0.030 | <0.001 |
| Right Isthmus of the Cingulate Cortex | 0.027 | 0.016 - 0.041 | 0.015 | <0.001 |
| Right Lateral Occipital Cortex | 0.107 | 0.083 - 0.133 | 0.058 | <0.001 |
| Right Lateral Orbitofrontal Cortex | 0.106 | 0.083 - 0.130 | 0.058 | <0.001 |
| Right Lingual Gyrus | 0.039 | 0.025 - 0.056 | 0.021 | <0.001 |
| Right Medial Orbitofrontal Cortex | 0.073 | 0.053 - 0.094 | 0.040 | <0.001 |
| Right Middle Temporal Gyrus | 0.141 | 0.115 - 0.168 | 0.077 | <0.001 |
| Right Paracentral Lobule | 0.030 | 0.018 - 0.043 | 0.016 | <0.001 |
| Right Parahippocampal Gyrus | 0.051 | 0.035 - 0.071 | 0.028 | <0.001 |
| Right Pars Opercularis | 0.042 | 0.026 - 0.057 | 0.023 | <0.001 |
| Right Pars Orbitalis | 0.068 | 0.049 - 0.090 | 0.037 | <0.001 |
| Right Pars Triangularis | 0.015 | 0.007 - 0.025 | 0.008 | <0.001 |
| Right Pericalcarine Cortex | 0.018 | 0.009 - 0.030 | 0.010 | <0.001 |
| Right Postcentral Gyrus | 0.077 | 0.058 - 0.099 | 0.042 | <0.001 |
| Right Posterior Cingulate Cortex | 0.045 | 0.030 - 0.062 | 0.025 | <0.001 |
| Right Precentral Gyrus | 0.098 | 0.074 - 0.119 | 0.053 | <0.001 |
| Right Precuneus | 0.086 | 0.065 - 0.110 | 0.047 | <0.001 |
| Right Rostral Anterior Cingulate Cortex | 0.043 | 0.029 - 0.061 | 0.024 | <0.001 |
| Right Rostral Middle Frontal Gyrus | 0.058 | 0.040 - 0.078 | 0.032 | <0.001 |
| Right Superior Frontal Gyrus | 0.096 | 0.073 - 0.120 | 0.052 | <0.001 |
| Right Superior Parietal Lobule | 0.071 | 0.053 - 0.091 | 0.039 | <0.001 |
| Right Superior Temporal Gyrus | 0.093 | 0.070 - 0.117 | 0.051 | <0.001 |
| Right Supramarginal Gyrus | 0.048 | 0.033 - 0.065 | 0.026 | <0.001 |
| Right Temporal Pole | 0.022 | 0.012 - 0.035 | 0.012 | <0.001 |
| Right Transverse Temporal Gyrus (Heschl’s Gyrus) | 0.057 | 0.040 - 0.076 | 0.031 | <0.001 |

**Supplementary Table 30: Regions mediating the associations between COI 2.0 education scores and Picture Vocabulary scores at baseline**

The table summarises the mediation analyses ran across regions that showed both environment-brain and brain-behaviour associations. The ACME (Average Causal Mediation Effect) estimate is reported, which reflects the indirect effect of the predictor on the outcome that is transmitted through the mediator, as well as proportion mediated values, which indicate the estimated fraction of the total effect explained by the mediator. Associated ACME confidence intervals and FDR-corrected p values are also reported. P values are false discovery rate (FDR) corrected to account for multiple comparisons. Analyses controlled for age and sex and included random intercepts for testing site and family ID (to account for sibling pairs).

| **Region** | **ACME estimate** | **ACME CI** | **Proportion mediated** | ***p*_FDR_** |
| --- | --- | --- | --- | --- |
| Left Banks of the Superior Temporal Sulcus | 0.037 | 0.023 - 0.054 | 0.022 | 0.000 |
| Left Caudal Anterior Cingulate Cortex | 0.029 | 0.017 - 0.044 | 0.017 | 0.000 |
| Left Caudal Middle Frontal Gyrus | 0.061 | 0.044 - 0.081 | 0.036 | 0.000 |
| Left Cuneus | 0.035 | 0.020 - 0.051 | 0.021 | 0.000 |
| Left Entorhinal Cortex | 0.062 | 0.045 - 0.083 | 0.037 | 0.000 |
| Left Frontal Pole | 0.031 | 0.017 - 0.045 | 0.018 | 0.000 |
| Left Fusiform Gyrus | 0.118 | 0.094 - 0.145 | 0.071 | 0.000 |
| Left Inferior Parietal Lobule | 0.039 | 0.024 - 0.055 | 0.023 | 0.000 |
| Left Inferior Temporal Gyrus | 0.116 | 0.092 - 0.144 | 0.069 | 0.000 |
| Left Insular Cortex | 0.075 | 0.056 - 0.094 | 0.045 | 0.000 |
| Left Isthmus of the Cingulate Cortex | 0.058 | 0.041 - 0.078 | 0.035 | 0.000 |
| Left Lateral Occipital Cortex | 0.087 | 0.068 - 0.108 | 0.052 | 0.000 |
| Left Lateral Orbitofrontal Cortex | 0.120 | 0.094 - 0.147 | 0.071 | 0.000 |
| Left Lingual Gyrus | 0.055 | 0.037 - 0.075 | 0.033 | 0.000 |
| Left Medial Orbitofrontal Cortex | 0.049 | 0.032 - 0.067 | 0.029 | 0.000 |
| Left Middle Temporal Gyrus | 0.117 | 0.093 - 0.144 | 0.070 | 0.000 |
| Left Paracentral Lobule | 0.042 | 0.027 - 0.058 | 0.025 | 0.000 |
| Left Parahippocampal Gyrus | 0.058 | 0.040 - 0.079 | 0.035 | 0.000 |
| Left Pars Opercularis | 0.040 | 0.024 - 0.055 | 0.024 | 0.000 |
| Left Pars Orbitalis | 0.070 | 0.051 - 0.091 | 0.042 | 0.000 |
| Left Pars Triangularis | 0.021 | 0.011 - 0.033 | 0.013 | 0.000 |
| Left Pericalcarine Cortex | 0.022 | 0.011 - 0.035 | 0.013 | 0.000 |
| Left Postcentral Gyrus | 0.103 | 0.079 - 0.129 | 0.061 | 0.000 |
| Left Posterior Cingulate Cortex | 0.045 | 0.031 - 0.062 | 0.027 | 0.000 |
| Left Precentral Gyrus | 0.112 | 0.088 - 0.138 | 0.067 | 0.000 |
| Left Precuneus | 0.082 | 0.059 - 0.104 | 0.049 | 0.000 |
| Left Rostral Anterior Cingulate Cortex | 0.050 | 0.035 - 0.068 | 0.030 | 0.000 |
| Left Rostral Middle Frontal Gyrus | 0.084 | 0.061 - 0.108 | 0.050 | 0.000 |
| Left Superior Frontal Gyrus | 0.078 | 0.056 - 0.101 | 0.047 | 0.000 |
| Left Superior Parietal Lobule | 0.072 | 0.053 - 0.094 | 0.043 | 0.000 |
| Left Superior Temporal Gyrus | 0.090 | 0.066 - 0.113 | 0.054 | 0.000 |
| Left Supramarginal Gyrus | 0.071 | 0.054 - 0.093 | 0.043 | 0.000 |
| Left Temporal Pole | 0.034 | 0.022 - 0.049 | 0.020 | 0.000 |
| Left Transverse Temporal Gyrus (Heschl’s Gyrus) | 0.056 | 0.040 - 0.075 | 0.034 | 0.000 |
| Right Banks of the Superior Temporal Sulcus | 0.051 | 0.035 - 0.069 | 0.030 | 0.000 |
| Right Caudal Anterior Cingulate Cortex | 0.026 | 0.014 - 0.041 | 0.016 | 0.000 |
| Right Caudal Middle Frontal Gyrus | 0.061 | 0.043 - 0.082 | 0.036 | 0.000 |
| Right Cuneus | 0.041 | 0.023 - 0.059 | 0.024 | 0.000 |
| Right Entorhinal Cortex | 0.038 | 0.023 - 0.055 | 0.023 | 0.000 |
| Right Frontal Pole | 0.051 | 0.033 - 0.070 | 0.030 | 0.000 |
| Right Fusiform Gyrus | 0.115 | 0.091 - 0.143 | 0.069 | 0.000 |
| Right Inferior Parietal Lobule | 0.063 | 0.045 - 0.082 | 0.037 | 0.000 |
| Right Inferior Temporal Gyrus | 0.103 | 0.079 - 0.128 | 0.061 | 0.000 |
| Right Insular Cortex | 0.059 | 0.041 - 0.079 | 0.035 | 0.000 |
| Right Isthmus of the Cingulate Cortex | 0.028 | 0.016 - 0.042 | 0.017 | 0.000 |
| Right Lateral Occipital Cortex | 0.111 | 0.087 - 0.139 | 0.066 | 0.000 |
| Right Lateral Orbitofrontal Cortex | 0.110 | 0.087 - 0.135 | 0.065 | 0.000 |
| Right Lingual Gyrus | 0.044 | 0.029 - 0.062 | 0.026 | 0.000 |
| Right Medial Orbitofrontal Cortex | 0.080 | 0.061 - 0.101 | 0.048 | 0.000 |
| Right Middle Temporal Gyrus | 0.152 | 0.121 - 0.180 | 0.090 | 0.000 |
| Right Paracentral Lobule | 0.031 | 0.019 - 0.044 | 0.018 | 0.000 |
| Right Parahippocampal Gyrus | 0.060 | 0.042 - 0.081 | 0.036 | 0.000 |
| Right Pars Opercularis | 0.039 | 0.025 - 0.055 | 0.023 | 0.000 |
| Right Pars Orbitalis | 0.071 | 0.053 - 0.092 | 0.042 | 0.000 |
| Right Pars Triangularis | 0.016 | 0.008 - 0.027 | 0.010 | 0.000 |
| Right Pericalcarine Cortex | 0.020 | 0.010 - 0.032 | 0.012 | 0.000 |
| Right Postcentral Gyrus | 0.083 | 0.063 - 0.106 | 0.049 | 0.000 |
| Right Posterior Cingulate Cortex | 0.041 | 0.027 - 0.058 | 0.025 | 0.000 |
| Right Precentral Gyrus | 0.096 | 0.073 - 0.123 | 0.058 | 0.000 |
| Right Precuneus | 0.089 | 0.067 - 0.112 | 0.053 | 0.000 |
| Right Rostral Anterior Cingulate Cortex | 0.039 | 0.024 - 0.055 | 0.023 | 0.000 |
| Right Rostral Middle Frontal Gyrus | 0.059 | 0.040 - 0.077 | 0.035 | 0.000 |
| Right Superior Frontal Gyrus | 0.094 | 0.073 - 0.119 | 0.056 | 0.000 |
| Right Superior Parietal Lobule | 0.075 | 0.054 - 0.097 | 0.045 | 0.000 |
| Right Superior Temporal Gyrus | 0.101 | 0.078 - 0.126 | 0.060 | 0.000 |
| Right Supramarginal Gyrus | 0.055 | 0.037 - 0.073 | 0.033 | 0.000 |
| Right Temporal Pole | 0.024 | 0.012 - 0.036 | 0.014 | 0.000 |
| Right Transverse Temporal Gyrus (Heschl’s Gyrus) | 0.058 | 0.041 - 0.077 | 0.034 | 0.000 |

**Supplementary Table 31: Regions mediating the associations between micro-environment scores and Picture Vocabulary scores at baseline**

The table summarises the mediation analyses ran across regions that showed both environment-brain and brain-behaviour associations. The ACME (Average Causal Mediation Effect) estimate is reported, which reflects the indirect effect of the predictor on the outcome that is transmitted through the mediator, as well as proportion mediated values, which indicate the estimated fraction of the total effect explained by the mediator. Associated ACME confidence intervals and FDR-corrected p values are also reported. P values are false discovery rate (FDR) corrected to account for multiple comparisons. Analyses controlled for age and sex and included random intercepts for testing site and family ID (to account for sibling pairs).

| **Region** | **Proportion mediated** | **95% confidence interval** | ***p*_FDR_** |
| --- | --- | --- | --- |
| Left Banks of the Superior Temporal Sulcus | 0.022 | 0.014 - 0.031 | 0.000 |
| Left Caudal Anterior Cingulate Cortex | 0.017 | 0.010 - 0.025 | 0.000 |
| Left Caudal Middle Frontal Gyrus | 0.035 | 0.026 - 0.046 | 0.000 |
| Left Cuneus | 0.017 | 0.008 - 0.025 | 0.000 |
| Left Entorhinal Cortex | 0.034 | 0.024 - 0.045 | 0.000 |
| Left Frontal Pole | 0.016 | 0.009 - 0.024 | 0.000 |
| Left Fusiform Gyrus | 0.065 | 0.051 - 0.080 | 0.000 |
| Left Inferior Parietal Lobule | 0.024 | 0.015 - 0.033 | 0.000 |
| Left Inferior Temporal Gyrus | 0.061 | 0.048 - 0.074 | 0.000 |
| Left Insular Cortex | 0.040 | 0.030 - 0.051 | 0.000 |
| Left Isthmus of the Cingulate Cortex | 0.030 | 0.021 - 0.041 | 0.000 |
| Left Lateral Occipital Cortex | 0.043 | 0.031 - 0.056 | 0.000 |
| Left Lateral Orbitofrontal Cortex | 0.064 | 0.050 - 0.079 | 0.000 |
| Left Lingual Gyrus | 0.028 | 0.019 - 0.040 | 0.000 |
| Left Medial Orbitofrontal Cortex | 0.029 | 0.020 - 0.039 | 0.000 |
| Left Middle Temporal Gyrus | 0.066 | 0.051 - 0.081 | 0.000 |
| Left Paracentral Lobule | 0.027 | 0.017 - 0.039 | 0.000 |
| Left Parahippocampal Gyrus | 0.030 | 0.020 - 0.041 | 0.000 |
| Left Pars Opercularis | 0.020 | 0.013 - 0.028 | 0.000 |
| Left Pars Orbitalis | 0.039 | 0.028 - 0.051 | 0.000 |
| Left Pars Triangularis | 0.011 | 0.006 - 0.017 | 0.000 |
| Left Pericalcarine Cortex | 0.011 | 0.005 - 0.018 | 0.000 |
| Left Postcentral Gyrus | 0.056 | 0.042 - 0.071 | 0.000 |
| Left Posterior Cingulate Cortex | 0.028 | 0.019 - 0.038 | 0.000 |
| Left Precentral Gyrus | 0.065 | 0.052 - 0.081 | 0.000 |
| Left Precuneus | 0.043 | 0.032 - 0.056 | 0.000 |
| Left Rostral Anterior Cingulate Cortex | 0.033 | 0.024 - 0.043 | 0.000 |
| Left Rostral Middle Frontal Gyrus | 0.046 | 0.034 - 0.058 | 0.000 |
| Left Superior Frontal Gyrus | 0.049 | 0.037 - 0.062 | 0.000 |
| Left Superior Parietal Lobule | 0.037 | 0.026 - 0.049 | 0.000 |
| Left Superior Temporal Gyrus | 0.053 | 0.041 - 0.066 | 0.000 |
| Left Supramarginal Gyrus | 0.040 | 0.029 - 0.051 | 0.000 |
| Left Temporal Pole | 0.019 | 0.012 - 0.027 | 0.000 |
| Left Transverse Temporal Gyrus (Heschl’s Gyrus) | 0.030 | 0.021 - 0.040 | 0.000 |
| Right Banks of the Superior Temporal Sulcus | 0.032 | 0.022 - 0.042 | 0.000 |
| Right Caudal Anterior Cingulate Cortex | 0.015 | 0.009 - 0.023 | 0.000 |
| Right Caudal Middle Frontal Gyrus | 0.038 | 0.027 - 0.050 | 0.000 |
| Right Cuneus | 0.019 | 0.011 - 0.027 | 0.000 |
| Right Entorhinal Cortex | 0.020 | 0.011 - 0.030 | 0.000 |
| Right Frontal Pole | 0.025 | 0.016 - 0.034 | 0.000 |
| Right Fusiform Gyrus | 0.063 | 0.050 - 0.079 | 0.000 |
| Right Inferior Parietal Lobule | 0.038 | 0.027 - 0.049 | 0.000 |
| Right Inferior Temporal Gyrus | 0.059 | 0.046 - 0.074 | 0.000 |
| Right Insular Cortex | 0.035 | 0.025 - 0.046 | 0.000 |
| Right Isthmus of the Cingulate Cortex | 0.019 | 0.011 - 0.027 | 0.000 |
| Right Lateral Occipital Cortex | 0.057 | 0.043 - 0.072 | 0.000 |
| Right Lateral Orbitofrontal Cortex | 0.059 | 0.046 - 0.074 | 0.000 |
| Right Lingual Gyrus | 0.023 | 0.015 - 0.033 | 0.000 |
| Right Medial Orbitofrontal Cortex | 0.044 | 0.034 - 0.056 | 0.000 |
| Right Middle Temporal Gyrus | 0.083 | 0.068 - 0.100 | 0.000 |
| Right Paracentral Lobule | 0.018 | 0.011 - 0.027 | 0.000 |
| Right Parahippocampal Gyrus | 0.032 | 0.023 - 0.041 | 0.000 |
| Right Pars Opercularis | 0.021 | 0.014 - 0.030 | 0.000 |
| Right Pars Orbitalis | 0.038 | 0.028 - 0.050 | 0.000 |
| Right Pars Triangularis | 0.009 | 0.005 - 0.015 | 0.000 |
| Right Pericalcarine Cortex | 0.011 | 0.005 - 0.018 | 0.000 |
| Right Postcentral Gyrus | 0.045 | 0.034 - 0.058 | 0.000 |
| Right Posterior Cingulate Cortex | 0.025 | 0.017 - 0.035 | 0.000 |
| Right Precentral Gyrus | 0.058 | 0.046 - 0.073 | 0.000 |
| Right Precuneus | 0.048 | 0.036 - 0.061 | 0.000 |
| Right Rostral Anterior Cingulate Cortex | 0.026 | 0.018 - 0.036 | 0.000 |
| Right Rostral Middle Frontal Gyrus | 0.033 | 0.024 - 0.045 | 0.000 |
| Right Superior Frontal Gyrus | 0.056 | 0.043 - 0.070 | 0.000 |
| Right Superior Parietal Lobule | 0.039 | 0.029 - 0.051 | 0.000 |
| Right Superior Temporal Gyrus | 0.055 | 0.044 - 0.068 | 0.000 |
| Right Supramarginal Gyrus | 0.031 | 0.021 - 0.041 | 0.000 |
| Right Temporal Pole | 0.014 | 0.008 - 0.021 | 0.000 |
| Right Transverse Temporal Gyrus (Heschl’s Gyrus) | 0.029 | 0.021 - 0.039 | 0.000 |

**Supplementary Table 32: Regions mediating the associations between COI 2.0 health/environment scores and Picture Sequence Memory scores at baseline**

The table summarises the mediation analyses ran across regions that showed both environment-brain and brain-behaviour associations. The ACME (Average Causal Mediation Effect) estimate is reported, which reflects the indirect effect of the predictor on the outcome that is transmitted through the mediator, as well as proportion mediated values, which indicate the estimated fraction of the total effect explained by the mediator. Associated ACME confidence intervals and FDR-corrected p values are also reported. P values are false discovery rate (FDR) corrected to account for multiple comparisons. Analyses controlled for age and sex and included random intercepts for testing site and family ID (to account for sibling pairs).

| **Region** | **ACME estimate** | **ACME CI** | **Proportion mediated** | ***p*_FDR_** |
| --- | --- | --- | --- | --- |
| Left Banks of the Superior Temporal Sulcus | 0.022 | 0.006 - 0.040 | 0.022 | 0.000 |
| Left Caudal Anterior Cingulate Cortex | 0.021 | 0.005 - 0.040 | 0.014 | 0.000 |
| Left Caudal Middle Frontal Gyrus | 0.025 | 0.009 - 0.043 | 0.038 | 0.000 |
| Left Cuneus | 0.036 | 0.018 - 0.057 | 0.021 | 0.000 |
| Left Entorhinal Cortex | 0.015 | 0.002 - 0.029 | 0.032 | 0.000 |
| Left Frontal Pole | 0.058 | 0.033 - 0.084 | 0.018 | 0.000 |
| Left Fusiform Gyrus | 0.027 | 0.011 - 0.046 | 0.057 | 0.000 |
| Left Inferior Parietal Lobule | 0.013 | -0.001 - 0.027 | 0.021 | 0.000 |
| Left Inferior Temporal Gyrus | 0.031 | 0.011 - 0.052 | 0.063 | 0.000 |
| Left Insular Cortex | 0.046 | 0.025 - 0.070 | 0.038 | 0.000 |
| Left Isthmus of the Cingulate Cortex | 0.015 | 0.000 - 0.032 | 0.031 | 0.000 |
| Left Lateral Occipital Cortex | 0.049 | 0.025 - 0.074 | 0.047 | 0.000 |
| Left Lateral Orbitofrontal Cortex | 0.020 | 0.005 - 0.039 | 0.059 | 0.000 |
| Left Lingual Gyrus | 0.028 | 0.014 - 0.046 | 0.027 | 0.000 |
| Left Medial Orbitofrontal Cortex | 0.031 | 0.013 - 0.051 | 0.022 | 0.000 |
| Left Middle Temporal Gyrus | 0.011 | 0.000 - 0.023 | 0.065 | 0.000 |
| Left Paracentral Lobule | 0.040 | 0.016 - 0.066 | 0.025 | 0.000 |
| Left Parahippocampal Gyrus | 0.013 | 0.001 - 0.027 | 0.027 | 0.000 |
| Left Pars Opercularis | 0.047 | 0.024 - 0.073 | 0.023 | 0.000 |
| Left Pars Orbitalis | 0.031 | 0.011 - 0.055 | 0.040 | 0.000 |
| Left Pars Triangularis | 0.019 | 0.007 - 0.035 | 0.010 | 0.000 |
| Left Pericalcarine Cortex | 0.023 | 0.007 - 0.041 | 0.012 | 0.000 |
| Left Postcentral Gyrus | 0.026 | 0.008 - 0.044 | 0.059 | 0.000 |
| Left Posterior Cingulate Cortex | 0.039 | 0.020 - 0.061 | 0.026 | 0.000 |
| Left Precentral Gyrus | 0.018 | 0.005 - 0.033 | 0.066 | 0.000 |
| Left Precuneus | 0.030 | 0.014 - 0.049 | 0.047 | 0.000 |
| Left Rostral Anterior Cingulate Cortex | 0.014 | 0.004 - 0.028 | 0.025 | 0.000 |
| Left Rostral Middle Frontal Gyrus | 0.010 | -0.003 - 0.023 | 0.041 | 0.000 |
| Left Superior Frontal Gyrus | 0.015 | 0.004 - 0.028 | 0.047 | 0.000 |
| Left Superior Parietal Lobule | 0.022 | 0.006 - 0.041 | 0.038 | 0.000 |
| Left Superior Temporal Gyrus | 0.020 | 0.004 - 0.037 | 0.039 | 0.000 |
| Left Supramarginal Gyrus | 0.019 | 0.003 - 0.037 | 0.035 | 0.000 |
| Left Temporal Pole | 0.037 | 0.018 - 0.058 | 0.016 | 0.000 |
| Left Transverse Temporal Gyrus (Heschl’s Gyrus) | 0.023 | 0.007 - 0.041 | 0.029 | 0.000 |
| Right Banks of the Superior Temporal Sulcus | 0.055 | 0.033 - 0.078 | 0.028 | 0.000 |
| Right Caudal Anterior Cingulate Cortex | 0.024 | 0.010 - 0.040 | 0.016 | 0.000 |
| Right Caudal Middle Frontal Gyrus | 0.036 | 0.014 - 0.062 | 0.034 | 0.000 |
| Right Cuneus | 0.046 | 0.023 - 0.069 | 0.024 | 0.000 |
| Right Entorhinal Cortex | 0.015 | 0.000 - 0.030 | 0.024 | 0.000 |
| Right Frontal Pole | 0.050 | 0.024 - 0.079 | 0.026 | 0.000 |
| Right Fusiform Gyrus | 0.013 | 0.001 - 0.027 | 0.054 | 0.000 |
| Right Inferior Parietal Lobule | 0.011 | -0.003 - 0.025 | 0.036 | 0.000 |
| Right Inferior Temporal Gyrus | 0.024 | 0.006 - 0.046 | 0.056 | 0.000 |
| Right Insular Cortex | 0.008 | 0.000 - 0.020 | 0.030 | 0.000 |
| Right Isthmus of the Cingulate Cortex | 0.032 | 0.014 - 0.053 | 0.015 | 0.002 |
| Right Lateral Occipital Cortex | 0.013 | 0.000 - 0.029 | 0.062 | 0.000 |
| Right Lateral Orbitofrontal Cortex | 0.034 | 0.013 - 0.056 | 0.059 | 0.000 |
| Right Lingual Gyrus | 0.034 | 0.012 - 0.056 | 0.021 | 0.000 |
| Right Medial Orbitofrontal Cortex | 0.022 | 0.010 - 0.038 | 0.037 | 0.000 |
| Right Middle Temporal Gyrus | 0.017 | 0.001 - 0.034 | 0.085 | 0.000 |
| Right Paracentral Lobule | 0.035 | 0.017 - 0.058 | 0.018 | 0.000 |
| Right Parahippocampal Gyrus | 0.027 | 0.010 - 0.047 | 0.028 | 0.000 |
| Right Pars Opercularis | 0.028 | 0.010 - 0.047 | 0.026 | 0.000 |
| Right Pars Orbitalis | 0.021 | 0.008 - 0.037 | 0.042 | 0.000 |
| Right Pars Triangularis | 0.015 | 0.000 - 0.033 | 0.010 | 0.000 |
| Right Pericalcarine Cortex | 0.022 | 0.006 - 0.040 | 0.009 | 0.000 |
| Right Postcentral Gyrus | 0.021 | 0.005 - 0.040 | 0.045 | 0.000 |
| Right Posterior Cingulate Cortex | 0.025 | 0.009 - 0.043 | 0.024 | 0.000 |
| Right Precentral Gyrus | 0.036 | 0.018 - 0.057 | 0.054 | 0.000 |
| Right Precuneus | 0.015 | 0.002 - 0.029 | 0.052 | 0.000 |
| Right Rostral Anterior Cingulate Cortex | 0.058 | 0.033 - 0.084 | 0.022 | 0.000 |
| Right Rostral Middle Frontal Gyrus | 0.027 | 0.011 - 0.046 | 0.032 | 0.000 |
| Right Superior Frontal Gyrus | 0.013 | -0.001 - 0.027 | 0.050 | 0.000 |
| Right Superior Parietal Lobule | 0.031 | 0.011 - 0.052 | 0.040 | 0.000 |
| Right Superior Temporal Gyrus | 0.046 | 0.025 - 0.070 | 0.049 | 0.000 |
| Right Supramarginal Gyrus | 0.015 | 0.000 - 0.032 | 0.027 | 0.000 |
| Right Temporal Pole | 0.049 | 0.025 - 0.074 | 0.010 | 0.014 |
| Right Transverse Temporal Gyrus (Heschl’s Gyrus) | 0.020 | 0.005 - 0.039 | 0.034 | 0.000 |

**Supplementary Table 33: Regions mediating the associations between COI 2.0 social/economic scores and Picture Sequence Memory scores at baseline**

The table summarises the mediation analyses ran across regions that showed both environment-brain and brain-behaviour associations. The ACME (Average Causal Mediation Effect) estimate is reported, which reflects the indirect effect of the predictor on the outcome that is transmitted through the mediator, as well as proportion mediated values, which indicate the estimated fraction of the total effect explained by the mediator. Associated ACME confidence intervals and FDR-corrected p values are also reported. P values are false discovery rate (FDR) corrected to account for multiple comparisons. Analyses controlled for age and sex and included random intercepts for testing site and family ID (to account for sibling pairs).

| **Region** | **ACME estimate** | **ACME CI** | **Proportion mediated** | ***p*_FDR_** |
| --- | --- | --- | --- | --- |
| Left Banks of the Superior Temporal Sulcus | 0.022 | 0.000 - 0.047 | 0.019 | <0.001 |
| Left Caudal Anterior Cingulate Cortex | 0.023 | 0.001 - 0.047 | 0.017 | <0.001 |
| Left Caudal Middle Frontal Gyrus | 0.030 | 0.007 - 0.054 | 0.036 | <0.001 |
| Left Cuneus | 0.043 | 0.015 - 0.074 | 0.017 | <0.001 |
| Left Entorhinal Cortex | 0.016 | 0.002 - 0.033 | 0.032 | <0.001 |
| Left Frontal Pole | 0.067 | 0.037 - 0.098 | 0.015 | <0.001 |
| Left Fusiform Gyrus | 0.033 | 0.011 - 0.054 | 0.060 | <0.001 |
| Left Inferior Parietal Lobule | 0.013 | -0.006 - 0.033 | 0.019 | <0.001 |
| Left Inferior Temporal Gyrus | 0.032 | 0.003 - 0.062 | 0.059 | <0.001 |
| Left Insular Cortex | 0.058 | 0.025 - 0.091 | 0.038 | <0.001 |
| Left Isthmus of the Cingulate Cortex | 0.015 | -0.007 - 0.038 | 0.029 | <0.001 |
| Left Lateral Occipital Cortex | 0.056 | 0.026 - 0.087 | 0.045 | <0.001 |
| Left Lateral Orbitofrontal Cortex | 0.024 | 0.003 - 0.047 | 0.063 | <0.001 |
| Left Lingual Gyrus | 0.038 | 0.016 - 0.060 | 0.028 | <0.001 |
| Left Medial Orbitofrontal Cortex | 0.034 | 0.011 - 0.058 | 0.024 | <0.001 |
| Left Middle Temporal Gyrus | 0.012 | -0.003 - 0.028 | 0.061 | <0.001 |
| Left Paracentral Lobule | 0.041 | 0.009 - 0.074 | 0.025 | <0.001 |
| Left Parahippocampal Gyrus | 0.013 | -0.002 - 0.029 | 0.029 | <0.001 |
| Left Pars Opercularis | 0.051 | 0.020 - 0.085 | 0.020 | <0.001 |
| Left Pars Orbitalis | 0.032 | 0.002 - 0.063 | 0.036 | <0.001 |
| Left Pars Triangularis | 0.028 | 0.008 - 0.049 | 0.010 | <0.001 |
| Left Pericalcarine Cortex | 0.025 | 0.000 - 0.051 | 0.011 | <0.001 |
| Left Postcentral Gyrus | 0.027 | 0.004 - 0.052 | 0.054 | <0.001 |
| Left Posterior Cingulate Cortex | 0.049 | 0.021 - 0.077 | 0.023 | <0.001 |
| Left Precentral Gyrus | 0.023 | 0.000 - 0.048 | 0.062 | <0.001 |
| Left Precuneus | 0.040 | 0.015 - 0.066 | 0.043 | <0.001 |
| Left Rostral Anterior Cingulate Cortex | 0.019 | 0.003 - 0.038 | 0.029 | <0.001 |
| Left Rostral Middle Frontal Gyrus | 0.009 | -0.009 - 0.027 | 0.042 | <0.001 |
| Left Superior Frontal Gyrus | 0.017 | 0.003 - 0.035 | 0.043 | <0.001 |
| Left Superior Parietal Lobule | 0.025 | 0.003 - 0.049 | 0.037 | <0.001 |
| Left Superior Temporal Gyrus | 0.021 | -0.003 - 0.043 | 0.047 | <0.001 |
| Left Supramarginal Gyrus | 0.018 | -0.007 - 0.042 | 0.038 | <0.001 |
| Left Temporal Pole | 0.046 | 0.017 - 0.075 | 0.018 | <0.001 |
| Left Transverse Temporal Gyrus (Heschl’s Gyrus) | 0.025 | 0.004 - 0.048 | 0.030 | <0.001 |
| Right Banks of the Superior Temporal Sulcus | 0.070 | 0.037 - 0.102 | 0.027 | <0.001 |
| Right Caudal Anterior Cingulate Cortex | 0.029 | 0.009 - 0.050 | 0.015 | <0.001 |
| Right Caudal Middle Frontal Gyrus | 0.036 | 0.002 - 0.069 | 0.033 | <0.001 |
| Right Cuneus | 0.055 | 0.023 - 0.085 | 0.021 | <0.001 |
| Right Entorhinal Cortex | 0.015 | -0.007 - 0.036 | 0.020 | <0.001 |
| Right Frontal Pole | 0.052 | 0.018 - 0.087 | 0.024 | <0.001 |
| Right Fusiform Gyrus | 0.014 | -0.002 - 0.030 | 0.058 | <0.001 |
| Right Inferior Parietal Lobule | 0.011 | -0.007 - 0.030 | 0.034 | <0.001 |
| Right Inferior Temporal Gyrus | 0.024 | -0.001 - 0.051 | 0.056 | <0.001 |
| Right Insular Cortex | 0.011 | 0.000 - 0.026 | 0.030 | <0.001 |
| Right Isthmus of the Cingulate Cortex | 0.035 | 0.009 - 0.063 | 0.015 | <0.001 |
| Right Lateral Occipital Cortex | 0.015 | -0.004 - 0.033 | 0.058 | <0.001 |
| Right Lateral Orbitofrontal Cortex | 0.038 | 0.007 - 0.067 | 0.058 | <0.001 |
| Right Lingual Gyrus | 0.035 | 0.008 - 0.066 | 0.021 | <0.001 |
| Right Medial Orbitofrontal Cortex | 0.030 | 0.014 - 0.050 | 0.040 | <0.001 |
| Right Middle Temporal Gyrus | 0.017 | -0.005 - 0.042 | 0.077 | <0.001 |
| Right Paracentral Lobule | 0.043 | 0.014 - 0.073 | 0.016 | <0.001 |
| Right Parahippocampal Gyrus | 0.030 | 0.001 - 0.057 | 0.028 | <0.001 |
| Right Pars Opercularis | 0.033 | 0.008 - 0.058 | 0.023 | <0.001 |
| Right Pars Orbitalis | 0.025 | 0.007 - 0.044 | 0.037 | <0.001 |
| Right Pars Triangularis | 0.013 | -0.008 - 0.036 | 0.008 | <0.001 |
| Right Pericalcarine Cortex | 0.022 | 0.000 - 0.047 | 0.010 | <0.001 |
| Right Postcentral Gyrus | 0.023 | 0.001 - 0.047 | 0.042 | <0.001 |
| Right Posterior Cingulate Cortex | 0.030 | 0.007 - 0.054 | 0.025 | <0.001 |
| Right Precentral Gyrus | 0.043 | 0.015 - 0.074 | 0.053 | <0.001 |
| Right Precuneus | 0.016 | 0.002 - 0.033 | 0.047 | <0.001 |
| Right Rostral Anterior Cingulate Cortex | 0.067 | 0.037 - 0.098 | 0.024 | <0.001 |
| Right Rostral Middle Frontal Gyrus | 0.033 | 0.011 - 0.054 | 0.032 | <0.001 |
| Right Superior Frontal Gyrus | 0.013 | -0.006 - 0.033 | 0.052 | <0.001 |
| Right Superior Parietal Lobule | 0.032 | 0.003 - 0.062 | 0.039 | <0.001 |
| Right Superior Temporal Gyrus | 0.058 | 0.025 - 0.091 | 0.051 | <0.001 |
| Right Supramarginal Gyrus | 0.015 | -0.007 - 0.038 | 0.026 | <0.001 |
| Right Temporal Pole | 0.056 | 0.026 - 0.087 | 0.012 | <0.001 |
| Right Transverse Temporal Gyrus (Heschl’s Gyrus) | 0.024 | 0.003 - 0.047 | 0.031 | <0.001 |

**Supplementary Table 34: Regions mediating the associations between COI 2.0 education scores and Picture Sequence Memory scores at baseline**

The table summarises the mediation analyses ran across regions that showed both environment-brain and brain-behaviour associations. The ACME (Average Causal Mediation Effect) estimate is reported, which reflects the indirect effect of the predictor on the outcome that is transmitted through the mediator, as well as proportion mediated values, which indicate the estimated fraction of the total effect explained by the mediator. Associated ACME confidence intervals and FDR-corrected p values are also reported. P values are false discovery rate (FDR) corrected to account for multiple comparisons. Analyses controlled for age and sex and included random intercepts for testing site and family ID (to account for sibling pairs).

| **Region** | **ACME estimate** | **ACME CI** | **Proportion mediated** | ***p*_FDR_** |
| --- | --- | --- | --- | --- |
| Left Caudal Middle Frontal Gyrus | 0.024 | 0.003 - 0.046 | 0.024 | 0.038 |
| Left Cuneus | 0.027 | 0.003 - 0.052 | 0.027 | 0.039 |
| Left Entorhinal Cortex | 0.034 | 0.011 - 0.057 | 0.034 | 0.012 |
| Left Fusiform Gyrus | 0.051 | 0.019 - 0.082 | 0.050 | 0.000 |
| Left Inferior Parietal Lobule | 0.019 | 0.002 - 0.036 | 0.019 | 0.043 |
| Left Inferior Temporal Gyrus | 0.076 | 0.044 - 0.110 | 0.075 | 0.000 |
| Left Insular Cortex | 0.037 | 0.013 - 0.062 | 0.037 | 0.010 |
| Left Isthmus of the Cingulate Cortex | 0.016 | -0.004 - 0.036 | 0.015 | 0.135 |
| Left Lateral Occipital Cortex | 0.039 | 0.010 - 0.071 | 0.038 | 0.020 |
| Left Lateral Orbitofrontal Cortex | 0.066 | 0.032 - 0.099 | 0.065 | 0.000 |
| Left Lingual Gyrus | 0.019 | -0.003 - 0.043 | 0.018 | 0.117 |
| Left Middle Temporal Gyrus | 0.063 | 0.033 - 0.097 | 0.062 | 0.000 |
| Left Paracentral Lobule | 0.024 | 0.007 - 0.043 | 0.023 | 0.015 |
| Left Parahippocampal Gyrus | 0.044 | 0.022 - 0.069 | 0.043 | 0.000 |
| Left Pars Orbitalis | 0.039 | 0.015 - 0.066 | 0.038 | 0.000 |
| Left Pericalcarine Cortex | 0.014 | -0.001 - 0.031 | 0.014 | 0.082 |
| Left Postcentral Gyrus | 0.048 | 0.016 - 0.080 | 0.048 | 0.000 |
| Left Posterior Cingulate Cortex | 0.016 | -0.001 - 0.034 | 0.015 | 0.094 |
| Left Precentral Gyrus | 0.055 | 0.025 - 0.087 | 0.054 | 0.000 |
| Left Precuneus | 0.038 | 0.008 - 0.069 | 0.037 | 0.017 |
| Left Rostral Anterior Cingulate Cortex | 0.028 | 0.010 - 0.047 | 0.027 | 0.006 |
| Left Rostral Middle Frontal Gyrus | 0.031 | 0.004 - 0.060 | 0.030 | 0.038 |
| Left Superior Frontal Gyrus | 0.030 | 0.008 - 0.054 | 0.030 | 0.015 |
| Left Superior Parietal Lobule | 0.055 | 0.026 - 0.084 | 0.054 | 0.000 |
| Left Superior Temporal Gyrus | 0.027 | 0.001 - 0.051 | 0.026 | 0.043 |
| Left Supramarginal Gyrus | 0.043 | 0.020 - 0.069 | 0.042 | 0.000 |
| Left Temporal Pole | 0.021 | 0.005 - 0.041 | 0.021 | 0.025 |
| Right Banks of the Superior Temporal Sulcus | 0.011 | -0.006 - 0.029 | 0.011 | 0.212 |
| Right Caudal Anterior Cingulate Cortex | 0.018 | 0.003 - 0.034 | 0.017 | 0.023 |
| Right Caudal Middle Frontal Gyrus | 0.028 | 0.006 - 0.050 | 0.027 | 0.017 |
| Right Cuneus | 0.025 | 0.001 - 0.051 | 0.025 | 0.058 |
| Right Entorhinal Cortex | 0.021 | 0.001 - 0.045 | 0.021 | 0.052 |
| Right Fusiform Gyrus | 0.054 | 0.024 - 0.083 | 0.053 | 0.000 |
| Right Inferior Parietal Lobule | 0.028 | 0.008 - 0.049 | 0.027 | 0.006 |
| Right Inferior Temporal Gyrus | 0.073 | 0.043 - 0.108 | 0.072 | 0.000 |
| Right Insular Cortex | 0.032 | 0.013 - 0.053 | 0.032 | 0.000 |
| Right Lateral Occipital Cortex | 0.044 | 0.010 - 0.078 | 0.043 | 0.012 |
| Right Lateral Orbitofrontal Cortex | 0.060 | 0.032 - 0.094 | 0.059 | 0.000 |
| Right Medial Orbitofrontal Cortex | 0.019 | -0.005 - 0.046 | 0.019 | 0.135 |
| Right Middle Temporal Gyrus | 0.062 | 0.028 - 0.100 | 0.061 | 0.000 |
| Right Paracentral Lobule | 0.015 | -0.001 - 0.032 | 0.015 | 0.078 |
| Right Parahippocampal Gyrus | 0.014 | -0.007 - 0.037 | 0.014 | 0.222 |
| Right Pars Orbitalis | 0.028 | 0.003 - 0.054 | 0.028 | 0.028 |
| Right Pericalcarine Cortex | 0.013 | -0.002 - 0.027 | 0.012 | 0.103 |
| Right Postcentral Gyrus | 0.041 | 0.012 - 0.069 | 0.040 | 0.015 |
| Right Posterior Cingulate Cortex | 0.016 | 0.001 - 0.033 | 0.015 | 0.054 |
| Right Precentral Gyrus | 0.042 | 0.015 - 0.072 | 0.042 | 0.006 |
| Right Precuneus | 0.041 | 0.013 - 0.072 | 0.040 | 0.010 |
| Right Rostral Anterior Cingulate Cortex | 0.028 | 0.014 - 0.048 | 0.027 | 0.000 |
| Right Rostral Middle Frontal Gyrus | 0.020 | -0.002 - 0.042 | 0.020 | 0.082 |
| Right Superior Frontal Gyrus | 0.046 | 0.019 - 0.075 | 0.045 | 0.000 |
| Right Superior Parietal Lobule | 0.035 | 0.007 - 0.063 | 0.035 | 0.012 |
| Right Superior Temporal Gyrus | 0.039 | 0.012 - 0.064 | 0.038 | 0.012 |
| Right Supramarginal Gyrus | 0.030 | 0.010 - 0.054 | 0.030 | 0.010 |
| Right Transverse Temporal Gyrus (Heschl’s Gyrus) | 0.016 | -0.004 - 0.039 | 0.016 | 0.132 |

**Supplementary Table 35: Regions mediating the associations between micro-environment scores and Picture Sequence Memory scores at baseline**

The table summarises the mediation analyses ran across regions that showed both environment-brain and brain-behaviour associations. The ACME (Average Causal Mediation Effect) estimate is reported, which reflects the indirect effect of the predictor on the outcome that is transmitted through the mediator, as well as proportion mediated values, which indicate the estimated fraction of the total effect explained by the mediator. Associated ACME confidence intervals and FDR-corrected p values are also reported. P values are false discovery rate (FDR) corrected to account for multiple comparisons. Analyses controlled for age and sex and included random intercepts for testing site and family ID (to account for sibling pairs).

| **Region** | **ACME estimate** | **ACME CI** | **Proportion mediated** | ***p*_FDR_** |
| --- | --- | --- | --- | --- |
| Left Caudal Middle Frontal Gyrus | 0.030 | -0.008 - 0.068 | 0.013 | 0.215 |
| Left Cuneus | 0.031 | -0.007 - 0.072 | 0.013 | 0.165 |
| Left Entorhinal Cortex | 0.043 | -0.001 - 0.089 | 0.019 | 0.128 |
| Left Fusiform Gyrus | 0.060 | 0.006 - 0.111 | 0.026 | 0.093 |
| Left Inferior Parietal Lobule | 0.024 | -0.010 - 0.063 | 0.011 | 0.232 |
| Left Inferior Temporal Gyrus | 0.100 | 0.048 - 0.158 | 0.043 | 0.000 |
| Left Insular Cortex | 0.048 | 0.005 - 0.092 | 0.021 | 0.093 |
| Left Isthmus of the Cingulate Cortex | 0.015 | -0.016 - 0.049 | 0.007 | 0.352 |
| Left Lateral Occipital Cortex | 0.042 | -0.001 - 0.093 | 0.018 | 0.128 |
| Left Lateral Orbitofrontal Cortex | 0.083 | 0.027 - 0.140 | 0.036 | 0.000 |
| Left Lingual Gyrus | 0.017 | -0.021 - 0.059 | 0.007 | 0.396 |
| Left Middle Temporal Gyrus | 0.082 | 0.025 - 0.142 | 0.035 | 0.034 |
| Left Paracentral Lobule | 0.032 | -0.010 - 0.076 | 0.014 | 0.219 |
| Left Parahippocampal Gyrus | 0.059 | 0.019 - 0.105 | 0.026 | 0.034 |
| Left Pars Orbitalis | 0.050 | 0.006 - 0.093 | 0.022 | 0.072 |
| Left Pericalcarine Cortex | 0.017 | -0.012 - 0.045 | 0.007 | 0.317 |
| Left Postcentral Gyrus | 0.052 | -0.009 - 0.111 | 0.022 | 0.165 |
| Left Posterior Cingulate Cortex | 0.018 | -0.015 - 0.051 | 0.008 | 0.334 |
| Left Precentral Gyrus | 0.070 | 0.008 - 0.128 | 0.030 | 0.093 |
| Left Precuneus | 0.040 | -0.006 - 0.091 | 0.018 | 0.195 |
| Left Rostral Anterior Cingulate Cortex | 0.042 | 0.000 - 0.084 | 0.018 | 0.128 |
| Left Rostral Middle Frontal Gyrus | 0.031 | -0.017 - 0.076 | 0.014 | 0.218 |
| Left Superior Frontal Gyrus | 0.036 | -0.010 - 0.087 | 0.015 | 0.218 |
| Left Superior Parietal Lobule | 0.072 | 0.027 - 0.122 | 0.031 | 0.000 |
| Left Superior Temporal Gyrus | 0.028 | -0.020 - 0.075 | 0.012 | 0.293 |
| Left Supramarginal Gyrus | 0.059 | 0.017 - 0.104 | 0.025 | 0.034 |
| Left Temporal Pole | 0.029 | -0.001 - 0.062 | 0.013 | 0.128 |
| Right Banks of the Superior Temporal Sulcus | 0.006 | -0.032 - 0.042 | 0.002 | 0.802 |
| Right Caudal Anterior Cingulate Cortex | 0.027 | 0.000 - 0.057 | 0.012 | 0.128 |
| Right Caudal Middle Frontal Gyrus | 0.034 | -0.010 - 0.079 | 0.015 | 0.205 |
| Right Cuneus | 0.028 | -0.004 - 0.064 | 0.012 | 0.165 |
| Right Entorhinal Cortex | 0.022 | -0.020 - 0.066 | 0.009 | 0.334 |
| Right Fusiform Gyrus | 0.065 | 0.013 - 0.118 | 0.028 | 0.051 |
| Right Inferior Parietal Lobule | 0.036 | -0.002 - 0.077 | 0.015 | 0.128 |
| Right Inferior Temporal Gyrus | 0.108 | 0.054 - 0.166 | 0.047 | 0.000 |
| Right Insular Cortex | 0.047 | 0.012 - 0.086 | 0.020 | 0.034 |
| Right Lateral Occipital Cortex | 0.045 | -0.007 - 0.098 | 0.020 | 0.165 |
| Right Lateral Orbitofrontal Cortex | 0.079 | 0.027 - 0.126 | 0.034 | 0.000 |
| Right Medial Orbitofrontal Cortex | 0.013 | -0.032 - 0.058 | 0.006 | 0.645 |
| Right Middle Temporal Gyrus | 0.070 | 0.004 - 0.135 | 0.030 | 0.093 |
| Right Paracentral Lobule | 0.019 | -0.012 - 0.049 | 0.008 | 0.304 |
| Right Parahippocampal Gyrus | 0.009 | -0.028 - 0.046 | 0.004 | 0.690 |
| Right Pars Orbitalis | 0.030 | -0.011 - 0.072 | 0.013 | 0.224 |
| Right Pericalcarine Cortex | 0.017 | -0.005 - 0.045 | 0.007 | 0.218 |
| Right Postcentral Gyrus | 0.046 | -0.007 - 0.102 | 0.020 | 0.171 |
| Right Posterior Cingulate Cortex | 0.019 | -0.014 - 0.048 | 0.008 | 0.334 |
| Right Precentral Gyrus | 0.048 | -0.011 - 0.107 | 0.021 | 0.171 |
| Right Precuneus | 0.046 | -0.002 - 0.101 | 0.020 | 0.128 |
| Right Rostral Anterior Cingulate Cortex | 0.050 | 0.017 - 0.088 | 0.022 | 0.034 |
| Right Rostral Middle Frontal Gyrus | 0.019 | -0.023 - 0.061 | 0.008 | 0.469 |
| Right Superior Frontal Gyrus | 0.061 | 0.010 - 0.113 | 0.027 | 0.072 |
| Right Superior Parietal Lobule | 0.040 | -0.002 - 0.084 | 0.017 | 0.128 |
| Right Superior Temporal Gyrus | 0.047 | 0.002 - 0.094 | 0.020 | 0.123 |
| Right Supramarginal Gyrus | 0.040 | 0.002 - 0.079 | 0.017 | 0.112 |
| Right Transverse Temporal Gyrus (Heschl’s Gyrus) | 0.016 | -0.017 - 0.049 | 0.007 | 0.387 |

**Supplementary Table 36: Regions mediating the associations between COI 2.0 health/environment scores and Oral Reading Recognition scores at baseline**

The table summarises the mediation analyses ran across regions that showed both environment-brain and brain-behaviour associations. The ACME (Average Causal Mediation Effect) estimate is reported, which reflects the indirect effect of the predictor on the outcome that is transmitted through the mediator, as well as proportion mediated values, which indicate the estimated fraction of the total effect explained by the mediator. Associated ACME confidence intervals and FDR-corrected p values are also reported. P values are false discovery rate (FDR) corrected to account for multiple comparisons. Analyses controlled for age and sex and included random intercepts for testing site and family ID (to account for sibling pairs).

| **Region** | **ACME estimate** | **ACME CI** | **Proportion mediated** | ***p*_FDR_** |
| --- | --- | --- | --- | --- |
| Left Banks of the Superior Temporal Sulcus | 0.027 | 0.016 - 0.039 | 0.032 | 0.000 |
| Left Caudal Anterior Cingulate Cortex | 0.017 | 0.008 - 0.029 | 0.021 | 0.000 |
| Left Caudal Middle Frontal Gyrus | 0.042 | 0.028 - 0.058 | 0.050 | 0.000 |
| Left Cuneus | 0.026 | 0.015 - 0.038 | 0.030 | 0.000 |
| Left Entorhinal Cortex | 0.036 | 0.023 - 0.050 | 0.042 | 0.000 |
| Left Frontal Pole | 0.023 | 0.013 - 0.035 | 0.028 | 0.000 |
| Left Fusiform Gyrus | 0.068 | 0.050 - 0.089 | 0.081 | 0.000 |
| Left Inferior Parietal Lobule | 0.030 | 0.018 - 0.046 | 0.036 | 0.000 |
| Left Inferior Temporal Gyrus | 0.073 | 0.055 - 0.095 | 0.086 | 0.000 |
| Left Insular Cortex | 0.044 | 0.029 - 0.062 | 0.053 | 0.000 |
| Left Isthmus of the Cingulate Cortex | 0.031 | 0.018 - 0.043 | 0.037 | 0.000 |
| Left Lateral Occipital Cortex | 0.060 | 0.044 - 0.078 | 0.071 | 0.000 |
| Left Lateral Orbitofrontal Cortex | 0.068 | 0.049 - 0.088 | 0.080 | 0.000 |
| Left Lingual Gyrus | 0.033 | 0.020 - 0.047 | 0.039 | 0.000 |
| Left Medial Orbitofrontal Cortex | 0.029 | 0.016 - 0.044 | 0.034 | 0.000 |
| Left Middle Temporal Gyrus | 0.076 | 0.057 - 0.097 | 0.090 | 0.000 |
| Left Paracentral Lobule | 0.031 | 0.019 - 0.044 | 0.036 | 0.000 |
| Left Parahippocampal Gyrus | 0.033 | 0.020 - 0.047 | 0.039 | 0.000 |
| Left Pars Opercularis | 0.028 | 0.016 - 0.040 | 0.033 | 0.000 |
| Left Pars Orbitalis | 0.043 | 0.029 - 0.058 | 0.051 | 0.000 |
| Left Pars Triangularis | 0.014 | 0.006 - 0.024 | 0.017 | 0.000 |
| Left Pericalcarine Cortex | 0.013 | 0.006 - 0.022 | 0.016 | 0.000 |
| Left Postcentral Gyrus | 0.078 | 0.059 - 0.099 | 0.093 | 0.000 |
| Left Posterior Cingulate Cortex | 0.033 | 0.019 - 0.048 | 0.039 | 0.000 |
| Left Precentral Gyrus | 0.081 | 0.061 - 0.103 | 0.096 | 0.000 |
| Left Precuneus | 0.056 | 0.041 - 0.075 | 0.067 | 0.000 |
| Left Rostral Anterior Cingulate Cortex | 0.029 | 0.016 - 0.043 | 0.035 | 0.000 |
| Left Rostral Middle Frontal Gyrus | 0.049 | 0.033 - 0.066 | 0.058 | 0.000 |
| Left Superior Frontal Gyrus | 0.053 | 0.036 - 0.072 | 0.063 | 0.000 |
| Left Superior Parietal Lobule | 0.045 | 0.031 - 0.060 | 0.054 | 0.000 |
| Left Superior Temporal Gyrus | 0.048 | 0.031 - 0.068 | 0.058 | 0.000 |
| Left Supramarginal Gyrus | 0.039 | 0.024 - 0.054 | 0.046 | 0.000 |
| Left Temporal Pole | 0.019 | 0.009 - 0.031 | 0.023 | 0.000 |
| Left Transverse Temporal Gyrus (Heschl’s Gyrus) | 0.033 | 0.021 - 0.047 | 0.039 | 0.000 |
| Right Banks of the Superior Temporal Sulcus | 0.033 | 0.019 - 0.047 | 0.039 | 0.000 |
| Right Caudal Anterior Cingulate Cortex | 0.018 | 0.009 - 0.029 | 0.021 | 0.000 |
| Right Caudal Middle Frontal Gyrus | 0.036 | 0.022 - 0.051 | 0.043 | 0.000 |
| Right Cuneus | 0.033 | 0.022 - 0.046 | 0.040 | 0.000 |
| Right Entorhinal Cortex | 0.029 | 0.017 - 0.041 | 0.034 | 0.000 |
| Right Frontal Pole | 0.029 | 0.018 - 0.043 | 0.035 | 0.000 |
| Right Fusiform Gyrus | 0.065 | 0.047 - 0.086 | 0.078 | 0.000 |
| Right Inferior Parietal Lobule | 0.046 | 0.031 - 0.062 | 0.054 | 0.000 |
| Right Inferior Temporal Gyrus | 0.061 | 0.045 - 0.080 | 0.073 | 0.000 |
| Right Insular Cortex | 0.038 | 0.024 - 0.054 | 0.045 | 0.000 |
| Right Isthmus of the Cingulate Cortex | 0.017 | 0.008 - 0.029 | 0.021 | 0.000 |
| Right Lateral Occipital Cortex | 0.077 | 0.059 - 0.096 | 0.092 | 0.000 |
| Right Lateral Orbitofrontal Cortex | 0.068 | 0.050 - 0.088 | 0.081 | 0.000 |
| Right Lingual Gyrus | 0.024 | 0.013 - 0.037 | 0.029 | 0.000 |
| Right Medial Orbitofrontal Cortex | 0.042 | 0.028 - 0.058 | 0.050 | 0.000 |
| Right Middle Temporal Gyrus | 0.098 | 0.077 - 0.123 | 0.117 | 0.000 |
| Right Paracentral Lobule | 0.023 | 0.012 - 0.035 | 0.028 | 0.000 |
| Right Parahippocampal Gyrus | 0.034 | 0.021 - 0.048 | 0.041 | 0.000 |
| Right Pars Opercularis | 0.025 | 0.014 - 0.037 | 0.030 | 0.000 |
| Right Pars Orbitalis | 0.047 | 0.033 - 0.063 | 0.056 | 0.000 |
| Right Pars Triangularis | 0.015 | 0.006 - 0.025 | 0.018 | 0.000 |
| Right Pericalcarine Cortex | 0.013 | 0.004 - 0.022 | 0.015 | 0.002 |
| Right Postcentral Gyrus | 0.058 | 0.042 - 0.077 | 0.069 | 0.000 |
| Right Posterior Cingulate Cortex | 0.030 | 0.017 - 0.045 | 0.036 | 0.000 |
| Right Precentral Gyrus | 0.064 | 0.047 - 0.084 | 0.076 | 0.000 |
| Right Precuneus | 0.061 | 0.044 - 0.080 | 0.072 | 0.000 |
| Right Rostral Anterior Cingulate Cortex | 0.027 | 0.014 - 0.040 | 0.032 | 0.000 |
| Right Rostral Middle Frontal Gyrus | 0.042 | 0.028 - 0.058 | 0.051 | 0.000 |
| Right Superior Frontal Gyrus | 0.057 | 0.040 - 0.075 | 0.068 | 0.000 |
| Right Superior Parietal Lobule | 0.048 | 0.033 - 0.065 | 0.057 | 0.000 |
| Right Superior Temporal Gyrus | 0.058 | 0.039 - 0.077 | 0.069 | 0.000 |
| Right Supramarginal Gyrus | 0.033 | 0.019 - 0.047 | 0.039 | 0.000 |
| Right Temporal Pole | 0.010 | 0.002 - 0.019 | 0.012 | 0.010 |
| Right Transverse Temporal Gyrus (Heschl’s Gyrus) | 0.039 | 0.026 - 0.053 | 0.046 | 0.000 |

**Supplementary Table 37: Regions mediating the associations between COI 2.0 social/economic scores and Oral Reading Recognition scores at baseline**

The table summarises the mediation analyses ran across regions that showed both environment-brain and brain-behaviour associations. The ACME (Average Causal Mediation Effect) estimate is reported, which reflects the indirect effect of the predictor on the outcome that is transmitted through the mediator, as well as proportion mediated values, which indicate the estimated fraction of the total effect explained by the mediator. Associated ACME confidence intervals and FDR-corrected p values are also reported. P values are false discovery rate (FDR) corrected to account for multiple comparisons. Analyses controlled for age and sex and included random intercepts for testing site and family ID (to account for sibling pairs).

| **Region** | **ACME estimate** | **ACME CI** | **Proportion mediated** | ***p*_FDR_** |
| --- | --- | --- | --- | --- |
| Left Banks of the Superior Temporal Sulcus | 0.030 | 0.019 - 0.043 | 0.028 | 0.000 |
| Left Caudal Anterior Cingulate Cortex | 0.029 | 0.017 - 0.042 | 0.026 | 0.002 |
| Left Caudal Middle Frontal Gyrus | 0.051 | 0.037 - 0.067 | 0.046 | 0.000 |
| Left Cuneus | 0.030 | 0.018 - 0.044 | 0.027 | 0.000 |
| Left Entorhinal Cortex | 0.048 | 0.033 - 0.064 | 0.044 | 0.000 |
| Left Frontal Pole | 0.027 | 0.017 - 0.040 | 0.025 | 0.000 |
| Left Fusiform Gyrus | 0.096 | 0.075 - 0.119 | 0.087 | 0.000 |
| Left Inferior Parietal Lobule | 0.036 | 0.024 - 0.050 | 0.033 | 0.000 |
| Left Inferior Temporal Gyrus | 0.090 | 0.070 - 0.115 | 0.082 | 0.000 |
| Left Insular Cortex | 0.059 | 0.042 - 0.078 | 0.054 | 0.000 |
| Left Isthmus of the Cingulate Cortex | 0.039 | 0.026 - 0.054 | 0.035 | 0.000 |
| Left Lateral Occipital Cortex | 0.077 | 0.058 - 0.097 | 0.070 | 0.000 |
| Left Lateral Orbitofrontal Cortex | 0.095 | 0.074 - 0.117 | 0.087 | 0.000 |
| Left Lingual Gyrus | 0.045 | 0.033 - 0.062 | 0.041 | 0.000 |
| Left Medial Orbitofrontal Cortex | 0.041 | 0.028 - 0.057 | 0.037 | 0.000 |
| Left Middle Temporal Gyrus | 0.095 | 0.074 - 0.114 | 0.086 | 0.000 |
| Left Paracentral Lobule | 0.040 | 0.027 - 0.056 | 0.037 | 0.000 |
| Left Parahippocampal Gyrus | 0.048 | 0.034 - 0.065 | 0.044 | 0.000 |
| Left Pars Opercularis | 0.031 | 0.020 - 0.044 | 0.028 | 0.000 |
| Left Pars Orbitalis | 0.050 | 0.035 - 0.067 | 0.046 | 0.000 |
| Left Pars Triangularis | 0.018 | 0.010 - 0.030 | 0.017 | 0.000 |
| Left Pericalcarine Cortex | 0.016 | 0.008 - 0.026 | 0.015 | 0.000 |
| Left Postcentral Gyrus | 0.097 | 0.077 - 0.119 | 0.089 | 0.000 |
| Left Posterior Cingulate Cortex | 0.036 | 0.023 - 0.051 | 0.033 | 0.000 |
| Left Precentral Gyrus | 0.101 | 0.080 - 0.124 | 0.092 | 0.000 |
| Left Precuneus | 0.070 | 0.052 - 0.089 | 0.063 | 0.000 |
| Left Rostral Anterior Cingulate Cortex | 0.047 | 0.033 - 0.062 | 0.043 | 0.000 |
| Left Rostral Middle Frontal Gyrus | 0.066 | 0.049 - 0.087 | 0.060 | 0.000 |
| Left Superior Frontal Gyrus | 0.064 | 0.047 - 0.082 | 0.058 | 0.000 |
| Left Superior Parietal Lobule | 0.060 | 0.044 - 0.079 | 0.055 | 0.000 |
| Left Superior Temporal Gyrus | 0.078 | 0.060 - 0.098 | 0.071 | 0.000 |
| Left Supramarginal Gyrus | 0.055 | 0.041 - 0.074 | 0.051 | 0.000 |
| Left Temporal Pole | 0.030 | 0.019 - 0.043 | 0.027 | 0.000 |
| Left Transverse Temporal Gyrus (Heschl’s Gyrus) | 0.046 | 0.031 - 0.062 | 0.042 | 0.000 |
| Right Banks of the Superior Temporal Sulcus | 0.041 | 0.028 - 0.058 | 0.037 | 0.000 |
| Right Caudal Anterior Cingulate Cortex | 0.022 | 0.013 - 0.033 | 0.020 | 0.000 |
| Right Caudal Middle Frontal Gyrus | 0.046 | 0.032 - 0.062 | 0.042 | 0.000 |
| Right Cuneus | 0.041 | 0.027 - 0.058 | 0.037 | 0.000 |
| Right Entorhinal Cortex | 0.034 | 0.020 - 0.049 | 0.031 | 0.000 |
| Right Frontal Pole | 0.036 | 0.023 - 0.049 | 0.033 | 0.000 |
| Right Fusiform Gyrus | 0.094 | 0.074 - 0.116 | 0.086 | 0.000 |
| Right Inferior Parietal Lobule | 0.056 | 0.040 - 0.074 | 0.051 | 0.000 |
| Right Inferior Temporal Gyrus | 0.081 | 0.062 - 0.101 | 0.074 | 0.000 |
| Right Insular Cortex | 0.051 | 0.036 - 0.068 | 0.046 | 0.000 |
| Right Isthmus of the Cingulate Cortex | 0.023 | 0.013 - 0.034 | 0.021 | 0.000 |
| Right Lateral Occipital Cortex | 0.100 | 0.079 - 0.124 | 0.091 | 0.000 |
| Right Lateral Orbitofrontal Cortex | 0.088 | 0.067 - 0.108 | 0.080 | 0.000 |
| Right Lingual Gyrus | 0.032 | 0.020 - 0.045 | 0.029 | 0.000 |
| Right Medial Orbitofrontal Cortex | 0.060 | 0.043 - 0.078 | 0.054 | 0.000 |
| Right Middle Temporal Gyrus | 0.117 | 0.094 - 0.141 | 0.107 | 0.000 |
| Right Paracentral Lobule | 0.027 | 0.017 - 0.039 | 0.025 | 0.000 |
| Right Parahippocampal Gyrus | 0.045 | 0.032 - 0.061 | 0.041 | 0.000 |
| Right Pars Opercularis | 0.029 | 0.017 - 0.042 | 0.026 | 0.000 |
| Right Pars Orbitalis | 0.054 | 0.040 - 0.072 | 0.049 | 0.000 |
| Right Pars Triangularis | 0.015 | 0.007 - 0.025 | 0.014 | 0.000 |
| Right Pericalcarine Cortex | 0.019 | 0.010 - 0.030 | 0.017 | 0.000 |
| Right Postcentral Gyrus | 0.073 | 0.056 - 0.092 | 0.066 | 0.000 |
| Right Posterior Cingulate Cortex | 0.040 | 0.027 - 0.055 | 0.036 | 0.000 |
| Right Precentral Gyrus | 0.086 | 0.066 - 0.106 | 0.078 | 0.000 |
| Right Precuneus | 0.073 | 0.055 - 0.094 | 0.066 | 0.000 |
| Right Rostral Anterior Cingulate Cortex | 0.038 | 0.025 - 0.051 | 0.034 | 0.000 |
| Right Rostral Middle Frontal Gyrus | 0.056 | 0.041 - 0.073 | 0.051 | 0.000 |
| Right Superior Frontal Gyrus | 0.080 | 0.062 - 0.100 | 0.073 | 0.000 |
| Right Superior Parietal Lobule | 0.062 | 0.047 - 0.080 | 0.057 | 0.000 |
| Right Superior Temporal Gyrus | 0.079 | 0.062 - 0.098 | 0.072 | 0.000 |
| Right Supramarginal Gyrus | 0.043 | 0.030 - 0.057 | 0.039 | 0.000 |
| Right Temporal Pole | 0.017 | 0.008 - 0.027 | 0.015 | 0.000 |
| Right Transverse Temporal Gyrus (Heschl’s Gyrus) | 0.046 | 0.032 - 0.062 | 0.042 | 0.000 |

**Supplementary Table 38: Regions mediating the associations between COI 2.0 education scores and Oral Reading Recognition scores at baseline**

The table summarises the mediation analyses ran across regions that showed both environment-brain and brain-behaviour associations. The ACME (Average Causal Mediation Effect) estimate is reported, which reflects the indirect effect of the predictor on the outcome that is transmitted through the mediator, as well as proportion mediated values, which indicate the estimated fraction of the total effect explained by the mediator. Associated ACME confidence intervals and FDR-corrected p values are also reported. P values are false discovery rate (FDR) corrected to account for multiple comparisons. Analyses controlled for age and sex and included random intercepts for testing site and family ID (to account for sibling pairs).

| **Region** | **ACME estimate** | **ACME CI** | **Proportion mediated** | ***p*_FDR_** |
| --- | --- | --- | --- | --- |
| Left Banks of the Superior Temporal Sulcus | 0.032 | 0.020 - 0.046 | 0.030 | 0.000 |
| Left Caudal Anterior Cingulate Cortex | 0.026 | 0.015 - 0.038 | 0.024 | 0.000 |
| Left Caudal Middle Frontal Gyrus | 0.046 | 0.031 - 0.061 | 0.042 | 0.000 |
| Left Cuneus | 0.031 | 0.017 - 0.044 | 0.028 | 0.000 |
| Left Entorhinal Cortex | 0.049 | 0.033 - 0.066 | 0.045 | 0.000 |
| Left Frontal Pole | 0.029 | 0.018 - 0.042 | 0.027 | 0.000 |
| Left Fusiform Gyrus | 0.100 | 0.080 - 0.124 | 0.092 | 0.000 |
| Left Inferior Parietal Lobule | 0.041 | 0.028 - 0.056 | 0.037 | 0.000 |
| Left Inferior Temporal Gyrus | 0.093 | 0.074 - 0.117 | 0.086 | 0.000 |
| Left Insular Cortex | 0.062 | 0.045 - 0.082 | 0.057 | 0.000 |
| Left Isthmus of the Cingulate Cortex | 0.041 | 0.027 - 0.056 | 0.037 | 0.000 |
| Left Lateral Occipital Cortex | 0.078 | 0.061 - 0.098 | 0.072 | 0.000 |
| Left Lateral Orbitofrontal Cortex | 0.096 | 0.076 - 0.118 | 0.089 | 0.000 |
| Left Lingual Gyrus | 0.047 | 0.033 - 0.063 | 0.043 | 0.000 |
| Left Medial Orbitofrontal Cortex | 0.045 | 0.032 - 0.061 | 0.042 | 0.000 |
| Left Middle Temporal Gyrus | 0.096 | 0.075 - 0.120 | 0.089 | 0.000 |
| Left Paracentral Lobule | 0.035 | 0.023 - 0.049 | 0.032 | 0.000 |
| Left Parahippocampal Gyrus | 0.051 | 0.035 - 0.068 | 0.047 | 0.000 |
| Left Pars Opercularis | 0.033 | 0.021 - 0.046 | 0.030 | 0.000 |
| Left Pars Orbitalis | 0.052 | 0.036 - 0.068 | 0.047 | 0.000 |
| Left Pars Triangularis | 0.021 | 0.011 - 0.032 | 0.019 | 0.000 |
| Left Pericalcarine Cortex | 0.018 | 0.008 - 0.029 | 0.016 | 0.000 |
| Left Postcentral Gyrus | 0.097 | 0.077 - 0.123 | 0.090 | 0.000 |
| Left Posterior Cingulate Cortex | 0.039 | 0.026 - 0.054 | 0.036 | 0.000 |
| Left Precentral Gyrus | 0.095 | 0.075 - 0.117 | 0.087 | 0.000 |
| Left Precuneus | 0.069 | 0.049 - 0.088 | 0.063 | 0.000 |
| Left Rostral Anterior Cingulate Cortex | 0.042 | 0.028 - 0.058 | 0.039 | 0.000 |
| Left Rostral Middle Frontal Gyrus | 0.071 | 0.054 - 0.091 | 0.065 | 0.000 |
| Left Superior Frontal Gyrus | 0.062 | 0.045 - 0.077 | 0.057 | 0.000 |
| Left Superior Parietal Lobule | 0.062 | 0.044 - 0.081 | 0.057 | 0.000 |
| Left Superior Temporal Gyrus | 0.079 | 0.060 - 0.101 | 0.073 | 0.000 |
| Left Supramarginal Gyrus | 0.055 | 0.040 - 0.072 | 0.050 | 0.000 |
| Left Temporal Pole | 0.030 | 0.019 - 0.043 | 0.028 | 0.000 |
| Left Transverse Temporal Gyrus (Heschl’s Gyrus) | 0.045 | 0.031 - 0.061 | 0.042 | 0.000 |
| Right Banks of the Superior Temporal Sulcus | 0.041 | 0.027 - 0.055 | 0.037 | 0.000 |
| Right Caudal Anterior Cingulate Cortex | 0.020 | 0.011 - 0.031 | 0.018 | 0.000 |
| Right Caudal Middle Frontal Gyrus | 0.045 | 0.031 - 0.062 | 0.041 | 0.000 |
| Right Cuneus | 0.043 | 0.028 - 0.057 | 0.039 | 0.000 |
| Right Entorhinal Cortex | 0.032 | 0.019 - 0.046 | 0.030 | 0.000 |
| Right Frontal Pole | 0.040 | 0.026 - 0.057 | 0.037 | 0.000 |
| Right Fusiform Gyrus | 0.100 | 0.079 - 0.123 | 0.092 | 0.000 |
| Right Inferior Parietal Lobule | 0.056 | 0.040 - 0.074 | 0.051 | 0.000 |
| Right Inferior Temporal Gyrus | 0.079 | 0.059 - 0.100 | 0.072 | 0.000 |
| Right Insular Cortex | 0.053 | 0.036 - 0.070 | 0.048 | 0.000 |
| Right Isthmus of the Cingulate Cortex | 0.023 | 0.013 - 0.035 | 0.021 | 0.000 |
| Right Lateral Occipital Cortex | 0.099 | 0.077 - 0.125 | 0.091 | 0.000 |
| Right Lateral Orbitofrontal Cortex | 0.088 | 0.068 - 0.109 | 0.081 | 0.000 |
| Right Lingual Gyrus | 0.035 | 0.022 - 0.049 | 0.032 | 0.000 |
| Right Medial Orbitofrontal Cortex | 0.064 | 0.048 - 0.082 | 0.059 | 0.000 |
| Right Middle Temporal Gyrus | 0.123 | 0.100 - 0.147 | 0.113 | 0.000 |
| Right Paracentral Lobule | 0.027 | 0.017 - 0.040 | 0.025 | 0.000 |
| Right Parahippocampal Gyrus | 0.052 | 0.037 - 0.069 | 0.048 | 0.000 |
| Right Pars Opercularis | 0.026 | 0.015 - 0.037 | 0.024 | 0.000 |
| Right Pars Orbitalis | 0.055 | 0.039 - 0.073 | 0.050 | 0.000 |
| Right Pars Triangularis | 0.017 | 0.008 - 0.027 | 0.015 | 0.000 |
| Right Pericalcarine Cortex | 0.020 | 0.011 - 0.031 | 0.018 | 0.000 |
| Right Postcentral Gyrus | 0.075 | 0.057 - 0.096 | 0.069 | 0.000 |
| Right Posterior Cingulate Cortex | 0.035 | 0.023 - 0.050 | 0.033 | 0.000 |
| Right Precentral Gyrus | 0.081 | 0.062 - 0.102 | 0.074 | 0.000 |
| Right Precuneus | 0.072 | 0.053 - 0.093 | 0.066 | 0.000 |
| Right Rostral Anterior Cingulate Cortex | 0.033 | 0.020 - 0.047 | 0.030 | 0.000 |
| Right Rostral Middle Frontal Gyrus | 0.055 | 0.040 - 0.072 | 0.051 | 0.000 |
| Right Superior Frontal Gyrus | 0.076 | 0.058 - 0.096 | 0.070 | 0.000 |
| Right Superior Parietal Lobule | 0.063 | 0.047 - 0.083 | 0.058 | 0.000 |
| Right Superior Temporal Gyrus | 0.084 | 0.066 - 0.105 | 0.078 | 0.000 |
| Right Supramarginal Gyrus | 0.048 | 0.035 - 0.065 | 0.045 | 0.000 |
| Right Temporal Pole | 0.017 | 0.009 - 0.028 | 0.016 | 0.000 |
| Right Transverse Temporal Gyrus (Heschl’s Gyrus) | 0.045 | 0.030 - 0.060 | 0.041 | 0.000 |

**Supplementary Table 39: Regions mediating the associations between micro-environment scores and Oral Reading Recognition scores at baseline**

The table summarises the mediation analyses ran across regions that showed both environment-brain and brain-behaviour associations. The ACME (Average Causal Mediation Effect) estimate is reported, which reflects the indirect effect of the predictor on the outcome that is transmitted through the mediator, as well as proportion mediated values, which indicate the estimated fraction of the total effect explained by the mediator. Associated ACME confidence intervals and FDR-corrected p values are also reported. P values are false discovery rate (FDR) corrected to account for multiple comparisons. Analyses controlled for age and sex and included random intercepts for testing site and family ID (to account for sibling pairs).

| **Region** | **ACME estimate** | **ACME CI** | **Proportion mediated** | ***p*_FDR_** |
| --- | --- | --- | --- | --- |
| Left Banks of the Superior Temporal Sulcus | 0.057 | 0.037 - 0.082 | 0.028 | 0.000 |
| Left Caudal Anterior Cingulate Cortex | 0.046 | 0.028 - 0.068 | 0.023 | 0.000 |
| Left Caudal Middle Frontal Gyrus | 0.077 | 0.055 - 0.100 | 0.038 | 0.000 |
| Left Cuneus | 0.043 | 0.022 - 0.065 | 0.021 | 0.000 |
| Left Entorhinal Cortex | 0.079 | 0.054 - 0.107 | 0.039 | 0.000 |
| Left Frontal Pole | 0.044 | 0.026 - 0.064 | 0.022 | 0.000 |
| Left Fusiform Gyrus | 0.161 | 0.127 - 0.198 | 0.080 | 0.000 |
| Left Inferior Parietal Lobule | 0.076 | 0.055 - 0.101 | 0.038 | 0.000 |
| Left Inferior Temporal Gyrus | 0.143 | 0.110 - 0.178 | 0.071 | 0.000 |
| Left Insular Cortex | 0.097 | 0.073 - 0.125 | 0.048 | 0.000 |
| Left Isthmus of the Cingulate Cortex | 0.060 | 0.040 - 0.083 | 0.030 | 0.000 |
| Left Lateral Occipital Cortex | 0.113 | 0.086 - 0.145 | 0.056 | 0.000 |
| Left Lateral Orbitofrontal Cortex | 0.148 | 0.114 - 0.184 | 0.074 | 0.000 |
| Left Lingual Gyrus | 0.071 | 0.048 - 0.094 | 0.035 | 0.000 |
| Left Medial Orbitofrontal Cortex | 0.081 | 0.058 - 0.107 | 0.040 | 0.000 |
| Left Middle Temporal Gyrus | 0.159 | 0.124 - 0.195 | 0.079 | 0.000 |
| Left Paracentral Lobule | 0.070 | 0.045 - 0.102 | 0.035 | 0.000 |
| Left Parahippocampal Gyrus | 0.079 | 0.056 - 0.104 | 0.039 | 0.000 |
| Left Pars Opercularis | 0.047 | 0.030 - 0.069 | 0.024 | 0.000 |
| Left Pars Orbitalis | 0.082 | 0.057 - 0.109 | 0.041 | 0.000 |
| Left Pars Triangularis | 0.033 | 0.018 - 0.051 | 0.016 | 0.000 |
| Left Pericalcarine Cortex | 0.026 | 0.011 - 0.044 | 0.013 | 0.000 |
| Left Postcentral Gyrus | 0.160 | 0.124 - 0.197 | 0.080 | 0.000 |
| Left Posterior Cingulate Cortex | 0.071 | 0.048 - 0.095 | 0.035 | 0.000 |
| Left Precentral Gyrus | 0.165 | 0.131 - 0.200 | 0.082 | 0.000 |
| Left Precuneus | 0.106 | 0.079 - 0.138 | 0.053 | 0.000 |
| Left Rostral Anterior Cingulate Cortex | 0.084 | 0.061 - 0.111 | 0.042 | 0.000 |
| Left Rostral Middle Frontal Gyrus | 0.113 | 0.083 - 0.142 | 0.056 | 0.000 |
| Left Superior Frontal Gyrus | 0.113 | 0.086 - 0.141 | 0.056 | 0.000 |
| Left Superior Parietal Lobule | 0.094 | 0.065 - 0.123 | 0.047 | 0.000 |
| Left Superior Temporal Gyrus | 0.137 | 0.107 - 0.168 | 0.068 | 0.000 |
| Left Supramarginal Gyrus | 0.088 | 0.062 - 0.116 | 0.044 | 0.000 |
| Left Temporal Pole | 0.050 | 0.031 - 0.072 | 0.025 | 0.000 |
| Left Transverse Temporal Gyrus (Heschl’s Gyrus) | 0.069 | 0.047 - 0.092 | 0.034 | 0.000 |
| Right Banks of the Superior Temporal Sulcus | 0.074 | 0.052 - 0.100 | 0.037 | 0.000 |
| Right Caudal Anterior Cingulate Cortex | 0.035 | 0.018 - 0.055 | 0.017 | 0.000 |
| Right Caudal Middle Frontal Gyrus | 0.081 | 0.055 - 0.108 | 0.040 | 0.000 |
| Right Cuneus | 0.057 | 0.037 - 0.080 | 0.028 | 0.000 |
| Right Entorhinal Cortex | 0.050 | 0.028 - 0.075 | 0.025 | 0.000 |
| Right Frontal Pole | 0.056 | 0.036 - 0.078 | 0.028 | 0.000 |
| Right Fusiform Gyrus | 0.161 | 0.130 - 0.196 | 0.080 | 0.000 |
| Right Inferior Parietal Lobule | 0.100 | 0.075 - 0.128 | 0.050 | 0.000 |
| Right Inferior Temporal Gyrus | 0.132 | 0.100 - 0.166 | 0.066 | 0.000 |
| Right Insular Cortex | 0.093 | 0.068 - 0.121 | 0.046 | 0.000 |
| Right Isthmus of the Cingulate Cortex | 0.045 | 0.028 - 0.066 | 0.022 | 0.000 |
| Right Lateral Occipital Cortex | 0.147 | 0.114 - 0.184 | 0.073 | 0.000 |
| Right Lateral Orbitofrontal Cortex | 0.139 | 0.107 - 0.172 | 0.069 | 0.000 |
| Right Lingual Gyrus | 0.054 | 0.033 - 0.077 | 0.027 | 0.000 |
| Right Medial Orbitofrontal Cortex | 0.104 | 0.075 - 0.132 | 0.052 | 0.000 |
| Right Middle Temporal Gyrus | 0.196 | 0.160 - 0.238 | 0.098 | 0.000 |
| Right Paracentral Lobule | 0.047 | 0.030 - 0.068 | 0.023 | 0.000 |
| Right Parahippocampal Gyrus | 0.080 | 0.057 - 0.104 | 0.040 | 0.000 |
| Right Pars Opercularis | 0.040 | 0.023 - 0.058 | 0.020 | 0.000 |
| Right Pars Orbitalis | 0.084 | 0.059 - 0.111 | 0.042 | 0.000 |
| Right Pars Triangularis | 0.028 | 0.016 - 0.044 | 0.014 | 0.000 |
| Right Pericalcarine Cortex | 0.032 | 0.018 - 0.050 | 0.016 | 0.000 |
| Right Postcentral Gyrus | 0.124 | 0.094 - 0.156 | 0.062 | 0.000 |
| Right Posterior Cingulate Cortex | 0.065 | 0.043 - 0.089 | 0.032 | 0.000 |
| Right Precentral Gyrus | 0.145 | 0.112 - 0.180 | 0.072 | 0.000 |
| Right Precuneus | 0.113 | 0.083 - 0.144 | 0.056 | 0.000 |
| Right Rostral Anterior Cingulate Cortex | 0.066 | 0.046 - 0.093 | 0.033 | 0.000 |
| Right Rostral Middle Frontal Gyrus | 0.094 | 0.068 - 0.122 | 0.047 | 0.000 |
| Right Superior Frontal Gyrus | 0.132 | 0.101 - 0.165 | 0.066 | 0.000 |
| Right Superior Parietal Lobule | 0.096 | 0.068 - 0.124 | 0.048 | 0.000 |
| Right Superior Temporal Gyrus | 0.134 | 0.102 - 0.168 | 0.067 | 0.000 |
| Right Supramarginal Gyrus | 0.080 | 0.057 - 0.107 | 0.040 | 0.000 |
| Right Temporal Pole | 0.029 | 0.016 - 0.046 | 0.014 | 0.000 |
| Right Transverse Temporal Gyrus (Heschl’s Gyrus) | 0.066 | 0.046 - 0.090 | 0.033 | 0.000 |

**Supplementary Table 40: Longitudinal associations between regional centile change scores and COI 2.0 health and environment summary change scores**

Values shown are unstandardised and standardised beta coefficients with associated standard errors (SE) for each region. P values are false discovery rate (FDR) corrected to account for multiple comparisons. All analyses controlled for age and included random intercepts for testing site and family ID (to account for sibling pairs).

| **Region** | **Unstandardised estimate (SE)** | **Standardised estimate (SE)** | **P value (FDR corrected)** |
| --- | --- | --- | --- |
| Left Banks of the Superior Temporal Sulcus | 0.001 (0.001) | 0.020 (0.028) | 0.524 |
| Left Caudal Anterior Cingulate Cortex | 0.000 (0.001) | 0.011 (0.029) | 0.712 |
| Left Caudal Middle Frontal Gyrus | 0.002 (0.001) | 0.069 (0.028) | 0.021 |
| Left Cuneus | 0.002 (0.001) | 0.070 (0.028) | 0.021 |
| Left Entorhinal Cortex | 0.001 (0.002) | 0.018 (0.027) | 0.540 |
| Left Frontal Pole | 0.001 (0.002) | 0.020 (0.028) | 0.520 |
| Left Fusiform Gyrus | 0.002 (0.001) | 0.064 (0.028) | 0.032 |
| Left Inferior Parietal Lobule | 0.004 (0.001) | 0.107 (0.028) | 0.000 |
| Left Inferior Temporal Gyrus | 0.002 (0.001) | 0.049 (0.028) | 0.103 |
| Left Insular Cortex | 0.003 (0.002) | 0.058 (0.028) | 0.055 |
| Left Isthmus of the Cingulate Cortex | 0.001 (0.001) | 0.023 (0.028) | 0.457 |
| Left Lateral Occipital Cortex | 0.002 (0.001) | 0.065 (0.028) | 0.028 |
| Left Lateral Orbitofrontal Cortex | -0.000 (0.001) | -0.001 (0.028) | 0.976 |
| Left Lingual Gyrus | 0.002 (0.001) | 0.081 (0.028) | 0.006 |
| Left Medial Orbitofrontal Cortex | 0.002 (0.002) | 0.034 (0.028) | 0.283 |
| Left Middle Temporal Gyrus | 0.002 (0.001) | 0.057 (0.028) | 0.057 |
| Left Paracentral Lobule | 0.001 (0.001) | 0.028 (0.027) | 0.347 |
| Left Parahippocampal Gyrus | 0.001 (0.001) | 0.015 (0.029) | 0.633 |
| Left Pars Opercularis | 0.001 (0.001) | 0.027 (0.028) | 0.382 |
| Left Pars Orbitalis | 0.002 (0.001) | 0.046 (0.028) | 0.136 |
| Left Pars Triangularis | 0.002 (0.001) | 0.051 (0.029) | 0.103 |
| Left Pericalcarine Cortex | 0.001 (0.001) | 0.015 (0.029) | 0.618 |
| Left Postcentral Gyrus | 0.001 (0.001) | 0.028 (0.028) | 0.369 |
| Left Posterior Cingulate Cortex | 0.001 (0.001) | 0.029 (0.028) | 0.347 |
| Left Precentral Gyrus | 0.002 (0.001) | 0.044 (0.028) | 0.152 |
| Left Precuneus | 0.002 (0.001) | 0.046 (0.028) | 0.126 |
| Left Rostral Anterior Cingulate Cortex | 0.001 (0.001) | 0.024 (0.029) | 0.447 |
| Left Rostral Middle Frontal Gyrus | 0.002 (0.001) | 0.055 (0.028) | 0.065 |
| Left Superior Frontal Gyrus | 0.002 (0.001) | 0.040 (0.027) | 0.179 |
| Left Superior Parietal Lobule | 0.002 (0.001) | 0.040 (0.028) | 0.195 |
| Left Superior Temporal Gyrus | 0.002 (0.001) | 0.063 (0.028) | 0.034 |
| Left Supramarginal Gyrus | 0.003 (0.001) | 0.073 (0.028) | 0.016 |
| Left Temporal Pole | -0.001 (0.002) | -0.008 (0.027) | 0.763 |
| Left Transverse Temporal Gyrus (Heschl’s Gyrus) | 0.002 (0.001) | 0.051 (0.029) | 0.106 |
| Right Banks of the Superior Temporal Sulcus | 0.000 (0.001) | 0.001 (0.028) | 0.976 |
| Right Caudal Anterior Cingulate Cortex | 0.001 (0.001) | 0.041 (0.028) | 0.179 |
| Right Caudal Middle Frontal Gyrus | 0.003 (0.001) | 0.062 (0.028) | 0.039 |
| Right Cuneus | 0.003 (0.001) | 0.081 (0.028) | 0.006 |
| Right Entorhinal Cortex | -0.002 (0.002) | -0.025 (0.028) | 0.406 |
| Right Frontal Pole | -0.001 (0.002) | -0.012 (0.026) | 0.661 |
| Right Fusiform Gyrus | 0.001 (0.001) | 0.025 (0.028) | 0.422 |
| Right Inferior Parietal Lobule | 0.002 (0.001) | 0.048 (0.028) | 0.113 |
| Right Inferior Temporal Gyrus | 0.003 (0.001) | 0.072 (0.028) | 0.015 |
| Right Insular Cortex | 0.005 (0.002) | 0.078 (0.028) | 0.009 |
| Right Isthmus of the Cingulate Cortex | 0.002 (0.001) | 0.056 (0.028) | 0.065 |
| Right Lateral Occipital Cortex | 0.002 (0.001) | 0.067 (0.028) | 0.027 |
| Right Lateral Orbitofrontal Cortex | 0.002 (0.002) | 0.029 (0.029) | 0.366 |
| Right Lingual Gyrus | 0.003 (0.001) | 0.093 (0.027) | 0.001 |
| Right Medial Orbitofrontal Cortex | 0.001 (0.002) | 0.018 (0.028) | 0.541 |
| Right Middle Temporal Gyrus | 0.001 (0.001) | 0.040 (0.028) | 0.191 |
| Right Paracentral Lobule | 0.001 (0.001) | 0.033 (0.028) | 0.299 |
| Right Parahippocampal Gyrus | 0.002 (0.001) | 0.031 (0.028) | 0.306 |
| Right Pars Opercularis | 0.001 (0.001) | 0.025 (0.029) | 0.425 |
| Right Pars Orbitalis | -0.001 (0.001) | -0.020 (0.029) | 0.514 |
| Right Pars Triangularis | -0.000 (0.001) | -0.003 (0.029) | 0.934 |
| Right Pericalcarine Cortex | 0.002 (0.001) | 0.049 (0.029) | 0.114 |
| Right Postcentral Gyrus | 0.001 (0.001) | 0.031 (0.029) | 0.330 |
| Right Posterior Cingulate Cortex | 0.001 (0.001) | 0.027 (0.028) | 0.383 |
| Right Precentral Gyrus | 0.002 (0.001) | 0.049 (0.028) | 0.107 |
| Right Precuneus | 0.004 (0.001) | 0.092 (0.028) | 0.002 |
| Right Rostral Anterior Cingulate Cortex | 0.002 (0.001) | 0.066 (0.028) | 0.030 |
| Right Rostral Middle Frontal Gyrus | 0.002 (0.001) | 0.037 (0.028) | 0.225 |
| Right Superior Frontal Gyrus | 0.003 (0.001) | 0.062 (0.028) | 0.035 |
| Right Superior Parietal Lobule | 0.001 (0.001) | 0.029 (0.028) | 0.354 |
| Right Superior Temporal Gyrus | 0.001 (0.001) | 0.038 (0.028) | 0.214 |
| Right Supramarginal Gyrus | 0.003 (0.001) | 0.067 (0.028) | 0.028 |
| Right Temporal Pole | -0.002 (0.002) | -0.026 (0.026) | 0.369 |
| Right Transverse Temporal Gyrus (Heschl’s Gyrus) | 0.001 (0.001) | 0.018 (0.028) | 0.553 |

**Supplementary Table 41: Longitudinal associations between regional centile change scores and COI 2.0 social and economic summary change scores**

Values shown are unstandardised and standardised beta coefficients with associated standard errors (SE) for each region. P values are false discovery rate (FDR) corrected to account for multiple comparisons. All analyses controlled for age and included random intercepts for testing site and family ID (to account for sibling pairs).

| **Region** | **Unstandardised estimate (SE)** | **Standardised estimate (SE)** | **P value (FDR corrected)** |
| --- | --- | --- | --- |
| Left Banks of the Superior Temporal Sulcus | 0.002 (0.001) | 0.052 (0.026) | 0.060 |
| Left Caudal Anterior Cingulate Cortex | 0.001 (0.001) | 0.028 (0.026) | 0.316 |
| Left Caudal Middle Frontal Gyrus | 0.002 (0.001) | 0.061 (0.026) | 0.027 |
| Left Cuneus | 0.002 (0.001) | 0.059 (0.026) | 0.032 |
| Left Entorhinal Cortex | 0.003 (0.002) | 0.041 (0.026) | 0.134 |
| Left Frontal Pole | 0.000 (0.002) | 0.005 (0.026) | 0.870 |
| Left Fusiform Gyrus | 0.002 (0.001) | 0.062 (0.025) | 0.021 |
| Left Inferior Parietal Lobule | 0.003 (0.001) | 0.099 (0.025) | 0.000 |
| Left Inferior Temporal Gyrus | 0.002 (0.001) | 0.062 (0.025) | 0.024 |
| Left Insular Cortex | 0.002 (0.002) | 0.038 (0.026) | 0.178 |
| Left Isthmus of the Cingulate Cortex | 0.001 (0.001) | 0.037 (0.025) | 0.166 |
| Left Lateral Occipital Cortex | 0.002 (0.001) | 0.053 (0.025) | 0.047 |
| Left Lateral Orbitofrontal Cortex | 0.001 (0.001) | 0.011 (0.026) | 0.689 |
| Left Lingual Gyrus | 0.002 (0.001) | 0.057 (0.025) | 0.032 |
| Left Medial Orbitofrontal Cortex | 0.002 (0.002) | 0.036 (0.026) | 0.206 |
| Left Middle Temporal Gyrus | 0.002 (0.001) | 0.061 (0.025) | 0.024 |
| Left Paracentral Lobule | 0.000 (0.001) | 0.003 (0.026) | 0.902 |
| Left Parahippocampal Gyrus | 0.002 (0.001) | 0.052 (0.027) | 0.070 |
| Left Pars Opercularis | 0.001 (0.001) | 0.040 (0.025) | 0.139 |
| Left Pars Orbitalis | 0.003 (0.001) | 0.060 (0.026) | 0.029 |
| Left Pars Triangularis | 0.002 (0.001) | 0.055 (0.026) | 0.050 |
| Left Pericalcarine Cortex | 0.001 (0.001) | 0.043 (0.026) | 0.127 |
| Left Postcentral Gyrus | -0.000 (0.001) | -0.003 (0.026) | 0.913 |
| Left Posterior Cingulate Cortex | 0.001 (0.001) | 0.044 (0.025) | 0.106 |
| Left Precentral Gyrus | 0.002 (0.001) | 0.049 (0.025) | 0.074 |
| Left Precuneus | 0.001 (0.001) | 0.029 (0.025) | 0.285 |
| Left Rostral Anterior Cingulate Cortex | 0.001 (0.001) | 0.019 (0.026) | 0.502 |
| Left Rostral Middle Frontal Gyrus | 0.002 (0.001) | 0.044 (0.025) | 0.101 |
| Left Superior Frontal Gyrus | 0.002 (0.001) | 0.057 (0.025) | 0.032 |
| Left Superior Parietal Lobule | 0.002 (0.001) | 0.049 (0.026) | 0.078 |
| Left Superior Temporal Gyrus | 0.002 (0.001) | 0.070 (0.025) | 0.010 |
| Left Supramarginal Gyrus | 0.003 (0.001) | 0.076 (0.026) | 0.005 |
| Left Temporal Pole | -0.002 (0.002) | -0.021 (0.026) | 0.443 |
| Left Transverse Temporal Gyrus (Heschl’s Gyrus) | 0.001 (0.001) | 0.041 (0.026) | 0.145 |
| Right Banks of the Superior Temporal Sulcus | -0.000 (0.001) | -0.001 (0.026) | 0.973 |
| Right Caudal Anterior Cingulate Cortex | 0.001 (0.001) | 0.018 (0.026) | 0.502 |
| Right Caudal Middle Frontal Gyrus | 0.001 (0.001) | 0.022 (0.026) | 0.433 |
| Right Cuneus | 0.002 (0.001) | 0.061 (0.026) | 0.026 |
| Right Entorhinal Cortex | 0.001 (0.002) | 0.018 (0.026) | 0.510 |
| Right Frontal Pole | -0.001 (0.002) | -0.019 (0.026) | 0.497 |
| Right Fusiform Gyrus | 0.000 (0.001) | 0.003 (0.025) | 0.902 |
| Right Inferior Parietal Lobule | 0.002 (0.001) | 0.065 (0.025) | 0.016 |
| Right Inferior Temporal Gyrus | 0.002 (0.001) | 0.060 (0.025) | 0.027 |
| Right Insular Cortex | 0.005 (0.002) | 0.076 (0.026) | 0.006 |
| Right Isthmus of the Cingulate Cortex | 0.003 (0.001) | 0.073 (0.026) | 0.007 |
| Right Lateral Occipital Cortex | 0.002 (0.001) | 0.073 (0.026) | 0.007 |
| Right Lateral Orbitofrontal Cortex | 0.002 (0.001) | 0.037 (0.026) | 0.193 |
| Right Lingual Gyrus | 0.002 (0.001) | 0.061 (0.024) | 0.017 |
| Right Medial Orbitofrontal Cortex | 0.001 (0.001) | 0.023 (0.026) | 0.424 |
| Right Middle Temporal Gyrus | 0.001 (0.001) | 0.032 (0.025) | 0.228 |
| Right Paracentral Lobule | 0.001 (0.001) | 0.020 (0.026) | 0.474 |
| Right Parahippocampal Gyrus | 0.002 (0.001) | 0.044 (0.026) | 0.116 |
| Right Pars Opercularis | 0.001 (0.001) | 0.035 (0.026) | 0.208 |
| Right Pars Orbitalis | 0.000 (0.001) | 0.012 (0.026) | 0.675 |
| Right Pars Triangularis | 0.002 (0.001) | 0.047 (0.027) | 0.099 |
| Right Pericalcarine Cortex | 0.002 (0.001) | 0.065 (0.026) | 0.020 |
| Right Postcentral Gyrus | 0.001 (0.001) | 0.026 (0.026) | 0.367 |
| Right Posterior Cingulate Cortex | 0.000 (0.001) | 0.012 (0.025) | 0.658 |
| Right Precentral Gyrus | 0.002 (0.001) | 0.044 (0.026) | 0.105 |
| Right Precuneus | 0.003 (0.001) | 0.082 (0.025) | 0.002 |
| Right Rostral Anterior Cingulate Cortex | 0.002 (0.001) | 0.048 (0.026) | 0.090 |
| Right Rostral Middle Frontal Gyrus | 0.002 (0.001) | 0.049 (0.026) | 0.078 |
| Right Superior Frontal Gyrus | 0.001 (0.001) | 0.029 (0.026) | 0.289 |
| Right Superior Parietal Lobule | 0.002 (0.001) | 0.041 (0.026) | 0.137 |
| Right Superior Temporal Gyrus | 0.001 (0.001) | 0.038 (0.025) | 0.153 |
| Right Supramarginal Gyrus | 0.004 (0.001) | 0.085 (0.026) | 0.002 |
| Right Temporal Pole | -0.001 (0.002) | -0.015 (0.026) | 0.579 |
| Right Transverse Temporal Gyrus (Heschl’s Gyrus) | 0.001 (0.001) | 0.025 (0.027) | 0.388 |

**Supplementary Table 42: Longitudinal associations between regional centile change scores and COI 2.0 education summary change scores**

Values shown are unstandardised and standardised beta coefficients with associated standard errors (SE) for each region. P values are false discovery rate (FDR) corrected to account for multiple comparisons. All analyses controlled for age and included random intercepts for testing site and family ID (to account for sibling pairs).

| **Region** | **Unstandardised estimate (SE)** | **Standardised estimate (SE)** | **P value (FDR corrected)** |
| --- | --- | --- | --- |
| Left Banks of the Superior Temporal Sulcus | 0.001 (0.001) | 0.037 (0.025) | 0.170 |
| Left Caudal Anterior Cingulate Cortex | 0.000 (0.001) | 0.008 (0.025) | 0.756 |
| Left Caudal Middle Frontal Gyrus | 0.002 (0.001) | 0.057 (0.025) | 0.032 |
| Left Cuneus | 0.001 (0.001) | 0.027 (0.025) | 0.323 |
| Left Entorhinal Cortex | 0.002 (0.002) | 0.038 (0.025) | 0.170 |
| Left Frontal Pole | 0.003 (0.002) | 0.042 (0.025) | 0.124 |
| Left Fusiform Gyrus | 0.001 (0.001) | 0.039 (0.024) | 0.131 |
| Left Inferior Parietal Lobule | 0.003 (0.001) | 0.089 (0.024) | 0.001 |
| Left Inferior Temporal Gyrus | 0.002 (0.001) | 0.060 (0.024) | 0.020 |
| Left Insular Cortex | 0.001 (0.001) | 0.011 (0.025) | 0.694 |
| Left Isthmus of the Cingulate Cortex | 0.001 (0.001) | 0.035 (0.024) | 0.181 |
| Left Lateral Occipital Cortex | 0.002 (0.001) | 0.068 (0.024) | 0.007 |
| Left Lateral Orbitofrontal Cortex | -0.000 (0.001) | -0.002 (0.025) | 0.957 |
| Left Lingual Gyrus | 0.002 (0.001) | 0.055 (0.024) | 0.031 |
| Left Medial Orbitofrontal Cortex | 0.000 (0.002) | 0.007 (0.025) | 0.790 |
| Left Middle Temporal Gyrus | 0.002 (0.001) | 0.064 (0.024) | 0.012 |
| Left Paracentral Lobule | 0.001 (0.001) | 0.031 (0.025) | 0.249 |
| Left Parahippocampal Gyrus | 0.002 (0.001) | 0.052 (0.026) | 0.058 |
| Left Pars Opercularis | 0.001 (0.001) | 0.022 (0.024) | 0.405 |
| Left Pars Orbitalis | 0.001 (0.001) | 0.024 (0.025) | 0.380 |
| Left Pars Triangularis | 0.001 (0.001) | 0.037 (0.025) | 0.170 |
| Left Pericalcarine Cortex | 0.002 (0.001) | 0.047 (0.025) | 0.081 |
| Left Postcentral Gyrus | 0.000 (0.001) | 0.003 (0.025) | 0.909 |
| Left Posterior Cingulate Cortex | 0.001 (0.001) | 0.028 (0.024) | 0.294 |
| Left Precentral Gyrus | 0.002 (0.001) | 0.046 (0.024) | 0.078 |
| Left Precuneus | 0.000 (0.001) | 0.013 (0.024) | 0.616 |
| Left Rostral Anterior Cingulate Cortex | -0.000 (0.001) | 0.000 (0.025) | 0.996 |
| Left Rostral Middle Frontal Gyrus | 0.002 (0.001) | 0.061 (0.024) | 0.018 |
| Left Superior Frontal Gyrus | 0.002 (0.001) | 0.048 (0.024) | 0.059 |
| Left Superior Parietal Lobule | 0.001 (0.001) | 0.014 (0.025) | 0.600 |
| Left Superior Temporal Gyrus | 0.002 (0.001) | 0.062 (0.024) | 0.015 |
| Left Supramarginal Gyrus | 0.002 (0.001) | 0.053 (0.025) | 0.046 |
| Left Temporal Pole | 0.002 (0.002) | 0.023 (0.025) | 0.411 |
| Left Transverse Temporal Gyrus (Heschl’s Gyrus) | 0.000 (0.001) | 0.012 (0.025) | 0.660 |
| Right Banks of the Superior Temporal Sulcus | 0.001 (0.001) | 0.013 (0.025) | 0.634 |
| Right Caudal Anterior Cingulate Cortex | 0.001 (0.001) | 0.030 (0.025) | 0.265 |
| Right Caudal Middle Frontal Gyrus | 0.002 (0.001) | 0.044 (0.025) | 0.104 |
| Right Cuneus | 0.002 (0.001) | 0.068 (0.025) | 0.009 |
| Right Entorhinal Cortex | -0.000 (0.002) | -0.001 (0.026) | 0.969 |
| Right Frontal Pole | -0.002 (0.002) | -0.027 (0.025) | 0.323 |
| Right Fusiform Gyrus | -0.001 (0.001) | -0.014 (0.024) | 0.594 |
| Right Inferior Parietal Lobule | 0.003 (0.001) | 0.078 (0.024) | 0.002 |
| Right Inferior Temporal Gyrus | 0.001 (0.001) | 0.026 (0.024) | 0.314 |
| Right Insular Cortex | 0.005 (0.002) | 0.080 (0.025) | 0.003 |
| Right Isthmus of the Cingulate Cortex | 0.002 (0.001) | 0.066 (0.024) | 0.011 |
| Right Lateral Occipital Cortex | 0.002 (0.001) | 0.071 (0.024) | 0.006 |
| Right Lateral Orbitofrontal Cortex | 0.001 (0.001) | 0.016 (0.025) | 0.561 |
| Right Lingual Gyrus | 0.001 (0.001) | 0.045 (0.023) | 0.068 |
| Right Medial Orbitofrontal Cortex | 0.001 (0.001) | 0.014 (0.026) | 0.621 |
| Right Middle Temporal Gyrus | 0.001 (0.001) | 0.031 (0.024) | 0.239 |
| Right Paracentral Lobule | 0.002 (0.001) | 0.051 (0.025) | 0.057 |
| Right Parahippocampal Gyrus | 0.001 (0.001) | 0.021 (0.025) | 0.457 |
| Right Pars Opercularis | 0.001 (0.001) | 0.019 (0.025) | 0.485 |
| Right Pars Orbitalis | 0.000 (0.001) | 0.001 (0.025) | 0.975 |
| Right Pars Triangularis | 0.001 (0.001) | 0.029 (0.026) | 0.291 |
| Right Pericalcarine Cortex | 0.001 (0.001) | 0.037 (0.025) | 0.170 |
| Right Postcentral Gyrus | 0.001 (0.001) | 0.029 (0.025) | 0.288 |
| Right Posterior Cingulate Cortex | 0.000 (0.001) | 0.016 (0.024) | 0.562 |
| Right Precentral Gyrus | 0.002 (0.001) | 0.046 (0.024) | 0.078 |
| Right Precuneus | 0.003 (0.001) | 0.074 (0.024) | 0.003 |
| Right Rostral Anterior Cingulate Cortex | 0.002 (0.001) | 0.047 (0.025) | 0.084 |
| Right Rostral Middle Frontal Gyrus | 0.002 (0.001) | 0.051 (0.025) | 0.062 |
| Right Superior Frontal Gyrus | 0.002 (0.001) | 0.050 (0.025) | 0.059 |
| Right Superior Parietal Lobule | 0.002 (0.001) | 0.033 (0.025) | 0.227 |
| Right Superior Temporal Gyrus | 0.001 (0.001) | 0.030 (0.024) | 0.249 |
| Right Supramarginal Gyrus | 0.003 (0.001) | 0.063 (0.025) | 0.019 |
| Right Temporal Pole | -0.001 (0.002) | -0.011 (0.025) | 0.697 |
| Right Transverse Temporal Gyrus (Heschl’s Gyrus) | 0.001 (0.001) | 0.028 (0.025) | 0.315 |

**Supplementary Table 43: Longitudinal associations between regional centile change scores and micro-environment latent factor change scores**

Values shown are unstandardised and standardised beta coefficients with associated standard errors (SE) for each region. P values are false discovery rate (FDR) corrected to account for multiple comparisons. All analyses controlled for age and included random intercepts for testing site and family ID (to account for sibling pairs).

| **Region** | **Unstandardised estimate (SE)** | **Standardised estimate (SE)** | **P value (FDR corrected)** |
| --- | --- | --- | --- |
| Left Banks of the Superior Temporal Sulcus | 0.002 (0.002) | 0.038 (0.025) | 0.150 |
| Left Caudal Anterior Cingulate Cortex | 0.001 (0.001) | 0.013 (0.025) | 0.632 |
| Left Caudal Middle Frontal Gyrus | 0.002 (0.001) | 0.029 (0.025) | 0.278 |
| Left Cuneus | 0.005 (0.001) | 0.084 (0.025) | 0.001 |
| Left Entorhinal Cortex | 0.001 (0.002) | 0.012 (0.025) | 0.653 |
| Left Frontal Pole | 0.002 (0.003) | 0.017 (0.026) | 0.532 |
| Left Fusiform Gyrus | 0.001 (0.001) | 0.009 (0.024) | 0.730 |
| Left Inferior Parietal Lobule | 0.004 (0.001) | 0.071 (0.024) | 0.006 |
| Left Inferior Temporal Gyrus | 0.002 (0.001) | 0.031 (0.025) | 0.227 |
| Left Insular Cortex | 0.001 (0.002) | 0.012 (0.026) | 0.648 |
| Left Isthmus of the Cingulate Cortex | 0.003 (0.001) | 0.052 (0.024) | 0.044 |
| Left Lateral Occipital Cortex | 0.003 (0.001) | 0.065 (0.024) | 0.010 |
| Left Lateral Orbitofrontal Cortex | 0.002 (0.002) | 0.034 (0.025) | 0.195 |
| Left Lingual Gyrus | 0.004 (0.001) | 0.086 (0.024) | 0.001 |
| Left Medial Orbitofrontal Cortex | 0.002 (0.002) | 0.022 (0.026) | 0.414 |
| Left Middle Temporal Gyrus | 0.004 (0.001) | 0.065 (0.024) | 0.011 |
| Left Paracentral Lobule | 0.003 (0.002) | 0.050 (0.025) | 0.058 |
| Left Parahippocampal Gyrus | 0.003 (0.002) | 0.046 (0.026) | 0.093 |
| Left Pars Opercularis | 0.002 (0.001) | 0.044 (0.024) | 0.094 |
| Left Pars Orbitalis | 0.000 (0.002) | 0.001 (0.025) | 0.968 |
| Left Pars Triangularis | 0.000 (0.001) | 0.001 (0.025) | 0.960 |
| Left Pericalcarine Cortex | 0.005 (0.001) | 0.089 (0.025) | 0.001 |
| Left Postcentral Gyrus | 0.001 (0.001) | 0.027 (0.025) | 0.304 |
| Left Posterior Cingulate Cortex | 0.001 (0.001) | 0.019 (0.024) | 0.470 |
| Left Precentral Gyrus | 0.002 (0.001) | 0.037 (0.024) | 0.149 |
| Left Precuneus | 0.001 (0.001) | 0.025 (0.024) | 0.330 |
| Left Rostral Anterior Cingulate Cortex | -0.000 (0.002) | -0.005 (0.025) | 0.853 |
| Left Rostral Middle Frontal Gyrus | 0.003 (0.002) | 0.044 (0.024) | 0.089 |
| Left Superior Frontal Gyrus | 0.002 (0.001) | 0.043 (0.024) | 0.094 |
| Left Superior Parietal Lobule | 0.003 (0.002) | 0.051 (0.025) | 0.052 |
| Left Superior Temporal Gyrus | 0.004 (0.001) | 0.073 (0.024) | 0.004 |
| Left Supramarginal Gyrus | 0.003 (0.001) | 0.057 (0.025) | 0.029 |
| Left Temporal Pole | -0.001 (0.003) | -0.005 (0.025) | 0.865 |
| Left Transverse Temporal Gyrus (Heschl’s Gyrus) | 0.002 (0.001) | 0.039 (0.025) | 0.149 |
| Right Banks of the Superior Temporal Sulcus | 0.003 (0.002) | 0.051 (0.025) | 0.055 |
| Right Caudal Anterior Cingulate Cortex | 0.001 (0.001) | 0.016 (0.025) | 0.538 |
| Right Caudal Middle Frontal Gyrus | 0.004 (0.002) | 0.061 (0.025) | 0.021 |
| Right Cuneus | 0.005 (0.001) | 0.092 (0.025) | 0.000 |
| Right Entorhinal Cortex | 0.004 (0.003) | 0.043 (0.026) | 0.117 |
| Right Frontal Pole | -0.003 (0.003) | -0.026 (0.025) | 0.330 |
| Right Fusiform Gyrus | 0.001 (0.001) | 0.016 (0.024) | 0.545 |
| Right Inferior Parietal Lobule | 0.002 (0.001) | 0.043 (0.024) | 0.096 |
| Right Inferior Temporal Gyrus | 0.004 (0.001) | 0.082 (0.024) | 0.001 |
| Right Insular Cortex | 0.005 (0.002) | 0.054 (0.025) | 0.047 |
| Right Isthmus of the Cingulate Cortex | 0.005 (0.001) | 0.080 (0.025) | 0.002 |
| Right Lateral Occipital Cortex | 0.004 (0.001) | 0.073 (0.025) | 0.005 |
| Right Lateral Orbitofrontal Cortex | 0.003 (0.002) | 0.036 (0.025) | 0.175 |
| Right Lingual Gyrus | 0.003 (0.001) | 0.054 (0.023) | 0.028 |
| Right Medial Orbitofrontal Cortex | 0.003 (0.002) | 0.041 (0.026) | 0.136 |
| Right Middle Temporal Gyrus | 0.002 (0.001) | 0.041 (0.024) | 0.109 |
| Right Paracentral Lobule | 0.003 (0.002) | 0.052 (0.025) | 0.051 |
| Right Parahippocampal Gyrus | 0.003 (0.002) | 0.039 (0.026) | 0.152 |
| Right Pars Opercularis | 0.003 (0.002) | 0.043 (0.025) | 0.110 |
| Right Pars Orbitalis | 0.001 (0.002) | 0.023 (0.025) | 0.392 |
| Right Pars Triangularis | 0.003 (0.002) | 0.045 (0.026) | 0.105 |
| Right Pericalcarine Cortex | 0.005 (0.001) | 0.087 (0.025) | 0.001 |
| Right Postcentral Gyrus | 0.001 (0.001) | 0.014 (0.025) | 0.612 |
| Right Posterior Cingulate Cortex | 0.002 (0.001) | 0.038 (0.024) | 0.144 |
| Right Precentral Gyrus | 0.004 (0.002) | 0.062 (0.025) | 0.017 |
| Right Precuneus | 0.004 (0.001) | 0.066 (0.024) | 0.010 |
| Right Rostral Anterior Cingulate Cortex | 0.001 (0.001) | 0.022 (0.026) | 0.414 |
| Right Rostral Middle Frontal Gyrus | 0.002 (0.002) | 0.028 (0.025) | 0.295 |
| Right Superior Frontal Gyrus | 0.004 (0.002) | 0.059 (0.025) | 0.025 |
| Right Superior Parietal Lobule | 0.003 (0.002) | 0.037 (0.025) | 0.169 |
| Right Superior Temporal Gyrus | 0.003 (0.001) | 0.054 (0.024) | 0.037 |
| Right Supramarginal Gyrus | 0.003 (0.002) | 0.042 (0.025) | 0.118 |
| Right Temporal Pole | 0.001 (0.003) | 0.008 (0.025) | 0.781 |
| Right Transverse Temporal Gyrus (Heschl’s Gyrus) | 0.002 (0.002) | 0.033 (0.026) | 0.226 |

**Supplementary Table 44: Longitudinal associations between regional centile change scores and CBCL internalising change scores**

Values shown are unstandardised and standardised beta coefficients with associated standard errors (SE) for each region. P values are false discovery rate (FDR) corrected to account for multiple comparisons. All analyses controlled for age and baseline internalising scores, included random intercepts for testing site and family ID (to account for sibling pairs).

| **Region** | **Unstandardised estimate (SE)** | **Standardised estimate** | **P value (FDR corrected)** |
| --- | --- | --- | --- |
| Left Banks of the Superior Temporal Sulcus | -0.169 (0.591) | -0.007 | 0.808 |
| Left Caudal Anterior Cingulate Cortex | 0.341 (0.802) | 0.010 | 0.716 |
| Left Caudal Middle Frontal Gyrus | 0.519 (0.671) | 0.018 | 0.496 |
| Left Cuneus | -1.853 (0.667) | -0.064 | 0.007 |
| Left Entorhinal Cortex | -0.213 (0.363) | -0.013 | 0.606 |
| Left Frontal Pole | -0.382 (0.335) | -0.026 | 0.300 |
| Left Fusiform Gyrus | -1.397 (0.649) | -0.051 | 0.040 |
| Left Inferior Parietal Lobule | -0.453 (0.745) | -0.015 | 0.593 |
| Left Inferior Temporal Gyrus | -0.323 (0.687) | -0.011 | 0.686 |
| Left Insular Cortex | -0.993 (0.394) | -0.057 | 0.015 |
| Left Isthmus of the Cingulate Cortex | 0.217 (0.672) | 0.008 | 0.781 |
| Left Lateral Occipital Cortex | -1.251 (0.711) | -0.042 | 0.098 |
| Left Lateral Orbitofrontal Cortex | -0.330 (0.508) | -0.015 | 0.569 |
| Left Lingual Gyrus | -2.507 (0.831) | -0.072 | 0.003 |
| Left Medial Orbitofrontal Cortex | -0.262 (0.370) | -0.016 | 0.533 |
| Left Middle Temporal Gyrus | -1.004 (0.653) | -0.037 | 0.151 |
| Left Paracentral Lobule | 0.113 (0.521) | 0.005 | 0.838 |
| Left Parahippocampal Gyrus | 0.118 (0.486) | 0.005 | 0.830 |
| Left Pars Opercularis | -0.281 (0.703) | -0.009 | 0.733 |
| Left Pars Orbitalis | -0.875 (0.540) | -0.037 | 0.129 |
| Left Pars Triangularis | -1.765 (0.729) | -0.056 | 0.020 |
| Left Pericalcarine Cortex | -1.767 (0.682) | -0.059 | 0.013 |
| Left Postcentral Gyrus | -0.153 (0.680) | -0.005 | 0.834 |
| Left Posterior Cingulate Cortex | -0.292 (0.765) | -0.009 | 0.743 |
| Left Precentral Gyrus | 0.160 (0.631) | 0.006 | 0.824 |
| Left Precuneus | -0.751 (0.658) | -0.028 | 0.300 |
| Left Rostral Anterior Cingulate Cortex | 0.319 (0.545) | 0.013 | 0.606 |
| Left Rostral Middle Frontal Gyrus | 0.069 (0.581) | 0.003 | 0.910 |
| Left Superior Frontal Gyrus | 0.953 (0.649) | 0.035 | 0.173 |
| Left Superior Parietal Lobule | -0.582 (0.552) | -0.024 | 0.338 |
| Left Superior Temporal Gyrus | -0.906 (0.689) | -0.031 | 0.226 |
| Left Supramarginal Gyrus | -1.265 (0.650) | -0.045 | 0.066 |
| Left Temporal Pole | -0.309 (0.314) | -0.022 | 0.373 |
| Left Transverse Temporal Gyrus (Heschl’s Gyrus) | 0.417 (0.662) | 0.015 | 0.579 |
| Right Banks of the Superior Temporal Sulcus | -0.150 (0.591) | -0.006 | 0.824 |
| Right Caudal Anterior Cingulate Cortex | -0.555 (0.737) | -0.017 | 0.507 |
| Right Caudal Middle Frontal Gyrus | 0.397 (0.572) | 0.016 | 0.539 |
| Right Cuneus | -1.998 (0.668) | -0.069 | 0.004 |
| Right Entorhinal Cortex | -0.298 (0.348) | -0.019 | 0.444 |
| Right Frontal Pole | -0.117 (0.328) | -0.008 | 0.759 |
| Right Fusiform Gyrus | -1.116 (0.670) | -0.040 | 0.119 |
| Right Inferior Parietal Lobule | -0.711 (0.712) | -0.024 | 0.366 |
| Right Inferior Temporal Gyrus | 0.180 (0.688) | 0.006 | 0.824 |
| Right Insular Cortex | -0.734 (0.377) | -0.044 | 0.066 |
| Right Isthmus of the Cingulate Cortex | -1.524 (0.612) | -0.057 | 0.016 |
| Right Lateral Occipital Cortex | -1.393 (0.724) | -0.046 | 0.068 |
| Right Lateral Orbitofrontal Cortex | -0.023 (0.413) | -0.001 | 0.955 |
| Right Lingual Gyrus | -2.271 (0.811) | -0.068 | 0.007 |
| Right Medial Orbitofrontal Cortex | -0.050 (0.426) | -0.003 | 0.910 |
| Right Middle Temporal Gyrus | -0.664 (0.718) | -0.022 | 0.404 |
| Right Paracentral Lobule | 0.615 (0.549) | 0.026 | 0.308 |
| Right Parahippocampal Gyrus | -1.175 (0.432) | -0.061 | 0.009 |
| Right Pars Opercularis | 0.279 (0.609) | 0.011 | 0.692 |
| Right Pars Orbitalis | -0.792 (0.563) | -0.032 | 0.192 |
| Right Pars Triangularis | -0.626 (0.594) | -0.024 | 0.338 |
| Right Pericalcarine Cortex | -1.022 (0.649) | -0.036 | 0.142 |
| Right Postcentral Gyrus | -0.471 (0.650) | -0.017 | 0.525 |
| Right Posterior Cingulate Cortex | 1.430 (0.805) | 0.042 | 0.095 |
| Right Precentral Gyrus | -0.285 (0.533) | -0.012 | 0.640 |
| Right Precuneus | -0.650 (0.615) | -0.025 | 0.338 |
| Right Rostral Anterior Cingulate Cortex | -0.772 (0.632) | -0.028 | 0.265 |
| Right Rostral Middle Frontal Gyrus | 0.397 (0.555) | 0.017 | 0.530 |
| Right Superior Frontal Gyrus | 0.647 (0.563) | 0.027 | 0.297 |
| Right Superior Parietal Lobule | -0.174 (0.490) | -0.008 | 0.759 |
| Right Superior Temporal Gyrus | -0.167 (0.706) | -0.006 | 0.831 |
| Right Supramarginal Gyrus | -0.537 (0.556) | -0.022 | 0.382 |
| Right Temporal Pole | -0.474 (0.324) | -0.033 | 0.174 |
| Right Transverse Temporal Gyrus (Heschl’s Gyrus) | -0.136 (0.591) | -0.005 | 0.834 |

**Supplementary Table 45: Longitudinal associations between regional centile change scores and CBCL externalising change scores**

Values shown are unstandardised and standardised beta coefficients with associated standard errors (SE) for each region. P values are false discovery rate (FDR) corrected to account for multiple comparisons. All analyses controlled for age and baseline externalising scores, included random intercepts for testing site and family ID (to account for sibling pairs).

| **Region** | **Unstandardised estimate (SE)** | **Standardised estimate** | **P value (FDR corrected)** |
| --- | --- | --- | --- |
| Left Banks of the Superior Temporal Sulcus | -0.661 (0.454) | -0.032 | 0.243 |
| Left Caudal Anterior Cingulate Cortex | -0.394 (0.614) | -0.014 | 0.632 |
| Left Caudal Middle Frontal Gyrus | -0.643 (0.515) | -0.028 | 0.337 |
| Left Cuneus | -1.039 (0.512) | -0.045 | 0.078 |
| Left Entorhinal Cortex | -0.438 (0.280) | -0.034 | 0.201 |
| Left Frontal Pole | -0.107 (0.258) | -0.009 | 0.707 |
| Left Fusiform Gyrus | -0.648 (0.496) | -0.030 | 0.308 |
| Left Inferior Parietal Lobule | -0.795 (0.571) | -0.032 | 0.270 |
| Left Inferior Temporal Gyrus | -0.897 (0.527) | -0.039 | 0.156 |
| Left Insular Cortex | -0.743 (0.303) | -0.054 | 0.027 |
| Left Isthmus of the Cingulate Cortex | -0.066 (0.516) | -0.003 | 0.912 |
| Left Lateral Occipital Cortex | -0.869 (0.542) | -0.037 | 0.188 |
| Left Lateral Orbitofrontal Cortex | -0.970 (0.390) | -0.055 | 0.025 |
| Left Lingual Gyrus | -1.140 (0.632) | -0.041 | 0.126 |
| Left Medial Orbitofrontal Cortex | -0.324 (0.285) | -0.025 | 0.395 |
| Left Middle Temporal Gyrus | -1.252 (0.497) | -0.058 | 0.023 |
| Left Paracentral Lobule | -0.303 (0.402) | -0.017 | 0.583 |
| Left Parahippocampal Gyrus | 0.061 (0.373) | 0.004 | 0.890 |
| Left Pars Opercularis | -0.518 (0.538) | -0.021 | 0.482 |
| Left Pars Orbitalis | -0.862 (0.413) | -0.046 | 0.069 |
| Left Pars Triangularis | -1.438 (0.558) | -0.057 | 0.019 |
| Left Pericalcarine Cortex | -1.664 (0.521) | -0.070 | 0.003 |
| Left Postcentral Gyrus | -0.573 (0.521) | -0.025 | 0.417 |
| Left Posterior Cingulate Cortex | -0.669 (0.585) | -0.026 | 0.394 |
| Left Precentral Gyrus | -0.715 (0.482) | -0.034 | 0.233 |
| Left Precuneus | -0.546 (0.502) | -0.025 | 0.419 |
| Left Rostral Anterior Cingulate Cortex | 0.054 (0.418) | 0.003 | 0.912 |
| Left Rostral Middle Frontal Gyrus | -0.847 (0.446) | -0.043 | 0.103 |
| Left Superior Frontal Gyrus | -0.251 (0.497) | -0.012 | 0.673 |
| Left Superior Parietal Lobule | -0.881 (0.422) | -0.047 | 0.069 |
| Left Superior Temporal Gyrus | -0.574 (0.528) | -0.025 | 0.419 |
| Left Supramarginal Gyrus | -0.219 (0.497) | -0.010 | 0.692 |
| Left Temporal Pole | -0.328 (0.242) | -0.029 | 0.287 |
| Left Transverse Temporal Gyrus (Heschl’s Gyrus) | -0.008 (0.505) | -0.000 | 0.991 |
| Right Banks of the Superior Temporal Sulcus | -0.235 (0.455) | -0.011 | 0.667 |
| Right Caudal Anterior Cingulate Cortex | -1.122 (0.567) | -0.044 | 0.087 |
| Right Caudal Middle Frontal Gyrus | -0.199 (0.439) | -0.010 | 0.690 |
| Right Cuneus | -1.274 (0.514) | -0.055 | 0.025 |
| Right Entorhinal Cortex | -0.200 (0.268) | -0.016 | 0.586 |
| Right Frontal Pole | -0.030 (0.253) | -0.003 | 0.917 |
| Right Fusiform Gyrus | -0.864 (0.512) | -0.039 | 0.159 |
| Right Inferior Parietal Lobule | -1.312 (0.547) | -0.056 | 0.031 |
| Right Inferior Temporal Gyrus | -0.319 (0.527) | -0.014 | 0.645 |
| Right Insular Cortex | -0.312 (0.291) | -0.024 | 0.426 |
| Right Isthmus of the Cingulate Cortex | -0.748 (0.470) | -0.035 | 0.191 |
| Right Lateral Occipital Cortex | -1.608 (0.553) | -0.066 | 0.007 |
| Right Lateral Orbitofrontal Cortex | -0.224 (0.318) | -0.015 | 0.609 |
| Right Lingual Gyrus | -1.914 (0.614) | -0.072 | 0.004 |
| Right Medial Orbitofrontal Cortex | -0.552 (0.328) | -0.037 | 0.159 |
| Right Middle Temporal Gyrus | -1.000 (0.549) | -0.042 | 0.122 |
| Right Paracentral Lobule | 0.007 (0.422) | 0.000 | 0.991 |
| Right Parahippocampal Gyrus | -0.451 (0.333) | -0.030 | 0.287 |
| Right Pars Opercularis | -0.425 (0.467) | -0.020 | 0.502 |
| Right Pars Orbitalis | -0.169 (0.433) | -0.009 | 0.723 |
| Right Pars Triangularis | -0.498 (0.456) | -0.024 | 0.419 |
| Right Pericalcarine Cortex | -0.958 (0.497) | -0.042 | 0.097 |
| Right Postcentral Gyrus | -0.449 (0.498) | -0.020 | 0.502 |
| Right Posterior Cingulate Cortex | -0.101 (0.616) | -0.004 | 0.890 |
| Right Precentral Gyrus | -0.951 (0.407) | -0.052 | 0.037 |
| Right Precuneus | -0.272 (0.470) | -0.013 | 0.660 |
| Right Rostral Anterior Cingulate Cortex | 0.001 (0.486) | 0.000 | 0.998 |
| Right Rostral Middle Frontal Gyrus | -0.160 (0.427) | -0.008 | 0.732 |
| Right Superior Frontal Gyrus | -0.230 (0.433) | -0.012 | 0.661 |
| Right Superior Parietal Lobule | -0.567 (0.377) | -0.034 | 0.224 |
| Right Superior Temporal Gyrus | -0.197 (0.541) | -0.008 | 0.738 |
| Right Supramarginal Gyrus | -0.497 (0.428) | -0.026 | 0.384 |
| Right Temporal Pole | -0.333 (0.250) | -0.029 | 0.295 |
| Right Transverse Temporal Gyrus (Heschl’s Gyrus) | -0.259 (0.454) | -0.013 | 0.660 |

**Supplementary Table 46: Longitudinal associations between regional centile change scores and Flanker Inhibitory Control and Attention Task change scores**

Values shown are unstandardised and standardised beta coefficients with associated standard errors (SE) for each region. P values are false discovery rate (FDR) corrected to account for multiple comparisons. All analyses controlled for age and baseline task scores, included random intercepts for testing site and family ID (to account for sibling pairs).

| **Region** | **Unstandardised estimate (SE)** | **Standardised estimate** | **P value (FDR corrected)** |
| --- | --- | --- | --- |
| Left Banks of the Superior Temporal Sulcus | -0.927 (0.810) | -0.021 | 0.304 |
| Left Caudal Anterior Cingulate Cortex | -1.621 (1.099) | -0.027 | 0.179 |
| Left Caudal Middle Frontal Gyrus | 0.388 (0.918) | 0.008 | 0.731 |
| Left Cuneus | 1.275 (0.916) | 0.026 | 0.205 |
| Left Entorhinal Cortex | -0.067 (0.499) | -0.002 | 0.910 |
| Left Frontal Pole | 0.628 (0.459) | 0.025 | 0.213 |
| Left Fusiform Gyrus | -1.464 (0.885) | -0.032 | 0.127 |
| Left Inferior Parietal Lobule | -0.154 (1.015) | -0.003 | 0.906 |
| Left Inferior Temporal Gyrus | -0.328 (0.937) | -0.007 | 0.778 |
| Left Insular Cortex | 0.837 (0.539) | 0.029 | 0.154 |
| Left Isthmus of the Cingulate Cortex | -0.387 (0.920) | -0.008 | 0.731 |
| Left Lateral Occipital Cortex | 1.172 (0.967) | 0.024 | 0.274 |
| Left Lateral Orbitofrontal Cortex | -1.453 (0.694) | -0.039 | 0.048 |
| Left Lingual Gyrus | 2.096 (1.130) | 0.036 | 0.084 |
| Left Medial Orbitofrontal Cortex | -0.550 (0.506) | -0.020 | 0.330 |
| Left Middle Temporal Gyrus | -0.657 (0.888) | -0.014 | 0.523 |
| Left Paracentral Lobule | -0.999 (0.715) | -0.026 | 0.204 |
| Left Parahippocampal Gyrus | 0.726 (0.662) | 0.020 | 0.327 |
| Left Pars Opercularis | 0.245 (0.963) | 0.005 | 0.839 |
| Left Pars Orbitalis | -0.959 (0.737) | -0.024 | 0.239 |
| Left Pars Triangularis | -1.425 (0.995) | -0.027 | 0.191 |
| Left Pericalcarine Cortex | 1.610 (0.935) | 0.032 | 0.111 |
| Left Postcentral Gyrus | -0.259 (0.926) | -0.005 | 0.822 |
| Left Posterior Cingulate Cortex | -1.501 (1.046) | -0.028 | 0.191 |
| Left Precentral Gyrus | 0.095 (0.860) | 0.002 | 0.926 |
| Left Precuneus | -2.067 (0.889) | -0.045 | 0.027 |
| Left Rostral Anterior Cingulate Cortex | -0.331 (0.746) | -0.008 | 0.718 |
| Left Rostral Middle Frontal Gyrus | -0.381 (0.796) | -0.009 | 0.694 |
| Left Superior Frontal Gyrus | -0.866 (0.888) | -0.019 | 0.387 |
| Left Superior Parietal Lobule | -0.945 (0.753) | -0.024 | 0.257 |
| Left Superior Temporal Gyrus | 0.033 (0.941) | 0.001 | 0.979 |
| Left Supramarginal Gyrus | -0.474 (0.886) | -0.010 | 0.655 |
| Left Temporal Pole | -0.521 (0.431) | -0.022 | 0.275 |
| Left Transverse Temporal Gyrus (Heschl’s Gyrus) | -0.783 (0.900) | -0.016 | 0.446 |
| Right Banks of the Superior Temporal Sulcus | -1.000 (0.812) | -0.023 | 0.266 |
| Right Caudal Anterior Cingulate Cortex | 0.030 (1.016) | 0.001 | 0.980 |
| Right Caudal Middle Frontal Gyrus | -0.998 (0.786) | -0.024 | 0.252 |
| Right Cuneus | 0.179 (0.918) | 0.004 | 0.877 |
| Right Entorhinal Cortex | 0.172 (0.476) | 0.007 | 0.772 |
| Right Frontal Pole | 0.075 (0.449) | 0.003 | 0.897 |
| Right Fusiform Gyrus | -0.721 (0.912) | -0.015 | 0.495 |
| Right Inferior Parietal Lobule | -1.596 (0.970) | -0.032 | 0.129 |
| Right Inferior Temporal Gyrus | -1.781 (0.940) | -0.036 | 0.076 |
| Right Insular Cortex | 0.437 (0.518) | 0.016 | 0.462 |
| Right Isthmus of the Cingulate Cortex | 0.513 (0.841) | 0.012 | 0.601 |
| Right Lateral Occipital Cortex | 0.085 (0.983) | 0.002 | 0.941 |
| Right Lateral Orbitofrontal Cortex | -0.279 (0.566) | -0.009 | 0.685 |
| Right Lingual Gyrus | 0.365 (1.099) | 0.007 | 0.787 |
| Right Medial Orbitofrontal Cortex | -0.361 (0.581) | -0.011 | 0.596 |
| Right Middle Temporal Gyrus | 0.025 (0.979) | 0.000 | 0.980 |
| Right Paracentral Lobule | -0.278 (0.754) | -0.007 | 0.768 |
| Right Parahippocampal Gyrus | 0.613 (0.591) | 0.019 | 0.353 |
| Right Pars Opercularis | -0.620 (0.830) | -0.014 | 0.522 |
| Right Pars Orbitalis | -0.155 (0.777) | -0.004 | 0.877 |
| Right Pars Triangularis | 0.114 (0.810) | 0.003 | 0.908 |
| Right Pericalcarine Cortex | 0.202 (0.887) | 0.004 | 0.858 |
| Right Postcentral Gyrus | -0.949 (0.883) | -0.020 | 0.334 |
| Right Posterior Cingulate Cortex | -1.007 (1.101) | -0.018 | 0.421 |
| Right Precentral Gyrus | -0.516 (0.726) | -0.013 | 0.538 |
| Right Precuneus | -0.618 (0.835) | -0.014 | 0.523 |
| Right Rostral Anterior Cingulate Cortex | 0.541 (0.862) | 0.012 | 0.594 |
| Right Rostral Middle Frontal Gyrus | -0.476 (0.757) | -0.012 | 0.594 |
| Right Superior Frontal Gyrus | -0.549 (0.771) | -0.014 | 0.538 |
| Right Superior Parietal Lobule | -1.182 (0.667) | -0.033 | 0.100 |
| Right Superior Temporal Gyrus | -0.143 (0.964) | -0.003 | 0.906 |
| Right Supramarginal Gyrus | -0.222 (0.759) | -0.005 | 0.815 |
| Right Temporal Pole | -0.482 (0.444) | -0.020 | 0.330 |
| Right Transverse Temporal Gyrus (Heschl’s Gyrus) | 0.277 (0.806) | 0.006 | 0.780 |

**Supplementary Table 47: Longitudinal associations between regional centile change scores and Picture Sequence Memory Task change scores**

Values shown are unstandardised and standardised beta coefficients with associated standard errors (SE) for each region. P values are false discovery rate (FDR) corrected to account for multiple comparisons. All analyses controlled for age and baseline task scores, included random intercepts for testing site and family ID (to account for sibling pairs).

| **Region** | **Unstandardised estimate (SE)** | **Standardised estimate (SE)** | **P value (FDR corrected)** |
| --- | --- | --- | --- |
| Left Banks of the Superior Temporal Sulcus | 1.608 (1.658) | 0.022 (0.021) | 0.499 |
| Left Caudal Anterior Cingulate Cortex | 3.989 (2.267) | 0.041 (0.027) | 0.129 |
| Left Caudal Middle Frontal Gyrus | 2.743 (1.877) | 0.034 (0.008) | 0.227 |
| Left Cuneus | 4.682 (1.876) | 0.058 (0.026) | 0.023 |
| Left Entorhinal Cortex | 0.289 (1.014) | 0.006 (0.002) | 0.905 |
| Left Frontal Pole | -1.222 (0.934) | -0.030 (0.025) | 0.297 |
| Left Fusiform Gyrus | 2.114 (1.834) | 0.028 (0.032) | 0.381 |
| Left Inferior Parietal Lobule | 1.209 (2.069) | 0.014 (0.003) | 0.776 |
| Left Inferior Temporal Gyrus | 3.910 (1.920) | 0.049 (0.007) | 0.072 |
| Left Insular Cortex | 2.144 (1.099) | 0.044 (0.029) | 0.086 |
| Left Isthmus of the Cingulate Cortex | 3.172 (1.895) | 0.040 (0.008) | 0.153 |
| Left Lateral Occipital Cortex | 6.632 (1.994) | 0.081 (0.024) | 0.002 |
| Left Lateral Orbitofrontal Cortex | 1.970 (1.430) | 0.032 (0.039) | 0.263 |
| Left Lingual Gyrus | 6.335 (2.342) | 0.066 (0.036) | 0.013 |
| Left Medial Orbitofrontal Cortex | 0.960 (1.037) | 0.021 (0.020) | 0.524 |
| Left Middle Temporal Gyrus | 4.688 (1.832) | 0.062 (0.014) | 0.019 |
| Left Paracentral Lobule | 3.816 (1.466) | 0.060 (0.026) | 0.017 |
| Left Parahippocampal Gyrus | 1.146 (1.355) | 0.019 (0.020) | 0.580 |
| Left Pars Opercularis | 3.813 (1.987) | 0.045 (0.005) | 0.091 |
| Left Pars Orbitalis | -0.662 (1.515) | -0.010 (0.024) | 0.843 |
| Left Pars Triangularis | 1.732 (2.050) | 0.020 (0.027) | 0.580 |
| Left Pericalcarine Cortex | 5.626 (1.919) | 0.068 (0.032) | 0.006 |
| Left Postcentral Gyrus | 4.408 (1.901) | 0.055 (0.005) | 0.036 |
| Left Posterior Cingulate Cortex | 3.326 (2.157) | 0.037 (0.028) | 0.195 |
| Left Precentral Gyrus | 3.452 (1.770) | 0.047 (0.002) | 0.086 |
| Left Precuneus | 3.910 (1.846) | 0.052 (0.045) | 0.060 |
| Left Rostral Anterior Cingulate Cortex | 0.683 (1.532) | 0.010 (0.008) | 0.843 |
| Left Rostral Middle Frontal Gyrus | 0.936 (1.632) | 0.014 (0.009) | 0.781 |
| Left Superior Frontal Gyrus | 1.903 (1.825) | 0.025 (0.019) | 0.451 |
| Left Superior Parietal Lobule | 3.549 (1.544) | 0.054 (0.024) | 0.038 |
| Left Superior Temporal Gyrus | 5.969 (1.926) | 0.075 (0.001) | 0.004 |
| Left Supramarginal Gyrus | 4.279 (1.819) | 0.056 (0.010) | 0.034 |
| Left Temporal Pole | 0.337 (0.878) | 0.009 (0.022) | 0.860 |
| Left Transverse Temporal Gyrus (Heschl’s Gyrus) | 3.046 (1.856) | 0.038 (0.016) | 0.162 |
| Right Banks of the Superior Temporal Sulcus | -0.049 (1.654) | -0.001 (0.023) | 0.988 |
| Right Caudal Anterior Cingulate Cortex | -0.388 (2.076) | -0.004 (0.001) | 0.931 |
| Right Caudal Middle Frontal Gyrus | 1.153 (1.633) | 0.016 (0.024) | 0.685 |
| Right Cuneus | 7.807 (1.872) | 0.097 (0.004) | 0.000 |
| Right Entorhinal Cortex | 1.725 (0.972) | 0.040 (0.007) | 0.125 |
| Right Frontal Pole | -0.642 (0.911) | -0.016 (0.003) | 0.685 |
| Right Fusiform Gyrus | 1.745 (1.883) | 0.022 (0.015) | 0.524 |
| Right Inferior Parietal Lobule | 4.703 (1.983) | 0.057 (0.032) | 0.032 |
| Right Inferior Temporal Gyrus | 1.496 (1.940) | 0.019 (0.036) | 0.638 |
| Right Insular Cortex | 2.673 (1.055) | 0.058 (0.016) | 0.021 |
| Right Isthmus of the Cingulate Cortex | 3.528 (1.721) | 0.048 (0.012) | 0.070 |
| Right Lateral Occipital Cortex | 7.922 (2.012) | 0.094 (0.002) | 0.000 |
| Right Lateral Orbitofrontal Cortex | 2.592 (1.167) | 0.052 (0.009) | 0.046 |
| Right Lingual Gyrus | 4.614 (2.312) | 0.050 (0.007) | 0.079 |
| Right Medial Orbitofrontal Cortex | 2.061 (1.186) | 0.040 (0.011) | 0.134 |
| Right Middle Temporal Gyrus | 6.306 (2.008) | 0.076 (0.000) | 0.003 |
| Right Paracentral Lobule | 1.608 (1.545) | 0.024 (0.007) | 0.017 |
| Right Parahippocampal Gyrus | 0.688 (1.205) | 0.013 (0.019) | 0.781 |
| Right Pars Opercularis | 0.532 (1.698) | 0.007 (0.014) | 0.891 |
| Right Pars Orbitalis | 5.457 (1.584) | 0.080 (0.004) | 0.001 |
| Right Pars Triangularis | 3.479 (1.651) | 0.049 (0.003) | 0.061 |
| Right Pericalcarine Cortex | 5.054 (1.820) | 0.064 (0.004) | 0.010 |
| Right Postcentral Gyrus | 2.152 (1.815) | 0.028 (0.020) | 0.365 |
| Right Posterior Cingulate Cortex | 2.147 (2.272) | 0.023 (0.018) | 0.515 |
| Right Precentral Gyrus | 1.750 (1.502) | 0.028 (0.013) | 0.375 |
| Right Precuneus | 4.803 (1.760) | 0.068 (0.014) | 0.012 |
| Right Rostral Anterior Cingulate Cortex | -0.765 (1.759) | -0.010 (0.012) | 0.843 |
| Right Rostral Middle Frontal Gyrus | 0.690 (1.541) | 0.011 (0.012) | 0.843 |
| Right Superior Frontal Gyrus | 1.436 (1.573) | 0.022 (0.014) | 0.531 |
| Right Superior Parietal Lobule | 2.723 (1.375) | 0.046 (0.033) | 0.081 |
| Right Superior Temporal Gyrus | 3.281 (1.979) | 0.040 (0.003) | 0.157 |
| Right Supramarginal Gyrus | 0.341 (1.547) | 0.005 (0.005) | 0.923 |
| Right Temporal Pole | 1.460 (0.901) | 0.036 (0.020) | 0.168 |
| Right Transverse Temporal Gyrus (Heschl’s Gyrus) | 0.550 (1.642) | 0.008 (0.006) | 0.889 |

**Supplementary Table 48: Longitudinal associations between regional centile change scores and Oral Reading Recognition Task change scores**

Values shown are unstandardised and standardised beta coefficients with associated standard errors (SE) for each region. P values are false discovery rate (FDR) corrected to account for multiple comparisons. All analyses controlled for age and baseline task scores, included random intercepts for testing site and family ID (to account for sibling pairs).

| **Region** | **Unstandardised estimate (SE)** | **Standardised estimate (SE)** | **P value (FDR corrected)** |
| --- | --- | --- | --- |
| Left Banks of the Superior Temporal Sulcus | -0.173 (0.596) | -0.007 (0.007) | 0.817 |
| Left Caudal Anterior Cingulate Cortex | -0.217 (0.816) | -0.007 (0.007) | 0.834 |
| Left Caudal Middle Frontal Gyrus | -0.743 (0.693) | -0.027 (0.027) | 0.487 |
| Left Cuneus | -0.324 (0.675) | -0.012 (0.012) | 0.727 |
| Left Entorhinal Cortex | -0.812 (0.364) | -0.053 (0.053) | 0.050 |
| Left Frontal Pole | -0.286 (0.335) | -0.021 (0.021) | 0.566 |
| Left Fusiform Gyrus | -0.968 (0.661) | -0.038 (0.038) | 0.264 |
| Left Inferior Parietal Lobule | -1.242 (0.750) | -0.043 (0.043) | 0.185 |
| Left Inferior Temporal Gyrus | -1.096 (0.687) | -0.040 (0.040) | 0.208 |
| Left Insular Cortex | 0.681 (0.398) | 0.042 (0.042) | 0.166 |
| Left Isthmus of the Cingulate Cortex | -0.269 (0.682) | -0.010 (0.010) | 0.760 |
| Left Lateral Occipital Cortex | 0.288 (0.729) | 0.010 (0.010) | 0.760 |
| Left Lateral Orbitofrontal Cortex | -0.010 (0.511) | -0.001 (0.001) | 0.984 |
| Left Lingual Gyrus | -0.374 (0.851) | -0.012 (0.012) | 0.748 |
| Left Medial Orbitofrontal Cortex | 0.078 (0.372) | 0.005 (0.005) | 0.865 |
| Left Middle Temporal Gyrus | 0.085 (0.664) | 0.003 (0.003) | 0.922 |
| Left Paracentral Lobule | -0.542 (0.525) | -0.025 (0.025) | 0.511 |
| Left Parahippocampal Gyrus | -0.174 (0.490) | -0.009 (0.009) | 0.780 |
| Left Pars Opercularis | -1.625 (0.717) | -0.057 (0.057) | 0.046 |
| Left Pars Orbitalis | -0.141 (0.548) | -0.006 (0.006) | 0.837 |
| Left Pars Triangularis | -0.807 (0.741) | -0.027 (0.027) | 0.479 |
| Left Pericalcarine Cortex | -0.167 (0.697) | -0.006 (0.006) | 0.845 |
| Left Postcentral Gyrus | -1.211 (0.686) | -0.045 (0.045) | 0.150 |
| Left Posterior Cingulate Cortex | -0.017 (0.776) | -0.001 (0.001) | 0.984 |
| Left Precentral Gyrus | -0.050 (0.647) | -0.002 (0.002) | 0.952 |
| Left Precuneus | -1.505 (0.667) | -0.059 (0.059) | 0.047 |
| Left Rostral Anterior Cingulate Cortex | -0.049 (0.552) | -0.002 (0.002) | 0.947 |
| Left Rostral Middle Frontal Gyrus | -0.081 (0.586) | -0.003 (0.003) | 0.918 |
| Left Superior Frontal Gyrus | -0.662 (0.664) | -0.026 (0.026) | 0.518 |
| Left Superior Parietal Lobule | -0.692 (0.576) | -0.031 (0.031) | 0.409 |
| Left Superior Temporal Gyrus | 0.038 (0.694) | 0.001 (0.001) | 0.963 |
| Left Supramarginal Gyrus | -0.507 (0.685) | -0.019 (0.019) | 0.586 |
| Left Temporal Pole | -0.549 (0.316) | -0.041 (0.041) | 0.158 |
| Left Transverse Temporal Gyrus (Heschl’s Gyrus) | 0.527 (0.669) | 0.020 (0.020) | 0.566 |
| Right Banks of the Superior Temporal Sulcus | -0.936 (0.593) | -0.001 (0.001) | 0.214 |
| Right Caudal Anterior Cingulate Cortex | -0.688 (0.745) | -0.004 (0.004) | 0.561 |
| Right Caudal Middle Frontal Gyrus | -0.270 (0.585) | 0.016 (0.016) | 0.740 |
| Right Cuneus | -1.040 (0.679) | 0.097 (0.097) | 0.233 |
| Right Entorhinal Cortex | -0.025 (0.350) | 0.040 (0.040) | 0.954 |
| Right Frontal Pole | -0.292 (0.328) | -0.016 (0.016) | 0.561 |
| Right Fusiform Gyrus | -0.915 (0.684) | 0.022 (0.022) | 0.330 |
| Right Inferior Parietal Lobule | -0.603 (0.715) | 0.057 (0.057) | 0.566 |
| Right Inferior Temporal Gyrus | -0.292 (0.701) | 0.019 (0.019) | 0.752 |
| Right Insular Cortex | -0.234 (0.380) | 0.058 (0.058) | 0.659 |
| Right Isthmus of the Cingulate Cortex | 0.701 (0.619) | 0.048 (0.048) | 0.449 |
| Right Lateral Occipital Cortex | -0.901 (0.730) | 0.094 (0.094) | 0.392 |
| Right Lateral Orbitofrontal Cortex | 0.227 (0.418) | 0.052 (0.052) | 0.699 |
| Right Lingual Gyrus | -0.278 (0.843) | 0.050 (0.050) | 0.797 |
| Right Medial Orbitofrontal Cortex | 0.160 (0.423) | 0.040 (0.040) | 0.767 |
| Right Middle Temporal Gyrus | -0.577 (0.720) | 0.076 (0.076) | 0.566 |
| Right Paracentral Lobule | -1.463 (0.554) | 0.024 (0.024) | 0.017 |
| Right Parahippocampal Gyrus | -0.434 (0.434) | 0.013 (0.013) | 0.518 |
| Right Pars Opercularis | -0.363 (0.608) | 0.007 (0.007) | 0.665 |
| Right Pars Orbitalis | -0.245 (0.577) | 0.080 (0.080) | 0.752 |
| Right Pars Triangularis | -0.294 (0.604) | 0.049 (0.049) | 0.725 |
| Right Pericalcarine Cortex | -0.190 (0.657) | 0.064 (0.064) | 0.817 |
| Right Postcentral Gyrus | 0.260 (0.652) | 0.028 (0.028) | 0.760 |
| Right Posterior Cingulate Cortex | -0.311 (0.818) | 0.023 (0.023) | 0.767 |
| Right Precentral Gyrus | 0.238 (0.548) | 0.028 (0.028) | 0.749 |
| Right Precuneus | -0.730 (0.634) | 0.068 (0.068) | 0.438 |
| Right Rostral Anterior Cingulate Cortex | -0.121 (0.633) | -0.010 (0.010) | 0.877 |
| Right Rostral Middle Frontal Gyrus | -0.577 (0.553) | 0.011 (0.011) | 0.505 |
| Right Superior Frontal Gyrus | -0.457 (0.564) | 0.022 (0.022) | 0.566 |
| Right Superior Parietal Lobule | -0.650 (0.494) | 0.046 (0.046) | 0.341 |
| Right Superior Temporal Gyrus | -0.179 (0.711) | 0.040 (0.040) | 0.838 |
| Right Supramarginal Gyrus | -0.331 (0.552) | 0.005 (0.005) | 0.665 |
| Right Temporal Pole | -0.258 (0.326) | 0.036 (0.036) | 0.566 |
| Right Transverse Temporal Gyrus (Heschl’s Gyrus) | -0.310 (0.589) | 0.008 (0.008) | 0.705 |

**Supplementary Table 49: Longitudinal associations between regional centile change scores and Pattern Comparison Processing Speed change scores**

Values shown are unstandardised and standardised beta coefficients with associated standard errors (SE) for each region. P values are false discovery rate (FDR) corrected to account for multiple comparisons. All analyses controlled for age and baseline task scores, included random intercepts for testing site and family ID (to account for sibling pairs).

| **Region** | **Unstandardised estimate (SE)** | **Standardised estimate (SE)** | **P value (FDR corrected)** |
| --- | --- | --- | --- |
| Left Banks of the Superior Temporal Sulcus | -2.053 (1.821) | -0.026 | 0.260 |
| Left Caudal Anterior Cingulate Cortex | 2.810 (2.471) | 0.026 | 0.256 |
| Left Caudal Middle Frontal Gyrus | -1.271 (2.042) | -0.014 | 0.534 |
| Left Cuneus | 1.480 (2.051) | 0.017 | 0.471 |
| Left Entorhinal Cortex | -0.718 (1.107) | -0.014 | 0.517 |
| Left Frontal Pole | -1.191 (1.020) | -0.026 | 0.243 |
| Left Fusiform Gyrus | -0.572 (2.000) | -0.007 | 0.775 |
| Left Inferior Parietal Lobule | -2.563 (2.278) | -0.027 | 0.261 |
| Left Inferior Temporal Gyrus | 0.631 (2.137) | 0.007 | 0.768 |
| Left Insular Cortex | -1.374 (1.197) | -0.026 | 0.251 |
| Left Isthmus of the Cingulate Cortex | 0.315 (2.048) | 0.004 | 0.878 |
| Left Lateral Occipital Cortex | 1.853 (2.169) | 0.020 | 0.393 |
| Left Lateral Orbitofrontal Cortex | -0.846 (1.572) | -0.012 | 0.590 |
| Left Lingual Gyrus | 2.676 (2.548) | 0.025 | 0.294 |
| Left Medial Orbitofrontal Cortex | 0.086 (1.130) | 0.002 | 0.939 |
| Left Middle Temporal Gyrus | 0.881 (2.002) | 0.011 | 0.660 |
| Left Paracentral Lobule | -0.683 (1.590) | -0.010 | 0.667 |
| Left Parahippocampal Gyrus | 0.635 (1.473) | 0.010 | 0.666 |
| Left Pars Opercularis | 0.663 (2.149) | 0.007 | 0.758 |
| Left Pars Orbitalis | -1.712 (1.646) | -0.024 | 0.298 |
| Left Pars Triangularis | -3.076 (2.220) | -0.032 | 0.166 |
| Left Pericalcarine Cortex | 1.604 (2.108) | 0.017 | 0.447 |
| Left Postcentral Gyrus | -3.891 (2.055) | -0.044 | 0.059 |
| Left Posterior Cingulate Cortex | -1.560 (2.390) | -0.016 | 0.514 |
| Left Precentral Gyrus | -2.074 (1.920) | -0.025 | 0.280 |
| Left Precuneus | -1.600 (2.043) | -0.019 | 0.434 |
| Left Rostral Anterior Cingulate Cortex | 0.784 (1.669) | 0.011 | 0.638 |
| Left Rostral Middle Frontal Gyrus | -0.320 (1.777) | -0.004 | 0.857 |
| Left Superior Frontal Gyrus | -3.733 (1.985) | -0.045 | 0.060 |
| Left Superior Parietal Lobule | -2.770 (1.685) | -0.038 | 0.100 |
| Left Superior Temporal Gyrus | -0.236 (2.150) | -0.003 | 0.913 |
| Left Supramarginal Gyrus | 0.984 (1.986) | 0.012 | 0.620 |
| Left Temporal Pole | -0.287 (0.964) | -0.007 | 0.766 |
| Left Transverse Temporal Gyrus (Heschl’s Gyrus) | -2.070 (2.007) | -0.024 | 0.302 |
| Right Banks of the Superior Temporal Sulcus | -4.406 (1.806) | -0.055 | 0.015 |
| Right Caudal Anterior Cingulate Cortex | 2.052 (2.268) | 0.021 | 0.366 |
| Right Caudal Middle Frontal Gyrus | -2.625 (1.756) | -0.034 | 0.135 |
| Right Cuneus | 1.215 (2.056) | 0.014 | 0.555 |
| Right Entorhinal Cortex | 0.007 (1.061) | 0.000 | 0.995 |
| Right Frontal Pole | -0.032 (0.997) | -0.001 | 0.975 |
| Right Fusiform Gyrus | -0.480 (2.045) | -0.006 | 0.814 |
| Right Inferior Parietal Lobule | -2.981 (2.165) | -0.033 | 0.169 |
| Right Inferior Temporal Gyrus | 0.215 (2.147) | 0.002 | 0.920 |
| Right Insular Cortex | -0.841 (1.157) | -0.016 | 0.468 |
| Right Isthmus of the Cingulate Cortex | -0.208 (1.874) | -0.003 | 0.912 |
| Right Lateral Occipital Cortex | 2.962 (2.192) | 0.032 | 0.177 |
| Right Lateral Orbitofrontal Cortex | -0.150 (1.268) | -0.003 | 0.906 |
| Right Lingual Gyrus | 1.584 (2.490) | 0.015 | 0.525 |
| Right Medial Orbitofrontal Cortex | 0.898 (1.292) | 0.016 | 0.487 |
| Right Middle Temporal Gyrus | -1.544 (2.181) | -0.017 | 0.479 |
| Right Paracentral Lobule | -2.263 (1.677) | -0.031 | 0.178 |
| Right Parahippocampal Gyrus | -0.258 (1.315) | -0.004 | 0.845 |
| Right Pars Opercularis | -3.675 (1.845) | -0.046 | 0.047 |
| Right Pars Orbitalis | -2.289 (1.726) | -0.030 | 0.185 |
| Right Pars Triangularis | -1.066 (1.824) | -0.013 | 0.559 |
| Right Pericalcarine Cortex | 1.188 (1.981) | 0.014 | 0.549 |
| Right Postcentral Gyrus | -0.287 (1.965) | -0.003 | 0.884 |
| Right Posterior Cingulate Cortex | -2.596 (2.454) | -0.025 | 0.290 |
| Right Precentral Gyrus | -0.442 (1.658) | -0.006 | 0.790 |
| Right Precuneus | 2.367 (1.903) | 0.030 | 0.214 |
| Right Rostral Anterior Cingulate Cortex | 0.741 (1.918) | 0.009 | 0.699 |
| Right Rostral Middle Frontal Gyrus | -3.085 (1.674) | -0.042 | 0.066 |
| Right Superior Frontal Gyrus | -2.914 (1.709) | -0.040 | 0.088 |
| Right Superior Parietal Lobule | -1.561 (1.499) | -0.024 | 0.298 |
| Right Superior Temporal Gyrus | -1.662 (2.147) | -0.018 | 0.439 |
| Right Supramarginal Gyrus | 0.678 (1.685) | 0.009 | 0.687 |
| Right Temporal Pole | 1.405 (0.985) | 0.032 | 0.154 |
| Right Transverse Temporal Gyrus (Heschl’s Gyrus) | -0.110 (1.789) | -0.001 | 0.951 |

**Supplementary Table 50: Longitudinal associations between regional centile change scores and Picture Vocabulary Task change scores**

Values shown are unstandardised and standardised beta coefficients with associated standard errors (SE) for each region. P values are false discovery rate (FDR) corrected to account for multiple comparisons. All analyses controlled for age and baseline task scores, included random intercepts for testing site and family ID (to account for sibling pairs).

| **Region** | **Unstandardised estimate (SE)** | **Standardised estimate** | **P value (FDR corrected)** |
| --- | --- | --- | --- |
| Left Banks of the Superior Temporal Sulcus | -0.463 (0.728) | -0.015 | 0.604 |
| Left Caudal Anterior Cingulate Cortex | 0.019 (0.993) | 0.000 | 0.985 |
| Left Caudal Middle Frontal Gyrus | 1.060 (0.846) | 0.031 | 0.367 |
| Left Cuneus | 0.504 (0.825) | 0.015 | 0.616 |
| Left Entorhinal Cortex | 0.455 (0.448) | 0.024 | 0.473 |
| Left Frontal Pole | 0.343 (0.412) | 0.020 | 0.532 |
| Left Fusiform Gyrus | -0.865 (0.806) | -0.027 | 0.446 |
| Left Inferior Parietal Lobule | 0.658 (0.924) | 0.018 | 0.566 |
| Left Inferior Temporal Gyrus | 0.080 (0.843) | 0.002 | 0.945 |
| Left Insular Cortex | 0.461 (0.490) | 0.023 | 0.498 |
| Left Isthmus of the Cingulate Cortex | -0.425 (0.833) | -0.013 | 0.675 |
| Left Lateral Occipital Cortex | 1.292 (0.890) | 0.037 | 0.268 |
| Left Lateral Orbitofrontal Cortex | 0.828 (0.624) | 0.032 | 0.333 |
| Left Lingual Gyrus | 0.918 (1.038) | 0.023 | 0.511 |
| Left Medial Orbitofrontal Cortex | 0.262 (0.457) | 0.014 | 0.635 |
| Left Middle Temporal Gyrus | 0.837 (0.812) | 0.026 | 0.465 |
| Left Paracentral Lobule | -0.229 (0.645) | -0.009 | 0.777 |
| Left Parahippocampal Gyrus | 0.718 (0.600) | 0.029 | 0.396 |
| Left Pars Opercularis | -0.022 (0.873) | -0.001 | 0.983 |
| Left Pars Orbitalis | 0.986 (0.669) | 0.036 | 0.260 |
| Left Pars Triangularis | 0.053 (0.906) | 0.001 | 0.964 |
| Left Pericalcarine Cortex | 1.736 (0.849) | 0.050 | 0.081 |
| Left Postcentral Gyrus | -0.728 (0.839) | -0.021 | 0.517 |
| Left Posterior Cingulate Cortex | 0.198 (0.947) | 0.005 | 0.866 |
| Left Precentral Gyrus | 0.787 (0.790) | 0.025 | 0.482 |
| Left Precuneus | -0.343 (0.815) | -0.011 | 0.728 |
| Left Rostral Anterior Cingulate Cortex | 0.099 (0.675) | 0.004 | 0.906 |
| Left Rostral Middle Frontal Gyrus | 0.159 (0.715) | 0.006 | 0.863 |
| Left Superior Frontal Gyrus | 0.350 (0.808) | 0.011 | 0.724 |
| Left Superior Parietal Lobule | 0.295 (0.704) | 0.011 | 0.728 |
| Left Superior Temporal Gyrus | 1.428 (0.850) | 0.042 | 0.180 |
| Left Supramarginal Gyrus | 1.186 (0.836) | 0.036 | 0.283 |
| Left Temporal Pole | 0.330 (0.387) | 0.020 | 0.526 |
| Left Transverse Temporal Gyrus (Heschl’s Gyrus) | 0.727 (0.819) | 0.022 | 0.511 |
| Right Banks of the Superior Temporal Sulcus | 0.236 (0.728) | 0.008 | 0.796 |
| Right Caudal Anterior Cingulate Cortex | 0.498 (0.910) | 0.013 | 0.652 |
| Right Caudal Middle Frontal Gyrus | 0.832 (0.714) | 0.028 | 0.410 |
| Right Cuneus | 0.983 (0.828) | 0.029 | 0.401 |
| Right Entorhinal Cortex | 0.093 (0.429) | 0.005 | 0.863 |
| Right Frontal Pole | -0.107 (0.404) | -0.006 | 0.835 |
| Right Fusiform Gyrus | -0.741 (0.834) | -0.023 | 0.511 |
| Right Inferior Parietal Lobule | 0.138 (0.875) | 0.004 | 0.902 |
| Right Inferior Temporal Gyrus | -0.561 (0.854) | -0.016 | 0.592 |
| Right Insular Cortex | 0.371 (0.465) | 0.019 | 0.534 |
| Right Isthmus of the Cingulate Cortex | 2.286 (0.754) | 0.074 | 0.005 |
| Right Lateral Occipital Cortex | 0.665 (0.897) | 0.019 | 0.549 |
| Right Lateral Orbitofrontal Cortex | 0.412 (0.512) | 0.019 | 0.534 |
| Right Lingual Gyrus | -0.275 (1.016) | -0.007 | 0.832 |
| Right Medial Orbitofrontal Cortex | 0.601 (0.522) | 0.027 | 0.413 |
| Right Middle Temporal Gyrus | 1.501 (0.881) | 0.043 | 0.172 |
| Right Paracentral Lobule | -0.896 (0.678) | -0.032 | 0.334 |
| Right Parahippocampal Gyrus | -0.179 (0.534) | -0.008 | 0.790 |
| Right Pars Opercularis | 0.732 (0.747) | 0.024 | 0.486 |
| Right Pars Orbitalis | 1.113 (0.702) | 0.039 | 0.215 |
| Right Pars Triangularis | -0.165 (0.743) | -0.005 | 0.863 |
| Right Pericalcarine Cortex | 0.597 (0.801) | 0.018 | 0.549 |
| Right Postcentral Gyrus | -0.242 (0.799) | -0.007 | 0.810 |
| Right Posterior Cingulate Cortex | 1.768 (0.996) | 0.044 | 0.149 |
| Right Precentral Gyrus | 1.016 (0.668) | 0.038 | 0.239 |
| Right Precuneus | 0.040 (0.759) | 0.001 | 0.965 |
| Right Rostral Anterior Cingulate Cortex | 1.240 (0.779) | 0.038 | 0.214 |
| Right Rostral Middle Frontal Gyrus | 1.038 (0.682) | 0.037 | 0.239 |
| Right Superior Frontal Gyrus | 0.468 (0.693) | 0.017 | 0.586 |
| Right Superior Parietal Lobule | -0.368 (0.601) | -0.015 | 0.616 |
| Right Superior Temporal Gyrus | 1.263 (0.869) | 0.036 | 0.268 |
| Right Supramarginal Gyrus | -0.045 (0.679) | -0.002 | 0.961 |
| Right Temporal Pole | -0.081 (0.401) | -0.005 | 0.868 |
| Right Transverse Temporal Gyrus (Heschl’s Gyrus) | 0.062 (0.726) | 0.002 | 0.949 |

**Supplementary Table 51: Regions mediating the associations between environmental variables and changes in the Picture Sequence Memory Task**

The table summarises the mediation analyses ran across regions that showed both environment-brain and brain-behaviour associations. The ACME (Average Causal Mediation Effect) estimate is reported, which reflects the indirect effect of the predictor on the outcome that is transmitted through the mediator, as well as proportion mediated values, which indicate the estimated fraction of the total effect explained by the mediator. Associated ACME confidence intervals and FDR-corrected p values are also reported. P values are false discovery rate (FDR) corrected to account for multiple comparisons. Analyses controlled for age and sex and included random intercepts for testing site and family ID (to account for sibling pairs).

| **Predictor** | **Region** | **ACME estimate** | **ACME CI** | **Proportion mediated** | ***p*_FDR_** |
| --- | --- | --- | --- | --- | --- |
| Health/environment | Left cuneus | 0.009 | 0.000 – 0.022 | 0.025 | 0.062 |
|  | Left Lateral Occipital Gyrus | 0.014 | 0.002 – 0.028 | 0.036 | 0.030 |
|  | Left Lingual Gyrus | 0.011 | -0.001 – 0.026 | 0.029 | 0.076 |
|  | Left Middle Temporal Gyrus | 0.008 | -0.001 – 0.020 | 0.022 | 0.088 |
|  | Left Superior Temporal Gyrus | 0.011 | 0.001 – 0.025 | 0.029 | 0.053 |
|  | Left Supramarginal Gyrus | 0.007 | 0.000 – 0.017 | 0.020 | 0.058 |
|  | Right Cuneus | 0.020 | 0.005 – 0.038 | 0.055 | 0.012 |
|  | Right Inferior Parietal Gyrus | 0.008 | 0.000 – 0.020 | 0.022 | 0.058 |
|  | Right insula | 0.008 | 0.000 – 0.020 | 0.022 | 0.058 |
|  | Right Lateral Occipital Gyrus | 0.015 | 0.003 – 0.029 | 0.040 | 0.012 |
|  | Right Pericalcarine Gyrus | 0.009 | 0.000 – 0.020 | 0.024 | 0.062 |
|  | Right Precuneus | 0.010 | 0.000 – 0.022 | 0.026 | 0.058 |
| Social/economic | Left Cuneus | 0.009 | 0.001 – 0.021 | 0.033 | 0.060 |
|  | Left Lateral Occipital Gyrus | 0.014 | 0.003 – 0.029 | 0.051 | 0.040 |
|  | Left Lingual Gyrus | 0.007 | -0.001 – 0.018 | 0.026 | 0.143 |
|  | Right Cuneus | 0.003 | -0.009 – 0.015 | 0.009 | 0.732 |
|  | Right Pericalcarine Gyrus | 0.002 | -0.006 – 0.011 | 0.007 | 0.732 |
| Education | Left Caudal Middle Frontal Gyrus | 0.007 | -0.001 – -0.020 | 0.035 | 0.096 |
|  | Left Cuneus | 0.003 | -0.004 – 0.013 | 0.015 | 0.148 |
|  | Left Lateral Occipital Gyrus | 0.009 | -0.000 – 0.013 | 0.042 | 0.148 |
|  | Left Lingual Gyrus | 0.007 | -0.000 – 0.022 | 0.033 | 0.148 |
|  | Left Superior Frontal Gyrus | 0.003 | -0.000 – 0.019 | 0.013 | 0.240 |
|  | Right Cuneus | 0.015 | -0.003 – 0.032 | 0.078 | 0.397 |
|  | Right Inferior Temporal Gyrus | -0.000 | -0.005 – 0.003 | -0.001 | 0.448 |
|  | Right Pericalcarine Gyrus | 0.006 | -0.002 – 0.017 | 0.028 | 0.892 |
| General LF | Left Caudal Middle Frontal Gyrus | 0.008 | -0.001 – 0.020 | 0.035 | 0.180 |
|  | Left Cuneus | 0.003 | -0.004 – 0.014 | 0.015 | 0.434 |
|  | Left Lateral Occipital Gyrus | 0.009 | -0.001 – 0.022 | 0.042 | 0.180 |
|  | Left Lingual Gyrus | 0.007 | 0.000 – 0.018 | 0.033 | 0.180 |
|  | Left Superior Temporal Gyrus | 0.003 | -0.002 – 0.010 | 0.013 | 0.434 |
|  | Right Cuneus | 0.015 | 0.003 – 0.030 | 0.068 | 0.048 |
|  | Right Inferior Temporal Gyrus | 0.000 | -0.005 – 0.004 | -0.001 | 0.884 |
|  | Right Pericalcarine Gyrus | 0.006 | -0.002 – 0.017 | 0.028 | 0.246 |

**Supplementary Table 52: Regions mediating the associations between environmental variables and changes on the Picture Vocabulary Task**

The table summarises the mediation analyses ran across regions that showed both environment-brain and brain-behaviour associations. The ACME (Average Causal Mediation Effect) estimate is reported, which reflects the indirect effect of the predictor on the outcome that is transmitted through the mediator, as well as proportion mediated values, which indicate the estimated fraction of the total effect explained by the mediator. Associated ACME confidence intervals and FDR-corrected p values are also reported. P values are false discovery rate (FDR) corrected to account for multiple comparisons. Analyses controlled for age and sex and included random intercepts for testing site and family ID (to account for sibling pairs).

| **Predictor** | **Region** | **ACME estimate** | **ACME CI** | **Proportion mediated** | ***p*_FDR_** |
| --- | --- | --- | --- | --- | --- |
| Education | Right Isthmus Cingulate Gyrus | 0.003 | -0.000 – -0.010 | 0.050 | 0.086 |
| General LF | Right Isthmus Cingulate Gyrus | 0.007 | -0.000 – -0.020 | 0.017 | 0.024 |

**Supplementary Table 53: Correlations between COI 2.0 and ADI analysis**

Reported are Pearson’s correlations (Pearson’s r) between the standardised beta coefficients of the analyses using the COI 2.0 and the ADI. Correlations are reported separately for baseline and longitudinal analyses, with asterisks indicating statistical significance (***p < 0.001).

| **Measure** | **Baseline analysis (Pearson’s r)** | **Longitudinal analysis (Pearson’s r)** |
| --- | --- | --- |
| COI 2.0 Social & Economic | 0.974^***^ | 0.865^***^ |
| COI 2.0 Education | 0.958^***^ | 0.764^***^ |

**Supplementary Table 54: Correlations between main analysis and puberty-controlled analysis**

Reported are Pearson’s correlations (Pearson’s r) between the standardised beta coefficients of the original analyses and the sensitivity analyses controlling for puberty. Correlations are reported separately for baseline and longitudinal analyses across environmental and behavioural measures, with asterisks indicating statistical significance (***p < 0.001).

| Measure | Baseline analysis (Pearson’s r) | Longitudinal analysis (Pearson’s r) |
| --- | --- | --- |
| COI 2.0 Health & Environment | 0.981^***^ | 0.917^***^ |
| COI 2.0 Social & Economic | 0.988^***^ | 0.891^***^ |
| COI 2.0 Education | 0.985^***^ | 0.930^***^ |
| Micro-environment | 0.987^***^ | 0.861^***^ |
| Externalising | 0.960^***^ | 0.947^***^ |
| Internalising | 0.918^***^ | 0.868^***^ |
| Flanker Inhibitory Control & Attention | 0.907^***^ | 0.862^***^ |
| Picture Sequence Memory | 0.956^***^ | 0.883^***^ |
| Pattern Comparison Processing Speed | 0.890^***^ | 0.903^***^ |
| Picture Vocabulary | 0.990^***^ | 0.903^***^ |
| Oral Reading Recognition | 0.987^***^ | 0.883^***^ |

**Supplementary Figure 1: ADI Baseline Figure**

Brain plots depicting baseline environment-brain associations using the ADI. Regions are colour-coded according to the magnitude and direction of the standardised beta values. Statistically significant regions (p_FDR_ < 0.05) are outlined.

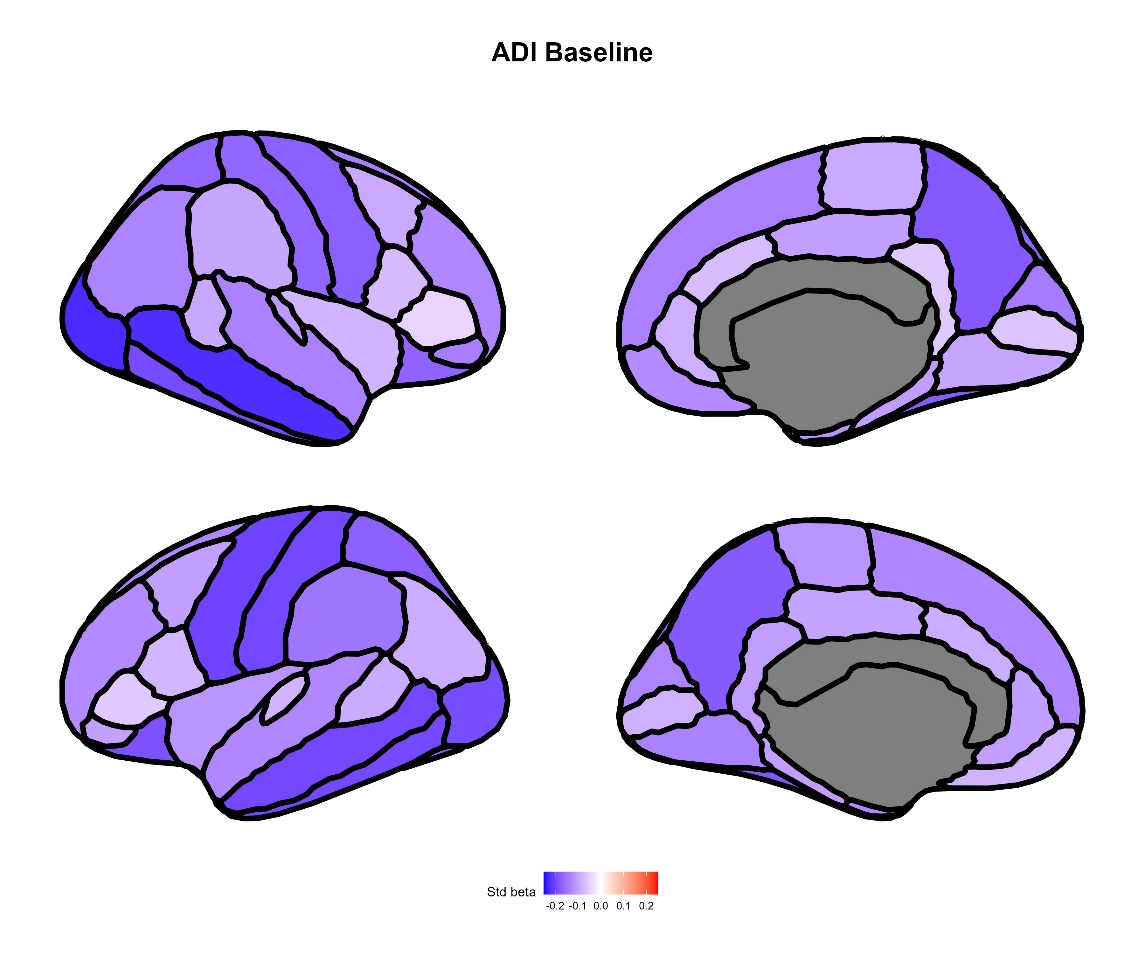

**Supplementary Figure 2: ADI Longitudinal Figure**

Brain plots depicting longitudinal environment-brain associations using the ADI. Regions are colour-coded according to the magnitude and direction of the standardised beta values. Statistically significant regions (p_FDR_ < 0.05) are outlined.

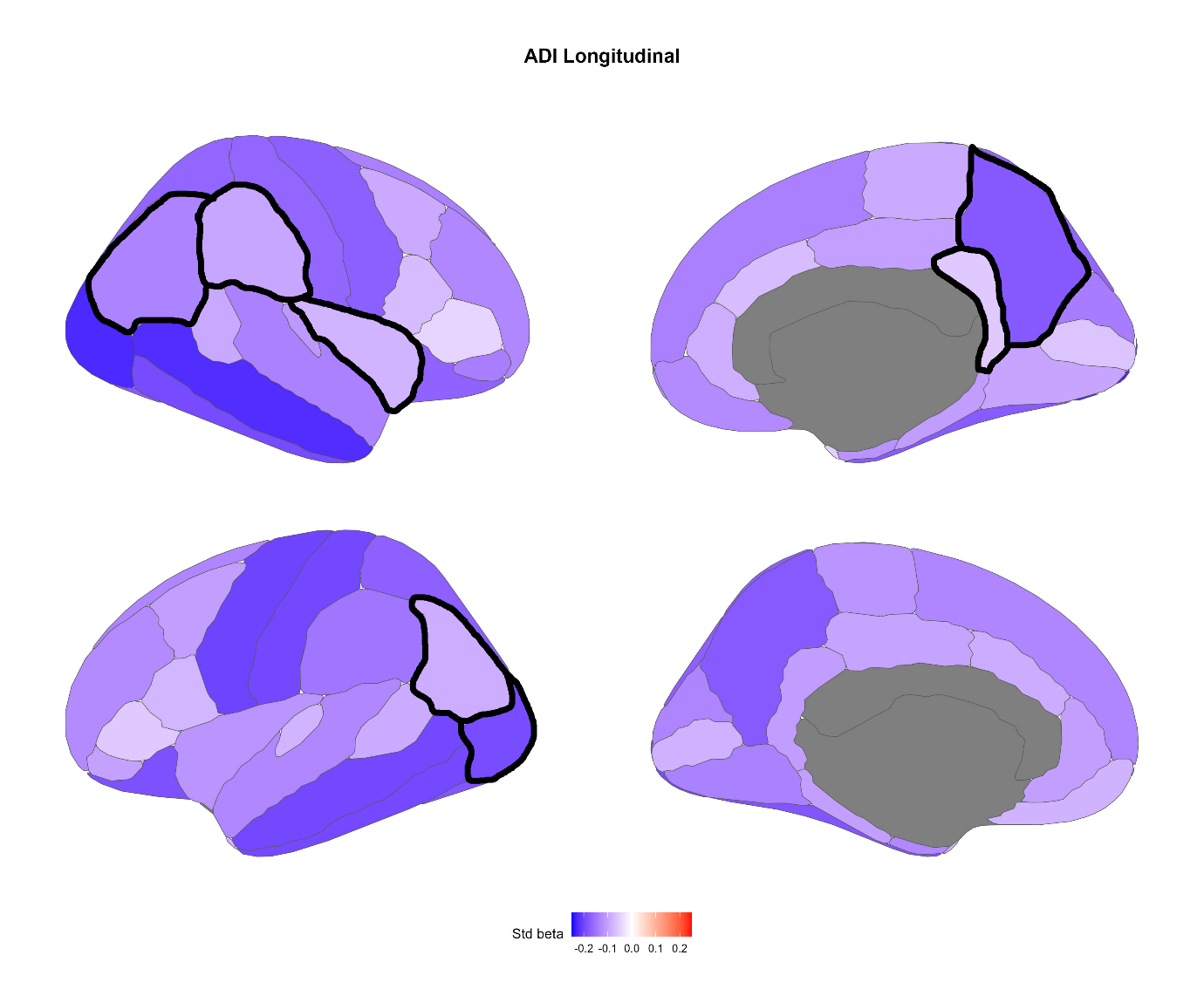

**Supplementary Figure 3: Puberty Baseline Figure**

Brain plots depicting regional standardised beta coefficients for environment-brain and brain-behaviour associations after controlling for puberty. Regions are colour-coded according to the magnitude and direction of the standardised beta values. Statistically significant regions (p_FDR_ < 0.05) are outlined.

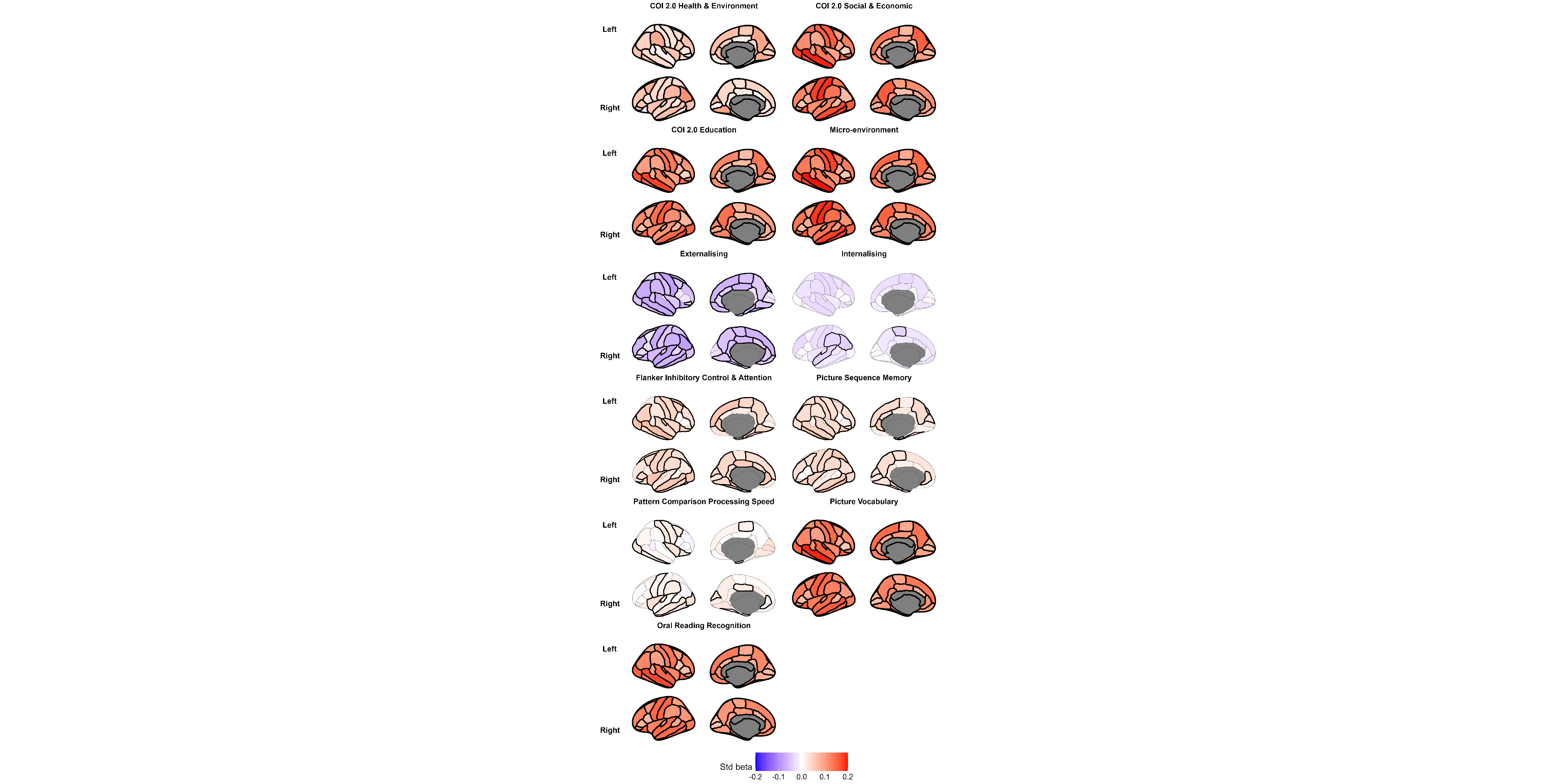

**Supplementary Figure 4: Puberty Longitudinal Figure**

Brain plots depicting the longitudinal environment-brain and brain-behaviour associations after controlling for the change in pubertal stage across the studied period. Regions are colour-coded according to the magnitude and direction of the standardised beta values. Statistically significant regions (p_FDR_ < 0.05) are outlined.

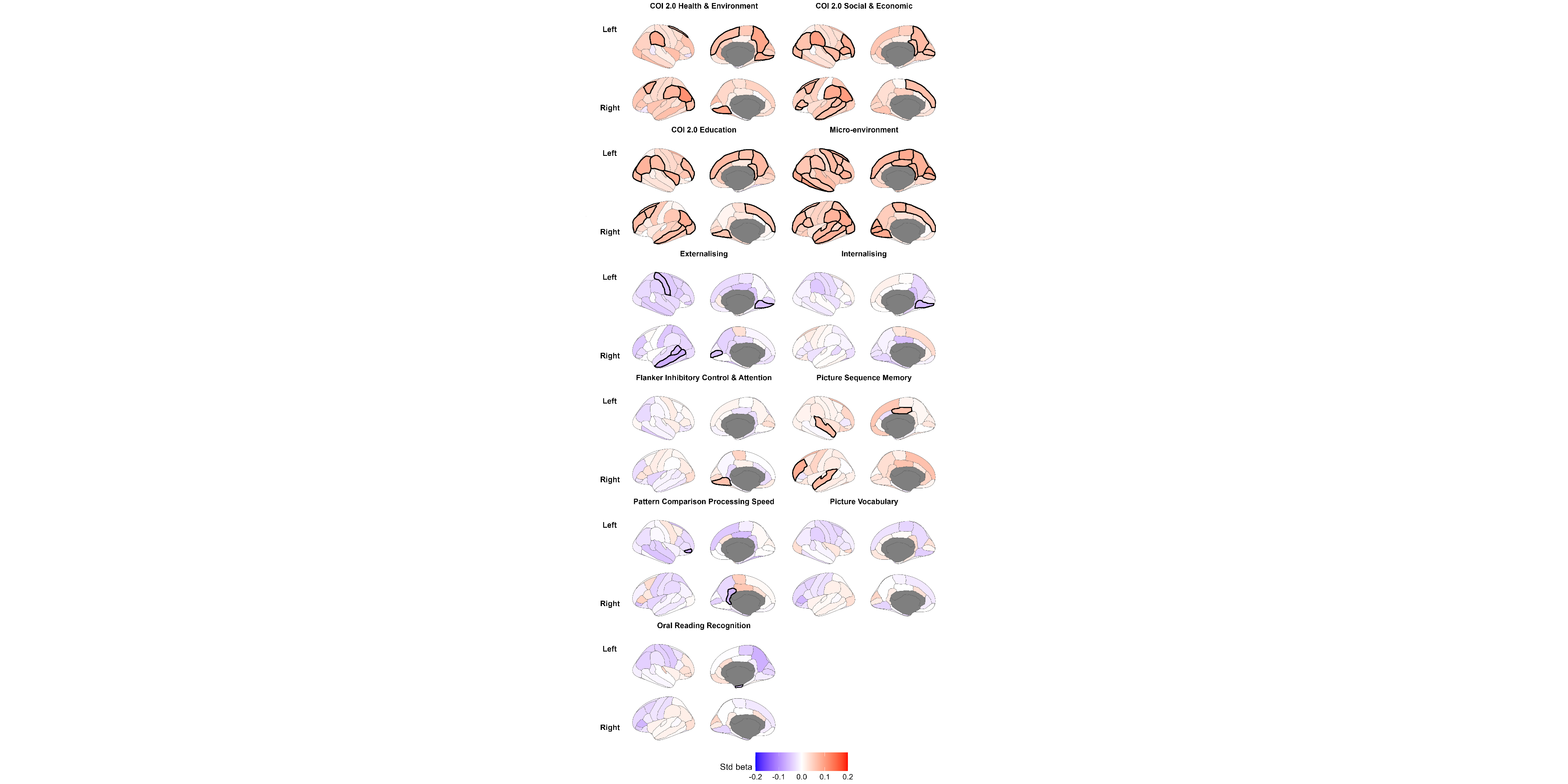
